# Identification and Functional Characterization of CXCL17 Orthologs in Amphibians

**DOI:** 10.64898/2026.01.18.700233

**Authors:** Jie Yu, Hao-Zheng Li, Juan-Juan Wang, Jun-Jie Yao, Wen-Feng Hu, Ya-Li Liu, Zhan-Yun Guo

**Affiliations:** Research Center for Translational Medicine at East Hospital, School of Life Sciences and Technology, Tongji University, Shanghai, China; National Institute of Metrology of China, Beijing, China

**Keywords:** Activity, Amphibian, CXCL17, GPR25, Receptor

## Abstract

C-X-C motif chemokine ligand 17 (CXCL17) has recently been identified as an agonist of the poorly characterized G protein-coupled receptor 25 (GPR25). Although GPR25 orthologs are widely distributed across vertebrates, non-mammalian CXCL17 orthologs have only been identified in some fish species in our recent studies. In this study, we systematically searched public databases for amphibian CXCL17 orthologs based on conserved C-terminal motif, gene synteny, and genomic architecture. Using this approach, we identified up to eighteen CXCL17 orthologs from diverse amphibian species. These amphibian CXCL17s exhibit no significant overall sequence similarity to known mammalian or fish CXCL17s, thus they were previously classified as uncharacterized proteins or even unannotated. Compared with known mammalian or fish CXCL17s, most amphibian CXCL17s display distinctive features, including four cysteine residues in their mature peptide and an additional residue following the conserved C-terminal Xaa-Pro-Yaa motif. A representative ortholog from the tropical clawed frog (*Xenopus tropicalis*) was recombinantly expressed and functionally characterized using cell-based assays, inducing ligand-receptor binding, β-arrestin recruitment, and chemotactic cell migration. The recombinant amphibian CXCL17 directly bound to and efficiently activated its cognate GPR25 receptor and induced chemotactic migration of the transfected human embryonic kidney (HEK) 293T cells, but deletion of four C-terminal residues largely abolished its activity, indicating that all CXCL17 orthologs employ a conserved mechanism for receptor binding and activation. These findings establish the presence of a functional CXCL17–GPR25 signaling system in amphibians and provide new insights into the phylogenetic distribution and sequence diversity of CXCL17 orthologs across vertebrate lineages.

## 1. Introduction

The mucosal C-X-C motif chemokine ligand 17 (CXCL17) was discovered approximately two decades ago [1,2]. It is involved in the recruitment of multiple immune cell types, including T cells, monocytes, macrophages, and dendritic cells [3-16], and has also been implicated in tumor development, likely through modulation of tumor-associated immune responses [17-22]. Despite these well-documented functions, the identity of the CXCL17 receptor has remained controversial. The early report proposing the orphan G protein-coupled receptor 35 (GPR35) as its receptor could not be independently reproduced [23-26]. More recently, CXCL17 has been suggested to act either as a modulator of the chemokine receptor CXCR4 [26] or as an agonist of the MAS-related receptor MRGPRX2 [27]. However, our recent work demonstrated that although human CXCL17 can activate MRGPRX2, as well as MRGPRX1 and MAS1, at micromolar concentrations, this activity does not depend on its conserved C-terminal fragment [28], indicating that these MAS-related receptors are unlikely to represent the evolutionarily conserved receptors for CXCL17.

Most recently, Ocón’s group and our team independently demonstrated that CXCL17 activates the poorly characterized G protein-coupled receptor 25 (GPR25) through its conserved C-terminal fragment [29,30], suggesting that CXCL17 and GPR25 constitute an evolutionarily conserved ligand–receptor pair. However, their phylogenetic distributions appear discordant: GPR25 orthologs are widely present across vertebrates, from fishes to mammals, whereas CXCL17 orthologs were initially thought to be restricted to mammals. In our recent studies, we searched for non-mammalian CXCL17 orthologs based on key structural and genomic features of mammalian CXCL17s and successfully identified fish CXCL17 orthologs in the lobe-finned coelacanth (*Latimeria chalumnae*) and in several ray-finned teleost fishes, including zebrafish (*Danio rerio*) [31-33]. These fish CXCL17s exhibit no significant overall amino acid sequence similarity to mammalian CXCL17s, rendering them undetectable by conventional homology-based approaches and leaving them annotated as uncharacterized proteins with unknown functions in public databases.

Amphibians represent an important vertebrate lineage, comprising approximately 8,000 extant species. Despite the presence of GPR25 orthologs in amphibians, CXCL17 orthologs have not previously been identified in this group. In this study, we searched public databases for putative amphibian CXCL17s based on the essential C-terminal Xaa-Pro-Yaa motif, as well as the gene synteny and genomic architecture of mammalian CXCL17s. Using this strategy, we identified up to 18 amphibian CXCL17 orthologs from frogs, toads, newts, and caecilians. Among them, four have been annotated in the National Center for Biotechnology Information (NCBI) databases but remain classified as uncharacterized proteins with unknown functions, whereas fourteen have not yet been annotated, although RNA sequencing data in the NCBI reference genomes support their existence. These amphibian CXCL17 orthologs exhibit no significant overall amino acid sequence similarity to known mammalian or fish CXCL17s and display several distinctive features, including four cysteine residues in the mature peptide and an additional residue following the conserved C-terminal Xaa-Pro-Yaa motif. A representative ortholog from tropical clawed frog (*Xenopus tropicalis*), designated as Xt-CXCL17, was recombinantly expressed in *Escherichia coli* and showed robust activity toward its cognate receptor, Xt-GPR25, in multiple cell-based functional assays, including NanoLuc Binary Technology (NanoBiT)-based β-arrestin recruitment, NanoBiT-based homogeneous ligand–receptor binding, and chemotaxis assays performed in transfected human embryonic kidney (HEK) 293T cells. In contrast, deletion of four C-terminal residues almost completely abolished activity in these assays, indicating that Xt-CXCL17 engages Xt-GPR25 through a mechanism conserved for all CXCL17-GPR25 pairs. These results demonstrate the existence of a functional CXCL17–GPR25 signaling system in amphibians and provide new insights into the phylogenetic distribution and sequence diversity of CXCL17 orthologs across vertebrate lineages.

## 2. Materials and methods

### 2.1. Recombinant expression of Xt-CXCL17

A DNA fragment encoding an N-terminally 6×His-tagged mature peptide of Xt-CXCL17 was chemically synthesized using *Escherichia coli*-biased codons and ligated into a pET vector via Gibson assembly, resulting in the expression construct pET/6×His-Xt-CXCL17 (Fig. S1). The coding region of the small NanoLuc fragment for NanoBiT (SmBiT)-fused Xt-CXCL17 was amplified by polymerase chain reaction (PCR) using pET/6×His-Xt-CXCL17 as the template and ligated into a pET vector via Gibson assembly, resulting in the expression construct pET/6×His-SmBiT-Xt-CXCL17 (Fig. S1). The expression construct for the C-terminally truncated mutant, 6×His-Xt-[desC4]CXCL17, was generated via the QuikChange approach using pET/6×His-Xt-CXCL17 as a mutagenesis template. The coding region of Xt-CXCL17 in these constructs was confirmed by DNA sequencing.

The recombinant wild-type (WT) or mutant Xt-CXCL17s were prepared according to our previous procedure developed for human or fish CXCL17s [30-33]. Briefly, the overexpressed Xt-CXCL17s were solubilized from inclusion bodies via an *S*-sulfonation approach, purified by immobilized metal ion affinity chromatography (Ni^2+^ column), refolded *in vitro*, and further purified by high performance liquid chromatography (HPLC) using C_18_ reverse-phase columns. After lyophilization, the purified Xt-CXCL17 proteins were dissolved in 1.0 mM aqueous hydrochloride (pH 3.0) and quantified by ultra-violet absorbance at 280 nm according to their calculated extinction coefficient (6990 M^-1^ cm^-1^ for WT or truncated 6×His-Xt-CXCL17; 8480 M^-1^ cm^-1^ for 6×His-SmBiT-Xt-CXCL17). During preparation, samples were analyzed by sodium dodecyl sulfate-polyacrylamide gel electrophoresis (SDS-PAGE). The refolded 6×His-Xt-CXCL17 was also subjected to mass spectroscopy analysis that was conducted on a Q Exactive mass spectrometer (ThermoFisher Scientific, Waltham, MA, USA).

### 2.2. Generation of the expression constructs for Xt-GPR25

The coding region of Xt-GPR25 was chemically synthesized according to its cDNA sequence (XM_002941165) and ligated into a pcDNA3.1 vector, resulting in the expression construct pcDNA3.1/Xt-GPR25 (Table S1 and Fig. S2). Thereafter, the coding region of Xt-GPR25 was PCR amplified using pcDNA3.1/Xt-GPR25 as the template and appropriate oligoes as primers and ligated into appropriate doxycycline (Dox)-inducible expression vectors for functional assays (Table S1 and Fig. S2). The construct pTRE3G-BI/Xt-GPR25-LgBiT:SmBiT-ARRB2 coexpresses a C-terminally large NanoLuc fragment for NanoBiT (LgBiT)-fused Xt-GPR25 (Xt-GPR25-LgBiT) and an N-terminally SmBiT-fused human β-arrestin 2 (SmBiT-ARRB2) for NanoBiT-based β-arrestin recruitment assays; the construct PB-TRE/sLgBiT-Xt-GPR25 encodes an N-terminally secretory LgBiT (sLgBiT)-fused Xt-GPR25 (sLgBiT-Xt-GPR25) for NanoBiT-based ligand-receptor binding assays; the construct PB-TRE/Xt-GPR25 encodes an untagged Xt-GPR25 for chemotaxis assays.

### 2.3. The NanoBiT-based β-arrestin recruitment assays

The NanoBiT-based β-arrestin recruitment assays for Xt-GPR25 were conducted on transfected living HEK293T cells according to our previous procedure [30-34]. Briefly, HEK293T cells were transiently cotransfected with the Dox-inducible expression construct pTRE3G-BI/Xt-GPR25-LgBiT:SmBiT-ARRB2 and the expression control vector pCMV-Tet3G (Clontech, Mountain View, CA, USA) using Lipo8000 transfection reagent (Beyotime Biotechnology, Shanghai, China). Next day, the transfected cells were trypsinized, suspended in induction medium (complete DMEM plus 1.0 ng/mL of Dox), seeded into white opaque 96-well plates, and cultured for ∼24 h to ∼90% confluence. To conduct the activation assay, the induction medium was removed, pre-warmed activation solution (serum-free DMEM plus 1% bovine serum albumin) was added (40 μL/well, containing 0.5 μL of NanoLuc substrate stock from Promega, Madison, WI, USA), and bioluminescence data were collected for ∼4 min on a SpectraMax iD3 plate reader (Molecular Devices, Sunnyvale, CA, USA). Thereafter, indicated concentrations of WT or mutant Xt-CXCL17 were added (10 μL/well), and bioluminescence data were continuously collected for ∼10 min. The inter-well variability of the measured bioluminescence was corrected by forcing curves after addition of NanoLuc substrate to same level and the corrected bioluminescence data were plotted using the SigmaPlot 10.0 software (SYSTAT software, Chicago, IL, USA). To obtain the dose-response curve, the measured bioluminescence data at highest point were plotted with the peptide concentrations using the SigmaPlot 10.0 software (SYSTAT software).

### 2.4. The NanoBiT-based ligand-receptor binding assays

The NanoBiT-based homogenous binding assays for Xt-GPR25 were conducted on transfected living HEK293T cells according to our previous procedures [32-36]. Briefly, HEK293T cells were transiently transfected with the Dox-inducible expression construct PB-TRE/sLgBiT-Xt-GPR25 with or without cotransfection of pTRE3G-BI/TPST1:TPST2, a Dox-inducible construct coexpressing human tyrosylprotein sulfotransferase TPST1 and TPST2. Next day, the transfected cells were trypsinized, suspended in induction medium (complete DMEM plus 20 ng/mL of Dox), seeded into white opaque 96-well plates, and cultured for ∼24 h to ∼90% confluence. To conduct the binding assay, the induction medium was removed and pre-warmed binding solution (serum-free DMEM plus 0.1% bovine serum albumin and 0.01% Tween-20) was added (50 μL/well). For saturation binding assays, the binding solution contains different concentrations of 6×His-SmBiT-Xt-CXCL17. For competition binding assays, the binding solution contains a constant concentration of 6×His-SmBiT-Xt-CXCL17 and different concentrations of competitors. After incubation at room temperature for ∼1 h, diluted NanoLuc substrate (30-fold dilution in the binding solution) was added (10 μL/well), and bioluminescence data were immediately measured on a SpectraMax iD3 plate reader (Molecular Devices). The measured bioluminescence data were expressed as mean ± standard deviation (SD, *n* = 3) and fitted to one-site binging model using the SigmaPlot 10.0 software (SYSTAT software).

### 2.5. Chemotaxis assays

The chemotaxis assays were conducted on transfected living HEK293T cells using a transwell apparatus according to our previous procedure [30-33]. Briefly, HEK293T cells were transiently transfected with the Dox-inducible expression construct PB-TRE/Xt-GPR25 and cultured the induction medium (complete DMEM plus 1.0 ng/mL of Dox) for ∼24 h. Thereafter, the cells were trypsinized, suspended in serum-free DMEM at the density of ∼5×10^5^ cells/mL, and seeded into polyethylene terephthalate membrane (8 μm pore size)-coated permeable transwell inserts that were pretreated with serum-free DMEM. The inserts were put into a 24-well plate containing chemotactic agent (WT or truncated Xt-CXCL17 diluted in serum-free DMEM plus 0.2% bovine serum albumin, 500 μL/well) and incubated at 37°C for ∼5 h. Thereafter, solution in the inserts was removed and cells on the upper face of the permeable membrane were wiped off using cotton swaps, and cells on the lower face of the permeable membrane were fixed with 4% paraformaldehyde solution, stained with crystal violet staining solution (Beyotime Technology), and observed under an Olympus APX100 microscope (Tokyo, Japan). The migrated cells were quantitated using the ImageJ software and the results were expressed as mean ± SD (*n* = 3).

## 3. Results

### 3.1. Identification of possible CXCL17 orthologs in amphibians

In mammals, *CXCL17* gene is typically located adjacent to the *CNFN*, *LIPE*, *CEACAM1*, and *CEACAM8* genes and share a conserved genomic organization consisting of four exons and three introns. We therefore hypothesized that amphibian *CXCL17* gene might display similar gene synteny and exon–intron architecture. Based on this premise, we searched the NCBI gene database for putative amphibian CXCL17 orthologs using the conserved C-terminal Xaa-Pro-Yaa motif (in which Xaa and Yaa are typically large aliphatic residues), together with conserved gene synteny and gene structure. Using this approach, we identified three annotated CXCL17 candidates from *Xenopus tropicalis*, *Ranitomeya variabilis*, and *Lithobates pipiens* that are present in the NCBI database but remain classified as uncharacterized proteins with unknown functions (Fig. 1A, Table S2, and Fig. S3–S5). In addition, we identified 14 putative amphibian CXCL17 orthologs that have not yet been annotated in the NCBI database, although RNA sequencing data from reference genomes support their existence (Fig. 1A, Table S2, and Fig. S6–S19). By manually assembling their four exons, we confirmed that these loci encode small secretory proteins containing a C-terminal Xaa-Pro-Yaa motif, consistent with the defining features of CXCL17s (Fig. 1A, Table S2, and Fig. S6–S19). Sequence BLAST analysis of these amphibian CXCL17s further retrieved a homolog from *Xenopus laevis* (Fig. 1A and Table S2). Thus, we identified up to 18 putative amphibian CXCL17 orthologs across frogs, toads, newts, and caecilians (Fig. 1A and Table S2).

**Fig. 1.**
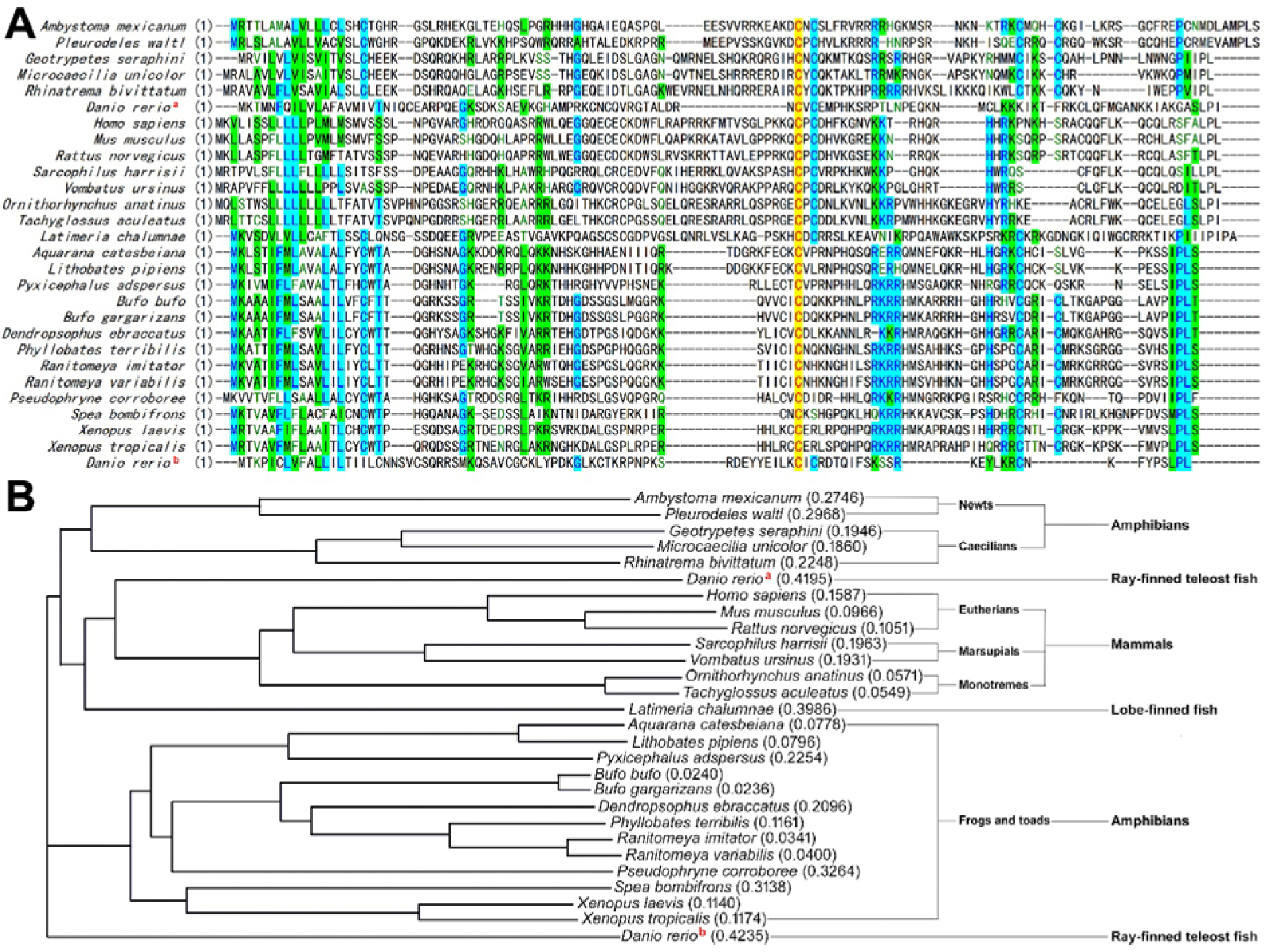
Identification of possible CXCL17 orthologs in amphibians. (**A**) Amino acid sequence alignment of CXCL17 orthologs from amphibians, mammals, or fishes. Their information is listed in supplementary Table S2 and S3. (**B**) Phylogenetic tree of the CXCL17 orthologs aligned in panel A. These sequences were aligned via AlignX algorithm using the Vector NTI 11.5.1 software. ^a^ zebrafish CXCL17, ^b^ zebrafish CXCL17-like.

All identified amphibian CXCL17s are small secretory proteins harboring an N-terminal signal peptide and a C-terminal Xaa-Pro-Yaa motif, which are hallmark features of mammalian and fish CXCL17s (Fig. 1A and Table S2). However, these amphibian orthologs show no significant overall amino acid sequence similarity to known mammalian or fish CXCL17s (Fig. 1A), explaining why they cannot be detected using conventional homology-based sequence searches. Moreover, substantial sequence divergence was observed among amphibian subgroups, including frogs/toads, newts, and caecilians (Fig. 1A,B). All amphibian CXCL17s are highly basic proteins, with predicted isoelectric point (pI) values ranging from 10.7 to 12.4 (Table S2). This property may be functionally relevant, as basic chemokines can bind negatively charged extracellular glycosaminoglycans after secretion and form localized concentration gradients to recruit immune cells. AlphaFold3 predictions indicated that mature amphibian CXCL17s adopt highly flexible conformations, with predicted pTM values of approximately 0.2. For ligand–receptor interaction modeling, AlphaFold3 yielded ipTM values below the reliability threshold of 0.6 in most cases, although the predicted models consistently suggested insertion of the CXCL17 C-terminal fragment into the orthosteric ligand-binding pocket of GPR25.

Compared with mammalian and fish CXCL17s, amphibian CXCL17s exhibit two notable features. First, most amphibian CXCL17s contain four cysteine residues in the mature peptide (Fig. 1A and Table S2), suggesting the formation of two intramolecular disulfide bonds, whereas mammalian and fish CXCL17s typically contain six cysteine residues and form three intramolecular disulfide bonds. Second, most amphibian CXCL17s possess an additional residue, commonly serine or threonine, immediately following the conserved Xaa-Pro-Yaa motif (Fig. 1A and Table S2). As the C-terminal Xaa-Pro-Yaa motif is predicted to insert into the orthosteric ligand-binding pocket of the GPR25 receptor, the functional impact of this additional C-terminal residue remains to be determined.

### 3.2. Recombinant preparation of Xt-CXCL17

To assess the biological activity of amphibian CXCL17s, the *X. tropicalis*–derived Xt-CXCL17 was selected as a representative, as it exhibits the distinctive features of amphibian CXCL17 orthologs. Following overexpression of the N-terminally 6×His-tagged Xt-CXCL17 in *Escherichia coli*, an approximately 12 kDa band (indicated by an asterisk) was detected by SDS-PAGE only after induction with isopropyl-β-D-thiogalactopyranoside (IPTG), confirming successful overexpression of the expected 6×His-Xt-CXCL17 (Fig. 2A). After bacterial lysis and solubilization of inclusion bodies, 6×His-Xt-CXCL17 was purified by immobilized metal ion affinity chromatography (Ni^2+^ column). SDS-PAGE analysis of the eluted fraction revealed the expected ∼12 kDa band along with several higher-molecular-weight species (Fig. 2A), indicating a tendency of Xt-CXCL17 toward intermolecular crosslinking, consistent with previous observations for recombinant human or fish CXCL17s [30,32,33].

**Fig. 2.**
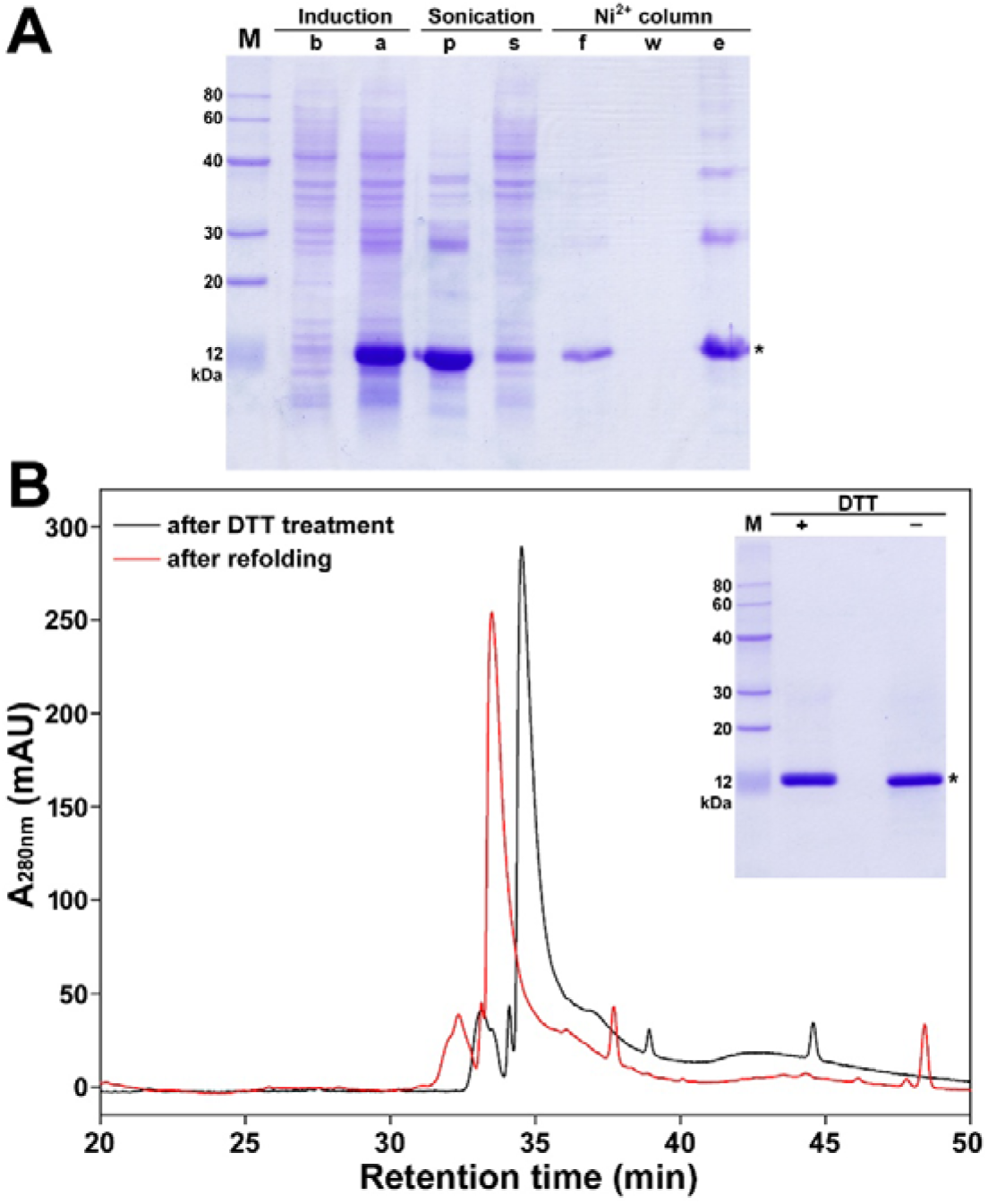
Preparation of recombinant Xt-CXCL17. (**A**) SDS-PAGE analysis of the recombinant 6×His-Xt-CXCL17. Samples at different preparation stages were loaded onto a 15% SDS-gel and subjected to electrophoresis. The gel was stained by Coomassie brilliant blue R250, and band of the monomeric 6×His-Xt-CXCL17 was indicated by an asterisk. Lane (M), protein ladder; lane (b), before IPTG induction; lane (a), after IPTG induction; lane (p), pellet after sonication; lane (s), supernatant after sonication; lane (f), flowthrough from the Ni^2+^ column; lane (w), washing fraction by 30 mM imidazole; lane (e), eluted fraction by 250 mM imidazole. (**B**) HPLC analysis of the recombinant 6×His-Xt-CXCL17. The eluted fraction from the Ni^2+^ column was analyzed by HPLC via a C_18_ reverse-phase column either after DTT treatment or after refolding. **Inner panel**, SDS-PAGE analysis of the refolded 6×His-Xt-CXCL17. The major refolding peak was collected and analyzed by SDS-PAGE. Lane (M), protein ladder; lane (+), with DTT treatment; lane (-), without DTT treatment.

The eluted fraction from the Ni^2+^ column was further analyzed by HPLC, which showed a sharp major peak eluting from a C_18_ reverse-phase column (Fig. 2B, black trace). After *in vitro* refolding, a dominant peak with a slightly shorter retention time was observed (Fig. 2B, red trace). This refolded peak was collected and analyzed by SDS-PAGE, revealing a single ∼12 kDa band under both reducing and non-reducing conditions (Fig. 2B, inner panel), suggesting that the refolded 6×His-Xt-CXCL17 was homogeneous. Mass spectrometric analysis showed that the refolded protein had a measured molecular mass of 11,547.1 Da, in agreement with its calculated average molecular mass of 11,547.1.

### 3.3. Activation of Xt-GPR25 by recombinant Xt-CXCL17

To determine whether recombinant 6×His-Xt-CXCL17 is functionally active, we employed a NanoBiT-based β-arrestin recruitment assay in which C-terminally LgBiT-fused Xt-GPR25 (Xt-GPR25-LgBiT) was coexpressed with N-terminally SmBiT-fused human β-arrestin 2 (SmBiT-ARRB2) in transfected HEK293T cells. Upon activation of Xt-GPR25-LgBiT by 6×His-Xt-CXCL17, SmBiT-ARRB2 is recruited to the receptor, restoring NanoLuc luciferase activity through proximity-induced complementation of SmBiT and LgBiT.

Addition of NanoLuc substrate to HEK293T cells coexpressing Xt-GPR25-LgBiT and SmBiT-ARRB2 produced only low basal bioluminescence, but subsequent addition of recombinant 6×His-Xt-CXCL17 resulted in a rapid, dose-dependent increase in bioluminescence (Fig. 3A), with a calculated EC_50_ value of approximately 120 nM. These results indicate that Xt-CXCL17 effectively activates Xt-GPR25 despite the presence of an additional serine residue at the C-terminus of the Xaa-Pro-Yaa motif. In contrast, removal of four C-terminal residues almost completely abolished activity in the β-arrestin recruitment assay, as the truncated 6×His-Xt-[desC4]CXCL17 exhibited only minimal effects even at concentrations up to 1.0 μM (Fig. 3B). These findings demonstrate that the C-terminal fragment is essential for Xt-CXCL17 activating Xt-GPR25, indicating that amphibian CXCL17s engage GPR25 through a mechanism conserved in all CXCL17-GPR25 pairs.

**Fig. 3.**
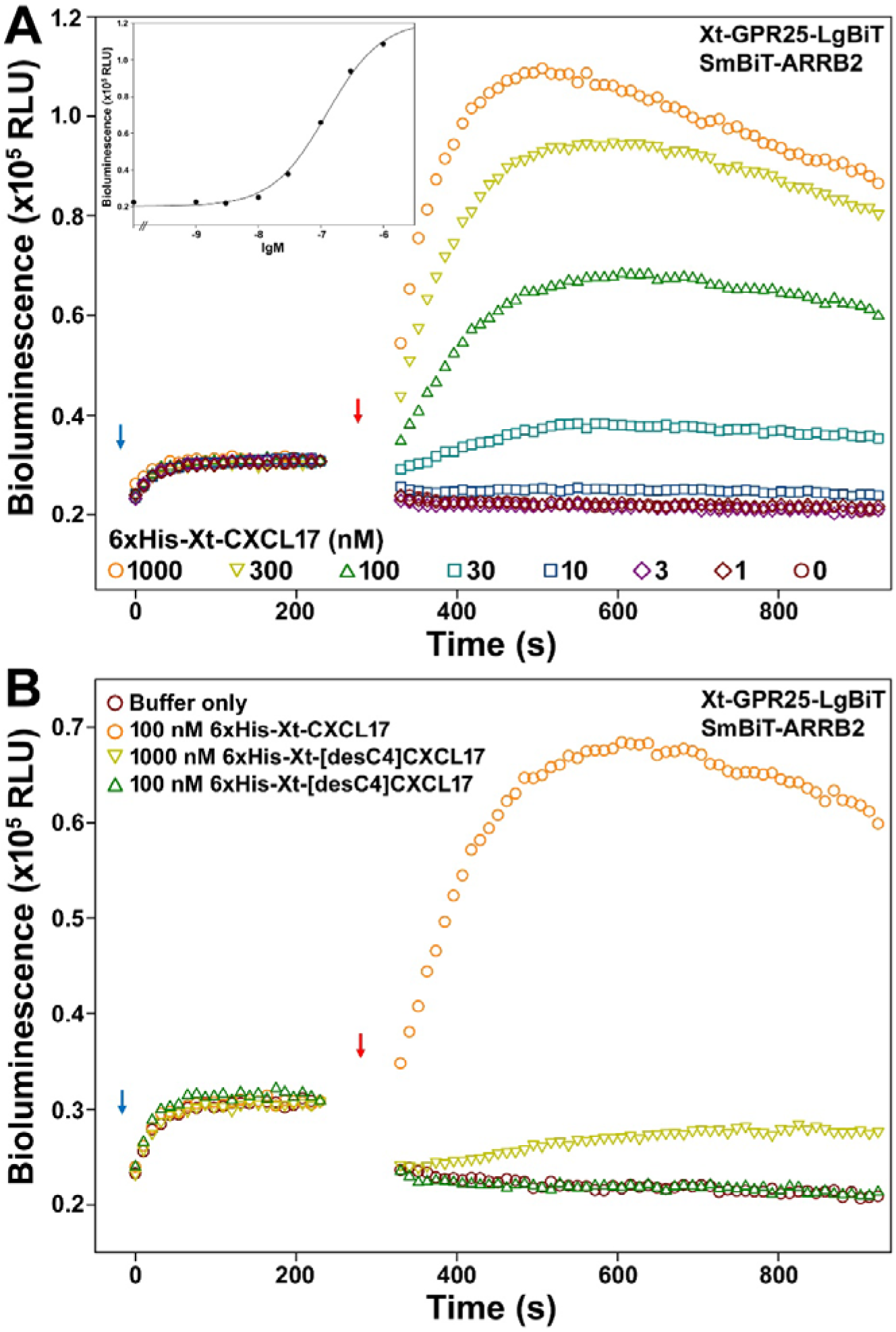
Activation of Xt-GPR25 by recombinant Xt-CXCL17 measured via NanoBiT-based β-arrestin recruitment assays. (**A**) Bioluminescence change caused by different concentrations of WT 6×His-Xt-CXCL17. **Inner panel**, dose response curve. (**B**) Bioluminescence change caused by the C-terminally truncated 6×His-Xt-[desC4]CXCL17. In these assays, NanoLuc substrate and recombinant WT or mutant Xt-CXCL17 were sequentially added to living HEK293T cells coexpressing Xt-GPR25-LgBiT and SmBiT-ARRB2, and bioluminescence was measured on a plate reader. The blue arrows indicate the addition of NanoLuc substrate, and red arrows indicate the addition of WT or mutant Xt-CXCL17.

### 3.4. Direct binding of recombinant Xt-CXCL17 to Xt-GPR25

To examine direct binding between Xt-CXCL17 and Xt-GPR25, we employed a NanoBiT-based binding assay that uses a SmBiT-tagged ligand (tracer) and an N-terminally sLgBiT-fused receptor. Upon ligand–receptor interaction, proximity-induced complementation of the ligand-attached SmBiT with the receptor-fused extracellular LgBiT restores luciferase activity.

When a 6×His tag and a SmBiT tag were tandemly fused to the N-terminus of Xt-CXCL17, the resulting tracer induced a rapid increase in bioluminescence in the NanoBiT-based β-arrestin recruitment assay (Fig. 4A), with a calculated EC_50_ value of approximately 130 nM, indicating that the N-terminal SmBiT tag did not compromise the activity of Xt-CXCL17.

**Fig. 4.**
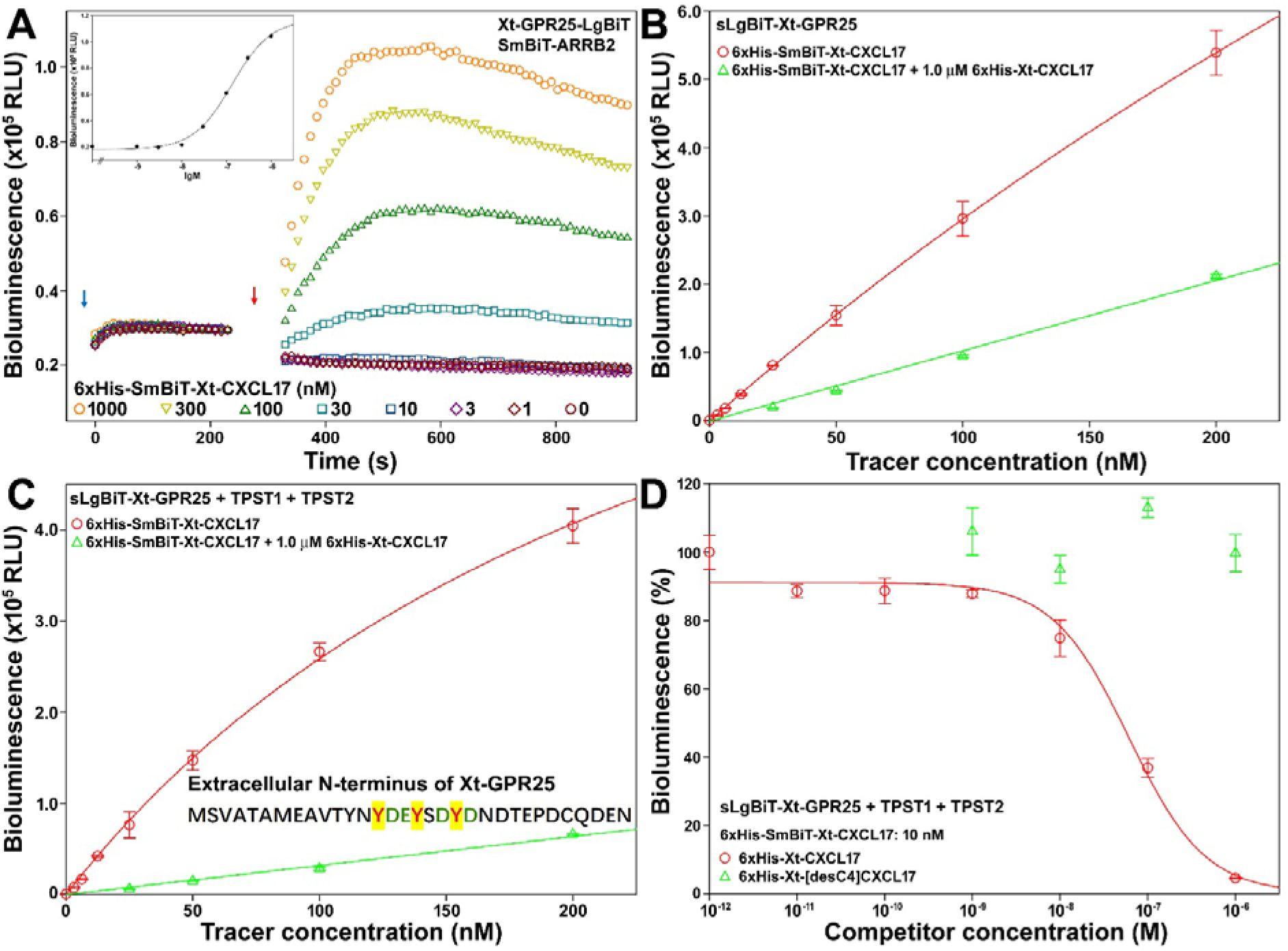
Direct binding of recombinant Xt-CXCL17 with Xt-GPR25 measured via the NanoBiT-based homogenous binding assays. (**A**) Activity of 6×His-SmBiT-Xt-CXCL17 towards Xt-GPR25 measured via the NanoBiT-based β-arrestin recruitment assay. The blue arrow indicates the addition of NanoLuc substrate, and the red arrow indicates the addition of 6×His-SmBiT-Xt-CXCL17. **Inner panel**, dose response curve. (**B,C**) Saturation binding of 6×His-SmBiT-Xt-CXCL17 with sLgBiT-Xt-GPR25 without (B) or with (C) coexpression of human tyrosylprotein sulfotransferases. The measured data are expressed as mean ± SD (*n* = 3) and plotted using the SigmaPlot10.0 software. Total binding data (red circles) were fitted with the function of Y = B_max_X/(K_d_+X) + k_non_X, the non-specific binding data (green triangles) were fitted with linear curves. (**D**) Competition binding of the WT or truncated Xt-CXCL17s with Xt-GPR25. The measured bioluminescence data are expressed as mean ± SD (*n* = 3) and fitted with sigmoidal curves using the SigmaPlot10.0 software.

Upon addition of 6×His-SmBiT-Xt-CXCL17 to living HEK293T cells overexpressing sLgBiT-Xt-GPR25, the bioluminescence signal increased in an almost linear manner (Fig. 4B). However, inclusion of 1.0 μM 6×His-Xt-CXCL17 as a competitor markedly reduced the signal (Fig. 4B), suggesting that the binding affinity of the tracer to sLgBiT-Xt-GPR25 was insufficient to reach saturation under these conditions. Notably, the extracellular N-terminus of Xt-GPR25 contains three tyrosine residues adjacent to negatively charged glutamate or aspartate residues (Fig. 4C, inner panel), implying potential post-translational sulfation. Consistent with this notion, coexpression of the tyrosylprotein sulfotransferases TPST1 and TPST2 resulted in a largely hyperbolic binding curve (Fig. 4C), yielding a calculated dissociation constant (K_d_) of approximately 270 nM, indicating that N-terminal tyrosine sulfation enhances binding between Xt-GPR25 and Xt-CXCL17.

In NanoBiT-based competition binding assays, 6×His-Xt-CXCL17 reduced the bioluminescence signal in a sigmoidal manner (Fig. 4D), with a calculated IC_50_ value of approximately 60 nM, consistent with its ability to bind sLgBiT-Xt-GPR25 and compete with the tracer. In contrast, the truncated 6×His-Xt-[desC4]CXCL17 failed to inhibit tracer binding (Fig. 4D), indicating minimal receptor binding, in agreement with its low activity in the β-arrestin recruitment assay.

### 3.5. Chemotactic movement of Xt-GPR25-expressing HEK293T cells induced by recombinant Xt-CXCL17

To determine whether Xt-CXCL17 induces cell migration through activation of Xt-GPR25, we performed chemotaxis assays using HEK293T cells transfected to express Xt-GPR25. In transwell assays, recombinant 6×His-Xt-CXCL17 induced migration of Xt-GPR25-expressing HEK293T cells in a dose-dependent manner (Fig. 5A,B), with significant chemotactic activity observed at concentrations as low as 10 nM. In contrast, the C-terminally truncated mutant exhibited a markedly reduced chemotactic effect, as concentrations up to 1,000 nM produced only minimal cell migration (Fig. 5A,B), consistent with its weak activity in the β-arrestin recruitment assay. These results demonstrate that recombinant Xt-CXCL17 promotes cell migration through activation of Xt-GPR25, providing functional evidence that the CXCL17–GPR25 signaling system is operative in amphibians.

**Fig. 5.**
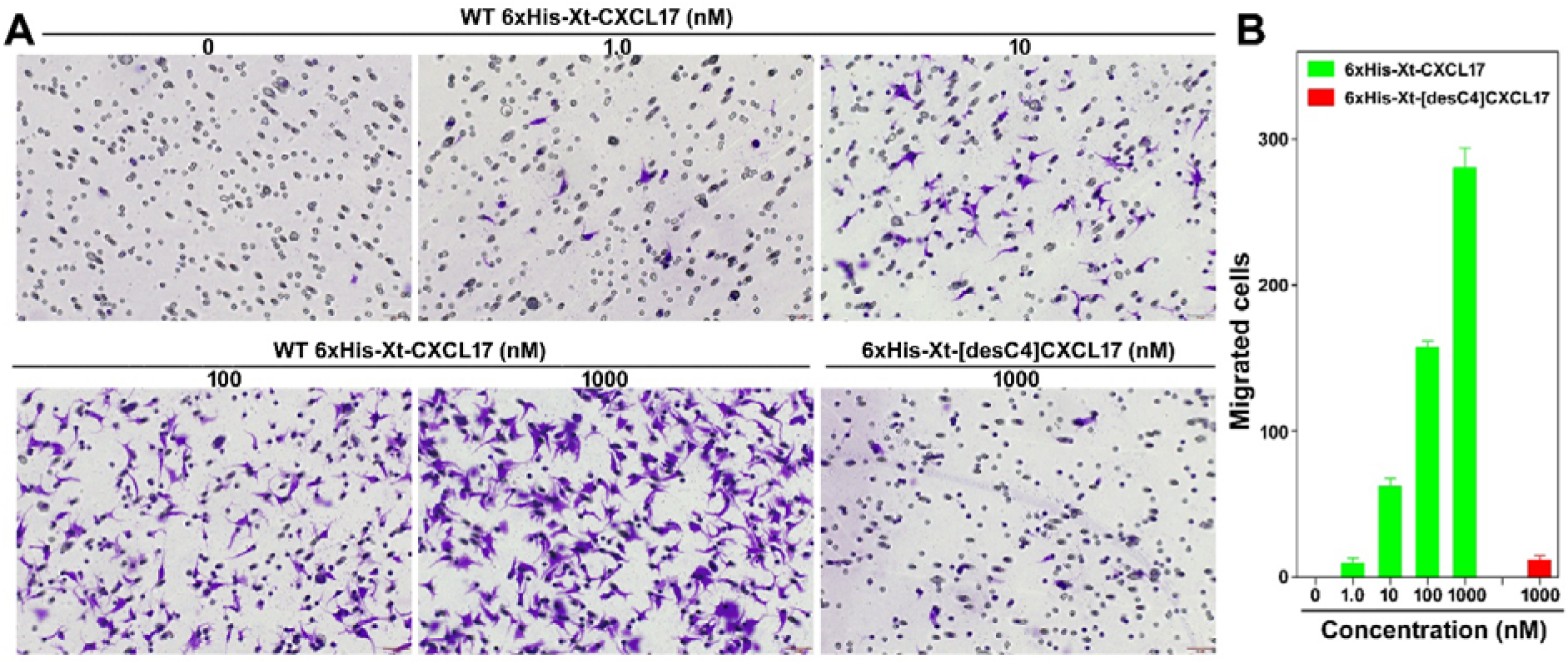
Chemotactic migration of Xt-GPR25-expressing HEK293T cells induced by recombinant Xt-CXCL17 measured via transwell assays. (**A**) Images of the migrated HEK293T cells induced by WT or mutant 6×His-Xt-CXCL17. The scale bar in these images is 50 μm. (**B**) Quantitative analysis of the migrated HEK293T cells induced by WT or mutant 6×His-Xt-CXCL17. Transfected HEK293T cells expressing Xt-GPR25 were seeded into the permeable membrane-coated inserts, and induced by chemotactic solution containing indicated concentrations of WT or mutant 6×His-Xt-CXCL17. After the transwell assay, cells on the upper face of the permeable membrane were wiped off, and cells on the lower face of the permeable membrane were fixed, stained, and observed under a microscope. The migrated cell numbers were calculated using the ImageJ software and the results are expressed as mean ± SD (*n* = 3).

## Discussion

In this study, we identified a group of CXCL17 orthologs in amphibians for the first time. These amphibian CXCL17s correspond either to previously uncharacterized proteins with unknown functions or to loci that remain entirely unannotated in public databases. Functional analyses demonstrated that a representative ortholog, Xt-CXCL17, is capable of binding to and activating its cognate receptor, Xt-GPR25, and of inducing chemotactic migration of transfected HEK293T cells, providing direct evidence for the existence of a functional CXCL17–GPR25 signaling system in amphibians.

Although the newly identified amphibian CXCL17s show no significant overall amino acid sequence similarity to known mammalian or fish CXCL17s, they retain several defining features of CXCL17s at both the protein and genomic levels. These include small secreted proteins with an N-terminal signal peptide and a typical length of fewer than 200 residues, a conserved C-terminal Xaa-Pro-Yaa motif in which Xaa and Yaa are generally large aliphatic residues, a high abundance of positively charged residues resulting in strongly basic proteins, conserved genomic organization relative to neighboring genes, and a shared exon–intron structure comprising four exons and three introns. On the basis of these conserved characteristics, additional CXCL17 orthologs are likely to be identified in other amphibian species and more broadly among non-mammalian vertebrates. Amphibian CXCL17s also exhibit distinctive features, including the presence of four cysteine residues in the mature peptide and an additional residue immediately following the C-terminal Xaa-Pro-Yaa motif. Importantly, these atypical properties do not appear to compromise receptor binding or biological activity, thereby refining our understanding of the essential structural determinants of CXCL17 function and providing a framework for the future identification of highly divergent CXCL17 orthologs.

CXCL17 orthologs from mammals, fishes, and amphibians exhibit exceptionally high sequence divergence (Fig. 1A), whereas their corresponding GPR25 orthologs display substantial sequence conservation (Fig. S20). This apparent mismatch raises the question of how such highly variable CXCL17 orthologs are able to recognize and activate their cognate GPR25 receptors. Our previous and present studies demonstrate that the conserved C-terminal Xaa-Pro-Yaa motif is indispensable for CXCL17 activity, as removal of this motif nearly abolished the activity of all representative CXCL17 orthologs examined to date [30-33]. However, a synthetic peptide corresponding to the short C-terminal fragment (10 residues) of human CXCL17 exhibited little to no activity (our unpublished data), indicating that the conserved C-terminal motif alone is insufficient for full receptor activation. These observations suggest that additional elements within the N-terminal region are required to achieve high-affinity binding and efficient receptor activation, although the critical determinants within this highly variable region remain to be identified.

To date, no experimentally determined structures of CXCL17 orthologs are available. AlphaFold predictions indicate that CXCL17 proteins adopt highly flexible conformations, limiting reliable structural modeling. Taken together with their extreme sequence variability, these findings raise the possibility that CXCL17 orthologs function as intrinsically disordered proteins lacking stable tertiary structures. Such structural plasticity may enable diverse CXCL17 sequences to engage a conserved receptor scaffold through a shared binding mechanism centered on the C-terminal motif. Nevertheless, this hypothesis remains speculative and will require experimental validation in future studies.

## CRediT authorship contribution statement

**Jie Yu**: Investigation, Methodology, Visualization. **Hao-Zheng Li**: Investigation, Methodology. **Juan-Juan Wang**: Investigation, Methodology. **Jun-Jie Yao**: Investigation. **Wen-Feng Hu**: Investigation. **Ya-Li Liu**: Formal analysis, Project administration. **Zhan-Yun Guo**: Supervision, Conceptualization, Writing - Original Draft, Writing - Review & Editing, Funding acquisition.

## Data availability statement

The data of this study are available in this manuscript, as well as the associated supplementary information.

## Declaration of competing interest

The authors declare no conflict of interest.

## Acknowledgments

This work was supported by grant from the National Natural Science Foundation of China (31971193).

## Supplementary data

Supplementary data for this article can be found online.

## Notes

### Competing Interest Statement

The authors have declared no competing interest.

