## Supplemental Table S1-S3 and Fig. S1-S20 for "Identification and Functional Characterization of CXCL17 Orthologs in Amphibians"

**Contents:**

**Table S1.** Information for generation of expression constructs for Xt-GPR25.

**Table S2.** Amphibian CXCL17 orthologs identified in this study.

**Table S3.** Information about other CXCL17 orthologs and GPR25 orthologs used in this study.

**Fig. S1.** The nucleotide sequence and amino acid sequence of the Xt-CXCL17 constructs overexpressed in *E. coli*.

**Fig. S2.** The nucleotide sequence and amino acid sequence of Xt-GPR25 constructs expressed in HEK293T cells.

**Fig. S3.** Gene position and architecture of the possible *cxcl17* gene (LOC116412198) from *Xenopus tropicalis* (tropical clawed frog).

**Fig. S4.** Gene position and architecture of the possible *cxcl17* gene (LOC143770339) from *Ranitomeya variabilis* (Zimmermann's poison frog).

**Fig. S5.** Gene position and architecture of the possible *cxcl17* gene (LOC143934019) from *Lithobates pipiens* (northern leopard frog).

**Fig. S6.** Gene position and architecture (**A**), genomic DNA sequence (**B**), cDNA sequence (**C**), and encoded protein sequence (**D**) of the unannotated *cxcl17* gene in the NCBI reference genome of *Pseudophryne corroboree* (corroboree frog).

**Fig. S7.** Gene position and architecture (**A**), genomic DNA sequence (**B**), cDNA sequence (**C**), and encoded protein sequence (**D**) of the unannotated *cxcl17* gene in the NCBI reference genome of *Ranitomeya imitator* (mimic poison frog).

**Fig. S8.** Gene position and architecture (**A**), genomic DNA sequence (**B**), cDNA sequence (**C**), and encoded protein sequence (**D**) of the unannotated *cxcl17* gene in the NCBI reference genome of *Dendropsophus ebraccatus* (hourglass treefrog).

**Fig. S9.** Gene position and architecture (**A**), genomic DNA sequence (**B**), cDNA sequence (**C**), and encoded protein sequence (**D**) of the unannotated *cxcl17* gene in the NCBI reference genome of *Aquarana catesbeiana* (American bullfrog).

**Fig. S10.** Gene position and architecture (**A**), genomic DNA sequence (**B**), cDNA sequence (**C**), and encoded protein sequence (**D**) of the unannotated *cxcl17* gene in the NCBI reference genome of *Phyllobates terribilis*.

**Fig. S11.** Gene position and architecture (**A**), genomic DNA sequence (**B**), cDNA sequence (**C**), and encoded protein sequence (**D**) of the unannotated *cxcl17* gene in the NCBI reference genome of *Pyxicephalus adspersus*.

**Fig. S12.** Gene position and architecture (**A**), genomic DNA sequence (**B**), cDNA sequence (**C**), and encoded protein sequence (**D**) of the unannotated *cxcl17* gene in the NCBI reference genome of *Bufo bufo* (common toad).

**Fig. S13.** Gene position and architecture (**A**), genomic DNA sequence (**B**), cDNA sequence (**C**), and encoded protein sequence (**D**) of the unannotated *cxcl17* gene in the NCBI reference genome of *Bufo gargarizans* (Asiatic toad).

**Fig. S14.** Gene position and architecture (**A**), genomic DNA sequence (**B**), cDNA sequence (**C**), and encoded protein sequence (**D**) of the unannotated *cxcl17* gene in the NCBI reference genome of *Spea bombifrons* (plains spadefoot toad).

**Fig. S15.** Gene position and architecture (**A**), genomic DNA sequence (**B**), cDNA sequence (**C**), and encoded protein sequence (**D**) of the unannotated *cxcl17* gene in the NCBI reference genome of *Pleurodeles waltl* (Iberian ribbed newt).

**Fig. S16.** Gene position and architecture (**A**), genomic DNA sequence (**B**), cDNA sequence (**C**), and encoded protein sequence (**D**) of the unannotated *cxcl17* gene in the NCBI reference genome of *Ambystoma mexicanum* (axolotl).

**Fig. S17.** Gene position and architecture (**A**), genomic DNA sequence (**B**), cDNA sequence (**C**), and encoded protein sequence (**D**) of the unannotated *cxcl17* gene in the NCBI reference genome of *Microcaecilia unicolor*.

**Fig. S18.** Gene position and architecture (**A**), genomic DNA sequence (**B**), cDNA sequence (**C**), and encoded protein sequence (**D**) of the unannotated *cxcl17* gene in the NCBI reference genome of *Rhinatrema bivittatum* (two-lined caecilian).

**Fig. S19.** Gene position and architecture (**A**), genomic DNA sequence (**B**), cDNA sequence (**C**), and encoded protein sequence (**D**) of the unannotated *cxcl17* gene in the NCBI reference genome of *Geotrypetes seraphini* (Gabon caecilian).

**Fig. S20.** GPR25 orthologs from mammals, fishes, and amphibians.

**Table S1.** Information for generation of expression constructs for Xt-GPR25.

| Expression constructs | Vectors for cloning | Restriction enzymes cleaving the vector | Primers for PCR amplification (5' to 3') | Template for PCR amplification | Approach for construct generation |
|---|---|---|---|---|---|
| pcDNA3.1/Xt-GPR25 | pcDNA3.1(+) | NheI, NotI | Chemical synthesis of the coding region of Xt-GPR25 (XM_002941165) | | Gibson assembly |
| pTRE3G-BI/ Xt-GPR25-LgBiT: SmBiT-ARRB2 | pTRE3G-BI/ GPR182-LgBiT: SmBiT-ARRB2 | NheI; AgeI (removing GPR182) | Forward: CCG TCA GAT CGC CTG GAG AAT TCG GGG AGA CCC AAG CTG GCT AGC<br>Reverse: GCT AGA CCC TCC GCC GGT ACC GAC CGG TGG GTA GCT GCG TGT GTA GGA GC | pcDNA3.1/Xt-GPR25 | Gibson assembly |
| PB-TRE/sLgBiT-Xt-GPR25 | PB-TRE/ sLgBiT-GPR182 | KpnI; PmeI (removing GPR182) | Forward: Forward: GGT GGC AGC GGC GGT GGT ACC AGT GTT GCT ACA GCC ATG GA<br>Reverse: GAG GCT GAT CAG CGG GTT TCA GTA GCT GCG TGT GTA GGA | pcDNA3.1/Xt-GPR25 | Gibson assembly |
| PB-TRE/Xt-GPR25 | PB-TRE/ dCas9-VPR | NheI; PmeI (removing dCas9-VPR) | Forward: TTC CTA CCC TCG TAA AGG TCT AGA G CTC ACT ATA GGG AGA CCC AAG CT<br>Reverse: GTT TCA GTT AGC CTC CCC CGT TT C ACA GTC GAG GCT GAT CAG CGG | pcDNA3.1/Xt-GPR25 | Gibson assembly |

PB-TRE/sLgBiT-GPR182 was generated in our laboratory based on PB-TRE/dCas9-VPR (Addgene cat#: 63800) via removal of the dCas9-VPR fragment by NheI and PmeI cleavage and then ligation of sLgBiT-GPR182 fragment, unpublished data.

pTRE3G-BI/GPR182-LgBiT:SmBiT-ARRB2 was generated in our laboratory, unpublished data.

For oligo primers, the sequence pairing with vector in Gibson assembly is highlighted in yellow, the sequence pairing with PCR template is underlined.

**Table S2.** Amphibian CXCL17 orthologs identified in this study. Their information is downloaded from the NCBI gene database (https://www.ncbi.nlm.nih.gov/gene). Those unannotated CXCL17 orthologs are manually generated according to RNA sequencing data in the NCBI gene database. The predicted signal peptide is shaded, cysteine residues in mature peptide are highlighted in yellow, the C-terminal Xaa-Pro-Yaa motif is shown in red and highlighted in yellow.

| Amphibian species | Information from NCBI | Amino acid sequence (signal peptide shaded) | pI of mature peptide |
|---|---|---|---|
| *Xenopus tropicalis*<br>(tropical clawed frog) | Gene ID: 116412198<br>mRNA ID: XM_031905867<br>Protein ID: XP_031761727<br>(Fig. S3) | MRTVAVFMFLAAITLCYCWTPQRQDSSGRTNENRGLAKRNGHKDALGSPLRPERHHLRCCERLSPQHPQRKR<br>RHMRAPRAHPIHQRRRCTTNCRGKKPSKFMVPLPLS | 12.4 |
| *Ranitomeya variabilis*<br>(Zimmermann's poison frog) | Gene ID: 143770339<br>mRNA ID: XM_077259867<br>Protein ID: XP_077115982<br>(Fig. S4) | MKVATIFMLSAVLILIYCLTTQGRHIPEKRHGKSGIARWSEHGESPGSPQGGKKTIICICNHKNGHIFSRKR<br>RHMSVHHKNGHHSPGCARICMRKGRRGGSVRSIPLS | 12.1 |
| *Lithobates pipiens*<br>(northern leopard frog) | Gene ID: 143934019<br>mRNA ID: XM_077457253<br>Protein ID: XP_077313379<br>(Fig. S5) | MKLSTIFMLAVALALFYCWTADGHSNAGKRENRRPLQKKNHHKGHHPDNITIQRKDDGKKFECKCVLRNPHQ<br>SQRERHQMNELQKRHLHGRKCHCKSLVKKPESSIPLS | 10.9 |
| *Pseudophryne corroboree*<br>(corroboree frog) | Unannotated<br>(Fig. S6) | MKVVTVFLLSAALLALCYCWTAHGHKSAGTRDDSRGLTKRIHHRDSLGSVQPGRQHALCVCDIDRHHLQRKK<br>RHMNGRRKPGIRSRHCCRRHFKQNTQPDVIIPLF | 12.0 |
| *Ranitomeya imitator*<br>(mimic poison frog) | Unannotated<br>(Fig. S7) | MKVATIFMLSAVLILIYCLTTQGHHIPEKRHGKSGVARWTQHGESPGSLQGRKKTIICICNHKNGHILSRKR<br>RHMSAHHKNGHHSPGCARICMRKGRRGGSVRSIPLS | 12.2 |
| *Dendropsophus ebraccatus*<br>(hourglass treefrog) | Unannotated<br>(Fig. S8) | MKAATIFLFSVVLILCYCWTTQGHYSAGKSHGKFIVARRTEHGDTPGSIQDGKKYLICVCDLKKANNLRKKR<br>HMRAQGKHGHHGRRCARICMQKGAHRGSQVSIPLT | 11.1 |
| *Aquarana catesbeiana*<br>(American bullfrog) | Unannotated<br>(Fig. S9) | MKLSTIFMLAVALALFYCWTADGHSNAGKKDDKRQLQKKNHSKGHHAENIIIQRTDGRKFECKCVPRNPHQS<br>QRERRQMNEFQKRHLHGRKCHCISLVGKPKSSIPLS | 11.1 |
| *Phyllobates terribilis* | Unannotated<br>(Fig. S10) | MKATTIFMLSAVLILFYCLTTQGRHNSGTWHGKSGVARRIEHGDSPGPHQGGRKSVICICNQKNGHNLSRKR<br>RHMSAHHKSGPHSPGCARICMRKSGRGGSVHSIPLS | 12.2 |
| *Pyxicephalus adspersus* | Unannotated<br>(Fig. S11) | MKIVMIFLFAVALTLFHCWTADGHNHTGKRGLQRKTHHRGHYVVPHSNEKRLLECTCVPRNPHHLQRKRRHM<br>SGAQKRNHRGRRCQCKQSKRNSELSIPLS | 12.2 |
| *Bufo bufo*<br>(common toad) | Unannotated<br>(Fig. S12) | MKAAAIFMLSAALILVFCFTTQGRKSSGRTSSIVKRTDHGDSSGSLMGGRKQVVCICDQKKPHNLPRKRRHM<br>KARRRHGHHRHVCGRICLTKGAPGGLAVPIPLT | 12.2 |
| *Bufo gargarizans*<br>(Asiatic toad) | Unannotated<br>(Fig. S13) | MKAAAIFMLSAALILLFCFTTQGRKSSGRTSSIVKRTDHGDSSGSLPGGRKHVVCICDQKKPHNLPRKRRHM<br>KARRRHGHHRSVCDRICLTKGAPGGLAVPIPLT | 12.0 |
| *Spea bombifrons*<br>(plains spadefoot toad) | Unannotated<br>(Fig. S14) | MKTVAVFLFLACFAICNCWTPHGQANAGKSEDSSLAIKNTNIDARGYERKIIRCNCKSHGPQKLHQKRRHKK<br>AVCSKPSHDHRCRHICNRIRLKHGNPFDVSMPLS | 10.7 |
| *Pleurodeles waltl*<br>(Iberian ribbed newt) | Unannotated<br>(Fig. S15) | MRLSLALAVLLVACVSLCWGHRGPQKDEKRLVKKHPSQWRQRRAHTALEDKRPRRMEEPVSSKGVKDCPCHV<br>LKRRRRHNRPSRNKHISQECRRQCRGQWKSRGCQHEPCRMEVAMPLS | 11.6 |
| *Ambystoma mexicanum*<br>(axolotl) | Unannotated<br>(Fig. S16) | MRTTLAMALVLLLCLSHCTGHRGSLRHEKGLTEHQSLPGRHHHGHGAIEQASPGLEESVVRRKEAKDCNCSL<br>FRVRRRRHGKMSRNKNKTRKCMQHCKGILKRSGCFREPCNMDLAMPLS | 11.1 |
| *Microcaecilia unicolor* | Unannotated<br>(Fig. S17) | MRALAVLVLVISAITVSLCHEEKDSQRQQHGLAGRPSEVSSTHGEQKIDSLGAGNQVTNELSHRRRERDIRC<br>YCQKTAKLTRRMKRNGKAPSKYNQMKCIKKCHRVKWKQPMIPL | 10.7 |
| *Rhinatrema bivittatum*<br>(two-lined caecilian) | Unannotated<br>(Fig. S18) | MRAVAVLFLVSAVIALSLCLEEKDSHRQAQELAGKHSEFLRRPGEQEIDTLGAGKWEVRNELNHQRRERAIR<br>CYCQKTPKHPRRRRRHVKSLIKKKQIKWLCTKKCQKYNIWEPPVIPL | 10.7 |
| *Geotrypetes seraphini*<br>(Gabon caecilian) | Unannotated<br>(Fig. S19) | MRVILVLVISVITVSLCHEEKDSQRQKHRLARRPLKVSSTHGQLEIDSLGAGNQMRNELSHQKRQRGIHCNC<br>QKMTKQSRRSRRHGRVAPKYRHMMCIKSCQAHLPNNLNWNGPIIPL | 11.6 |
| *Xenopus laevis*<br>(African clawed frog) | Genbank ID: OCT73022<br>Protein name: XELAEV_18036001mg | MRTVAAFIFLAAITLCHCWTPESQDSAGRTDEDRSLPKRSVRKDALGSPNRPERHHLKCCERLRPQHPQRKR<br>RHMKAPRAQSIHHRRRCNTLCRGKKPPKVMVSLPLS | 12.0 |

**Table S3.** Information about other CXCL17 orthologs and GPR25 orthologs used in this study. Their information is downloaded from the NCBI gene database (https://www.ncbi.nlm.nih.gov/gene). The predicted signal peptide of CXCL17s is shaded. [a] zebrafish CXCL17, [b] zebrafish CXCL17-like.

| Class | Species | Gene ID | mRNA ID | Protein ID | Amino acid sequence |
|---|---|---|---|---|---|
| CXCL17s | *Homo sapiens* | 284340 | NM_198477 | NP_940879 | MKVLISSLLLLLPLMLMSMVSSSLNPGVARGHRDRGQASRRWLQEGGQECECKDWFLRAPRRKFMTVSGLPKKQCPCDHFKGNVKKTRHQRHHRKPNKHSRACQQFLKQCQLRSFALPL |
| | *Mus musculus* | 232983 | NM_153576 | NP_705804 | MKLLASPFLLLLPVMLMSMVFSSPNPGVARSHGDQHLAPRRWLLEGGQECECKDWFLQAPKRKATAVLGPPRKQCPCDHVKGREKKNRHQKHHRKSQRPSRACQQFLKRCHLASFALPL |
| | *Rattus norvegicus* | 308436 | NM_001107491 | NP_001100961 | MKLLASPFLLLLTGMFTATVSSSPNQEVARHHGDQHQAPRRWLWEGGQECDCKDWSLRVSKRKTTAVLEPPRKQCPCDHVKGSEKKNRRQKHHRKSQRPSRTCQQFLKRCQLASFTLPL |
| | *Sarcophilus harrisii* | 116422895 | XM_031963247 | XP_031819107 | MRTPVLSFLLLFLLLLLSITSFSSDPEAAGGQRHHKLHAWRHPQGRRQLCRCEDVFQKIHERRKLQVAKSPASHCPCVRPKHKWKKPGHQKHWRQSCFQFLKQCQLQSILLPL |
| | *Vombatus ursinus* | 114040160 | XM_027858170 | XP_027713971 | MRAPVFFLLLLLLLPPLSVASSSPNPEDAEGQRNHKLPAKRHARGGRQVCRCQDVFQNIHGGKRVQRAKPPARQCPCDRLKYKQKKPGLGHRTHWRRSCLGFLKQCQLRDITLPL |
| | *Tachyglossus aculeatus* | 119946135 | XM_038767563 | XP_038623491 | MRLTTCSLLLLLLLLLTFATVTSVPQNPGDRRSHGERRLAARRRLGELTHKCRCPGSSQELQRESRARRLQSPRGECPCDDLKVNLKKRPMWHHKGKEGRVHYRRKEACRLFWKQCELEGLSLPI |
| | *Ornithorhynchus anatinus* | 103166888 | XM_029066640 | XP_028922473 | MQLSTWSLLLLLLLLLTFATVTSVPHNPGGSRSHGERRQEARRRLGQITHKCRCPGLSQELQRESRARRLQSPRGECPCDNLKVNLKKRPVWHHKGKEGRVHYRHKEACRLFWKQCELEGLSLPI |
| | *Danio rerio*[a] | 100151367 | NM_001144821 | NP_001138293 | MKTMNFQILVLAFAVMIVTNIQCEARPQEGKSDKSAEVKGHAMPRKCNCQVRGTALDRNCVCEMPHKSRPTLNPEQKNMCLKKKIKTFRKCLQFMGANKKIAKGASLPI |
| | *Danio rerio*[b] | 100536854 | NM_001386806<br>XM_073906074 | NP_001373735<br>XP_073762175 | MTKPICLVFALLILTIILCNNSVCSQRRSMKQSAVCGCKLYPDKGLKCTKRPNPKSRDEYYEILKCICRDTQIFSKSSRKEYLKRCNKFYPSLPL |
| | *Latimeria chalumnae* | 106704074 | XM_014490303 | XP_014345789 | MKVSDVLVLLCAFTLSSCLQNSGSSDQEEGRVPEEASTVGAVKPQAGSCSCGDPVGSLQNRLVSLKAGPSKHCDCRRSLKEAVNIKRPQAWAWKSKPSRKRCKRKGDNGKIQIWGCRRKTIKPIIIPIPA |
| GPR25s | *Homo sapiens* | 2848 | NM_005298 | NP_005289 | MAPTEPWSPSPGSAPWDYSGLDGLEELELCPAGDLPYGYVYIPALYLAAFAVGLLGNAFVVWLLAGRRGPRRLVDTFVLHLAAADLGFVLTLPLWAAAAALGGRWPFGDGLCKLSSFALAGTRCAGALLLAGMSVDRYLAVVKLLEARPLRTPRCALASCCGVWAVALLAGLPSLVYRGLQPLPGGQDSQCGEEPSHAFQGLSLLLLLLTFVLPLVVTLFCYCRISRRLRRPPHVGRARRNSLRIIFAIESTFVGSWLPFSALRAVFHLARLGALPLPCPLLLALRWGLTIATCLAFVNSCANPLIYLLLDRSFRARALDGACGRTGRLARRISSASSLSRDDSSVFRCRAQAANTASASW |
| | *Mus musculus* | 383563 | NM_001101516 | NP_001094986 | MQSTEPWSPSWGTLSWDYSGSGSLDQVELCPAWNLPYGHAIIPALYLAAFAVGLPGNAFVVWLLSRQRGPRRLVDTFVLHLAAADLGFVLTLPLWAAAEARGGLWPFGDGLCKVSSFALAVTRCAGALLLAGMSVDRYLAVGRPLSARPLRSARCVRAVCGAAWAAAFLAGLPALLYRGLQPSLDGVGSQCAEEPWEALQGVGLLLLLLTFALPLAVTLICYWRVSRRLPRVGRARSNSLRIIFTVESVFVGCWLPFGVLRSLFHLARLQALPLPCSLLLALRWGLTVTTCLAFVNSSANPVIYLLLDRSFRARARFGLCARAGRQVRRISSASSLSRDDSSVFRGRSPKVNSASATW |
| | *Rattus norvegicus* | 363993 | NM_001398594 | NP_001385523 | MQSTESWNSSWGTTMSWDYSGSGSLDQVELCPAWELPYSHAIISALYLAAFVVGLTGNTFVVWLLSRRRRPRRLVDTFVLHLAAADLGFVLTLPLWAVAEARGGLWPFGDGLCKISSFALAATRCAGALLLAGMSVDRYLAVGRPLNARPLRSARCVRASCSAVWAAAFLAGLPTLLYRRLQPSLDGEGSQCAEEPSDALQGVGLLLLLLTFALPLAVTLTCYWRVSRRLLRVRRARGNSLRIIFTVESVFVGSWLPFSILRALFYLARLRALPLPCSLLLALRWGLTVATCLAFVHSCANPVIYLLLDRSFRARVRFGPCTRVGRQVRRISSASSLSRDDSSVFWGRSSKANSALAAW |
| | *Sarcophilus harrisii* | 100926064 | XM_003770662 | XP_003770710 | MSTESWSPSPGEGDEGYEWSGDYEEPNVCPIVDLPYSHIYIPLLYLAAFGAGLMGNSFVVWLLARRQGPRRLVDTFVLHLAVADLVFVVTLPFWATAAGLGGRWTFGEDLCKVSSFLIAVTRCASTLLLTGMSVDRYLAVVKLLDSRSLRNQSCILTTCGVTWTISIVAGVPSLVYRKLLPMPVGPGSRCADDPTDIFQWLSLLVLLLTFIFPLGITLFCYFLISRRLWNPPHLGRGRKNSLRIIFAIVGTFTCSWLPFSILKTFFHIAHLRTLPLDCPLLLALHWGLTLSTCLAFVNSSANPIIYLFLDRSFRARALGSCKHTHPPLTSRVSSASSLSGGDSSIFQGRALKRGRNPHMCPRLSSPSTP |
| | *Vombatus ursinus* | 114029117 | XM_027843464 | XP_027699265 | MTTESWSPSPDEGDENYEWSGDYGEPALCPTPALPFGHVFIPLLYLAAFGAGLIGNSFVVWLLAGQQGPRRLVDTFVLHLAVADLMFVVTLPFWATAVGLHGQWIFGEELCKVSSFIIAVTRCASTLLLAGMSVDRYLAVVKLLDARPLRNRGCMLTTCAVIWIISILAGVPSLVYRKLLPMPVGPGSLCGDDPADIFQGLSLLVLSLTFVLPLGIIFFCYSLISQRLWNPLQLGRGRKNSLRIIFAIVGTFTCSWLPYSILKTFFHIAHLRALPLDCPLLLALRWGLTLSTCLAFVNSSANPIIYLLLDRSFRARARGSCRRTRPPLASRVSSASSLSGGDNSIFQGWALGRGRNTRLAPRLSLLQNP |
| | *Tachyglossus aculeatus* | 119930418 | XM_038748745 | XP_038604673 | MTTKISVKSDAVSEPWNSEAGAVVKGDSGSKIKKCTENGSDWQARGSDASKEQDSPIPHRTPLMIAVAAVRAAAVVAVTAASKGGGASATSAPGPAAAALSMTSQTWGPSSPGGLGPYSGWGDYEDVGSTVTPWCPTPELPYGYAYIPALYLAAFVVGLVGNGFVVALMVGQPGPRRLVDTFVLHLAVADLVFVCTLPLWAAAAGLGNHWPFGEGLCKLSSFAMAVNRCSSALLLAGMSLDRYLVVVKMLEVRPLRTPGCVVSACGVIWAISLLAGIPSLVFRKLQPAPRASGSLCGEEVSDAFQSLSLALLVFTFALPLAVILFCYCLISRRLRQHHGLGRQRKNSLRIIFAVVGTFVGSWLPLSLFRVFYHLSRLRVLELPCSLLLALRWGLTLSACLAFINSCANPIIYLSLDRSFRVRALRGACGKASRLVRRASSVSSSRDSQDSGSLFQARGRTRAGSGPCGPGWGPEDP |
| | *Ornithorhynchus anatinus* | 100093467 | XM_029069673 | XP_028925506 | MNFAAKNSYDLTEEVPELHPKALFNCPFLLEVASIKCTVSSPLLSLPPARRPSAHPPRGLLHLPLPREPVATAVAAARAVVTAAAAASAGSGASTTSAPGPAAAALSMTSQTWGPSSPGGLGPYSGWGDYEEVGGTVMPWCPTPELPYGYAYIPALYLAAFVVGLVGNGFVVALMVGQPGPRRLVDTFVLHLAVADLVFVCTLPLWAAAAGLGNHWPFGEGLCKLSSFAMAVNRCSSALLLAGMSLDRYLVVVKMLEVRPLRTPGCVVSACGVIWAISLLAGIPSLVFRKLQPAARASGSLCGEEASDAFQSLSLALLVFTFALPLAVILFCYCLISRRLRQHHGLGRHRKNSLRIIFAVVGTFVGSWLPLSLLRAFYHLSRLGVLELPCPLLLALRWGLTLSACLAFINSCANPIIYLSLDRSFRVRALRGACGKASRLVRRASSVSSSRDSRDSGSLFQARGRTRAGSAPGGPG |
| | *Danio rerio* | 795188 | XM_073916757 | XP_073772858 | MASSTEMAHSGITMSLTSEYDYDYPINSTDENPIYTLPDAELLPMSNIYIPVLYIIMFLTGSLGN |

| | | | | | |
|---|---|---|---|---|---|
| | | | | | LFVIVVIGKRRKKSGRLVDTFVLNLALADLVFVLTLPMWAISTRYDEWPFGEVLCKISSFIIAVNRFSNIFFLTCMSVDRYLAVVRLMDSRFLRSSNCAQITCGIVWVVSFFLGSPSLAYRHLINNSVCSEDSKSSFVQGMNLLTILLTFLLPVLILGLCYGSILVNLRRHCHNPANTRTDARRRHSVKIVFAIISAFLISWLPFNCFKAIHVALLIINGDLNEDTYVVIHRGLMLSCCLAFLNSCVNPAIYFFLDQHFRRRASMLCLSCLSQNDQAHQSYITSNSYSNGTSETCSGNTSTRGRLFSLTQKA |
| | *Latimeria chalumnae* | 102365624 | XM_005988473 | XP_005988535 | MPTESLQTASHDPSDFDYNADYSNFSTEDCDGDLPYAKIYIPIFYFVIFFTGLFGNVFVIAAMTLKQTTKRLVDIFVINLAVADLVFVFTLPLWSVSAAFDDQWLFGGVLCKLSSYVIAVNRYSSIFFMTGMSVDRYMAVVKLLDSKFIRTRRCILITCTIIWIISLVMGIPSLVYRDLSTQDSEHTYCIEDQDSIIFKGISLASLFLAFVLPVIIILFCYCSISARLYSHFHHANRYDQKRKKTLKIIFTIITAFVCSWLPFNTFKTLYLLFSFQGKMPPCRVGILRQGLTITACFAFLSSCVNPIIYTFLDNHFRKRAHRLLVKALGRYTERRNSFGESWASETSSTFVSRIRANSVKELQNMNKTQQNTIPT |
| | *Xenopus tropicalis* (tropical clawed frog) | 100486541 | XM_002941165 | XP_002941211 | MSVATAMEAVTYNYDEYSDYDNDTEPDCQDENLPHGHWILPTLYLIFFFFGLTGNAIVIAIVSRRSSRRADIFILHLAISDLLFVLTLPFWASSLALGGQWSFGVHMCRASGFIIAVTRCASSLLMAIMSVDRYLAVIGGHRIHPFRTRTCSLGACCAIWLLSLVTGIPPLVFRNLVENVCVESSESRLSVGIKLATLFLTFVLPLTVVAFCYSCMAKHLWNYFGGPQNVAGGKAKPRRRHSWLRIVSCVVAAYSLSWLPYNTLSTVRLVAELGEGFPCHTTVAINQALSATAAIAFANSCTNPLIYSILDAGFRHRTKQTLPSIFPICQAVFHLPFRGGSQQSTVASTESSSSYTRSY |
| | *Ranitomeya variabilis* (Zimmermann's poison frog) | 143818606 | XM_077299858 | **XP_077155973** | MTYPSDIKPIASLWSYATSMHHRHSPAVMANESLMDDYLYIDYDEKKTNYDYDYYDNETKCQPQELPHARWILPTLYSIFFLVGFFGNMVVITVVSKRSSRRADTFILNLAVSDLLFVLTLPLWASALAQGGHWIFGAFLCQASVFVIEVTRCASSLLMAVMSVDRYVAVIKGKKIHPLRTRSFSIGTCCGIWGTSILVGCPTFVIRHLNTDNSFCEESDESLFSLGFKMVVIFLTFVLPFAVVLFCYCRMAKYLWNYFGKQVRAMRSTVKPRRGHSWLRIVFCVVGAFCFSWLPYNTVKTVLVISNLGIDISSTTKGVLEQARSATAALAFANSCSNPLIYALLDAGFRRRAHLSMPGLFSKCRSMVPMPSWTVPTTSMSMESTSTYTGN |
| | *Lithobates pipiens* (northern leopard frog) | 143927393 | XM_077451446 | XP_077307572 | MATEPPVEDYSTFYYENENEPCHPKELPHARWILSTLYSVFFVIGLIGNIVVIMVVSRRTSRRADTFILNLAVSDLTFVLTLPFWISSVAQERYWIFGTFLCQASGFAMAVTRCASSLLMAIMSIDRYVAVIKGTKVHPLRTRSCSIGACCVIWAVSILVGCPSLKTRDVDAETFVCMDSDDSTVILCFRIAVIFLTFVFPFSVMVFSYCRMAKYLWNYFGRQVNKLSRKGRPRSRHSWLRIVTCLVGAYCFSWLPYNTLNTLLLMDRLVAYIPCSAWVAIEQGLHVASALAFANSCTNPLIYALLDVGFRRRAQLALPRLLHSCRSLLSTASLTMPTMSASVESTSTYTGNKQCH |
| | *Pseudophryne corroboree* (corroboree frog) | 134989986 | XM_063950642 | XP_063806712 | MATESWTENYYDVDYDEPVKACQFEELPHARWILPTLYSVFFLVGLAGNVVVIVVVSRRSSRRADTFILNLAVSDLLFVMTLPLWASSLAQGGYWPFGAALCRASGFIIAVTRCASSFLMAIMSVDRYFAVIKGVKVHPLRVRSCSLVVCCSIWALSVLVGCPSLVSRHLSNTTACVDSDSPISLGFKMAVILLTFVLPFAIVLYCYSRMTKYLWNYFGSHVSASMTAGKPRRGHSWLRIVSCVVGAYCFSWLPFNALNALWVMSQLGADMPCSVRVTIYQALNAAAALAFANSCTNPLIYALLDAGFRRRAQSALPGLCLACRSILPMPGWSVPTNSATIESTSSYTDNR |
| | *Ranitomeya imitator* (mimic poison frog) | 138671497 | XM_069759673 | XP_069615774 | MTYPSDIKPIASLWSYATSMHHRHSPAVMANESLMGDYYHDYDENKTNYDYDYYDNETEGRLQELPHTHWILPTLYSIFFLVGFFGNMVVITVVSKRSSRRADTFILNLAVSDLLFVLTLPLWASALAQGRHWIFGAFLCQASVFVIAVTRCASSLLMAVMSVDRYVAVIKGKKMHPLRTRSCSIGTCCGIWGTSILVGCPTLVSRHLNPSNPACEDSESSFSLGFKMVVIFLTFVLPFAVVLFCYCRMAKYLWNYFGKQVRAMRSTVKPRRGHSWLRIVSCVVGAFCFSWLPYNTVNTVLVISNLGIDISFTTKGALEQALSVTAALAFANSCSNPLIYALLDAGFRRRAHLSMPGLFSKCRSMVPMPSWTVPTTSMSMESTSTYTVN |
| | *Dendropsophus ebraccatus* (hourglass treefrog) | 138793393 | XM_069971953 | XP_069828054 | MTSETQMDTYDDLDEYDYLLYYGNDSDCIYQELPHAHWILPTLYSIFFLVGFLGNVVVIVVVSKRSSRRADTFILNLAVSDLLFILTLPLWACSLAQGYWTFGDFLCKASGFIITITRCASSLLMAVMSVDRYVAVMKGKKMHPLRTRSCSLGSCCAIWATSILVGCPTLVSKHLDVADSSCVDSNGSQFSLGYRLALIVLAFILPFAVVLFCYCSMAKYLWSYFGKQMRAITGPGKPRRGHSWLRIVTCVVGAFCLSWLPFTTLNTIDVISQMGIHMPCSTKVAARQALTATAALAFANSCSNPLIYSLLDDGFRRRAQLSLPGLFLKCRSMLPSPVWSVPTTSVSMDSTSTYTGI |
| | *Phyllobates terribilis* | 141244277 | XM_073669000 | XP_073525101 | MANESLMVDYFDYDDYYGNETEWPLQELSHARWILPTLYSIFFLVGFFGNMVVITVVSKRSSRRADTFILNLAVSDLLFVLTLPLWASALAQGGLWPFGAFLCKTSGFVIAVTRCASSWLMAVMSVDRYVTVIKGKKMHPLRTRSCSIGTCCGIWGTSILVGCPALVSRHLETCSSKCVDSDGSSFSLGFKMVVIFLTFVLPFAVVLFCYCRMAKYLWNYFGKQVRAMTSAGKPRRGHSWLRIVSCVVGAFCLSWLPYNTLNTVEVISKLGIDIPYTMDEAIDQALSATAALAFANSCSNPLIYALLDAGFRRHAHLSMPGLFYKCRSMVPMPSWTMPTTSMSMESTSTYTGN |
| | *Pyxicephalus adspersus* | 140342250 | XM_072428407 | XP_072284508 | MTTETLVDDYDYDYYFGNETEEPCYFEELPHAYWILPTLCSIFFVCGLIGNAVVIMVVSRRSSRRADTFILNLAVSDLTFVFTLPFWASSQAQGFWPFGTFLCQVSGFVIAITRCASSLLMAIMSIDRYVAVIRGTKMHPLRTRKYSIGCCCVIWTVSILIGCPSLLTREVNTETLACVDSNATFSLRFRIAVVFLTFVFPFAVIVFCYCRMAKYLWNYFGKQVNKVSNNGRPRSRHSWLRIVTCVVGAYCLSWLPFNTLNTVLLMAKLGVDIPCTVLVALRQGLDATATLAFANSCTNPLIYALLDAGFRRRAQLALPRLFHTCRSLMSTTGSTMPTTSASVESTSTYTGIR |
| | *Bufo bufo* (common toad) | 120994039 | XM_040422500 | XP_040278434 | MSLHQSPDATGNESWPEYYYDIEEECQFQELPHARWILPTLYSIFFLVGFLGNVVVIAMVSKRTSRRSDIFILNLAVSDLLFVFTLPLWASSSARSGYWPFGVYLCKASGFVIAVNRCASSFLMAVMSVDRYVTVIKGKKMHPLRTRSYSLAACCTVWAISILVGCPSLVSRDLDSNNFACVDSYQWLFTIQYKMVLTFLTFVLPFAVVLFCYCSMAKYLWNYFGKQVRAMTSTGKPQRGHSWLRIVLCVVGAFCLSWLPFNALNTVLVIHNLLETNMTCSTEVFIGQALSVAAALAFANSCSNPLIYALLDSGFRRRARRCLPGLFLTCCSALHIPTWTVPITITSMESTTPTLTTDDHASQPRVWNRLMQTVLRKICRQQPGT |
| | *Spea bombifrons* (plains spadefoot toad) | 128473933 | XM_053456155 | XP_053312130 | MWATLYLPWWGGQNHRKAQHKYVYRLQDLYKPRQPYTAHSFLSPEQGTEDRMMDVEYSSTAYYDYYDQNNETDHDCPPQDSAHARWVLPMLYLVFFVLGFIGNVMVIAVLSRRNARRADTFILNLAVSDLLFVLTLPLWATSLALSGFWPFGVHICRASGFLIAVTRCASSMLMAIMSVDRYLAVKKGQKVHLLRSPTCSFGTCCTVWAFSFVSGIPAIFSRRIDNRNLTCMESSASPISTALKAANIMVTFALPFSVVVFCYCSMAKHLWKYFGDQKNVFSGTTKRRYGHSWLRIVSCVVAAYCLSWFPFNTLSIVTVVAQLGPALPCHTLDAIRQAISAAAALAFANSCTNPLIYALLDAGFRRRAKQAIPWLLCVCCPLKLLSFRGWSFSGSPGSTESTSTYTDSAVRQPVK |
| | *Pleurodeles* | 138301999 | XM_069242427 | XP_069098528 | MMENQSQVYWADQTTTYDPYEDTTDYEDSDIWDCQDEKLPHGNIYLPILYSVSFFVGFLGNIFVI |

| | | | | | |
|---|---|---|---|---|---|
| | *waltl* (Iberian ribbed newt) | | | | VLMVRKQGTRRLVDTFVINLALADLVFVCTLPLWSVTAALDNQWHFGNALCKMSSFVIMVNRCSSILFLAGMSVDRYLAIVKLLDSRCVRTRKCVLTTCAAVWAVSFLMGIPSLIYRKVWTDKDTGSTYCMEDMKPEASIIKLVSLLFTFVLPLFTILFCYCSILVRLSTHVYQGKRTKNSLKIIFTIIAAFICSWLPFNLLSAIFLFSKLEWTALSCSTTTALKWSLTVTACFAFMNSCVNPIIYIFLDRYFRYQVLKNTFAFFPGAAFKGRRPSLGSSFSLPNDSFSIFSSKSKSTGMSTY |
| | *Ambystoma mexicanum* (axolotl) | 138522110 | XM_069635377 | XP_069491478 | MKSCGQRNDPSPFHPPHRDKRVCQPSTRKFANKCVPASAPYNKNPAVNRWSKLADIHLLPMTMRENVSEHYWAAEMTTEVWDNGEFVTGAYEDCPDQDLPYGNTYIPILYSIAFLIGFFGNIFVIVLMVRKQGARRLVDTFIINLAVADLVFVCTLPLWSATVASEQQWYFGDALCKISSFVIMVNRCSSILFLTGMSVDRYLAIVKLMDSRCVRTRKCVFTTCLTVWLISFLMGIPSLVFRKTWTEEDTNTSYCMEEMSPMVSGIKLTLLLFTFVLPMCTILFCYCSILFRLSTHIYPGKRTKNSLKIIFTIVGTFICCWLPFNLLSAIYLFSKMEWTNLTCSSATALKWSLSITACFAFMNSCVNPIIYIFLDRYFRYQVLRTTFTLLPRSLFNGRRPSLGSSCSLPSESFSIFSAKTKSTVI |
| | *Microcaecilia unicolor* | 115482117 | XM_030221686 | XP_030077546 | MAAPLPCTNGLSLETVVNDWNEIYDDYTTDENSMYHGVFSLPQCPQKELPYGHVYIPFLYSITFLVGVFGNLFVVVLMLRKQSSRRLVDTFVINLAIADLVFVCTLPFWAGAAAAGNQWLYGATLCIASSYSISVTRYSGTLFLMAMSLDRYLAIVRLHKSQALRTRGCIRTTSAGIWVISLLLGIPTLTSRRLLTDQETNRTVCSDICSEVQSGTKLALLFFTFLLPFAIILFCYWSVSARLRRHCQQGRNRRNSLKIIFTVVGTFLCSWLPFNTLSCIALFYKLEWVALSCRSSQALSWGLTLTACLAFTNSCVNPIIYTFLDRHFRQQILQKPVWLFQKPKAAETTPGSSITFSSQSSAFFPPRG |
| | *Rhinatrema bivittatum* (two-lined caecilian) | 115074284 | XM_029573602 | XP_029429462 | MDGLSLKMGVTAWSVAYEDYTTDAGLIGCPQEELPYSHVYIPVLYVISFFLGFFGNLFVVALMMGKRGPRRLVDTFVINLAVADLVFVLTLPFWAVSAALGNQWLFGAGLCTVSSYSITVTRYSGTLFLMGMSLDRYLAIVRPHRSRALRTPRCVLTTSGSIWTVSLLLGIPSLVSRRLSKETTHGVCGEASSVILSSTKLAMLLLTFLLPLVIILFCYCSILARLRRHCQQGRSRRNSLKIIFTIVGTFLCSWLPFNTLSSIAIFCSLDWVSLSCGSMQTLRWGLTLTACLAFTHSCVNPIIYIFLDRHFRQQLLRKSLQLFQKPRPVEATAGSSITFSSHSSALFPRRG |
| | *Geotrypetes seraphini* (Gabon caecilian) | 117347567 | XM_033918662 | XP_033774553 | MGAPLLYTDGISLETMLVDYWNASYEDYTTDENSRDVPEECLQEELPYGHVYIPILYSIAFLVGFFGNLFVVGLMLKKQGSRRLVDTFVVNLAIADLVFVCTLPFWARSAAVGNQWLYGATLCIASSYSITVTRYSGILFLMGMSLDRYLAIVRLHKSRAVRTRQCILLTSSGIWAISLLLGIPTLASRRLLTNTETNCSVCKDISSAVLSGTRLALLFLTFFLPLAIILFCYCAIVARLRQHCQQGRNRRNSLKIIFTIVGTFLCSWMPFNTLSCITLFYKLEWLSLSCRFSQALSWGLTLTNCLAFMNSCMNPVIYIFLDRHFRQQILQKSIQLLQKPRTAETTPGSSITFSSKSSALFPPRG |
| | *Xenopus laevis* (African clawed frog) | 108707616 | XM_018245607 | XP_018101096 | MADVTPFYDEYALYDNDTEPFCQNENLPHGQWILPALYLFFFFFGLTGNAVVIAIVSKRSSRRADIFILNLAISDLFFVLTLPFWASSLALGGQWPFGIHLCRASGFIIAVTRCASSLLMAIMSVDRYLAVIGGHRIHPFRTRTCSMGACCIIWFISLVSGIPPLMFRHLEQMVCVESSESVLSTGIKLATLFLTFVLPLTVVLFCYSCMAKHLWNYFGGHQNVATSKLKPRRRHSWLRIVSCVVAAYSLSWLPYNTLSAVTLVDQLGEGLSCHTLVVINQALSATAAIAFANSCTNPLIYSILDAGFRHRTKQALPRIFPMCTAVFNLSFPGWSQQSTVASTESSSSYTLNTSKAVANANKKEGSNTTEITTWHMPNTPASL |

**6×His-Xt-CXCL17**

```
1    CTT TAA GAA GGA GAT ATA ATG CAT CAT CAC CAC CAT CAC TGG ACT CCT CAA CGT CAG GAC AGT TCA GGT CGC ACC AAT GAG AAC CGT GGT
     GAA ATT CTT CCT CTA TAT TAC GTA GTA GTG GTG GTA GTG ACC TGA GGA GTT GCA GTC CTG TCA AGT CCA GCG TGG TTA CTC TTG GCA CCA
                              M   H   H   H   H   H   H   W   T   P   Q   R   Q   D   S   S   G   R   T   N   E   N   R   G

91   CTG GCC AAA CGC AAT GGC CAC AAA GAT GCT CTG GGT TCT CCA CTT CGT CCT GAA CGC CAC CAT CTG CGT TGC TGT GAA CGC CTC AGC CCG
     GAC CGG TTT GCG TTA CCG GTG TTT CTA CGA GAC CCA AGA GGT GAA GCA GGA CTT GCG GTG GTA GAC GCA ACG ACA CTT GCG GAG TCG GGC
      L   A   K   R   N   G   H   K   D   A   L   G   S   P   L   R   P   E   R   H   H   L   R   C   C   E   R   L   S   P

181  CAG CAT CCA CAG CGT AAG CGC CGT CAC ATG CGC GCA CCA CGT GCG CAC CCG ATT CAT CAG CGC CGT CGC TGC ACC ACG AAC TGC CGT GGC
     GTC GTA GGT GTC GCA TTC GCG GCA GTG TAC GCG CGT GGT GCA CGC GTG GGC TAA GTA GTC GCG GCA GCG ACG TGG TGC TTG ACG GCA CCG
      Q   H   P   Q   R   K   R   R   H   M   R   A   P   R   A   H   P   I   H   Q   R   R   R   C   T   T   N   C   R   G

271  AAG AAA CCG TCC AAA TTC ATG GTT CCG TTG CCA TTA TCG TAA GCG GCC GCA CTC GAG CAC CAC CAC CAC
     TTC TTT GGC AGG TTT AAG TAC CAA GGC AAC GGT AAT AGC ATT CGC CGG CGT GAG CTC GTG GTG GTG GTG
      K   K   P   S   K   F   M   V   P   L   P   L   S   *
```

**6×His-SmBiT-Xt-CXCL17**

```
1    CTT TAA GAA GGA GAT ATA ATG CAT CAT CAC CAC CAT CAC GGT GTG ACC GGC TAC CGT CTG TTT GAA GAA ATT CTG GGC GGC TGG ACT CCT
     GAA ATT CTT CCT CTA TAT TAC GTA GTA GTG GTG GTA GTG CCA CAC TGG CCG ATG GCA GAC AAA CTT CTT TAA GAC CCG CCG ACC TGA GGA
                              M   H   H   H   H   H   H   G   V   T   G   Y   R   L   F   E   E   I   L   G   G   W   T   P

91   CAA CGT CAG GAC AGT TCA GGT CGC ACC AAT GAG AAC CGT GGT CTG GCC AAA CGC AAT GGC CAC AAA GAT GCT CTG GGT TCT CCA CTT CGT
     GTT GCA GTC CTG TCA AGT CCA GCG TGG TTA CTC TTG GCA CCA GAC CGG TTT GCG TTA CCG GTG TTT CTA CGA GAC CCA AGA GGT GAA GCA
      Q   R   Q   D   S   S   G   R   T   N   E   N   R   G   L   A   K   R   N   G   H   K   D   A   L   G   S   P   L   R

181  CCT GAA CGC CAC CAT CTG CGT TGC TGT GAA CGC CTC AGC CCG CAG CAT CCA CAG CGT AAG CGC CGT CAC ATG CGC GCA CCA CGT GCG CAC
     GGA CTT GCG GTG GTA GAC GCA ACG ACA CTT GCG GAG TCG GGC GTC GTA GGT GTC GCA TTC GCG GCA GTG TAC GCG CGT GGT GCA CGC GTG
      P   E   R   H   H   L   R   C   C   E   R   L   S   P   Q   H   P   Q   R   K   R   R   H   M   R   A   P   R   A   H

271  CCG ATT CAT CAG CGC CGT CGC TGC ACC ACG AAC TGC CGT GGC AAG AAA CCG TCC AAA TTC ATG GTT CCG TTG CCA TTA TCG TAA GCG GCC
     GGC TAA GTA GTC GCG GCA GCG ACG TGG TGC TTG ACG GCA CCG TTC TTT GGC AGG TTT AAG TAC CAA GGC AAC GGT AAT AGC ATT CGC CGG
      P   I   H   Q   R   R   R   C   T   T   N   C   R   G   K   K   P   S   K   F   M   V   P   L   P   L   S   *

361  GCA CTC GAG CAC CAC CAC CAC
     CGT GAG CTC GTG GTG GTG GTG
```

**Fig. S1.** The nucleotide sequence and amino acid sequence of the Xt-CXCL17 constructs overexpressed in *E. coli*. The mature peptide of Xt-CXCL17 is shown in red, that of SmBiT in blue.

### Untagged Xt-GPR25 in pcDNA3.1 vector or PB-TRE vector

```
1     ATG AGT GTT GCT ACA GCC ATG GAG GCC GTT ACT TAT AAT TAC GAT GAA TAC AGT GAT TAT GAT AAC GAC ACG GAA CCC GAC TGC CAG GAT
      TAC TCA CAA CGA TGT CGG TAC CTC CGG CAA TGA ATA TTA ATG CTA CTT ATG TCA CTA ATA CTA TTG CTG TGC CTT GGG CTG ACG GTC CTA
       M   S   V   A   T   A   M   E   A   V   T   Y   N   Y   D   E   Y   S   D   Y   D   N   D   T   E   P   D   C   Q   D

91    GAG AAT TTG CCC CAT GGG CAC TGG ATT CTG CCC ACC CTC TAT CTG ATC TTT TTT TTC TTT GGC CTG ACT GGG AAC GCC ATA GTG ATC GCC
      CTC TTA AAC GGG GTA CCC GTG ACC TAA GAC GGG TGG GAG ATA GAC TAG AAA AAA AAG AAA CCG GAC TGA CCC TTG CGG TAT CAC TAG CGG
       E   N   L   P   H   G   H   W   I   L   P   T   L   Y   L   I   F   F   F   F   G   L   T   G   N   A   I   V   I   A

181   ATT GTG TCC AGG CGG AGT TCA CGT CGG GCA GAT ATT TTC ATT CTC CAC CTC GCC ATC TCT GAC TTG CTT TTT GTA TTG ACT CTG CCA TTT
      TAA CAC AGG TCC GCC TCA AGT GCA GCC CGT CTA TAA AAG TAA GAG GTG GAG CGG TAG AGA CTG AAC GAA AAA CAT AAC TGA GAC GGT AAA
       I   V   S   R   R   S   S   R   R   A   D   I   F   I   L   H   L   A   I   S   D   L   L   F   V   L   T   L   P   F

271   TGG GCT TCT TCG TTG GCG CTG GGG GGG CAG TGG AGC TTT GGG GTA CAC ATG TGC CGC GCC AGT GGG TTT ATT ATC GCG GTT ACC CGC TGT
      ACC CGA AGA AGC AAC CGC GAC CCC CCC GTC ACC TCG AAA CCC CAT GTG TAC ACG GCG CGG TCA CCC AAA TAA TAG CGC CAA TGG GCG ACA
       W   A   S   S   L   A   L   G   G   Q   W   S   F   G   V   H   M   C   R   A   S   G   F   I   I   A   V   T   R   C

361   GCC AGC TCC TTG CTA ATG GCC ATC ATG AGT GTT GAC CGT TAC CTT GCG GTG ATT GGG GGG CAT AGG ATT CAC CCT TTC CGC ACT AGG ACC
      CGG TCG AGG AAC GAT TAC CGG TAG TAC TCA CAA CTG GCA ATG GAA CGC CAC TAA CCC CCC GTA TCC TAA GTG GGA AAG GCG TGA TCC TGG
       A   S   S   L   L   M   A   I   M   S   V   D   R   Y   L   A   V   I   G   G   H   R   I   H   P   F   R   T   R   T

451   TGC TCC CTG GGG GCT TGC TGC GCC ATT TGG CTG CTC TCC CTG GTA ACT GGG ATC CCC CCT CTG GTT TTC CGG AAC CTG GTG GAG AAC GTG
      ACG AGG GAC CCC CGA ACG ACG CGG TAA ACC GAC GAG AGG GAC CAT TGA CCC TAG GGG GGA GAC CAA AAG GCC TTG GAC CAC CTC TTG CAC
       C   S   L   G   A   C   C   A   I   W   L   L   S   L   V   T   G   I   P   P   L   V   F   R   N   L   V   E   N   V

541   TGC GTG GAA TCC TCA GAA TCG AGA CTC AGC GTG GGC ATC AAG TTG GCC ACT TTG TTC TTG ACC TTC GTC CTT CCA CTT ACC GTG GTG GCG
      ACG CAC CTT AGG AGT CTT AGC TCT GAG TCG CAC CCG TAG TTC AAC CGG TGA AAC AAG AAC TGG AAG CAG GAA GGT GAA TGG CAC CAC CGC
       C   V   E   S   S   E   S   R   L   S   V   G   I   K   L   A   T   L   F   L   T   F   V   L   P   L   T   V   V   A

631   TTT TGC TAC AGC TGC ATG GCT AAG CAC CTC TGG AAT TAT TTT GGG GGC CCT CAG AAT GTT GCT GGG GGC AAA GCG AAG CCT CGG CGT AGG
      AAA ACG ATG TCG ACG TAC CGA TTC GTG GAG ACC TTA ATA AAA CCC CCG GGA GTC TTA CAA CGA CCC CCG TTT CGC TTC GGA GCC GCA TCC
       F   C   Y   S   C   M   A   K   H   L   W   N   Y   F   G   G   P   Q   N   V   A   G   G   K   A   K   P   R   R   R

721   CAC AGC TGG CTA CGT ATA GTC TCG TGT GTT GTG GCA GCG TAT TCA TTG TCT TGG TTG CCC TAT AAT ACC CTC AGC ACT GTA AGA CTG GTG
      GTG TCG ACC GAT GCA TAT CAG AGC ACA CAA CAC CGT CGC ATA AGT AAC AGA ACC AAC GGG ATA TTA TGG GAG TCG TGA CAT TCT GAC CAC
       H   S   W   L   R   I   V   S   C   V   V   A   A   Y   S   L   S   W   L   P   Y   N   T   L   S   T   V   R   L   V

811   GCC GAG CTT GGA GAA GGC TTT CCC TGC CAC ACC ACG GTC GCC ATT AAC CAA GCG CTC AGT GCC ACA GCC GCC ATT GCC TTT GCC AAC AGC
      CGG CTC GAA CCT CTT CCG AAA GGG ACG GTG TGG TGC CAG CGG TAA TTG GTT CGC GAG TCA CGG TGT CGG CGG TAA CGG AAA CGG TTG TCG
       A   E   L   G   E   G   F   P   C   H   T   T   V   A   I   N   Q   A   L   S   A   T   A   A   I   A   F   A   N   S

901   TGC ACC AAC CCC CTC ATC TAC TCC ATC CTC GAT GCC GGC TTT CGG CAC AGG ACA AAG CAG ACC TTG CCC AGC ATT TTC CCC ATA TGC CAA
      ACG TGG TTG GGG GAG TAG ATG AGG TAG GAG CTA CGG CCG AAA GCC GTG TCC TGT TTC GTC TGG AAC GGG TCG TAA AAG GGG TAT ACG GTT
       C   T   N   P   L   I   Y   S   I   L   D   A   G   F   R   H   R   T   K   Q   T   L   P   S   I   F   P   I   C   Q

991   GCA GTT TTC CAT CTC CCC TTC CGT GGT GGG AGC CAA CAG TCT ACA GTT GCC AGC ACC GAG AGT AGC AGC TCC TAC ACA CGC AGC TAC TGA
      CGT CAA AAG GTA GAG GGG AAG GCA CCA CCC TCG GTT GTC AGA TGT CAA CGG TCG TGG CTC TCA TCG TCG AGG ATG TGT GCG TCG ATG ACT
       A   V   F   H   L   P   F   R   G   G   S   Q   Q   S   T   V   A   S   T   E   S   S   S   S   Y   T   R   S   Y   *
```

### Xt-GPR25-LgBiT in MCS2 of pTRE3G-BI vector (SmBiT-ARRB2 in MCS1)

```
1     ATG AGT GTT GCT ACA GCC ATG GAG GCC GTT ACT TAT AAT TAC GAT GAA TAC AGT GAT TAT GAT AAC GAC ACG GAA CCC GAC TGC CAG GAT
      TAC TCA CAA CGA TGT CGG TAC CTC CGG CAA TGA ATA TTA ATG CTA CTT ATG TCA CTA ATA CTA TTG CTG TGC CTT GGG CTG ACG GTC CTA
       M   S   V   A   T   A   M   E   A   V   T   Y   N   Y   D   E   Y   S   D   Y   D   N   D   T   E   P   D   C   Q   D

91    GAG AAT TTG CCC CAT GGG CAC TGG ATT CTG CCC ACC CTC TAT CTG ATC TTT TTT TTC TTT GGC CTG ACT GGG AAC GCC ATA GTG ATC GCC
      CTC TTA AAC GGG GTA CCC GTG ACC TAA GAC GGG TGG GAG ATA GAC TAG AAA AAA AAG AAA CCG GAC TGA CCC TTG CGG TAT CAC TAG CGG
       E   N   L   P   H   G   H   W   I   L   P   T   L   Y   L   I   F   F   F   F   G   L   T   G   N   A   I   V   I   A

181   ATT GTG TCC AGG CGG AGT TCA CGT CGG GCA GAT ATT TTC ATT CTC CAC CTC GCC ATC TCT GAC TTG CTT TTT GTA TTG ACT CTG CCA TTT
      TAA CAC AGG TCC GCC TCA AGT GCA GCC CGT CTA TAA AAG TAA GAG GTG GAG CGG TAG AGA CTG AAC GAA AAA CAT AAC TGA GAC GGT AAA
       I   V   S   R   R   S   S   R   R   A   D   I   F   I   L   H   L   A   I   S   D   L   L   F   V   L   T   L   P   F

271   TGG GCT TCT TCG TTG GCG CTG GGG GGG CAG TGG AGC TTT GGG GTA CAC ATG TGC CGC GCC AGT GGG TTT ATT ATC GCG GTT ACC CGC TGT
      ACC CGA AGA AGC AAC CGC GAC CCC CCC GTC ACC TCG AAA CCC CAT GTG TAC ACG GCG CGG TCA CCC AAA TAA TAG CGC CAA TGG GCG ACA
       W   A   S   S   L   A   L   G   G   Q   W   S   F   G   V   H   M   C   R   A   S   G   F   I   I   A   V   T   R   C

361   GCC AGC TCC TTG CTA ATG GCC ATC ATG AGT GTT GAC CGT TAC CTT GCG GTG ATT GGG GGG CAT AGG ATT CAC CCT TTC CGC ACT AGG ACC
      CGG TCG AGG AAC GAT TAC CGG TAG TAC TCA CAA CTG GCA ATG GAA CGC CAC TAA CCC CCC GTA TCC TAA GTG GGA AAG GCG TGA TCC TGG
       A   S   S   L   L   M   A   I   M   S   V   D   R   Y   L   A   V   I   G   G   H   R   I   H   P   F   R   T   R   T

451   TGC TCC CTG GGG GCT TGC TGC GCC ATT TGG CTG CTC TCC CTG GTA ACT GGG ATC CCC CCT CTG GTT TTC CGG AAC CTG GTG GAG AAC GTG
      ACG AGG GAC CCC CGA ACG ACG CGG TAA ACC GAC GAG AGG GAC CAT TGA CCC TAG GGG GGA GAC CAA AAG GCC TTG GAC CAC CTC TTG CAC
       C   S   L   G   A   C   C   A   I   W   L   L   S   L   V   T   G   I   P   P   L   V   F   R   N   L   V   E   N   V

541   TGC GTG GAA TCC TCA GAA TCG AGA CTC AGC GTG GGC ATC AAG TTG GCC ACT TTG TTC TTG ACC TTC GTC CTT CCA CTT ACC GTG GTG GCG
      ACG CAC CTT AGG AGT CTT AGC TCT GAG TCG CAC CCG TAG TTC AAC CGG TGA AAC AAG AAC TGG AAG CAG GAA GGT GAA TGG CAC CAC CGC
       C   V   E   S   S   E   S   R   L   S   V   G   I   K   L   A   T   L   F   L   T   F   V   L   P   L   T   V   V   A

631   TTT TGC TAC AGC TGC ATG GCT AAG CAC CTC TGG AAT TAT TTT GGG GGC CCT CAG AAT GTT GCT GGG GGC AAA GCG AAG CCT CGG CGT AGG
      AAA ACG ATG TCG ACG TAC CGA TTC GTG GAG ACC TTA ATA AAA CCC CCG GGA GTC TTA CAA CGA CCC CCG TTT CGC TTC GGA GCC GCA TCC
       F   C   Y   S   C   M   A   K   H   L   W   N   Y   F   G   G   P   Q   N   V   A   G   G   K   A   K   P   R   R   R

721   CAC AGC TGG CTA CGT ATA GTC TCG TGT GTT GTG GCA GCG TAT TCA TTG TCT TGG TTG CCC TAT AAT ACC CTC AGC ACT GTA AGA CTG GTG
      GTG TCG ACC GAT GCA TAT CAG AGC ACA CAA CAC CGT CGC ATA AGT AAC AGA ACC AAC GGG ATA TTA TGG GAG TCG TGA CAT TCT GAC CAC
       H   S   W   L   R   I   V   S   C   V   V   A   A   Y   S   L   S   W   L   P   Y   N   T   L   S   T   V   R   L   V

811   GCC GAG CTT GGA GAA GGC TTT CCC TGC CAC ACC ACG GTC GCC ATT AAC CAA GCG CTC AGT GCC ACA GCC GCC ATT GCC TTT GCC AAC AGC
      CGG CTC GAA CCT CTT CCG AAA GGG ACG GTG TGG TGC CAG CGG TAA TTG GTT CGC GAG TCA CGG TGT CGG CGG TAA CGG AAA CGG TTG TCG
       A   E   L   G   E   G   F   P   C   H   T   T   V   A   I   N   Q   A   L   S   A   T   A   A   I   A   F   A   N   S

901   TGC ACC AAC CCC CTC ATC TAC TCC ATC CTC GAT GCC GGC TTT CGG CAC AGG ACA AAG CAG ACC TTG CCC AGC ATT TTC CCC ATA TGC CAA
      ACG TGG TTG GGG GAG TAG ATG AGG TAG GAG CTA CGG CCG AAA GCC GTG TCC TGT TTC GTC TGG AAC GGG TCG TAA AAG GGG TAT ACG GTT
       C   T   N   P   L   I   Y   S   I   L   D   A   G   F   R   H   R   T   K   Q   T   L   P   S   I   F   P   I   C   Q

991   GCA GTT TTC CAT CTC CCC TTC CGT GGT GGG AGC CAA CAG TCT ACA GTT GCC AGC ACC GAG AGT AGC AGC TCC TAC ACA CGC AGC TAC CCA
      CGT CAA AAG GTA GAG GGG AAG GCA CCA CCC TCG GTT GTC AGA TGT CAA CGG TCG TGG CTC TCA TCG TCG AGG ATG TGT GCG TCG ATG GGT
       A   V   F   H   L   P   F   R   G   G   S   Q   Q   S   T   V   A   S   T   E   S   S   S   S   Y   T   R   S   Y   P

1081  CCG GTC GGT ACC GGC GGA GGG TCT AGC AGT GGC GGT GGG ATG GTC TTC ACA CTC GAA GAT TTC GTT GGG GAC TGG GAA CAG ACA GCC GCC
      GGC CAG CCA TGG CCG CCT CCC AGA TCG TCA CCG CCA CCC TAC CAG AAG TGT GAG CTT CTA AAG CAA CCC CTG ACC CTT GTC TGT CGG CGG
       P   V   G   T   G   G   G   S   S   S   G   G   G   M   V   F   T   L   E   D   F   V   G   D   W   E   Q   T   A   A
```

1171 TAC AAC CTG GAC CAA GTC CTT GAA CAG GGA GGT GTG TCC AGT TTG CTG CAG AAT CTC GCC GTG TCC GTA ACT CCG ATC CAA AGG ATT GTC
ATG TTG GAC CTG GTT CAG GAA CTT GTC CCT CCA CAC AGG TCA AAC GAC GTC TTA GAG CGG CAC AGG CAT TGA GGC TAG GTT TCC TAA CAG
Y N L D Q V L E Q G G V S S L L Q N L A V S V T P I Q R I V

1261 CGG AGC GGT GAA AAT GCC CTG AAG ATC GAC ATC CAT GTC ATC ATC CCG TAT GAA GGT CTG AGC GCC GAC CAA ATG GCC CAG ATC GAA GAG
GCC TCG CCA CTT TTA CGG GAC TTC TAG CTG TAG GTA CAG TAG TAG GGC ATA CTT CCA GAC TCG CGG CTG GTT TAC CGG GTC TAG CTT CTC
R S G E N A L K I D I H V I I P Y E G L S A D Q M A Q I E E

1351 GTG TTT AAG GTG GTG TAC CCT GTG GAT GAT CAT CAC TTT AAG GTG ATC CTG CCC TAT GGC ACA CTG GTA ATC GAC GGG GTT ACG CCG AAC
CAC AAA TTC CAC CAC ATG GGA CAC CTA CTA GTA GTG AAA TTC CAC TAG GAC GGG ATA CCG TGT GAC CAT TAG CTG CCC CAA TGC GGC TTG
V F K V V Y P V D D H H F K V I L P Y G T L V I D G V T P N

1441 ATG CTG AAC TAT TTC GGA CGG CCG TAT GAA GGC ATC GCC GTG TTC GAC GGC AAA AAG ATC ACT GTA ACA GGG ACC CTG TGG AAC GGC AAC
TAC GAC TTG ATA AAG CCT GCC GGC ATA CTT CCG TAG CGG CAC AAG CTG CCG TTT TTC TAG TGA CAT TGT CCC TGG GAC ACC TTG CCG TTG
M L N Y F G R P Y E G I A V F D G K K I T V T G T L W N G N

1531 AAA ATT ATC GAC GAG CGC CTG ATC ACC CCC GAC GGC TCC ATG CTG TTC CGA GTA ACC ATC AAC AGT TAA
TTT TAA TAG CTG CTC GCG GAC TAG TGG GGG CTG CCG AGG TAC GAC AAG GCT CAT TGG TAG TTG TCA ATT
K I I D E R L I T P D G S M L F R V T I N S *

**sLgBiT-Xt-GPR25-LgBiT in PB-TRE vector**

1 ATG AAC TCC TTC TCC ACA AGC GCC TTC GGT CCA GTT GCC TTC TCC CTG GGC CTG CTC CTG GTG TTG CCT GCT GCC TTC CCT GCC CCA GTC
TAC TTG AGG AAG AGG TGT TCG CGG AAG CCA GGT CAA CGG AAG AGG GAC CCG GAC GAG GAC CAC AAC GGA CGA CGG AAG GGA CGG GGT CAG
M N S F S T S A F G P V A F S L G L L L V L P A A F P A P V

91 TTC ACA CTC GAA GAT TTC GTT GGG GAC TGG GAA CAG ACA GCC GCC TAC AAC CTG GAC CAA GTC CTT GAA CAG GGA GGT GTG TCC AGT TTG
AAG TGT GAG CTT CTA AAG CAA CCC CTG ACC CTT GTC TGT CGG CGG ATG TTG GAC CTG GTT CAG GAA CTT GTC CCT CCA CAC AGG TCA AAC
F T L E D F V G D W E Q T A A Y N L D Q V L E Q G G V S S L

181 CTG CAG AAT CTC GCC GTG TCC GTA ACT CCG ATC CAA AGG ATT GTC CGG AGC GGT GAA AAT GCC CTG AAG ATC GAC ATC CAT GTC ATC ATC
GAC GTC TTA GAG CGG CAC AGG CAT TGA GGC TAG GTT TCC TAA CAG GCC TCG CCA CTT TTA CGG GAC TTC TAG CTG TAG GTA CAG TAG TAG
L Q N L A V S V T P I Q R I V R S G E N A L K I D I H V I I

271 CCG TAT GAA GGT CTG AGC GCC GAC CAA ATG GCC CAG ATC GAA GAG GTG TTT AAG GTG GTG TAC CCT GTG GAT GAT CAT CAC TTT AAG GTG
GGC ATA CTT CCA GAC TCG CGG CTG GTT TAC CGG GTC TAG CTT CTC CAC AAA TTC CAC CAC ATG GGA CAC CTA CTA GTA GTG AAA TTC CAC
P Y E G L S A D Q M A Q I E E V F K V V Y P V D D H H F K V

361 ATC CTG CCC TAT GGC ACA CTG GTA ATC GAC GGG GTT ACG CCG AAC ATG CTG AAC TAT TTC GGA CGG CCG TAT GAA GGC ATC GCC GTG TTC
TAG GAC GGG ATA CCG TGT GAC CAT TAG CTG CCC CAA TGC GGC TTG TAC GAC TTG ATA AAG CCT GCC GGC ATA CTT CCG TAG CGG CAC AAG
I L P Y G T L V I D G V T P N M L N Y F G R P Y E G I A V F

451 GAC GGC AAA AAG ATC ACT GTA ACA GGG ACC CTG TGG AAC GGC AAC AAA ATT ATC GAC GAG CGC CTG ATC ACC CCC GAC GGC TCC ATG CTG
CTG CCG TTT TTC TAG TGA CAT TGT CCC TGG GAC ACC TTG CCG TTG TTT TAA TAG CTG CTC GCG GAC TAG TGG GGG CTG CCG AGG TAC GAC
D G K K I T V T G T L W N G N K I I D E R L I T P D G S M L

541 TTC CGA GTA ACC ATC AAC AGT GGT GGC GGC TCT GGT GGT GGC AGC GGC GGT GGT ACC AGT GTT GCT ACA GCC ATG GAG GCC GTT ACT TAT
AAG GCT CAT TGG TAG TTG TCA CCA CCG CCG AGA CCA CCA CCG TCG CCG CCA CCA TGG TCA CAA CGA TGT CGG TAC CTC CGG CAA TGA ATA
F R V T I N S G G G S G G G S G G G T S V A T A M E A V T Y

631 AAT TAC GAT GAA TAC AGT GAT TAT GAT AAC GAC ACG GAA CCC GAC TGC CAG GAT GAG AAT TTG CCC CAT GGG CAC TGG ATT CTG CCC ACC
TTA ATG CTA CTT ATG TCA CTA ATA CTA TTG CTG TGC CTT GGG CTG ACG GTC CTA CTC TTA AAC GGG GTA CCC GTG ACC TAA GAC GGG TGG
N Y D E Y S D Y D N D T E P D C Q D E N L P H G H W I L P T

721 CTC TAT CTG ATC TTT TTT TTC TTT GGC CTG ACT GGG AAC GCC ATA GTG ATC GCC ATT GTG TCC AGG CGG AGT TCA CGT CGG GCA GAT ATT
GAG ATA GAC TAG AAA AAA AAG AAA CCG GAC TGA CCC TTG CGG TAT CAC TAG CGG TAA CAC AGG TCC GCC TCA AGT GCA GCC CGT CTA TAA
L Y L I F F F F G L T G N A I V I A I V S R R S S R R A D I

811 TTC ATT CTC CAC CTC GCC ATC TCT GAC TTG CTT TTT GTA TTG ACT CTG CCA TTT TGG GCT TCT TCG TTG GCG CTG GGG GGG CAG TGG AGC
AAG TAA GAG GTG GAG CGG TAG AGA CTG AAC GAA AAA CAT AAC TGA GAC GGT AAA ACC CGA AGA AGC AAC CGC GAC CCC CCC GTC ACC TCG
F I L H L A I S D L L F V L T L P F W A S S L A L G G Q W S

901 TTT GGG GTA CAC ATG TGC CGC GCC AGT GGG TTT ATT ATC GCG GTT ACC CGC TGT GCC AGC TCC TTG CTA ATG GCC ATC ATG AGT GTT GAC
AAA CCC CAT GTG TAC ACG GCG CGG TCA CCC AAA TAA TAG CGC CAA TGG GCG ACA CGG TCG AGG AAC GAT TAC CGG TAG TAC TCA CAA CTG
F G V H M C R A S G F I I A V T R C A S S L L M A I M S V D

991 CGT TAC CTT GCG GTG ATT GGG GGG CAT AGG ATT CAC CCT TTC CGC ACT AGG ACC TGC TCC CTG GGG GCT TGC TGC GCC ATT TGG CTG CTC
GCA ATG GAA CGC CAC TAA CCC CCC GTA TCC TAA GTG GGA AAG GCG TGA TCC TGG ACG AGG GAC CCC CGA ACG ACG CGG TAA ACC GAC GAG
R Y L A V I G G H R I H P F R T R T C S L G A C C A I W L L

1081 TCC CTG GTA ACT GGG ATC CCC CCT CTG GTT TTC CGG AAC CTG GTG GAG AAC GTG TGC GTG GAA TCC TCA GAA TCG AGA CTC AGC GTG GGC
AGG GAC CAT TGA CCC TAG GGG GGA GAC CAA AAG GCC TTG GAC CAC CTC TTG CAC ACG CAC CTT AGG AGT CTT AGC TCT GAG TCG CAC CCG
S L V T G I P P L V F R N L V E N V C V E S S E S R L S V G

1171 ATC AAG TTG GCC ACT TTG TTC TTG ACC TTC GTC CTT CCA CTT ACC GTG GTG GCG TTT TGC TAC AGC TGC ATG GCT AAG CAC CTC TGG AAT
TAG TTC AAC CGG TGA AAC AAG AAC TGG AAG CAG GAA GGT GAA TGG CAC CAC CGC AAA ACG ATG TCG ACG TAC CGA TTC GTG GAG ACC TTA
I K L A T L F L T F V L P L T V V A F C Y S C M A K H L W N

1261 TAT TTT GGG GGC CCT CAG AAT GTT GCT GGG GGC AAA GCG AAG CCT CGG CGT AGG CAC AGC TGG CTA CGT ATA GTC TCG TGT GTT GTG GCA
ATA AAA CCC CCG GGA GTC TTA CAA CGA CCC CCG TTT CGC TTC GGA GCC GCA TCC GTG TCG ACC GAT GCA TAT CAG AGC ACA CAA CAC CGT
Y F G G P Q N V A G G K A K P R R R H S W L R I V S C V V A

1351 GCG TAT TCA TTG TCT TGG TTG CCC TAT AAT ACC CTC AGC ACT GTA AGA CTG GTG GCC GAG CTT GGA GAA GGC TTT CCC TGC CAC ACC ACG
CGC ATA AGT AAC AGA ACC AAC GGG ATA TTA TGG GAG TCG TGA CAT TCT GAC CAC CGG CTC GAA CCT CTT CCG AAA GGG ACG GTG TGG TGC
A Y S L S W L P Y N T L S T V R L V A E L G E G F P C H T T

1441 GTC GCC ATT AAC CAA GCG CTC AGT GCC ACA GCC GCC ATT GCC TTT GCC AAC AGC TGC ACC AAC CCC CTC ATC TAC TCC ATC CTC GAT GCC
CAG CGG TAA TTG GTT CGC GAG TCA CGG TGT CGG CGG TAA CGG AAA CGG TTG TCG ACG TGG TTG GGG GAG TAG ATG AGG TAG GAG CTA CGG
V A I N Q A L S A T A A I A F A N S C T N P L I Y S I L D A

1531 GGC TTT CGG CAC AGG ACA AAG CAG ACC TTG CCC AGC ATT TTC CCC ATA TGC CAA GCA GTT TTC CAT CTC CCC TTC CGT GGT GGG AGC CAA
CCG AAA GCC GTG TCC TGT TTC GTC TGG AAC GGG TCG TAA AAG GGG TAT ACG GTT CGT CAA AAG GTA GAG GGG AAG GCA CCA CCC TCG GTT
G F R H R T K Q T L P S I F P I C Q A V F H L P F R G G S Q

1621 CAG TCT ACA GTT GCC AGC ACC GAG AGT AGC AGC TCC TAC ACA CGC AGC TAC TGA
GTC AGA TGT CAA CGG TCG TGG CTC TCA TCG TCG AGG ATG TGT GCG TCG ATG ACT
Q S T V A S T E S S S S Y T R S Y *

**Fig. S2.** The nucleotide sequence and amino acid sequence of Xt-GPR25 constructs expressed in HEK293T cells. The amino acid sequence of Xt-GPR25 is shown in red, that of LgBiT in blue, and the signal peptide of sLgBiT shaded.

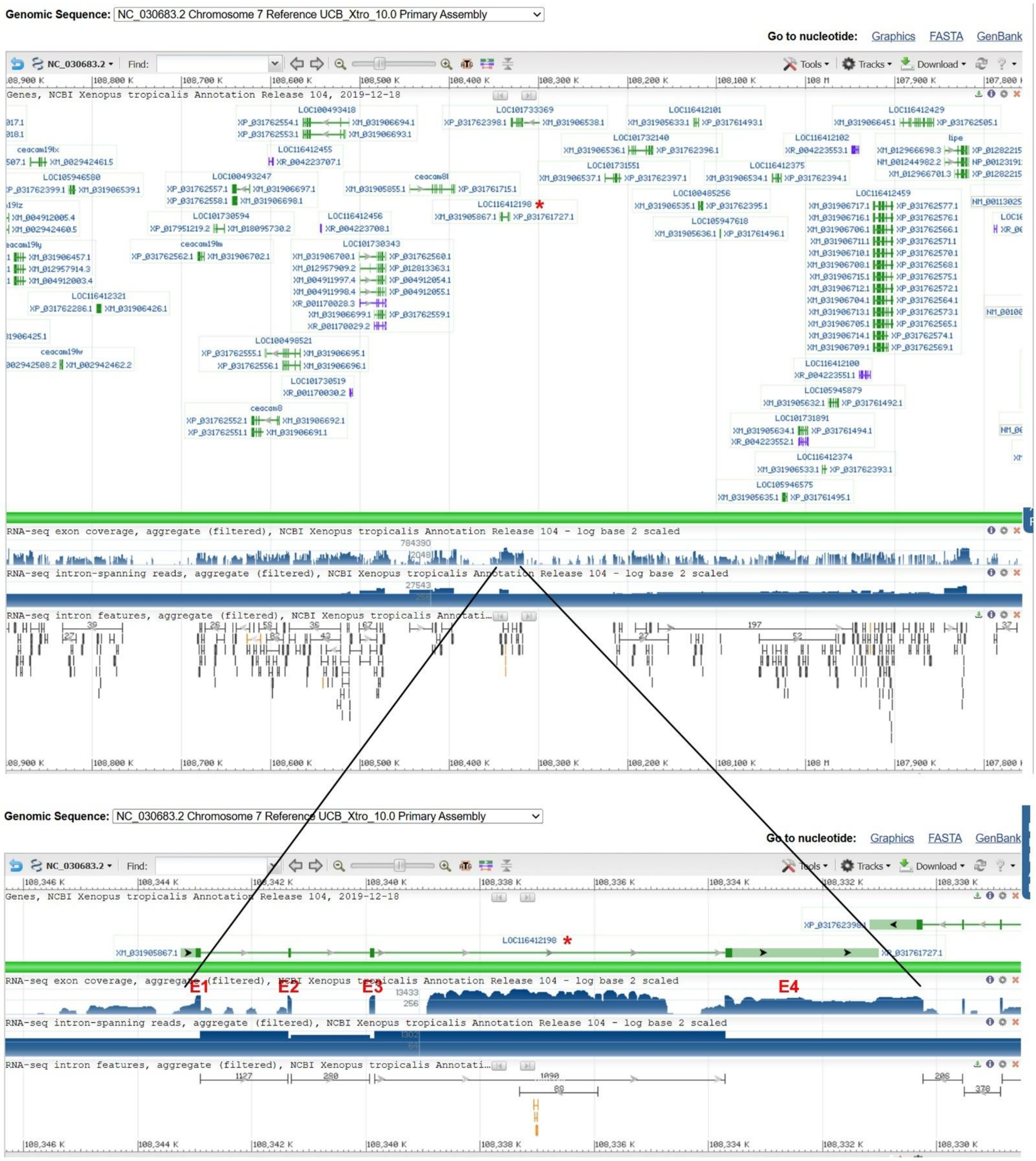


**Fig. S3.** Gene position and architecture of the possible *cxcl17* gene (LOC116412198) from *Xenopus tropicalis* (tropical clawed frog). The possible amphibian *cxcl17* gene is indicated by a red asterisk. The information was downloaded from the NCBI gene database.

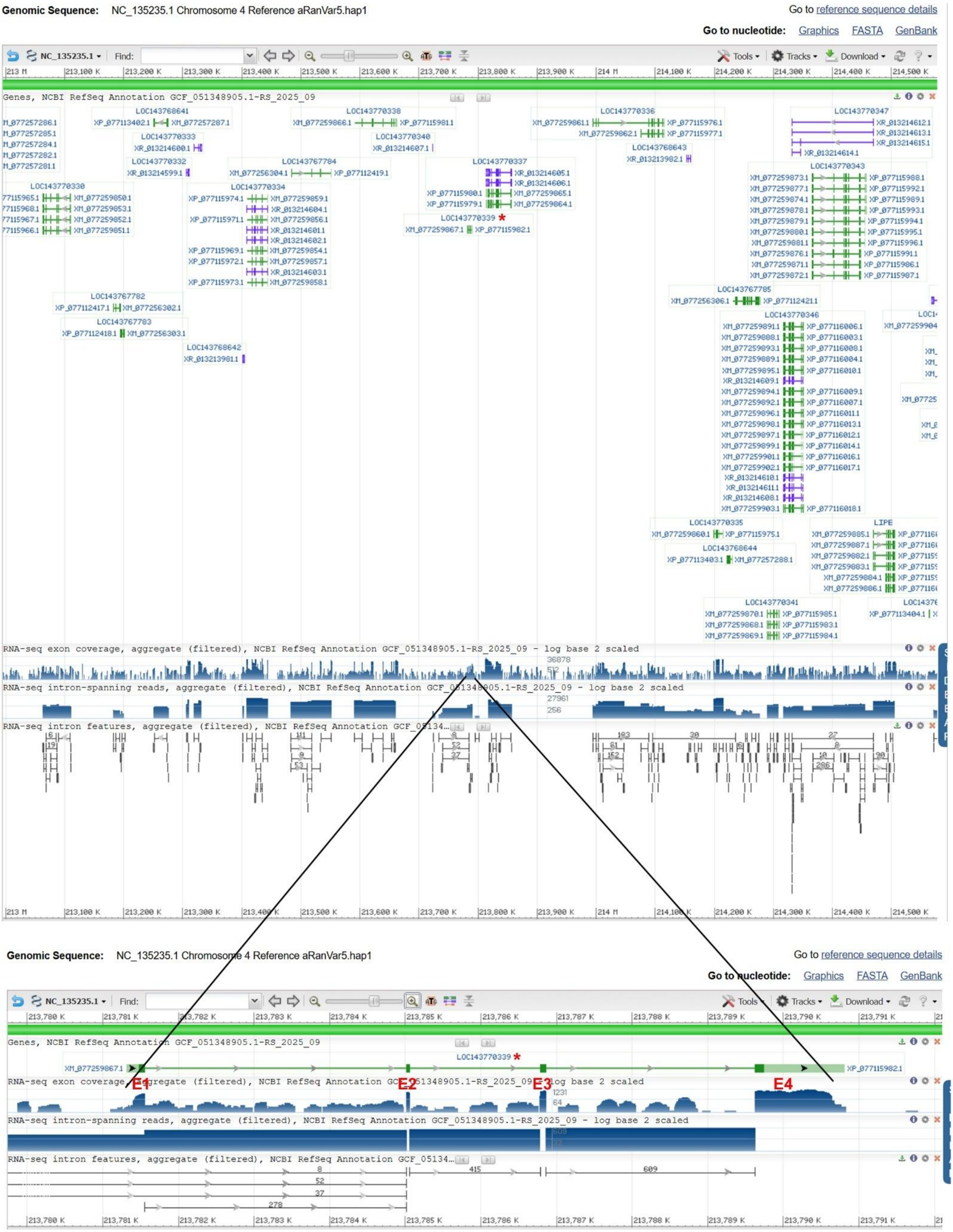


**Fig. S4.** Gene position and architecture of the possible *cxcl17* gene (LOC143770339) from *Ranitomeya variabilis* (Zimmermann's poison frog). The possible amphibian *cxcl17* gene is indicated by a red asterisk. The information was downloaded from the NCBI gene database.

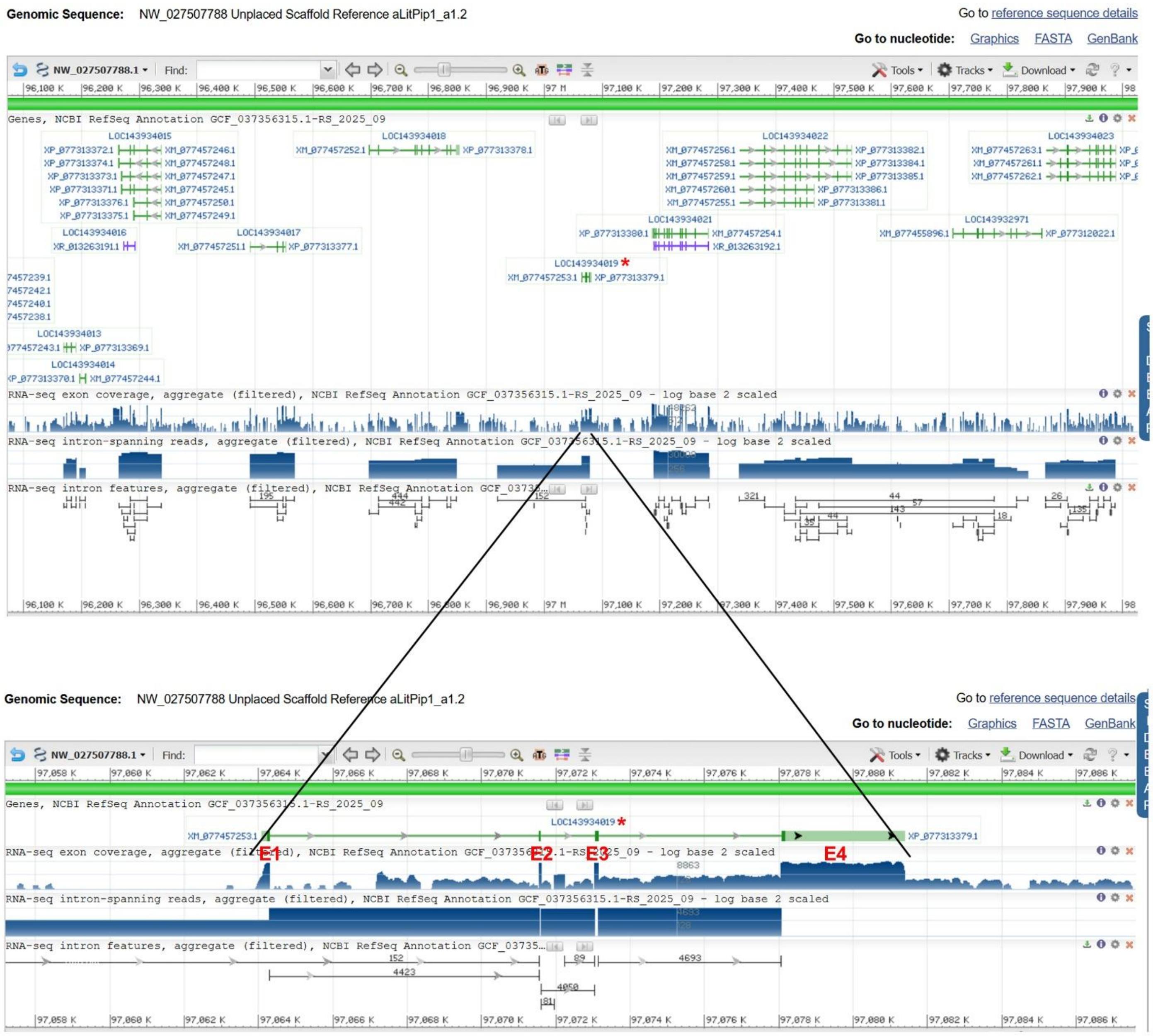


**Fig. S5.** Gene position and architecture of the possible *cxcl17* gene (LOC143934019) from *Lithobates pipiens* (northern leopard frog). The possible amphibian *cxcl17* gene is indicated by a red asterisk. The information was downloaded from the NCBI gene database.

**A:** Gene position and architecture

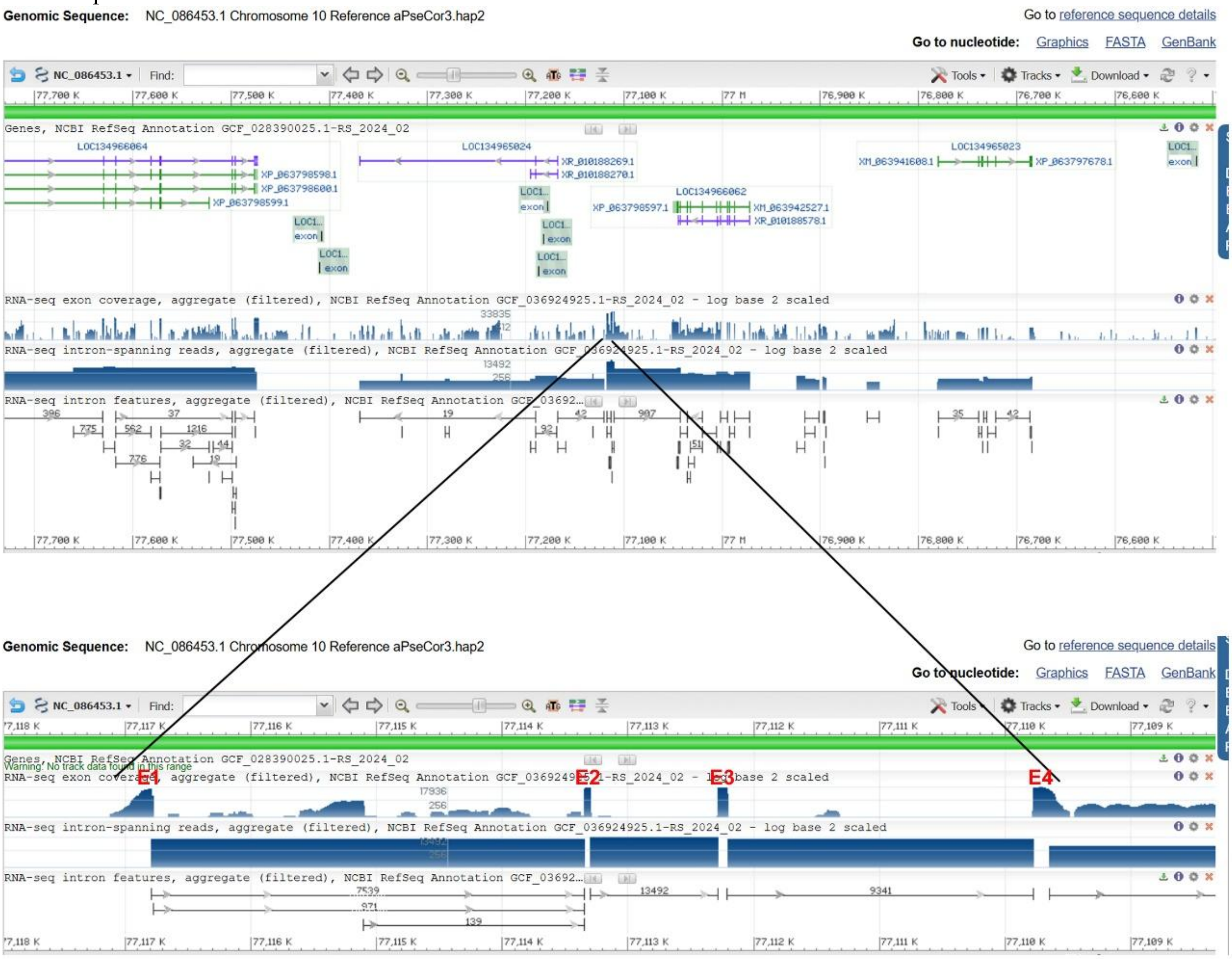


### **B:** Sequence of genomic DNA (exons in red and highlighted in yellow, coding region underlined)

atatctttgtagtgagagggttaagactaggggatacttgacattccttcaaagtaaaacctgttgcacggcccttccctgtgtgtatctcgagccaatgtgagtgcacggaaggtgtgggtgacagacaggtaagggggtagagcttctcctctaacagaagcctc
tctgaggatgaaggtggtcacggttttcctgctatccgcagccctgctggccctgtgctactgctggaccgctcatggccataagagtgcaggtaattcagtacctaaaggtaacagtcctgtaggataacgttttcaaagtaacattttcttatattctgtgtcacaag
caatggtctaaaatgtgctcctcagtggccaagtatagagctacctctttactattcctgatgaatgggttcgttgctctttcacgtattaaaacttagacaagggctacactagagacctgtgtctgggactgcatgcaacatgtcacatctgtattttcaatgcgccttc
aggggaacggtttgttttctctattttataccattctctgttgtagttcaacaaatcacatctcttattaaagtcagtttacataattcatcagacacttgttggaaagtaacagtcaccagtgcagctggctccatctgaaaacagatcacgtcacttggtcgccatttcttgc
agaattagtgtaggttgaccccttattcccttagggtcaaagcaaatagaaatttgaagtcaaaatgcagcaaaaaaggcaagtaaagtgcttgcgtgcataaaacggggcactgagacaagggacgaggatgtaatcctgccgctgtacaaatcattggtacgtc
cgcatctgaaatattgtgttcaattttgggcaccgtattataaaatagatatcggtgaactagaaagagttcaaaggcgagctactaaattgattaaaggaatagagacactggaatatgaggaaaggcttactaggttaaatatgtatacactagaaaagaggcgtct
aagaggagacaatattaatatcttcaaatatataaagggtcattatttattatttatccattatttatatggtgccacaagggttctgcagcgcccaattacagagtgcataaacaaataaccaaaacaggaaaacagcaacttacagttgacaatataggacaacttca
ggatgaataaacatagctacatcagcagatgacactggaataagtatcaggtggcagaagactgctggatttggtgcagttgaagattattaaagtaagaaaaatgataagcacatgagggaagagggccctgctcgtgagagcttacattctaaagggggaggg
gcagacagacaagggtgacacagacggggtatatagacagcgtacaacagagggtcaggatgagatttggctgggtttggtgaagaagttggtcttgagagcctgtttgaagttttgtagagaggtggagagtctgagggggagaggtaggtaattccagag
aaatggagcagcacgtgaaaaatactggaggtaggagtgggaggaagtaatcagtagtcaggagaggcggcgtgcattagcagagcgaagaggacgggtgggagtgtaaagggagataaggtcagagatgtagatgggagaggagtgggtgagggct
ttgtaagtgaatgtgagaaacttgaaatggattctgaaagggaaggggagccagtgaaggtctttggtgaggagaggtggacgtagtgcgtttggtgaggaaaatgagccgggtagcaacattgaggatattattacaaggagttagcagggggatttgtttatta
aaagaaccctgtatcggacacgtgggcactcgctgaggctagaggagagaaaacgtaggaagcggttcttcacagtaaggtcaataaagatttggaactccctaccggagaaggtaataatggcagactctgtaaatgcatttaaaaacggatcggacaaattt
ctaactggaaaaagtatccaggtttatagcatttaacatagtgacattaatatctgggagtggtaagaatcatagttgtcaattggtactaaatcattactccagcaggtgcattataatatgaacagcttaacacagaatacaggttgaacccgatgggcattttgcctc
ttttcaacctcactatgtaactatgactaggatgcctttcactgggaaccactggtgtggaatatagatgcctgccatttccttgtactccaacccaattctccttaaacattatttgcttgcagccgctgaatgatagatttgtgagttctgtacctgcatcagaccttttttact
accgtgtgtaaataatgtcaggcatcagactactggtgcataagggcatattccagtgaatctagttgcattgcccacttctaatgactacattactgaacatattttgcttctatccttttgctccacggtgcacactgatggcctgcaggctgagcagcacttgaaaaa
atagatttataccgtatttttactgtctttactttctcagtgaatgaataaaaagttataagaggtgcagggataacagccagtaacaatgcagcccaagtagtgccatgacattgggtagttatcgccagcatctgattggctgcttgcagtgaccaattagaagatagc
gatcaaagcatgagcagcactactattggggtggagtggtggggacagtggcaccagccagagcgtatgacatacagttctacagggcctgcaagtactggtggcgtattactgagcagtattgttactgctggcaaatcagtgggcattgggggtcattccga
gttgatcgcagccagcaactttttgctgctggtgcgatcaactaatccccgcctatgggagagtgtattttagcttagtagggctgtgttcgcttgtgcagccctgcttagctaaaaatgttttgtgctgttgaagagtaaggctggatctacttaccctgtgcgacggat
ccagcaacgttcctctcggctttgacatcagacatccgccctccgtttgtctggacacgcctacattcggacaaccactccttgaaaacggcatccaacggtgagtatccgccccagaacgcctctttcctgtcaatcttcttgcgatcggcgctgcgtcttctttttttct
ctatgcacgctgttgcccggcgacatcccgtttgcaccgcaacaaagaaccactgcgtgcggacgggtcggaatgacccccaatatgactgcagccataattttattgttctttacatgtattattggccattcattttcctgctggtatattgctgaaaacagttataca
gtaactgctgagactttatttggcttctttctgataggctgattgttcggtgttgtgttataatttatctatatatcacattcaattgaattctgattgctaaattgatcagtatctaatcattaaagcattgtcaactttctctctctccttaacctgagcatgttttacagttacagaa
acaggacagagagaatgtttcatggttgctaccctgatacctggcactggtgtgccttaggttggctcattatttgcatatccctagcttgtgctatgtttagtagaaactcatgcacacttctgtgtattccacctataaaccaatcttgtcttttagggacaagggatgac
agcagaggcttaaccaagaggattcatcatagaggtgagcagagataaaattaaacacaagaccagtgcgcatgaattatggaaaactgaaaacacaaaaaaacgctcaccatcatacattgtctagctgtctttatatgtattatatatatatatatgggtgggagga
gggattgcatgtccatgcagtaaaacccattgttgtttcaccacgattatgtacagtacatccaggcacattccatatctcattagccataaataattgtgcaaaccctggcatcctacagtaattgcacagacctcattgctaaatatattcatagacagattacaacaat
ttttgtgacacggtttccaatttacgaggggacaaggcatagtagggaagcagtgagttcagacagtaattaactgaaatcattggagtgtggtcaatacctaagtaaatagtgggggtggtattcaggagaccaacggtcacatgacctcccccaacatcctgcc
gcctcactatcccgacggtcggcatgccgaccaatagggactatttccactcttggatgtccacgacacccatacagtaggaatagagccggcagcgtggagagtgcagcgagcccgcaagaggcttgctgcattcgcccccccacccccgctggaatcccg
gcatcagtatactgaccggtggtcaggagatcgtcggtcagccatacctcatccaatagaggcatgggggagtaacagagtacaggatgagacaactgtagtgtaactgagtatgagtagatggctacagaaagaagcatcattactcacagaaataatgcaa
aactgagaataacataaatcctgataagtctgttcttcctccatctagtcccctgcagagtgctgaaaaagtaaacagtcttaggataattgccatacataggtcattttaatctgctattaaatgaattcatgccattaatcaccatatgtccattactggcggaggagttg
ttgcttcatttgatgtacttttatttattgcatttcctcattcagattctcttggatccgtacagcctgggaggcaacacgcactttgtgtctgtgacattgatcggcatcacttacagagtaagtcatagataagattaaacatcctggcagtcatcctgtatgttttccctcca
gcttgtccggaatcatggacagttttaggggcctacttaaaatgaatcccaatttttggcccatttatgatcatttattattggcactattcctagcgataattgatcaattaccagagctaggcctacctaaagctaggtacacacttgttcgatcccggctgatcaggtaa
aatgctcccgatcagccgggctctgtgtgtatctagctgtgacccctgtgacctcatgtgaggctccactatggggatcgggaagtcacatgtgcagcacataggatgcgacctcctggtcctgaccattacaggagcgttataagctgtttatcggctaacgctcc
tgcatcggtacagaatacacacctatataaacatggccattaacccaatatcgggagaacggcccgacaattgtataagtgtgtatccagcttaactgtaagctctttctggtacaacacattcaataagaggctgtcctggtgaagattttacagtatatatgatttcta
cctacttcgtgatgggtgctcatgcgcatggaaccattaaacacaggaggtagggaagatagtgcagatgggtgaaacaaataggtaaaccccataatgccatctcccattcccatttcatatggaactaagaggagggcaatgcttttgagactgcagtagaga

tctgtgacacattgggtaaaatatgcgggcggtcaagatcccgacagcagaatcgcgatggtcacaatgctcaccaaccccttttgtagcctaccccgtaaccctccccttcagcagcctaaccataaacctgccctctagtgcctaaccctaaaactgaccctaat actcaccgtcggtatccatcggcattgtgactgacagttattcaatcatcaggatcctgactagatcccattatctccagtattattgcataacaaatatgcccattacttaatgtttccataaccctatgaaaatgccagcaggatttattcagatgtgtcttcgttaattgg agtatctacaatgagtgctatgtgtgtggtgcacatgggcctttttgggttcagtaggcccacgctgcacccattttttcagtactcacctatccggagtacgtgtttgcagcggcagttctttagaagctcccggtaaatggtgcggtggccattttccaggcattttgca cgtgttcattgggggaaattgctgtgaaaatggccaccgtatcattttcccagtgctctgcacatgaatagtagactccagcatagtgctagagtctactagcaactgccagcaagagagagagaggagggggggccagatggagcctgccaggggtgttttaataga ggagagggcccgtgtgcagactccgagtgggccccctcctctctatggcgcgtaggctccagccttgtgccagagtctactgcgcatgcgcaggtctctggaaacatggcgcctgccatgatccagagataaattttgtactgcgcatgcgtggtggccattttg ggtgtgatttttttgctttggcacccatcccgggactccggagagttaagtattaaaacgggtgcagtgtgggccccccatggacccagtggcccatttgcattgcacacattgcatctatgatagaaacgtcaatggagcctgctcactggcccccttctctcctaat atgcccctggtccttatacactcttgtcacagccattccacacagacattcttgggactccaggtcccatgtgactcagcttgcacgagattcattttactcaaatgtgcatcttagtcacaaaaatgaagcaattagaagccacaacactgcttaatgagactgcactg agggcaccacacatctgtagcttaagcttctttcttaaatatgctctatagtcactgatggccaggctgaactttgcaaatgatgacccctgtgttatttctatcatgaggtccctatcccagaggaggggatgtggttattggtgctcacattgtctcctctcttccttcca cagcaccatccatcttgttactcataatagatgagttcttctctggttccctctcatctccccttctgtggttctatttcactgggacatactgtacactaccattttttttatataccaccagttattattcagagatgtagaggctacctggaaataacccatgcaatcacaa agagaacatacaaactacacatatgtatctatcacctagactattatttagtagtagactgatttaatgtacatggtccatctactctgtgcttcatcttatcttatctgtgtcattgcttgcagcttttcacattggatcatagtatgagctaattttctatatttcacagggaaaa agcgccacatgaatggacgccgcaaacctggcattcgtagtcggcattgctgcagacgacacttcaagcaaaatacccagccagacgtcatcattcctctgttctgaagttatgactcaatacaagtaaattcctgaaaatccacattatttctaataaacctcattgc agcttcaaatactgtaatagaaaaaaaaatcaacttctaatacagtctttttatggttctatatggcctttctaatatgtcttaatgttttcaaggcaggaatacagaagacaaaataggatggggcggggggggggggtgcaataaaa

**C:** Sequence of cDNA (coding region in red and highlighted in yellow)

Gcacggaaggtgtgggtgacagacaggtaagggggtagagcttctcctctaacagaagcctctctgaggatgaaggtggtcacggttttcctgctatccgcagccctgctggccctgtgctactgctggaccgctcatggccataagagtgcagggacaagg gatgacagcagaggcttaaccaagaggattcatcatagagattctcttggatccgtacagcctgggaggcaacacgcactttgtgtctgtgacattgatcggcatcacttacagaggaaaaagcgccacatgaatggacgccgcaaacctggcattcgtagtcg gcattgctgcagacgacacttcaagcaaaatacccagccagacgtcatcattcctctgttctgaagttatgactcaatacaagtaaattcctgaaaatccacattatttctaataaacctcattgcagcttcaaatactgtaata

**D:** Amino acid sequence (signal peptide shaded)

MKVVTVFLLSAALLALCYCWTAHGHKSAGTRDDSRGLTKRIHHRDSLGSVQPGRQHALCVCDIDRHHLQRKKRHMNGRRKPGIRSRHCCRRHFKQNTQP DVIIPLF

**Fig. S6.** Gene position and architecture (**A**), genomic DNA sequence (**B**), cDNA sequence (**C**), and encoded protein sequence (**D**) of the unannotated *cxcl17* gene in the NCBI reference genome of *Pseudophryne corroboree* (corroboree frog). The information was downloaded from the NCBI gene database (https://www.ncbi.nlm.nih.gov/gene).

**A:** Gene position and architecture

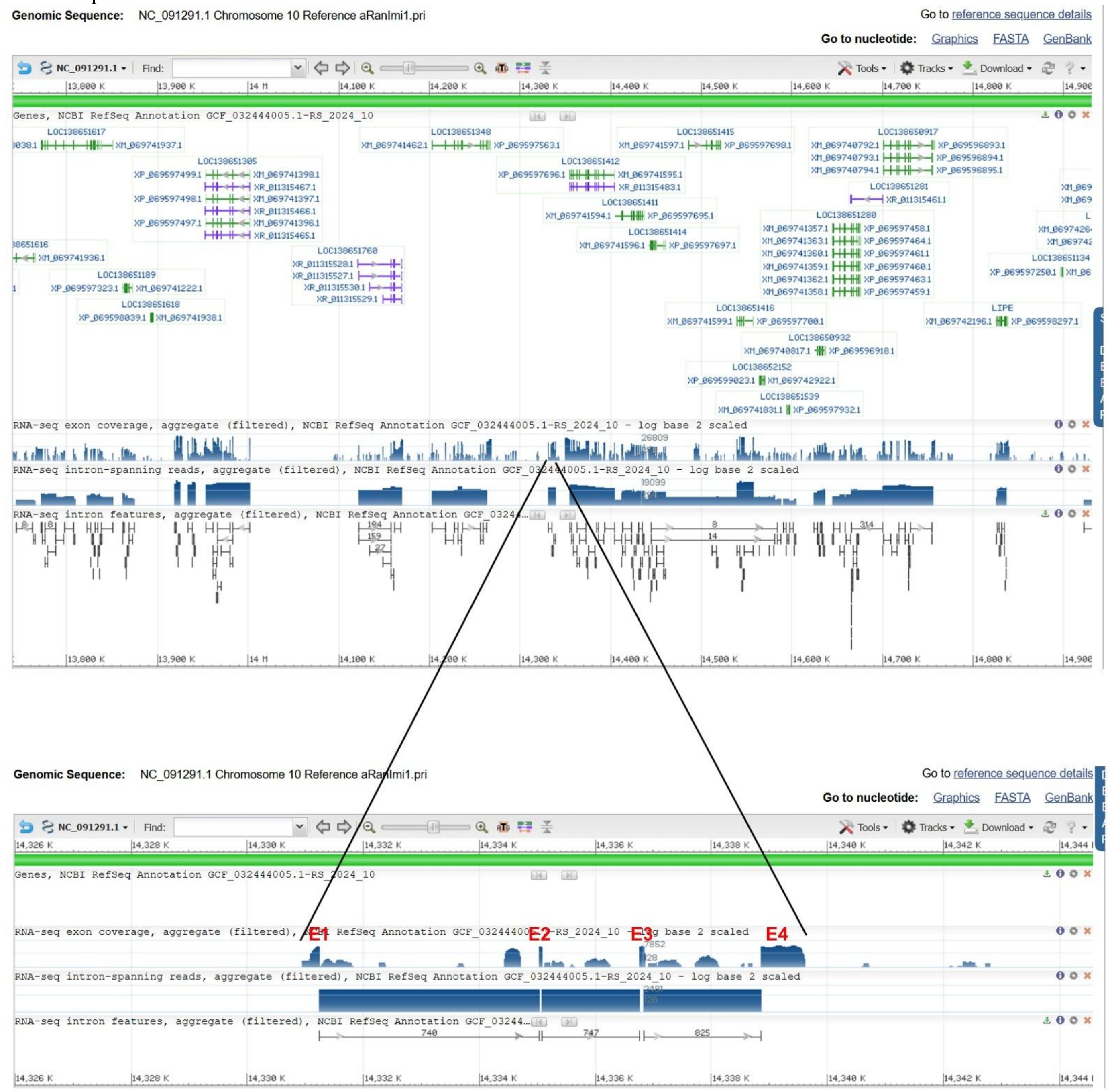


**B:** Sequence of genomic DNA (exons in red and highlighted in yellow, coding region underlined)

tatcacacctgatattggaaaccatagatccctgtaatgagagggttaagtctgggggacattttgacataacatcaaactcagacctggttacagccctaccctttatttacttggagccacggtgagaaggacggcggcagctgtgagtggcagaaaacaaaga
gatagagcgtccgctgtgagaggagccaacgtgaagatgaaggtggccacgatcttcatgttgtccgctgttctgatcctaatttactgcttgaccacacaaggccatcacattccaggtaactttaactacgaagatatattaaatccagattgacctctataggatc
tatgggactctgacttattgaattctatgctatcaatgaatggaaccttttgggttctatgggatatatgtaactctatgggattctttgggacacaatggcactctatgggacaccatggcactctatggaactctatgggacaccatggcactctatggaactctatggg
acaccatagcactctttggaactctatgggacactatgggacactatggcactctatggaacactatgggacactatggaacactatcggacactatgggacaccatggggctttatggaacaccattgaaccctgtgggactctatgaaacacaatgggacacta
tggcactctatggggacactatggcactctatgggacactatggcactctaaggaacactgtcggacactataggacaccatggggctttatggaatactactgaaccctatggggttctatgaaacactatgagcctctatgggacactatgaaacactatggga
cccataagactctatagtatctatgagacctatttgactctagcagatctattagaccaatgtgataatatggcacacaatgggactccttggcacatatgcatgtctaatagacactaggagacttcatagaatgtgcgggatcctaagggacctatgtgacctatcca
tgggatctatgtatctttattcaaaattatgagaccctataggatctatggaaccttatcggttctataggactctatagcatctacgagaactatgcgattctatgatatctgtaggactaatgtggctttatgtggcacactgtatctctgtgatatctgagggaccggtgc
aacactattggacattataagactccataggatctatgggaccctatggattctatatgactctatagtatctatgagacctatgtgactctgtaaactctataggtcctgtataatgattataggactctatcgtagctatggtaactatgtaattctatgagagctacaaga
cctatgtgactttatggtacactatctaattgtatagcatctatatgagacttatgtgactctatgagatccataggacctatgtgatattgttgtgcactatttaactttatagcatctacatgagacctatgcacctctgtgagatctatagggccagtgtgatactgtgaaa
cactaggggacactatgggactgtatgctatttataggacctatgagacaccactgggccctattggactccatgggatccatggagtctatggcttgctgtgagagctattggccctcaatgggtcaccatggaactccattgaacattattggattctatgcgatttt
cgtttttgcaggaagcaacagtgaacaattgcttctattcatgaaaggtcaggaattgcgataattcataagacagcatttagctcctcaatgcctttgttatgtaggaggtttacaactgtattataacaaaatagcagtaaatccatttgttatacctttacaagaataatc
cagaagtatcacgagtctcagttttgtccttgttgcatctagcaaccaatcagagctcagctttcattattctagagcagttacggaaattagagctgagccctgattagttgctagaggcaattaaagaagtctcactgttagacagcctgaggcctattaaagcaaac
aatatattttgccatttggacctttccctctctatatatacacccaaatatcgaaaaatgtttgtatcctaactatatacaggttcacaaacaaacctgtccgaaataggaaaatctctttaaattaggaataaatggtcataccccaggagagtcaaactatagggagttaa
gttatccaaaatctttgaatgatgtccaagatttgctccatgccccaatgagtagctccaaagtcatgaaaattggaaagtcccaacatgcctgtaaaaccaaacctcagcttcttgacgtgagttttcccagaacaaagaatgttccttgactatatcagacaatgtcct
cctccctaatcctggcccataatgcgtcagtagtcaacaaggcaagtgcacattccttgtaaaagagatatatttgcgctcggatcaacccccttgttagtgcgggtgctttagtgaaaaatatttgactaatgaaaagcggtcagggaaccttgacagcccgtaatga
tagtatagcaatatagcgtttggctgatggcaaagagttagaatcggactctgcctataccccgcgtagcagatgaatctgacatgcagtttgggagaataagggcacttgggttaaaaataaattgaaatacctgttccgcagcggtactgtacagatcactcagtt
cttcagtgccgctgtgagatccttccgttctttgtagtcggtgattctatgtttagataggaggactccgggcgctgctccggctggtggcaagcatcccccaggagtcttcacccgcgtggagcgggagtgtgatgccgctacagcaagtagccgctaggttcac
tgtaggtacaagaccttgcgtgacatcacggtcatgtgattgctcacgtgaatggctgtaggaagcagggtagaccaatgtcctacgcgtttcggaatagacatattccttcttcatggatggtcctctatacctttatagttagcggcaatcatgtggcctcctcacat
gttgacagcaaacagctctatctcgctgttgaggcccttagggattaaagtatctagtttgaaaatcattctactttcttcccttgacagttgctggatatagctgcccgttctcaagctcttttgaactattttcaggcctgtaatgatggtaccttttgaagaacctcagtga
tgaatcaaaaaagtgtacattccttggtaacccaagagagttatagtactggaagtgcagtaagagttatcccatctggacaactgaagtccacatggtatactagggtatgtgtccactgaagagtctgaagagcagtgtccacgctaccggactcagcatgacc

atttgatggcaagtgatgtaacactacatttcgaaaagccaaccacagcttccaccgacattgtcaacatgggcagcacggtggcgcagtggttagcactacagccttgcagcgctggggtcctgggttctaatcccacccaggacaacatctgcaaagagtttg
tatgttctctccgtgtttgcgtgggtttcctccgggcgctccggtttcctcccacattccaaagacatactgataggaattctagattgtgagcccaatcggggacagtgatggtaatgtgttcaaactgtaaagcgctgcggaatttgttagcgctatataaaaataaa
gattattattattattattataactgataactaaggcagcaatgatcagctgattgaagacgcagttgactattgaggtcaggttgtccttaaattatgcactccctggtggtccttgtcagttctggacatcagtatctcatctgatctagcgtccttggaggagaacacta
atgggatcagctattagtgtactgacctcaccattaattgtcaccaccatgtcaggaaccctacctatgtttctaggactaaaaggagatcctactatcacagaagacaaatatagagttattcatgtacaaataaaaatgttttctattttagaaaaacgacacgggaag
tccggcgtagccagatggacccaacatggaggtatgtgaatcagtgtcaatggaatgtgtcaattgtctaaacatatttttgacctgttgtaaatgccgccgttcttctgagtctggaattctttttctttagttcctgcacctctccatttctgaaatatgccaaactctatcc
tatatataagtctagttttcttatccaaccgggtgtggtcatcaagaagacgcctgtagggaacagttgaggactaagccccaggggcctaactatattggtttcaatactattaaagtgcagtcatagtcgggggtcttgctggagattttgcattgaagtttcaccact
tccacacccacttgcctaacaaaactagatttacttgcgggaagagtgagccatatctcaggaatggcgaggcacaggaacaaaagaaaaacattgacggattaagaagaacaaaggtgtgacgcgagtcactatagaaaatattagcaccagaagtggaga
ggactacaatgacagatagcaaaaatatacactaattagagaataagtaaaggtaggacaaacaattcgaggttgggtccaggaaatggcgcagtacaacaaggggaaatcagagacaaaatccacttcacgagccgccgaggtcaggaaaaccagaaag
acaagacagtttacagggagcaggcacaagagaagtcaggtcacaagcaaaaagtcaaagtccagaaagggaataaagtcaggctgggtcaggcacggagcacagagggatcagtcttacgcaggcagcagcaagtgtacgactgacaccggatggaa
gagctagcatgcttaagtagaagactaatcaagcaaagtgaacacagatgattacagtacagacccaactcaaccaaagcagactgacagcctctacacaaagtagcatatcagacaggatatcgtgtggagatcatgacaaaaggcatttgcaccagataaa
actgacacatttatggcaagtgataggtcctctttaaaggagtttttccaaaatagtaagctataagtgatcataagtgatctggtctaaatggagtgaaggccaagtttatactgtctattcattgactatggtgctgccaatggtactggactttctccggcaatccaata
gacaataaattttgcagaaattttgctggactctacagaatctgtttttggatcaaagggtgtttccagaagtcagacccccagctattagtaagttatcctctatcccaatagtaagggataagttagtgtccttggaaatcccctttaagtacaacagagtgtgatcggt
gggaatgaatggatgtactgttattctgccacaattattatttgtattatgtttattattaatactattgccaatgttcagcgcatatgagagttcccagaatcctcgcccccggctatgtccttctgtgcataagtaaatattcactgccagacatctatggcccaaggtgaat
gcagcatctactgcttctgtctgttatagtgtcttcatcagagatgctgtggtctaaggaggtaaggtttctccttgtggagtgagacatgcctgacatgccagtgtaaggagacttgtaggccatgcactgctatgatgtgatgttcttatactttgctttttaaacaattga
tataaaattgtatattgatgccttcaatctttttagaatctcctggatctctccaaggtagaaagaaaaccatcatctgtatttgtaaccataagaacgggcatattctttcaagtaaggacagaactaacacttttaggttcgattcgttattgcatttgcgcctttttttgtttagt
cttaggtgtttttcgtctatttttcatttgctccttattcttctatttattactattttatatgtagtcgcctagattttttttttaccttcaccatttgcggcctccagatgttttcgactaattcatcaattgcgacttttctaaaaaagtcactcaaattttcttgcagatcaactccagtcc
ttctctttggtgttttgaatgtaaaaaccagaatttttcttcttttttgcttaaattttgtgacgttttcgttgtccaaagggtaatttttctccttgtggaaaaacctggcatgtcagagtaaggagactttataggccacgcatgctcctctgggaaattaaatatgcaaattgtc
tctacacacttggaataaaagccaaactacaacttcatgtgacattttactagaacttatgactttttttagtgctaaaaacttgcagaaaaatataaagacaataataaaaagcatttttttgattttgcgcaaaaatcatgaaaacaaatatcgcaaaataaaacgaaaaa
gtgacttaaaatatttaatgtatggctgtcccctagtgtggctgctctatggttggctgcagattgaggagctggaccggtgaaaatgagctcatgaccaagtggccttaaaggggatttcggtcaatccatttaggctggtttcacacttgcgtttcaaaacgcatgc
gtttttttgcggcaaaaaaaacaccgcaaaaaaataataataataataataataataataatttttatttatatagcgccaacatattccgcagcgctttacaaattatagaggggacttgtacatacaatagacattacagcataacagaaatacagttcaaaacagatatca
agaggagtgagggccctgctcgtaagcttacaaactatgaggaaaaggggagacacgagaggtggatggtaacaattgctatagttattcggaccagccatagtgtaaggcttgggtgttcatgtaaagctgcatgaaccagttaattaattttttttttttttttttttttttt
ttttttttaatataggccacacagggatcgttaggttaatgcattgaggcggtaggccagtctgaacaaatgagtttttagggcacgcttaaaactgtggggattggggattaatcgtattatcctaggtagtgcattccaaagaatcggcgcagcacgtgtaaagtctt
ggagacaggagtgggaggttctgattattgaggatgctaacctgaggtcatcagcggagcggagggcacgggtagggtggtagactgagaccagaaaaaatactacatgttgcatttctgcaacacaacgcatgcagagaaaaaacgtatgcgttgcaaaaa
cgcgtcaaaaaaagcatgcgtttttaatgttaaatatagggggaaaaaaacacatgcttttttttggcgtttttaccgcaaaaacgcaaaaaaaaaaaacgcaaatgtgaaaccagccttagataacctatcaataggataggtcatcaacacttgatacgtgggtgtcc
atcacccattacacccttaaatatgttgaaaacattgataggagctgttctgctgtacctggcagcggctgctacatattgaatggagctgcattctttacatccgattaccactggagttaatacttgatcaatggggttgtcaggtgtcggacctccacaaatttcatat
tattgaccaattttgtagcctccataagtagttaatcaattttggtgcaatttgtcctattttaagactcctctttttttttgttttttcaaactgtccagcaaggtgtaaaaagtgtctaaaacataaaagaaatctggcacatgacatttacgaagagcctttctgtaacttaata
aatttatctagtttcttcatttgatggtttttttttgtaatgatccacaggaaaaagacgccatatgagtgcacatcacaaaaatggtcatcatagccccggatgcgccaggatctgcatgcgtaaaggtagaagaggaggatccgtacgttccatacctttgtcttgag
caaattttgaaaaattttctgaagaacccaggagcgtaaaaatttagagtaactctttttttttttcttatcgagccaccccagtttacagtttatatacagatctatatactgtaattatataaagttaagtcactttgtagatagttgacatggcccatcttaatcagagttgcg
gcctatattatgctccagagctgcgctcacaattctccttcttagttatagctttgaacaaactggttatcacacatttttgttttatgttaaagcttagcttgcatcatttagtagggtagataatagttggccaacgagtcgattttggtcaaccgtataggggtttatgggtt
gtggggtcctcatgactgtcccctgcggtgaaggaaagattaaaggttcatagagtaaagtgaactcttctgtaaagccattttcttgccttgtgtgttacatttttgtagtgtaggtttatcgctctatctcacagtatttcttcatagtgttgtaaccaccacggtcatattgt
cacttgcacttcttgttttattacaccatagtgtcagcacggcttaaactttttgtgatattgtttatattaaaataaagtttttgcatttttgtacttatttatattatatgattatgatatcatgatgaagaaacttcatctttggtttttggcaacaaaactcaaagacttcttgaga
cagcagcatggctgacgatggctgctggagatccttggaagaaaaaaatctggtgggtcaaattcccatct

### C: Sequence of cDNA (coding region in red and highlighted in yellow)

Acggtgagaaggacggcggcagctgtgagtggcagaaaacaaagagatagagcgtccgctgtgagaggagccaacgtgaagatgaaggtggccacgatcttcatgttgtccgctgttctgatcctaatttactgcttgaccacacaaggccatcacattcca
gaaaaacgacacgggaagtccggcgtagccagatggacccaacatggagaatctcctggatctctccaaggtagaaagaaaaccatcatctgtatttgtaaccataagaacgggcatattctttcaagaaaaagacgccatatgagtgcacatcacaaaaatgg
tcatcatagccccggatgcgccaggatctgcatgcgtaaaggtagaagaggaggatccgtacgttccatacctttgtcttgagcaaattttgaaaaattttctgaagaacccaggagcgtaaaaatttagagtaactctttttttttttcttatcgagccaccccagttta
cagtttatatacagatctatatactgtaattatataaagttaagtcactttgtagatagttgacatggcccatcttaatcagagttgcggcctatattatgctccagagctgcgctcacaattctccttcttagttatagctttgaacaaactggttatcacacatttttgttttatgt
taaagcttagcttgcatcatttagtagggtagataatagttggccaacgagtcgattttggtcaaccgtataggggtttatgggttgtggggtcctcatgactgtcccctgcggtgaaggaaagattaaaggttcatagagtaaagtgaactcttctgtaaagccattttc
ttgccttgtgtgttacatttttgtagtgtaggtttatcgctctatctcacagtatttcttcatagtgttgtaaccaccacggtcatattgtcacttgcacttcttgttttattacaccatagtgtcagcacggcttaaactttttgtgatattgtttatattaaaataaagtttttgcatttt
ttgta

### D: Amino acid sequence (signal peptide shaded)

MKVATIFMLSAVLILIYCLTTQGHHIPEKRHGKSGVARWTQHGESPGSLQGRKKTIICICNHKNGHILSRKRRHMSAHHKNGHHSPGCARICMRKGRRG
GSVRSIPLS

**Fig. S7.** Gene position and architecture (**A**), genomic DNA sequence (**B**), cDNA sequence (**C**), and encoded protein sequence (**D**) of the unannotated *cxcl17* gene in the NCBI reference genome of *Ranitomeya imitator* (mimic poison frog). The information was downloaded from the NCBI gene database (https://www.ncbi.nlm.nih.gov/gene).

**A:** Gene position and architecture

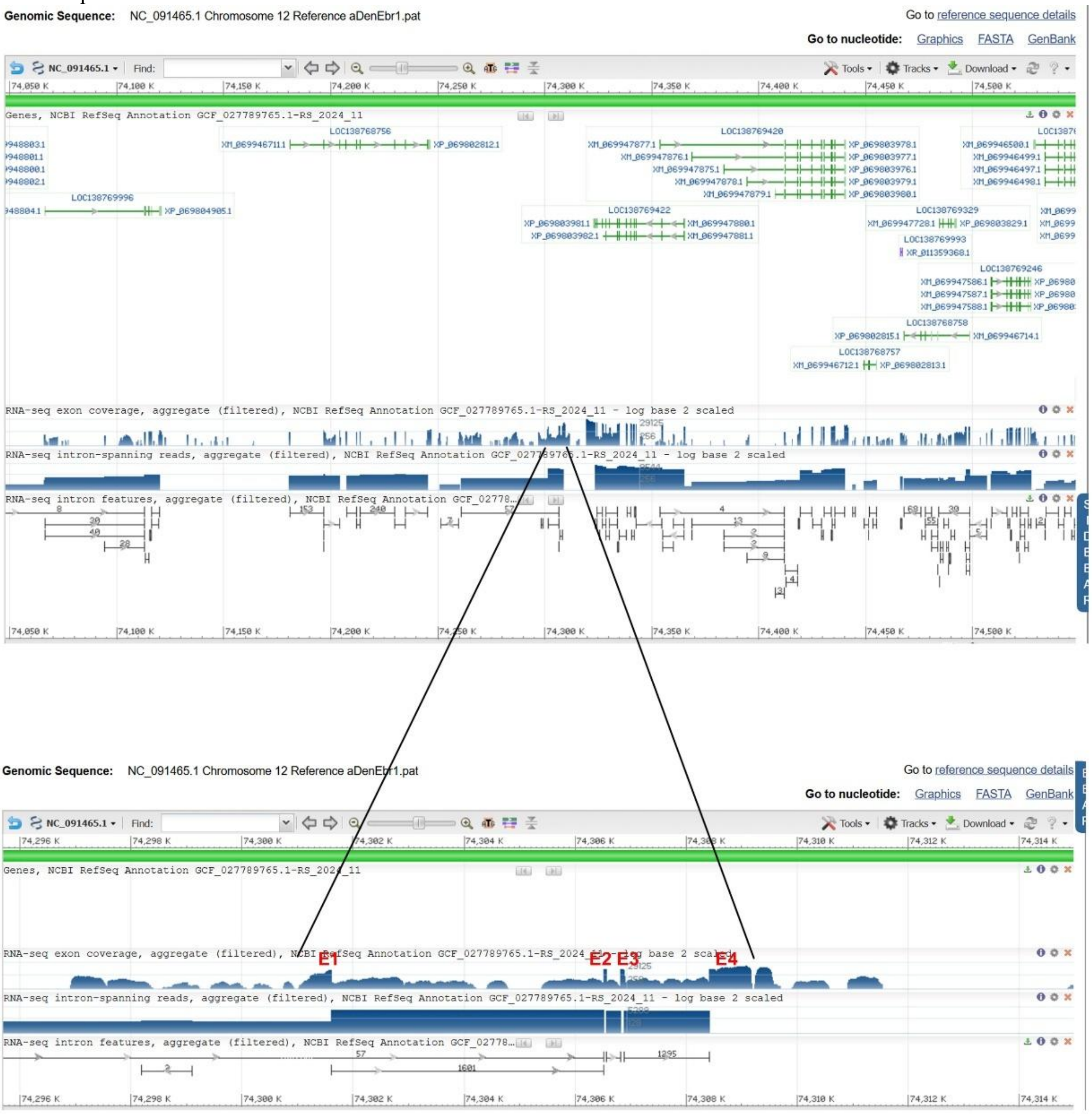


**B:** Sequence of genomic DNA (exons in red and highlighted in yellow, coding region underlined)

atcataatacaaagctctttaactcttcggagcttaacaggcagacggaaaacggattgttctttatatgctggtattagagatcgttgtgatggatcattgctgtattgagagggttaagtctggggaacacttgacattcttttaaactcaaacctgcttacagccctac
cctttatttacccgaagtcggcgtgagtttacggaaagtactggtgcctgacaacaaaaggctcaagcttctcctgtaacagaagccaaggtgaagatgaaggcagccacgattttcttgttctccgtagtcctgatcctatgctactgctggaccactcaaggccat
tacagtgcaggtaacttttaaccgggggtcatggttcaaggacgtatatttgacctatgggacctacatctttaggacccatggggttctatggcttctatgggatccttgaaactctataagataccatggtactccatgagtttttacgtcttcgacgatagactagaa
gttgttcctacatagaggccagcatagcgttcagcttttactttgtcttgtgactttttgtattacctttatatactaatatttacaatagctttatttaaacattgaagaaaatctactccttctatctctagcaaaactaagtatgttccaagtataacacagcaagtagcggcat
aactatcggctactcgcatacaattgtttctggcatgttctcctaaactggggttctttgatattaagcattttccataccgctcatgtacatgttactgcagatatagctgaaccgggctgcactaagtttatttggcatcacaattaaagggaattgaatgttgaaacaaa
actaagcaaagagtcagatctcggctgttggttccaaatgactttaagttatggctcctagatcttcctgagaacaaatggtcccaaagttatctgcccatatgatggacttattgtttggcccctctgtttctactgatcagttgtatgtctgctatgtgttgttttatgttttac
gagaagaacatttgtgcccatgacaacactccatttcagtgcttggcgtggataaatggacatcaaatagacccaaatcctcatccctttcaacgtccacctcctaattgctttcgttttagtgttagctgagtcaataaggatgtggttggatatttgaagcaatacaag
gcacttgtgaaagcaatgaacgtgactcagaggaggaagtcttatcctggagatacaagtactcaacctcatgggtgacatgttaagtcagttcctgataactccacacccttttacttctccatatgtggtcatcagggagggacagaacacttggcttcacatgtaa
aggaccaaattggatggagtttagtcatccaatttctcgtcaagtttgataaagtcctcaatcagtggcctcaagatcacaaaagtgccaacacctacagcctgctccaccctccgaaggtggaaattcccaagacttaggatatttcctgaccatatctgaccatatc
ctttcctccctcataatgctattcttgatagttagctgaacaaacatacatacattcctttgtgactttgtgtaattatatactagacctggaaattgatggtgtccatcataggcatgctttagatttatgtattggagactccccaaaatgtatttgtctaatggattctggtaaa
acagtgtgaacctagtctgaccaaaagagagatagataagatgtgatctatcccccccccttctccctgtacaatgaccattgcataggtcacataacatgcctagaaaactcaaccatagaactcaatagggtcctctgcagtctattgatcttttctccatggaac
cttttgtagaaacatacctctaaatgctgttctctaaaggtacatcaggcaagagatctgcccccataataatatacaaaacaatatataaaataaaataaaaaatacaatcagaaaatagaaacaaattagaaaacatttataaagaccagtttccgatgtggagaacat
ggcacaaccatgtgttacatcattacttatgtaaagtatctgaataaaagtcataacatatacacaacaggccaatcatagcaacccattgcaatgtcgaatgatcatggagttacctgtaggaccattgatgatcagctaataagaaatgcagttgactatttagatca
ggttgtcaagttaagaaaatccctttaaatttgccatccctggtgttctagaaaatgacatctagacttttgtttgtctattccagacaatgctatctatcatactcagtggaggaaaggcgcatttgtgcccatgacaacactccatttcagtgcttggcgttgataaatgg
acatcaaaaagacccaaatcatcatccctttcaatgtccacctcctaactgctttcgttttagtgttagctaagtcaattaggatgtggttggatatttgaatcaatacaaggcacttgtgaaagcaatgaacataactcagaggaggaagtcttatcctggagatacaag
tacagaacctcatgggtgacatgttaagtcagttcctgataactccacaccctttacttctccatatattgtcatcagggagtttagtcatccaatttctcgtcaagtttgataaagttctcaatcagtggcctcaagatcacaaaagtgccaacacctacagcctgctcca
ccctcctaaggtggaaattcccaagacttagggtcctattccattggccgatgggggcctgatcaacgatgtaaacgagcaccgatctgctagatcgacgctcgtttacagggcttattccatggctcgacaatcgtttagcgagggctgcaagggcattgttacc
gatgtccttgcagcccttgtttcatactttacctttccaggctgcaggccttctcccggtcccgcacgcagcagcttcggtgcagcctgactgagctgtcagaccgctcagccaatcactggccgcggcggtcccggccagtgtttggctgagtggtctgtcagttc
agacaggccgctccaaagctgctgcgcgggaccgggagaagacctgcgcctggacagataatgtatggtgcttgtgagatcgtcgggcacccatcgcgcatcgctattccacatagcgatgcgcgggtggcgaccgatgattttaggtttgaacctaaatgaa

cgatcagccgatgacaagatcagtggctgatcgttctctctattccacggagcgataatcagccgaatcaggccgattcggacaattatcgctccatggaataggccccttaggatatttcctgaccatatctgaccatattctttcctccctcataatgctattctcagt
agctgaacaaacatacatacattcctttgtgacttagtgtaatcatatattagacctataaatttaaagggagtgtgtcatcagaaaatcacaattgcttaaagcgactctgtacccacaatctgaccccccaaactgcctgtaccttcagatagctgcttttaatccaag
atctgtcctggggtccgttcggcaggtgatgcagttattgtcataaaaacaacttttaatccggcagcgctgtgtctaacggccggggcttacatttgtatatgcattaggctggcacaccctctctgtccttcctccccaccctcctcctcattaggaatgctccaggc
agattgcttcctattccccacctgtgtgtataatgaacatgggctggatcgttaatacgcctgtgcaaagctcaaacagcagtaaatgttcctggagcattcctaatgaggagggtggggaggaaggacggagaggtggtgccagcctaatgcacatacaatgta
atccccggccgttagacacagcgctgcaggattaaaagttgttttatgacaataactgcacccccctgtcgaacggaccccaggacagatcttggattaaaagcagctatctgaaggtacaagcagtttgggggggacagattgtgggtacagagtcgctttaaat
caagtttatgttaaacatttctttaagaagtttaccatttacaattttactatctatattataaaataatcctaaaattttgcaatttccattctggccactaagtctaataataagctgagcatgttaaagtattgtctggtaagctttatttaatacatagcaggtaccagcacag
gcgttaaaaggatccattaagccacatttacctgtgtaattttgacatttattctaaatgtatacactgtgctttgtcacatgaaatagtgcagactactgtctgagtcctaaaaaaatgtattccaagtgttgttgatgtcattttaacacatgttccaatgtatgacatatacc
agtatgtggtgtccatcttgggcgtgcattcgattcatttattgtagactccacaaaatgtatatgtctaatggatactggtaaaacagtgtggacctagtctgactatgaaaggtgacaccaaaagagagatagataagatgtgatgagttcatcccccacccccttct
ccctgcacaatgatcgtcgcataggtcacagaacatgcctagaaaactctaccacagaatagaatcctctgcagtctattgatcttttctccatagaaacatacctctaaatgctgttctctaaaggtagctcaggcaagaggtctgcccccataataataaacaaaac
aataaataaaataaaatacaatcagaaaatagaaacaaattagaaaacacttatggagaccagtttccgatgtggggaacatggcacaaccatgtgttacatcatttcttatgtatttgagagatgcggttgactatttagatcaggttgtcaagttaggaaaatcccttt
aaatttgccatctctggtgttctagagaatgaaacctagaccttggtttgtcaattccagacaatcctatctaccatacttagcggaggagaactctgtataggatcagccattactgtacgatcaccctcattcattgtcatagtaaacgatcacgtcaatgatcagtcct
cgcacagagtgttgggaactctaccattgttgttgtcactataactatgaggcgttcatatcctatcagcagatgaacatagtgccatagtgttgcacctttgttacaattacaaaactgttttgtcttttagggaaaagtcacggcaaattcatcgtagccaggcggact
gagcatggaggtatctgatcaatatgtgtactttatgcagtgggagtgactgtttgtcctgtaataaccattattaatataactcttataaagctttgtcccttgccccagttttatccccccttgcccagcacccagtgtacacaagccccattggcttccattccaagctata
gctccgagagcattagtagacatccattatcatagagttatgatccacgatatatgcagctactattgatgtggactttgatgctttctagatactccgggatccattcaagatgggaagaaatacttgatctgcgtctgtgacctgaagaaagccaataatctaagtaa
ggacaaaaatacatataaagagggcatgtagaatacagtgtaattccattattctggcaattacctccgaacagctgtctgcagatgggtagtcttatagagagatttctggctcggcatatgtccccagtcatgtcctattacatctagttactgcgaagcactgactt
catgggaagctgtctttaaatagtggtttgagagctactttgttatcctttttattttaaagaccacacgtagttacagtattatgagttacgttaacatttagttacggcatatggggtatacagtgttgagggtcctctctcagtaaccacacccatggccagtggcgtagc
taccatagaggcagggtaggcagttgctatggggcctgtgcaggaggggggccagggagagaaggtaagagcatgtgtccttctgcttaacccccttatgtactgcagtgtgtaagtgaaccaatgtttacttacagtacatgctgcaacacaaaaaagggttaac
aagcagaagacagagatctctgctttactgctctgataacctctgacctctgtatttcagctggcaattaagtaggaatgtaaaatctgcagaggtcagaggttatcagtgcaccagaggtaaagtatatatatatatatatatatatatatatatatatagtggcatggtg
gggctgtaagaggcagttctatgcctctggagtgggaattggagttcaggtggagctcctggtccagatactccactccagccttagagctaatgaggttctgaaatccacctggtggagctctattagagactccagagcttcccaggtgatggctcccaggtga
agactcccagggtgggagaacacagaagaagaagaagaagaagaagaagaagtctgggtgagaccagtggagctggtgatattgagcattaggctcataagtaacagctagggaagctgtggtgttagtcagtggctagacggcctaggttttattttaggc
agtgttgcctatgtcttgtgttatttgaacggataaataaagctgactgaggtcagtataaagcatacagcactgactggactgcaatttgtgatgtgccaaccgtctcaggcccaggagatggcgatcccagcaagtgagtaaccccccagattgtcactatatatat
acatgcagtgctttgtgttgtgcgtaggggagggggggcctttttgctatggggccccgtgaatcctacctacgcccctatccatggccaaatgcatagactgtatgtaatactccatttcacctgtagtggccgctgctgggtaaatgtatgacccagactcatgg
cgatcagctgatcaataagctagaattagtttgtcttgatgaggcaactccttaaagcatgccagcatatatattctgaaacctctgccttagctgtgtactgtataactttttaaataactttttttactgtatcctgtttggccacataatatatgactttttttaatcattcaca
ggaaaaaaacgtcatatgcgggcacaaggcaaacatggtcatcacgggcgcaggtgtgccaggatctgcatgcagaaaggtgcacacagaggatcgcaggtttccatacctttgacataagaaaatgttgactgaattgctgatgacaaacagaactcctgaa
aatgtttactatcatttctgtaatccacgaatataaaaatttacagtaactgtttttgtatagagccactccagaagacaggtgatacagtatataccaatactctttatacaaagctgactgactgtataactgttactttctcattacatcctgtatcatccttcagagctgcat
tcacaattctgctgcttgttgttgagaaccattaaaaaacgttgttgcatgctcttgtaagagctaagaatgcacaataccagatgtaaagatgatactttgtaaaagacaattgactttaagggattttagctctaaagccttattggccgatctcaccctgtgtgattggc
cgttgaacacaaggtcgttccagcccactgaaaaataccagactatatgatatatatctctctgtggaatgggaaatgtgtggatgatggctaatagcagagaagggccaccagtactcattgtgcattgtgaaggctctgtacatgagattggtgctggtttacata
caggatcgggccgactgatataaccctaacacctaaaactcagtgggggagatttatcaaacatggtgtaaagtgaaactggctcagttgcccctagcaaccaatcagattccagttttcattttccaaagagtctgtgaggaatgaaaggtgga

**C:** Sequence of cDNA (coding region in red and highlighted in yellow)

Agagggttaagtctggggaacacttgacattcttttaaactcaaacctgcttacagccctaccctttatttacccgaagtcggcgtgagtttacggaaagtactggtgcctgacaacaaaaggctcaagcttctcctgtaacagaagccaaggtgaagatgaaggc
agccacgattttcttgttctccgtagtcctgatcctatgctactgctggaccactcaaggccattacagtgcagggaaaagtcacggcaaattcatcgtagccaggcggactgagcatggagatactccgggatccattcaagatgggaagaaatacttgatctgc
gtctgtgacctgaagaaagccaataatctaagaaaaaaacgtcatatgcgggcacaaggcaaacatggtcatcacgggcgcaggtgtgccaggatctgcatgcagaaaggtgcacacagaggatcgcaggtttccatacctttgacataagaaaatgttgact
gaattgctgatgacaaacagaactcctgaaaatgtttactatcatttctgtaatccacgaatataaaaatttacagtaactgtttttgtatagagccactccagaagacaggtgatacagtatataccaatactctttatacaaagctgactgactgtataactgttactttct
cattacatcctgtatcatccttcagagctgcattcacaattctgctgcttgttgttgagaaccattaaaaaacgttgttgcatgctcttgtaagagctaagaatgcacaataccagatgtaaagatgatactttgtaaaagacaattgactttaagggattttagctctaaag
ccttattggccgatctcaccctgtgtgattggccgttgaacacaaggtcgttccagcccactgaaaaataccagactatatgatatatatctctctgtggaatgggaaatgtgtggatgatggctaatagcagagaagggccaccagtactcattgtgcattgtgaag
gctctgtacatgagattggtgctggtttacatacaggatcgggccgactgatataaccctaacacctaaaactcagtgggggagatttatcaaacatggtgtaaa

**D:** Amino acid sequence (signal peptide shaded)

MKAATIFLFSVVLILCYCWTTQGHYSAGKSHGKFIVARRTEHGDTPGSIQDGKKYLICVCDLKKANNLRKKRHMRAQGKHGHHGRRCARICMQKGAHRG
SQVSIPLT

**Fig. S8.** Gene position and architecture (**A**), genomic DNA sequence (**B**), cDNA sequence (**C**), and encoded protein sequence (**D**) of the unannotated *cxcl17* gene in the NCBI reference genome of *Dendropsophus ebraccatus* (hourglass treefrog). The information was downloaded from the NCBI gene database (https://www.ncbi.nlm.nih.gov/gene).

### A: Gene position and architecture

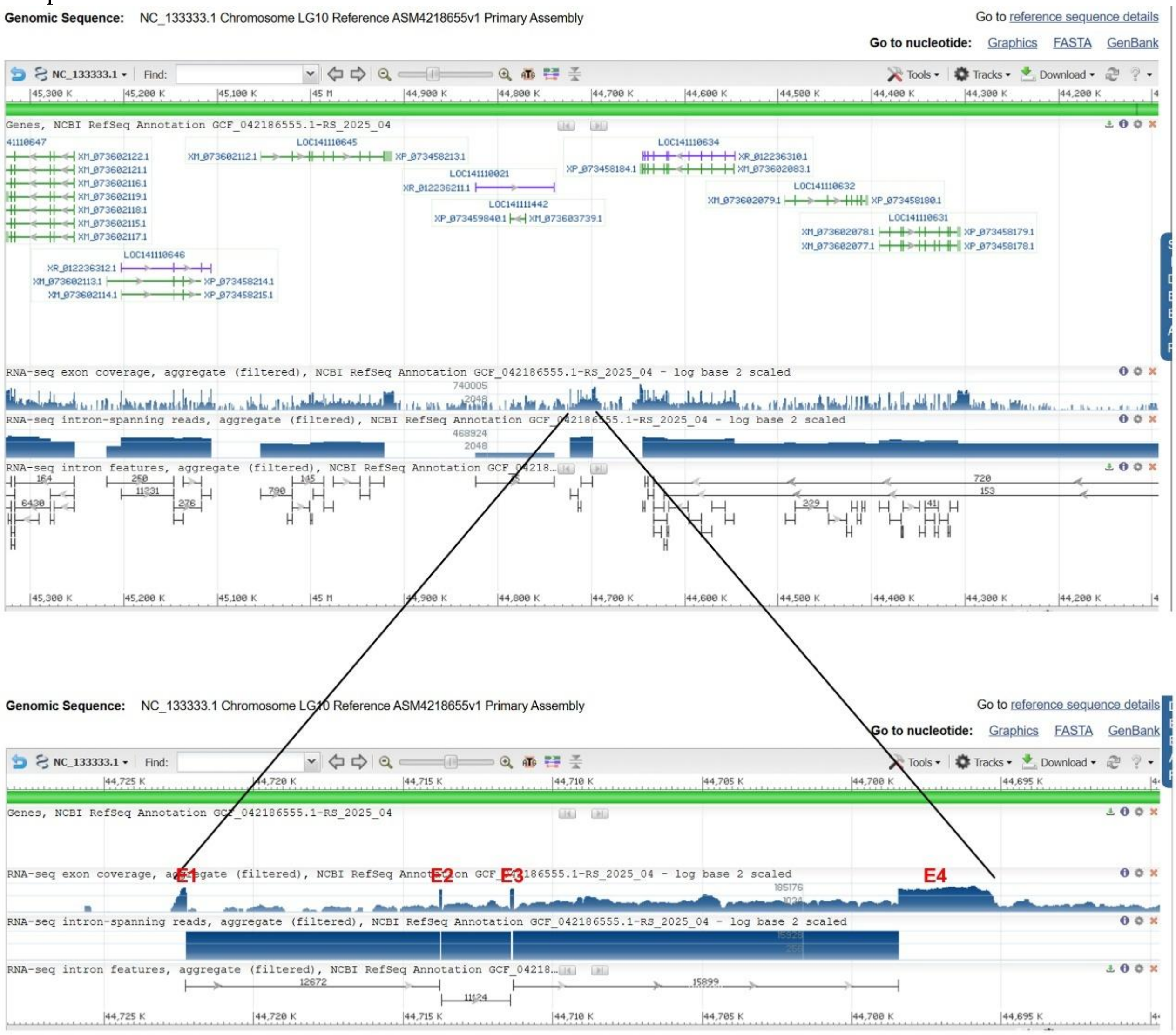


### B: Sequence of genomic DNA (exons in red and highlighted in yellow, coding region underlined)

aacgctgagggaggatctcaagcatgtggtgaaacagttaaagttggggggacacttgacattccatcagattttatacctgttgtacagccctttcctgtgtgagtcttaaactcatatgaaagcaaggcggaggtgagtaacagacggcaacggaatagtgct
gatcctcatacaggaacaattgaaaccatgaagttgtccacgattttcatgctagctgtggctctggccctgttctactgctggactgctgatggtcatagtaatgcaggtaagattgtaccagaatctgactgatgctaagacaggacactgtagcccaggcaagta
aagaagggcatcaaaactgattagaaagcccgtcctggacttaggaccttcacatacgattaaacagtgcaaggagcaataccactttgttggctttcaggtgcccgctggtcacaaacaacattgggttgactgcatttatatgagtaacacttaatttctgattggtt
gctaggagttaaagcatcattctcttagtgaccgagtcgggtggtttaggtaattattgggcaaacagtagtactgaatttgccaaacagaacacaaatctgaagagtgcctttatttaacattgatatattgtgaagatctgaagaacatacagtatctcaaaaaagtg
agtacacccctcacatttttgtaaacattttattatatcttttcatgtgacaacactgaagaaatttacactttgctacaatgtaaaatagtaagtgtacagcttgtataacagtgtaaattcactgtcccttcaaaataactcaacacacagtcattattgtctaaaccgctggc
aacaaaagtgagtacacccctaagtgaaaatgtccaaattgggcccaaagtgtcaatattttgtgtggccaccattatttgccagcactgccttaaccctcttgggcatggagtccaccagagcttcacaggttgccactggagccttcttccactcctccatgatga
catcacggagctggtggatgttagagaccttgcgctcctccaccttccatttgaggatgtcccacagatgctcaatagggtttagatctgaagacatggttggccagtccatcacctttactatcagcttatttagcaaggcagtggtcgccttggagtcgtgtttggg
gtcgttatgttggaatactgccctgcggcccagtctccgaaaggaggggatcatgctctgcttcaatatgtcacagtacatgttggcattcatggttccctcaatgaactgcagctccacagtgccggcagcactcatgcagccccagaccatgacacttccacca
ccatgcttgactgtggacaagacacacttgtctttgtactcctcacctggttgccgccacacactcttgacaccatctgaaccaaataagtttatcttggtctcatcagaccacaggacatggttccagtaatccatgcccttagtctgcttgttttcagcaaactgtttgtg
ggcattgtgcatcatctttagaagaggcttccttctgggatgacagccatgcagaccaaattgatgcagcgtacggcatatggtctgagcactgacaggctcaccctcacccccttcaacctctgcagcaatgctggcagcactcatacgtctatttcccaaagacaa
cctctgcatatgacgctgagcatgtgcacttaacctctttggtcgaccatggcgaggcctgttctgagtggaacctgtcctgttaaaccgctgtatggtcttggccaccatgctgcagctcagtttcagggtcttgacaatcttcttgtccatctttatgtagagcaactat
tttttttcagatcctcagagagttctttgcgatgatgtgccatattgaacttccagtgaccagtatgggagagtgagagtgataacatcaaatttaacacacctgctccccattcacacctgagatcttgtaacactaacgagtcacattgagacatcggggagggaaa
atggctaattgggcccaatttggacattttcacttaggggtgtactcacttttgttgccagcggtttagacattattggctgtgtgttgagttattttgaggggacagcaaatttacactgttatacaagctgtacactcactactttacattgtaggaaagtgtcatttcttcag
tgttgttacatgaaaagatataataatttttttttcaaaaatgtgaggggtgtactcacttttgtgagatgctgtacataaaaggcaaatatttgactataagagtaattagaaatacttaatttcattaattggacattttaagagtggtgcattgtctcatgtaaaatatatattta
gtccaaagaaagtcttcaactttttacccaataccactgaacctatacagttgcagctagtgttctgctgttgtggacttaaggtaaacctgccataagatcaaataaattaaattaacatgacctttaatgctacaatgttaaccttctgaaaataccagatgtctacttgtg
tgtggttttagtatttgataaccagacctatacggagcaaatgaaagtcagaaaaaagttagaatttcctatactcattttggctcagtaactcaaaatattgaaaccattggatcagcatattcatcagttaaatctttctgaaggttcaaattgcctgacatgctgtatattt
cttcaaaatgggtttagttataaaaaaaaaaaaaatgcatattggtttgttcagtgaatctgcatgttttgtgcaaactgggaaaaaaaaaaaaatgttcttttgaccatttttttcttctgactgaatgttcccttatttatttgcaataggaaaatactgcattacggccactaaatg
gagctgaggagcataggaaatacacatgtctttagaataagagatacaatttaacaattccagagaacattggagggaattagcaaataaatgtgaccaacggtctccagaattgtaactttgtaccttttgttacaaagatgtaccgtatatatttcctatgcttcgcag
ctccatctagtagctgtaatgaggtattatactactgcccataataaaagaacattcatgtagaattaaaaatagcctaaagaacatttggtttttgcccaattcacatataatgaatgtataaatgtatatgcagaatcactagaatttttggagaggccaagaatagtag
ggaagactacaactccatttttttaaattcctgttggtgtccctgttccattccttctcatgcctcctgatagtaaaatcattgtcaccgggacaatccttatacactctatccacagcttaagaaaaatgtttttggcttgtcaaatctcttggtatacattatcctgctctctttgt
attgctttatttgtatgaaatacctggtgatcctgccagtccccctactttactattttaaactgaccactctaagcaggagatcacatccatctggctgtgctggaaaaccagcctacctgtcctccaatggccaagacttgtgttgacaccctcctgcacagtcattaac
cgagaagatcagtgtgctgttgtttcttgaccaccagctctctaagccgcttatgcaacagaggacagagattatgtgatcacttttaaatataaagaaaaaaaaaaaagcagtgtttttttttttaatatatacactatattaccaaaagtattgggacacctgcctttacacg
cacatgacctttaatgatatcccagtcttagtccgtagggttccatattaagttggccgaccctttgcagctataacagcttcaactcttctgggaaggctgtccacaaggtttaggagtgtgtttatgggaatgtttgatcattcttccagaagtgcatttgtgaggtcag
gaactgatgttggatgaaaaggcctggctcacagtctccgctctaattcatcccaaagatgttctatcgggttgaggccagtcaatttcctccaccccaaacatgctgatccatgtctttatggaccttgctttgtgcactgttccaaatcatttggtggagggggggatta
tggtgtggggttgtttttcaggggttgggggttggcccctgagttccagtgaagggaactcttaaggtgtcaacataccaagacattttggacaatttcatgctcccaactttgtgggaacagtttggggatggccccttcctgttccaacatgactgcgcaccagtgc
acaaaacagggacagcgttcccagaagagttgaagctgttctagctgcaaagggtgggccaactcaatattgaaccctatgggctaagactgggatgccataaaagttaatgtgtgtgttaaggcaggcgtcccaataattttgacaatatagtgtataagtgaaa
gtgaaactgcagcagaaaataatcatatccccctctgacaggcaaggctgtactgtttagcacagttttgtatatttcaagactggatggacagatataaagatcccattggcagttaaaatagctctcatacagtatatgcatttcatatgtgtattaacagtttacttgg
cgttttaatttcccagcccctacaaatttgttataagacaatttccattgtgtatggtctctctgcaccatttatgcattgtataaaaatgtttgccgcatatatgtataatgactgtaccttctctttatataacgtggctccatactaaaaccattaaagcggagttccaccctg
aaatttttttttctgaatcagactcctctaaacctaaaaaaaaaaaaaacatttggaggaaaatttttttttatacttacaggaatcgctctgttgctatgtagtcctcctaatctgacacttcctggtccgcggagctcctcgtcacttcctccctcgcggagctcctcgtcact
tcctccctcgcagagctcctcatcacttcctccctcacctgtgctcctgggaaattatgtgtcataatttcccaggagtctgtgggcagtactgtgattatgccataacaacggggcgggaccttcctctgtcgccaaccgttgttatggcaactttagaaacaatcggc
gctacagccggtgaatcgcacatgcgcgagatcgcctctccaaaacgaatccaggaagatatctcaattgggcttcacataaccacaatcatgatggcaacggcaacaacaggaggcagtaaatattcagtaagaaattgttatgtatttagcaaaaatattaaac

attgtatactagtttttgaatagtattagaaataatgttaggatcagttaaaaaaaaactggaactctgctttaaggtgccccttgcttctccaatctccatttaaagctgggttcacactatgtacggcgcggggcacagcaggggggtccggtccctgttcactgttgtca
ggtccgaattcagacctgaaacagagcaaagacgatcagaacccttctgctgtgcgctccgtggccaccccgcagatgtgtgaatcggcctcattgagagctggtcacagtctcctgtcatgcgaattggatgcggggaaaccggcatccaattcgcataagtg
tgaaccctgcctaaaggcaattcctgtagtggttcagaatatcagatgtctttatagcaaacatacagaggttgtcaaagtatgtatttattgaatgtatcaatattggacaaattctacagctaattacaataaatcaatgtgcggcaagaaatcatagccccctgtgctt
ctgggccctatagcaatcatatggttacgaatacatcttaattccagtggaagttcaagatggaatccctgctgttgttattttctagcttttggaacctgggatggaaggtgctatgtgatttttatgtttattttccggttctttaaaaaaatgtggcccagttttggaaagcc
attcataagggtccattgaagatgatattaaaagatacagggccatcttaagagcattataggcccccgggcaatacagtgcactggggccctgtctacactatcgcgcacgagaataaaaatacaaattatcactaaggctatatttattggtacttccaacaaaat
cagtgtttaaacattaacacaacatttggataataacgttgcagtgtttgatgggcacagtggctgcgtttgatggtcacagtggcagcttttgacgggcacagtcagtggcgtctccagctttcatatttagggggggcacatgggggggacagggacaaaagtag
gggggcaactataaaatgcgtgtatatatatctatatagatatatatatatctatatagatatatatatatctatatagatatatagatatatatagatatatctatatagatagagatatatatagagatatatatatatatatatatatatatatatatatatatatatactg
gggccctttactacgaccccacaacggccctttcacatgttctgcagtgagctcccttcctactgtattggcgtcccccagggtggcagaaaacaagatatatatgtcaccagcataccaagaaaatataaggatccaaagcagtgggagaactatcagggttgc
aaaggttgtcttgcctccgggccctggtgttctgccactgtggggtttcccagcctcctcttgctgtcctgcccctgctattgactgctctggtctggcatctcttggaatttacaggttctcctgtcctaaggatgggatcaatcagtctgtttttttcagtaactgagagtc
ctggtggagtccttcctgcaactcgttcttctccctgccaatgaggatgcaggggaaggagcagatcgggcggccgtggttgtgtgagcgctcgcattatgcagtgtcagcagagggaggggggaggaatcacccacaggagctgactggggactctctccc
tcttagacattgatttttttttttaaataaataaataaaatcttttttttactgggggcacatggtggggcacaacataatgttgggggggtcagggaccccctggccccccctagggatgccattgggcacagtggcagctttgatgggcacagtggcagctttg
atgggcacagtggcagctttgatgggcacagtggctgtgtttgggcacagtggcagctttgatgggcaaaatggcagctttgggcacagcggcagctttgatgggcacagcggcagcttttgatgggcacagtggcagcttttgatgggcacagtggcagcttt
tgatgggcacagtggcagcttttgatgggcacagtggcagcttttgataggcacagtggcgtgtttgggcacagtggctgtgtttgatggacttacctgaggccgcctgcctggtgcagtgcagtgagtcacagcaaggcag
ctatcaatctcagggctgtctagatcaatatctttctttagagaccgaaagtttcgctccgctccacctttgctccctctgagccacgctgtctgaagaccatggcagaagtgggcatggagcttgttcactagctgcaaaggggggcagcgctgtgaatgctggggc
ggccgcgacctcagcctcagcctgagactgaggtcactggttgccagctagacgccaggcggccggcgattctagcaagtgacgagcaaccagtgtagtcagtcaacttcatgtcttctgcagcagattgcagcgctgaggatcagccctgggtggggccct
ccaggcaagtggggcccccaggcaaatgcccagcatgcccagtcaaaaaaaacggccctgaaaagatacatgttcaagcaccttcatccccaattccccccttcctgtgacagtgggctgggaatcttcttgtttgaaccgctgtctcaatttgatagcagcgtca
ctcattcacagtttcctgtgaatgaatagcctttaagtaccgtcagccgcttatagttctcagtgaattgtgaaggcgctagtgagcgctctcctagtctattgatcttcctgctctcaagtaactgcctgtccatacaaggtacaggcagacagctatcctgagagtctg
cagggcctgagattgcacctgtgctctgctgatcacgggtgcagtgctctgcaggctccttaaacccctttcacactggggcgctttgcaggtgctacagcgctaaaaatagcaccctccaagagccgctcctcacactccagtgtgggagcccgagggctttca
cactggagcggtgcgctggcaggacggtaaatacagtcttgctagcagcatctttggagtggtgagggagcgttgtgtgtataccgctcctgcccattgaaatgaatggggcagcgcggctataccgccggcaaagcgccactgcagcagcgctttgcggtg
gttttaaccctttctcggccgctagcaggtggtaaaaccaccccgctatcggccgaataccgccgctaaaacgacagtaaagcatcgctaaaaatagtgccgttttaccgccagcgcctgcaccgtcccagtgtgaaatggctcttagaaaaaaacaaatgcaca
ttttacctgcaaaaaaatttacatttatttttttcataatttattctgccagagcccaagaacagtacaatacattgcactgtcatttttctcttccaggaaaaaaagatgataagagacaactacagaagaagaatcactccaaaggtaagtcaaaagtggtgcaccacct
acaatataatcttaggccagccatacatggttcaaaccattaatgggcaggttgaatgtaccaaatgcttcttcttttgtccatacggacatgtgttcataaactgtcatcaataagcattacatgtcccaggagctttattatgtaactgatttactcaccaagttgaatctg
tatatgtatgtacagaaaagattatcatgacagcaaggcataaaaacaaaaaagctattttaaagcggggttaaacccaaaaacaaaaaatataaaatagtgcagctcatcaatctttaatgtggtggctgaatttgattttctttttttttaggcttctttccttttattttcac
ctggagatccatccagtaacacaatccctgtcacagcaaaagttatatttcctttcggaataaaggttttacataaaagaataaaagatcatcattgtaagcacccctgtcagagtttatggtttgtctgaaccacttaactgtcacttatgctggacagttcggttggttaa
aggaagttaaaaaataacagacttactgatcaatcaccaaattaaaaatatctaaaaagaaaagtaatgcatccatcacatctaaggattggtaagctgcaatataatacatttctgtttttgagtttaccctaaatgggttgtaaaggttcactaaaaaaaacaaaaaaa
aaagttatacctacctgctctgtgtgagggttttgcacagagcagccctgattctcctcttctgcggtgccccgctggggctcctggctgctccccttcaccaggtgcccccacagaaagccgctttccattgggccacgcgtgtgggctcgctcctgggtcatgct
gctgcgtccattgacacagacagcggaactcagccccgccccctgctcctgtgtcactggatttgattggccgatggctcctgctgctatcaatctgtccaatgaggacagagacagtggctggagctgctgggctcgtgcacatcgctggatcggttcgggctc
aggtaagaaaaagggggggcctgggggggcagctgcagcaaagaaagtttttcactttaatgcatagaatgcagactttacaatcactttagccacttctggactgccgcacgccgatatacgtccttattttgaagggagttatcgttgatatggcagcagctagc
tgccataacccccggtatcctcttgtttggccggcggtccgctttcagataaaagtggtctctgcggcagattccccgccatgctccagtgccctccgctgcttaccggagccatcggcagcggcggaagggatcggatcttcacccttgctgggcatgtagacg
agtgaggggaagatggcccccacccgtctccattacagggcggaagcgatgtcaaaacgtcacttccgcccatagctcttaaagagccatttttttttattgcatttaattgtaaatataagatctgaggtctttttgaccccagatctcatatttaagaggtcctgtcatg
ctttttctattacaagggatgtttatattccttgtaataggaataaaagtgacacaatttaaaaaaaaaacaaaaaaaacagtgtaaaaataaaaggtaaaataaataagaaaaaaaaggtttttttaaacgcgccccgtcccgccgagctcatgtgcagaagcaaac
gcatacgtgagtagcgcctgcatatgaaaactgtgttcaaaccacacgtgaggtattactgcgattggtagagtgagagcaacaattctagtcctagacctcctctgtaacttaaaacatgcaacctgtagattttttaaacatcgcctatagagatattcaagggtaa
acgtttatcgccattccacgagtgggcgcatatttgaagcgtgacatgttgggtatcaatttactcagcataacattatctttcacaatatacaaacaaaaatgtatgataccttttaaaatatatattcaatttcaaagtgtatagcattgttgtgggcttatgtaactgttgtat
tatacatgacttggatcccaaaacgcaatcactaggtttctgttgcaatgtgagtttcaaaataaagaatatagaaggatatggttagaatttacctttttattagcacatcaagttaccagaatatttatgttgtaatatacactacatcaaggaagtttattctatggattgc
ctatggtttatgttatgttaattttcatatcacacattcttcatctctttgcaggacatcatgcagaaaatataataattcaaaggaccgatggaaggaaatttgaatgtaaatgtgtgccgagaaaccctcatcaatcacagagtgagttctagaaaggaacaaggggca
tcctccttaactgatcatatagcataagacccaacagttggcttattatttgtgtgtaaggacctgtgtcaacaaaccttggttagttttaaatgggttttgcattcccatacaagtttagattagaagaaggtcaattgcacctatcaatgatttctgaatgatcttagaagca
tctcaaactgatgccattgaaaaggccaaagcccgttgcgccggccctaaagcttgtataagatgaccatattcctactgatgatgaaaaaaaacgtgtggttaaaaactgtacagtaaaaagccacctctatgctttggtcacctataggcacagggcttttatgcc
caacggtaaacccaggtctacaaatgtatatctagttattgaatccttaaacgttcaccttttccagaaaatggcctatgtagtcaataagccccagaagatatcaaaaatctatgcagcggaagcacaggtgcgaagctgctgagcagtcatgaatgcttagagctc
cgctgtcaccgggaccaaaaggccttctatcgacaagtcacggtcgaataccgcattaatacggtaacccacttattactgagactacggggcacaactacatataatgtttagcgggaaaaagacaaaacttaaacagccagcaggcaaagaaaagctgaca
agaaacgctcctaatatgacagcagcacaatcttcccgctctgcattacctcagagacagtcacaaaggggggaactctacatatttttcctcacaacttgctgactctgaccttgattatggattaacaccaatatctctacagaatattcctaataactcggatattctta
gtcagtttgaacgtgttttacaaaaagccctccaaaatacctcagacacaataacagaaaaactgacaagggaaattcgtgaggtagggcgccgcacatctatcttagaacacaaaatggaggatataaatgatatcactaatactcatactgacgagctagaagc
cttgaaagaagaaaattttactctacacacacgtctggaagatttcgaaaacagagcgagaagatccaacctgcgattgcgcggtatacctgaaactataatagatcttcaatcaaccattacagcactttttcaggagttagccccatcaatccccattgaacgattg
gagatggacagaattcatagaactttggcccctaaaaaagaagatggtcccccagggacattgtaataaaattacactatttagaacaaaggaacaactgatggaagcggcacgtaataaagatacattaacctttcaaggccatcgctaccaagtattccatg
atctctcgcagcaaactctctacagaagaaaagagatgaaacctcacctacaaatactccaataccaccacatcccctatcgttggagttttccatttgccctgcgattcacctataaaggtacaacatatacctgcaagaaattggacgacctactgactgcgatcc
aggagcttcgtttgtcaattccagataattcgactgaaggctccaaacgcagagcatcatcaaactcacccaaggctcagcaagacaaccaaactcaactagctcactccagtaccagtaaaaggagtcgttttgcatcaccacctcctaattccaaagatcaaat
gaactgaaagccactacgattcttttatttcattgaaaccttttacctagtgttactattataacggtaaccaatttcaggtagattcagagtagggatcgaataatccttcctctgtctttacctttatccttataatttcatttaagcagaagttaatcaatcctttgtaaaataat
aattgttaccaaactttattactccaatatcatttttaggtttagtttagtaaatgcaatgcccttttttataagtgcatgatgtaatgcaaagttatagaaacatatctctattacattatcctaataatagctattatattattagtgacttgtattattttgttattgtttgtatagcctt
tgaggaggctgcctttgtcccggagacaatccactgcgcctctcaatagatcctcgagtagccggacttactgaccggactctctttggaccatttgtacccaccggttggtgatatttcaccaccccttggatactagctccatataggagctattttcctctttatttaa
aaacaacttctcatctactgctctaaaacaggaattttacttattctacttgtcctattcatccccctttattctttttacccatcctttcctgcccccccccccctttttttttttttttttttttattgtgttttgtttttttgtttttctcgcccctccagagtataaccttcttggatattat
cacattaagaatttattataatccctaatacctttattcaaaaatgtctttaaatgtactaacacttaatgtccagggtctcaactccccaccaaaacgaacaaaagcctttcattccttttcttcactttctgcacatgtaatatgtctgcaggaaacacacttctctacacag
tcttcccctaaatttataagtcgccaatacccacaagtctatatggcctcggccactacaaaacaaagaggagtgctgattgcatttcataacacaacccccttcactttacatagagaaataaaagatcctgagggtcgctatataatccttacaggtctcctgttagat
aatgaaataactatagtttcttactatgctcccaacagaaatcctctttcgttcctctcacacttgttctcagtagttgaatctcataaaaaaggtaccctcatgatgtgtggagactcaaatcaagttttatacccaattttagacaaatcccctacaccactacaacctcaa
cgccttaatttcagacaactccttgaacaacattctcttattgactcttggcgtgaacaaaaccctagaagaaaacaatacacttactattcccatccccacaaaacattttctcgtatagaccaccaattcgtccacattggcacgttaccacaaatacataattccttaat
catcccatgcgcatggtcagaccacaatgcggtcatgacgtccttctcatccttgatcccgaggcctagccatagatcctggactataaatgataatttattagttgacccaacataccgtctaaaaatagaaacagccatacaagattacatcgctaataatttacca
acagatactaatccactgaatctatgggaagctcttaaaccttccattagaggagtctgtatcagccagtcgtcttaccgacgtaaattgaaacagcaaatgcaaaacaaattagaggaagcttattataaagcctctgaaattttccaaaatttgccctctcaaactac
aaagctagactatgataaagcaaagttggagcttgatctttttttaacagactcagctatacagattcttcgcaaaactaaacatacaatatatatgaaagctaataagccaaccacattcttggcattaactctacgcagagcagaccacattccgaaaccaattcggc
tacaaatcaccaaaaacaattttctagtaatccacagaaaattatcaatgagttccgtaggaaactcaaagatttataccaaaatacacaaacattcgactcaacaaaattagacgcactgttatcttctattcttctcccaaccctttctgttacccaacaagaaacttta
gataagcagatcactaatatggaagtcatatcagcaataaaatccctaaaattgcataaacgcccaggccctgatggatactcagccacattttataaacactttactacactactagccccacttatgacaaacgcatttaattctctattagaaggctgtgcctttaac
gcggaatccctactagcttcaatatccatgattccaaaaccgcatgcagatgacacgtcatggtccaattacagaccaatctcaatccttaatatagacataaaaatattagcaaagattctatctaatagacttaatccaattattggcacattaattcataaagaccaa
acgggtttcatacctttcagacaagcaggtgataacatacggcgtgctaccctactggctcatatagccaggtctcgcaaaatccctgcctgtttcctctctatcgatatacaaaaggcgtttgacactctgtcatggtcttatcttaaagccattcttcataaatggggat
ttggtgagcggttccagggatggatttcctctctttatcataatccaaaagcgtatataacctattcaggttaccgctcagacccctttaatattgaaagaggaaccaggcagggatgtccattatcccctcttctattctccttggctattgagcccctagctcaacttatt
agatccaatataaatataaaaggacttgaattaggtggtcaccaacataagttatgtttatttgccgatgacgtcttagtatttcttacttcgcccctactatcagctccaaacctcattgaaattcttacaaaattcggtaaaatatctggtcttaagataaatcagcaaaaat
caaaagccctaaatatatcgttatcacctgatttacagaaacagttacaagactcccttccatttacatggtcaacaacacatttaccatatttagggatctcactaacggctaatccagctgacctatactctgcaaattatcctcccatgctaactcaattatcaacctta
ctgagaaaatggtcttctttaccgttagcatggatgggtagaatcacagtagtaaaaatgtccatattgcctaaaatcctatacctgttccgtgtacttcctataaacgttccggcttattttttccgaatactacaacgcaaatcagcccaatttatttggggcaaacttaaa
ccacgattaccacaaaatacactacatatggcccgaacctgtggaggtttgggtgttccaaatttctcaaaatactattatgcttctcaattagcacaaattattaaataccatgccactaaagaaatccctctttgggtaactatagaatcggtggaatgtgatccacttt
ctatatcaaatctgttatggctttcaccaccccaacgacgttcaataacaaatctagtcacaaaacatagcctctcaatatgggatagacttaaattccgttttggtctccaatctccactaaaccctctcctctcctttctaaacaacccctccttctacccagcttggactt
taccttcctcctttaaagcttggataaatgtgggtcttacacgagttcatcacttctatgactctgataaacttatttcatttcctacaataagcaagacacacaacatacccagtaatgaaacatttagattcttacaaattaaaaactttttatcccaaaacatccccacaaa
cttgacaactccagaatatacttcatttgaacgatcatgtatgtcagatccacacagacgaggacttatctctgacatatatgctcaactactttcacaaacaaattcctctcccccgtcttatactaccaaatgggaagctgatttaaaccttcctccagactcactggatt
ggtcacaaatctggcaatctacaaaatcagccacacccaacatagttgccctagaaacaaattataaagttcttaccaggtggtatttagtgccagcacgtatatcaaaataccttccccaataccccccctcactgttttcgcggatgcagcgacattggtacacatct
gcatatttggtggacttgtccaatagtccaaacatattggtcaaaaatatttcaaattctctctacaatgtttgaaaccacattacaaccagatccattggtagccctcctaaataataagccacgaaacataactcaacgtcaatttaaacttttactatatgtcacaacag
ctgcaaaacaatctatagcaaaagcctggaaaaccaataagctttgtgacatatctgtaaaacaacgtgtaactcaagctatgatccatgctaaaatggaggcaatcatacttgataaactggacaaatttgaatctctttggcttccatggataacacacatactacc
accggatttgaactattcacttcttttaccctgatgagatctaccaccttcacggtctatattattacatgttcccataaatatttaacactcgcttctctctggagaccccaaattccctccataaccatttcagatcccaccacatattcccacttattaaattttatttttagtga
tgtataataataaaatgtgtctttatctatttctattaaattattatttggttaggcaaactatgtctttttcggatgcaaactatctataccctttgatggggatgctttggagggacacccatgcacccgggatatgccgggccaaatagccggaccttggccctggttgtttg
ggtgtctctctaggtcatctcattaatacccaatttatctaatttagttgcattttatgtctcactattgtgacatattgtatttttttttttttatatatatttatctacttacttatttatttatttttacaattacaaaagctctattaattattataaccatataattcagatatgtacactgg
ataatagattttcttactgatttctagaagataacttaagtattattattctaagttattattgagaagtttcttttcctcatattttgtaaccttacataaatttaaaaccagtactgtaaaattgtattgttgtattctcaataaattatttgaaaagaaagacaaaaatctatgcagct
ttgaaaatgatcatttcatctgtaaccattttggatccctcagaagcctggctaaaaaaacctcaaatttagcaacccccccatgtcctcccttttttatcaggatgtgaaaaccacttctcacttcttcaatgctgggagtgagcatccaatggttttcacatcctagtaacaa
gtcaggacatggggggtttctagctaggagtctttttagccaggcttctgaggggacccagatggcctataggtaaaacaataaaataatgatcattttacaaagttgcacagatttttaatatcttctagagctccttgactgcatacactactttctggaaaaggtgaa
ccttttaggtaatgaagtaatacatacatatttttttttttgccagggggaggggaggcatttttggcagtattgtcttgcaagaataattttctttctatgtaatctttagacaaaaagacataccaaaatgaaaaaaaaaaaaaaaaatttctgttttgagtagggttaattttgt
taaacaatgtttttattgatgttttacagagagaaagggtttacaaacagtcaagtggaacatattccattatcgaaaaatgtacagaaatgacatcagggcctagttaagtcattgcacacagacaagaacatgtaggataaaccatgagaactttgtgaatggggaa
ttaataacaagcacatgaagatgttgactgtgcgagcagggttaattttaatcgcttgacctccggaagcttttacccccttttatgaccaggccactggggtttttattgtttattatataaatgaggaaagaccaaaaaataaaaaaaaaatattttctactttctgctttaaa
acatatccaaacaaaaaatctagtataaaaatattggtaaacccaaatgaaaattatatatatatatatgtgtgaagctgccccttaacccccttgccgaccagccaccgtcattttacggcagcaggtcggcacaatcccgcgaactgtcgtagctaaacatcggctc
gcgggaccaggatagcaggcacgcgcacgccaccagaggcgggggtgccgatgcttgtctcgtggccgacagttgcgatgcccgcctgccacgagcgatcacgggcacgagaggcagaacagggaagtgtgtgtgtaaacacacaaatttctgttctga
gaggagtgacagattgtgagttcctattagctaggaaccacgatctgtcatctcctatagtcagtcccctcccccctacagttagaaacacatagcagggaacacacttaaccccttgatcgcccccctagtgttaaccccttccctgccagtgacattttacagtaatc
agtgcatttttatagcactgatcactgtataaatgccaatggtcccaaaaatgtgtcaaaagtgtccaatgtgtccgctataatgtcgcagtcacaataaaaatcgcagatcgccatcattactagtaaaaaaaaaaaaaataataaaaatgctataactatgccctatttt

gtagatgctataacttttgcgcaaaccaatcaatatacatttattttataaaaatggggatatttattatagcaaaaagtacaaaatattgcgttttttcaaaattgacgcaatttttttgtttatagcgcaaaaaataaaaacctcagaggcgatcaaataccaccaaaaga
aagctctatttgtggtaaaaaaaggatgtcatttttgtttgggtgcagcgtcgcaggacgcgcaattgtcagttaaagcgacgcagtgccgaatggcaaataatggcctggtcattcagcagccaaatcttccggggctgaagtggttaagtgagcaagtcaactct
gaaaaggaaaaacaatgaaaagcagggcgctaagacatatctgatatttgcatgtgaacaaaaaataaatgacaatccaaactaaacttagaagtccaaagatgagaaaaattatgaaaattatatattgcacagtccaaaaaaattcataaatcttgctgaagaaa
gctccaatcacagtgctcctggttttgcctcggatactcctcatataagtgctccaccacaaagtgtacaaaatgcacacttaccaacaacatcaacaatgtgcgcttttggtgtcttatggtaccaccttttttgcaagttgttttcacaatgtcccaccttggcagtggta
taggggaaactaccacaagcttttctcctcaggaatcaataactgagacccagggagagataatgatccaagatagtgtgataacattttattaaaagtcaaaaaaggattaaaatatgaattttccacttacatttaaaggtgctattatagtcagcaacaacacaaac
agcctggttcaattgtcccgcacacctctaatgccccgtacacatggtcggacattgatcggacattacgacaacaaaatcctaggattttttccgacggatgttggctcaaacttgtcttgcatacacatggtcacacaaagttgtcggaaaatcctatcattctgaac
gcggtgacgtaaaacacctacgtcgggactataaacggggcagtagccaatagctttcatctctttatttattctgagcatgcgtggcactttgtgcgtcagatttgtgtacacacgatcggaaattccgacaacggattttgttgtcggaaaactttgtgtgtcagaaa
atctgatggaaaaagtcagatggagcccacacacggtcggaatttccgacaacacgcttcgatcgcacattttccgtcggaaaatctgaccgtgtgtacggggcattagtccgagtcgcttggtacgaccgagcggtgtgcaggacaattgaaccaggctgtttg
tgttggtgctgataatagtacctttaaatgtaagtggaaaatttatatataatatgtatatatatatatagatatatatataatggtaaacatgttagtgtaaataaagtataaacagtatacttgagtttaggctctttccctaatggaacattgtcaaaaagagaaataagact
aactggctacagcgcttataattgcctcaacctatttgctgatgtggagctccacatcttaatacaggtgtccggagtgtatatattttgcttgtggtattcagtagggttttaatagcgatgataaaggggggttggtggtagtggttataagaggttttcgaaagggtagg
gggtgtcattaatggtcactattgggggggaagggggaaagaaataaggccagaagttctgaactattcttgtcagctgtatttttgggtaactaggtgatagatttggtgtattcaccagtagtgataaagtggaggaccatatttttcttaatttaggagtatcattcac
ctgatgtataagttactccattgctgaggttttccgtataagtcttacccagtctgttatggtcgcaggatctgaggagcgccatttggcgggtatgcacatatttgctgctttaactatatggatggctaacaagcggaggtatgcatcttttgaatgtatgtgtagatgttt
tcccattctatttctggtcaatatacagatcctgctcccacttgaggcttgctaggtttgatttggggttcagatccgcaaatagtagagcatatgtgccagaggctagatgtcttttgggttccatttttaggcacatatattcaactacaacactttcttgaaggatctcaa
ccccgaccggattggtcccgacagactacaccttttgaatctatgaccataatgggaacaattggttagacatccgggagaacaacaatgaaagagggatgaggcgacctctttcatagaattgtgctgtgctgatgtttggatgtggccatgtatccaagagaga
gtttgcggatatgcccggagaaaagtctgggttctgacgatttggcgtcatggggccaagagttgatgagattttgtatttgcgacaggcttcctggaaggttactaaggtgggattcattaatgggtgatttctgcaagtcggttgggtcaccagggaagttgatgta
gtgggattttggacatagcatgtttggaaaattcatattttaatccttgtttgacttttgataaaacgttatcaaactatcattgttcgttatagtcatatttgttgatgtttttaagatgtttctgaaatgaataaataactttttttatgtattacataccttctcctatatttatatatacgt
gcctttaaagtccaccactaggggaaccctctctctttttgtttgattatctaaatatcggatgtggcacacttttgcagttaatttaccgactaacaatccctgttcgaaatattattatgctaaggctgtacgactaaggctgttgccaaaacgtgtccactgttttgcttgt
cactttatgtaaccaaccattaaaatcgctatatggactacagagttgccggctcaatccttttacttgtttatcctgaattattatctttctctgtgtctcggttattgattcctgaggagaacagctggtggtagttttccctataccactgccaatgtgggtcattgtgaaaa
caacttgcaaaaagggtggtaccctaagacttcaaaaatgtaggccaaaatgtattttgctacatgtctttggtaaaaaaaaaaaatctctaagtggatattaattggtttctgtgaaagctatagtttctacaagctatagtgcatacatactggaatttacgcagctatgg
atgtagctaactgtaatggcagtgatcattgacttatagtgggactgtgacagggtggtgggcaatctggcgctaacagacgctggctgggagggtcactaactgatgtcgctagtgcaactaatacagtgaccagtgataatgctgtacacggacactgtactaa
tgacactggctggaaaagggttaacatttggggcaatcaaagggttaactgtgtgcccaacaatgtgtagcatgtgctgtttttactttctgatctgggtgtttttttcttcttttgctaacacagaactctgtgtgcgattggtttcagccgaaatcattggctgaaatatgct
gtgaaacttgcgctgtgtccaattagttggcaggggaggcgtgcatgctcaccccaaacctggaagaaaggtggtcatgtatatgtacatgattgtgcttgtacctgccgcctgctcgcagtatatgtagcggtcgtcagggggttaaataagtattattactgtgga
caagtactatcttattctataaaactaaaatcacttttctggtaaaaatttggatttatcaagtttatgaatgatgagaaatgatccttccctccttcactgtctattttggcaaggccagaaaatacagtgagggtagccgtactgcacatgctcagtttagtctctattgctg
gagagaacattttcctgtttcagttgaactgtttaggtcacatgatattgacatcacacatgtgggcgtgtatacaggctgtagtggaaatctcctcttccctgaaaactcagcactggacaaactgtgcagtttatcatgtgaccgtactatacagatcagcaaaaaag
atactgtatatagtggcataattttaatgtaaatggaagctttataagatgtgggcactgtgggggggggggggggggcagaggcactattttcatattgctgagtttagtaacactttaatgatactttactttatgtggtacgcataattcttatgttattgccattacataat
tctcccttacataattctccctctgacattgcagtattttatgtagtatgacatcgcagtattttatgttgtatgaactgctcttctgtgcttcacaggagaaaggcgccaaatgaatgaattccagaagcgccatcttcatggccgaaaatgtcactgcatatcattggttgg
gaaaccaaagagctccataccgttatcttgaggaaatgtcttcctgcctgcacagccaaacgccaaagaacagttacctttcagtgtcatatagccgtctgataattcttttctatgctgcgcttttactaaagttgaaccagccgaaattcggctggttcaggacagac
tagccgaattttgatctgtttgtggcctaccctgtttgacaaaagttgattgttgaacaattggaaaactttcattgattagtggactgcaactgctggtaagtgtatcctgacagaggaggaatgctcgctgacagaatacaatagcatagttgggggggtttcctcaccc
acctccaatgtgtggatggagaaatctgttcattttttttttttttccattcacacctcaggcagatataaaaaaacaaaacaaaaaaacaatcatgtactgtatacccagcttaagcgttgccctaaatcaatagggcaacatacggatcaatgatggtgataaagtgc
catgtctcatattagcaattctgaagtcctttgttaactgaaaccattataaggtgctatttaattggttgatataggttactacacttgatacttcattgtttctaaattgcagacctggtgttggaagacctaatatgtgaatgcttgttctgggaatctaacaaagtaagtgtt
aaaacaagagtcaggatgatagtaatcactaatcaataccagctctcataaacttatcccaaaatattgtcttttatttccaacaaatctaattagttccatattattacgtgatacttggcaaatgtgtacagtttactttatttcatttatcttcagaataaaataaatgaaactt
tgaggccgggttcacatatgtgcgagttggatgcgggtttttgaagctggttcacacctgtctgaggcaaccgcggtccgcattccaaaagggtccccgtacgtctttgggttcagttcaggtgtgaattcaggcaaaaattaggacctgattcacacctgatccgg
tgaacaggaacgcagcggaccccatgctgggaactgcggccgcacatatgtgaacccgctcttactgtataaagtgctactctttgatttatgtgacctgaactgtggtagaacatgtgttattggcaaatactcccactcagcaaaccttcaagtaagggcttcaaa
cataacttttgtgaataaagcataaagtagggtgccttgttttaaaatatcatgaaaaagtgtatatgagctgctggactggatctttccacctttagtcggggaagtgttggttgggcatgatggctgaaccttcggtagctccattgggtcccccaattaaaatggtaa
tggtttccactgggaaactcaagtgaaactcatataactttcttaaagaggaacttcactttttgcttttctcctatcccatctttcctcctctgatgagttatgtttaaaaacacatggcatccacccactaagggaatccagttttgctggctgcattccaatgctccactgc
cccaagctaggtcctgtgctaccatcttcttttcctacctctttagggggctggcttccttttctgtaggcccctgatgaaataccaacatgcatgtgagatacatcacgacacatgggcagactttacggtgctggattacataacattcttatcccaagaggctacag
gttgctggagatatgtaattttcattttggcgtaattttcaccttggtgagaaaggtggaagtagaattctttaaaaaaaaaaaaaaaaaaaaaagttcccaaatgtagtatgggggtaggagttaaaaaaaaaaagtaaaaggaagactgaaggtgtggaagattgaga
gtgaacacccatgcgatctgattctggtgtggatctaaaaaaagggtcctgcatgattttgtatggctgaatttgcatcgcacagacatcgcatgtgatttgcactgctggggctccaggctcctcctgctagctgcttgcgatgagggcacccgaccgatgttctgt
gtgtccacacatggagcacggctcagcccactaccggactttcctcattggctcactagctgtaattcacagtaacagaagccagtggctaccgctgctgtcttggccaatgaggagaaagaggctcttgtgcacttcgttggattgagatggggtaagttttagg
ggagcttgggcgctgttgcacactaaattttttttttttttttacctgcatgcatagaatgcatgaagataaaaaccccttgagcttttacaaccaatttaagcacacctccattatgactattatttatccttctctcatttagaacactttccaggagtcccatagtaagtttttata
ctttttatttgagcccagataaagggaaattgcttggcctccaaagcatgttggtctttttaaatctttctcttctacagttaaacagtacaactacaatatctcccaccccaagcccatgactagaccacagagaagcaacaccagaatgatgaggtcatcactgttagc
agattccttgttagtgcatgtgaatgcagaacactcaatctgtcctagtgcgacccttccatatctcagattgaaagaagtccatcaagttcatccaaactaactatggtactaatttagtaaaataggctaaaatcaagtcaaattcttatacctactctagcaaatgtgct
tttaaagattaaaaaaaaaaatacataactgcattcattgttttttttgttaatatttctatgaagttcagcattaaatatttgtcttgtcatgtgaaatatatagggatttactgagaaacagtgtaataatccatattagccaattaactgtcacctttccatgcttctttaaacaaat
gacaggtgaaatctcattcgttgctccttacattttctgccagacatattattatgcttcacagatatatatagatatatatatatgacatgcatttttttttaactgcacaaaatgtaacactagcagtatacacatcaagtacgtatgcctatttttatattattttgtaataaaaat
cctgattgcacaagctatgttttgaagttttgtgttgaaatcatttttttgccgtctcattacaaaactattttctaagtgagtgatttagaaagtattggggtctttgaatcatcttcattacagctagggagctaata

**C:** Sequence of cDNA (coding region in red and highlighted in yellow)

caaggcggaggtgagtaacagacggcaacggaatagtgctgatcctcatacaggaacaattgaaaccatgaagttgtccacgattttcatgctagctgtggctctggccctgttctactgctggactgctgatggtcatagtaatgcaggaaaaaaagatgataag
agacaactacagaagaagaatcactccaaaggacatcatgcagaaaatataataattcaaaggaccgatggaaggaaatttgaatgtaaatgtgtgccgagaaaccctcatcaatcacagagagaaaggcgccaaatgaatgaattccagaagcgccatcttc
atggccgaaaatgtcactgcatatcattggttgggaaaccaaagagctccataccgttatcttgaggaaatgtcttcctgcctgcacagccaaacgccaaagaacagttacctttcagtgtcatatagccgtctgataattcttttctatgctgcgcttttactaaagttga
accagccgaaattcggctggttcaggacagactagccgaattttgatctgtttgtggcctaccctgtttgacaaaagttgattgttgaacaattggaaaactttcattgattagtggactgcaactgctggtaagtgtatcctgacagaggaggaatgctcgctgacag
aatacaatagcatagttgggggggtttcctcacccacctccaatgtgtggatggagaaatctgttcattttttttttttttccattcacacctcaggcagatataaaaaaacaaaacaaaaaaacaatcatgtactgtatacccagcttaagcgttgccctaaatcaatagg
gcaacatacggatcaatgatggtgataaagtgccatgtctcatattagcaattctgaagtcctttgttaactgaaaccattataaggtgctatttaattggttgatataggttactacacttgatacttcattgtttctaaattgcagacctggtgttggaagacctaatatgtg
aatgcttgttctgggaatctaacaaagtaagtgttaaaacaagagtcaggatgatagtaatcactaatcaataccagctctcataaacttatcccaaaatattgtcttttatttccaacaaatctaattagttccatattattacgtgatacttggcaaatgtgtacagtttactt
tatttcatttatcttcagaataaaataaatgaaactttgaggccgggttcacatatgtgcgagttggatgcgggtttttgaagctggttcacacctgtctgaggcaaccgcggtccgcattccaaaagggtccccgtacgtctttgggttcagttcaggtgtgaattcag
gcaaaaattaggacctgattcacacctgatccggtgaacaggaacgcagcggaccccatgctgggaactgcggccgcacatatgtgaacccgctcttactgtataaagtgctactctttgatttatgtgacctgaactgtggtagaacatgtgttattggcaaatact
cccactcagcaaaccttcaagtaagggcttcaaacataacttttgtgaataaagcataaagtagggtgccttgttttaaaatatcatgaaaaagtgtatatgagctgctggactggatctttccacctttagtcggggaagtgttggttgggcatgatggctgaaccttc
ggtagctccattgggtcccccaattaaaatggtaatggtttccactgggaaactcaagtgaaactcatataactttcttaaagaggaacttcactttttgcttttctcctatcccatctttcctcctctgatgagttatgtttaaaaacacatggcatccacccactaagggaa
tccagttttgctggctgcattccaatgctccactgccccaagctaggtcctgtgctaccatcttcttttcctacctctttagggggctggcttccttttctgtaggcccctgatgaaataccaacatgcatgtgagatacatcacgacacatgggcagactttacggtgct
ggattacataacattcttatcccaagaggctacaggttgctggagatatgtaattttcattttggcgtaattttcaccttggtgagaaaggtggaagtagaattctttaaaaaaaaaaaaaaaaaaaaaagttcccaaatgtagtatgggggtaggagttaaaaaaaaaaag
taaaaggaagactgaaggtgtggaagattgagagtgaacacccatgcgatctgattctggtgtggatctaaaaaaagggtcctgcatgattttgtatggctgaatttgcatcgcacagacatcgcatgtgatttgcactgctggggctccaggctcctcctgctagct
gcttgcgatgagggcacccgaccgatgttctgtgtgtccacacatggagcacggctcagcccactaccggactttcctcattggctcactagctgtaattcacagtaacagaagccagtggctaccgctgctgtcttggccaatgaggagaaagaggctcttgtg
cacttcgttggattgagatggggtaagttttaggggagcttgggcgctgttgcacactaaattttttttttttttttacctgcatgcatagaatgcatgaagataaaaaccccttgagcttttacaaccaatttaagcacacctccattatgactattatttatccttctctcatttag
aacactttccaggagtcccatagtaagtttttatactttttatttgagcccagataaagggaaattgcttggcctccaaagcatgttggtctttttaaatctttctcttctacagttaaacagtacaactacaatatctcccaccccaagcccatgactagaccacagagaag
caacaccagaatgatgaggtcatcactgttagcagattccttgttagtgcatgtgaatgcagaacactcaatctgtcctagtgcgacccttccatatctcagattgaaagaagtccatcaagttcatccaaactaactatggtactaatttagtaaaataggctaaaatca
agtcaaattcttatacctactctagcaaatgtgcttttaaagattaaaaaaaaaaatacataactgcattcattgttttttttgttaatatttctatgaagttcagcattaaatatttgtcttgtcatgtgaaatatatagggatttactgagaaacagtgtaataatccatattagcca
attaactgtcacctttccatgcttctttaaacaaatgacaggtgaaatctcattcgttgctccttacattttctgccagacatattattatgcttcacagatatatatagatatatatatatgacatgcatttttttttaactgcacaaaatgtaacactagcagtatacacatcaag
tacgtatgcctatttttatattattttgtaataaaaatcctgattgcac

**D:** Amino acid sequence (signal peptide shaded)

MKLSTIFMLAVALALFYCWTADGHSNAGKKDDKRQLQKKNHSKGHHAENIIIQRTDGRKFECKCVPRNPHQSQRERRQMNEFQKRHLHGRKCHCISLVG
KPKSSIPLS

**Fig. S9.** Gene position and architecture (**A**), genomic DNA sequence (**B**), cDNA sequence (**C**), and encoded protein sequence (**D**) of the unannotated *cxcl17* gene in the NCBI reference genome of *Aquarana catesbeiana* (American bullfrog). The information was downloaded from the NCBI gene database (https://www.ncbi.nlm.nih.gov/gene).

**A:** Gene position and architecture

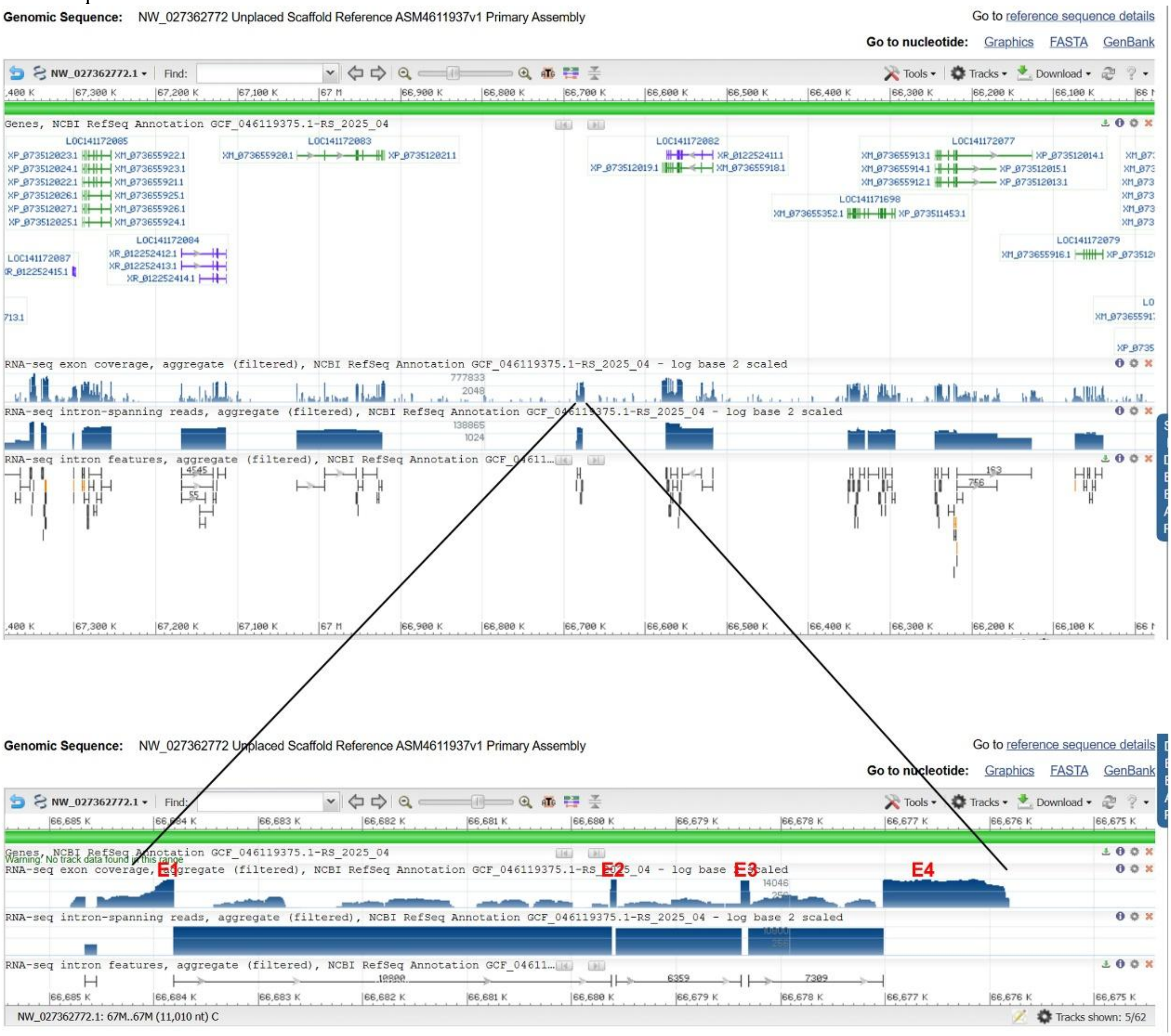


**B:** Sequence of genomic DNA (exons in red and highlighted in yellow, coding region underlined)

gaggttaccagtaagagagatgaaatcacacctggtattggaaaccatagtgatagatccatgtaaggagagggttaagactgggggacacttgacataacatcaaactcagacctggttacagccctacccttatttatctggagccaaggtgagagtacagc
ggctgtgagtggcagaaaacaaaagtatagagcttcccctgtgagaggagccaaagtgaagatgaaggcgaccacgatcttcatgttgtccgctgttctgatcctattctactgcttgaccactcaaggccgtcacaattcaggtaagttttcccaggaagctatatt
aattctggatagaactctgtatcatctgtgggactctgacctatcaaactctatggtatctatgaatgggacctttggggttctatgggatgtatagaacctatgtaactctatgggactctatggaacactatgggactgtatgggatacaatgggtccttatgaaacact
tttgaaccctgttggactctatgggacactatgggactctataagacactatgggtctttatggaacactattgaaccctatggaattctatggaacactatgggactctatgggacactatgggactctatgagacactatggaactttatggaacactattgggccat
gtaggactctatggaaattatggaactctatggaacactatggaactctatgggacactatgggtctttatggaacactatggcactatatgggacactatgggtctctaaacaacttatgatgggattctattctttcctcaggatgacacattaaatttagataggactc
cataggatctattgcagtctatagcatctatagggttttataatatatgctggacttaggtgactctatgagatctatagcaccaatgttgactatacggtatctatgggatctaactctaatggattctattggactccataggatctatgggatcctataagactctatacta
tttatgagacttatttgactctatgaaatctattagaccaatgtgataatatggcacactatgggaccccttgggacttatataactctattagacactaggggacttaatggaatgtgtgggatcctactatgcgactctatagtatatatgggatggaagcgactctatg
agatctgtaggactgatgtgactttatggcatattatgtgactgtatggtaatcatgggatctatttatctgtattcgacattacaccttcatcagttagtgcagactgctctaagaggtgtgaagacacattggcggggtcagaggaaagggtccattagaagagagaa
ggtggcatgcaaggcatgggatgcggtatccaccgctagtctccgagattgcctctttgagcagtctgcactaactgatgaaggtcttgcctatcgaaacgtctgctatgattgtgctgaactgctattaaaaatattctgtaaaaatgtttaatcaagttctatgcccag
ttctattttctattaatctggttttggaccctatgagatctacaggccccctatgatgggtcacttttgcacactattgaatattacaggactctatgagattttagtttttgcaggaatctacagggaacaattgcccctattctggaaaggtcagacattgcgattacgcata
agatagcattgggctcctcaatgttttcgttatgtagaaggtttacaattgttttataacaacatagtagtaactccatttattataccattacaagaataatccagaagtaacacgggcctcagttttcaaaactgtcttacggtaaaagtgtccttgttgcacctagcagcc
aatcagagctcagctttcattattttagagcagttacagaagtgagagctgagctctgattggttgctcggggcaacaaaagaagtctcactgttagatgactcgaggcctattaaatcaacagaatattctgccattggatctacctaaatgtttggcccctttccctct
atatctatatataaacccaaatctctttgggttttgtttctgttactttatttccatgccaacaatggcttctctgtgactatatggctaagtgctgttatagggacctaacatataatatacaggggcgtaataattgccctgggcacaaaattgttagggattcaaaagaaaa
aaaaatatcaatagtgtcgctagtaccaaggatggttccgaaaaagtgcacatgccacggtcttctgcctaggtgccccagatcaccgggcttgtgccagtttgaacagtatcggagtcaacatggcaccaaaacagttggattttgtggtaaaacatgtagctata
gtctccatcctctaggtcacagactacaagatgtcagcgtcccagcgcaggtggtgcaatgatgtcattgcactacaccacctgtgcccgcctgcgaggtgcacagggtagaagaagacttagtcagtcggaagtatttttttattggagatggcacaaagaggca
ttcatgatcaatatttttttatgagggcattaaaggaggctctcaggggtattatttgttgtaaggaggcactcagggggtattacttgttgtcaggggacactcggggtattattttctgtaaggaggctctcaaggggtattacttgttgtaatgagactctcagggggta
ttatttgttgtaaggaaactctcaggggtattatttgttgcaaggagaatatcgggggtattatttggtgtatgaaggttctcagggggtattatttgttgtaagggtgcactcagggggatattatttgttgtagtgaggctcttggggtattatttggtgtatagaggctctc
aggggctattatttgttgtaagggtgcacttagggggtattatttgttgtaaggagcactccagcagcattatcactggttttaagggcacaatacttgccttgccccaggcgttggaaaaccaaactaagccacttttattgctaagtcaattaggataggagtagatg
cacaaaattgccatagtgctaaggaggaagtgtactcatggccacaaaggtttagaaattcatgtcacgtgtagcctgtaacttcaattcctgataactcaaaactatggtatttactagtccatatgttgaccttaaaggggtttactggtttcgttaaaaccctgttccc
cgattgtagtatatttaaaaagaaaacatgtgcgatactccactcctctgaaggttgagtctccactgctgctcccgatgctactatctacagccatccacaggttcacagacaataacaggagcaacggcggagactcagagctggacctggagagggtgagta
aagcaaactgcaaccgggtaacagggagtttctgaaattggaaaatctctttaaattaggaagaaatggtcatatactacgagagtcaaactataggtagttaattcatccaaaatctctgattgattaacaagatttgctccatgctccagtgagcagctccaacgtc
atgaaaattggaaagtcacaacatgcctgtaaaacgaactgcagcttcctgacgtgagaattcccagaacaaagaatgtttctcgactatatcggacaatgtcctctcccctaatcctgccccataatactatactttagtagtcaactaggcaagtgtacatactttgg
ttacccaacactgttataaaactggaattacaataagagttatcccatctggacaactgaagtccacatgatctactagggtatgtgtacactgcttgggaaccccttttattcactggagtataaagaggtatgaagagtttccacattggaacatgaccatttgacgg
gaagttgtgtaacactacatttcgataagccaacctcagcttccgctcatattgctaactgataactaaggtagccattatcatttgattggagatgcagtagactattgagatccggttgtcctttaaatcagggttccccaactccagtccttaaagcccaccaacagg
ccgtgttttcaggatttccttagtatggcccagatgataattccatcacctgaacaggcaataattctatcacctgtgtaatactgaggaaatcctgaagacatgaactgttggtgggccttgaagactggagttggggaacactgctttaaatatacagttcctggtggt
ccttctcagttttagactatagtatctcatctgatctagcatccttggaggagaacactaatgggatcagctattactgtattgacctcatcattaccaccatgtacatgactgatcctctcaaaaattgtcaagaaccctacatatgtccctaggactaaaggagatcctt
caataacagaagaaaaatatagagttatatctgtgcaaataaaaatatcttctcttttaggaacatggcatgggaagtccggcgtagccagacggattgaacatggaggtatgtgaatgagtgttaaagggaatttatcaattgcgtaaatatataattttgacctgttc

taaatgccattattctcctgaatctggagttgtttttctttagttcctgcacctctccatttctgaaatatgccccccttctatctatatatataaatctagttttctgagccaactgggtgtggtcctcaagaagacacccaaagagaacagttgaggactaagcctcaagggcat
aactatattgggtgcagggcttgctgttgcgtcctggtcccaatactataaaagaacagtcatagttgagggtcttgctggagattttgcattgggggacacggcttcaagtttcaccacgccctcttccgtagcaaaactagattcacttctggaaagagagggcca
tatctcaggaacggagaggcgtaggaacaaaagaaaaacatcaccggattcaggagaacagcgacatttacaccagatgaaaattacatatttatggcaagggataggtcctctttaaatgggttatccagtgatcacaagtgatcttgtctaaatggagtgaagg
ccaagcttgtaaagtctattcattgcttatggtgtttccaatggtattggactttctctggcaatcccatacacaatggattttgcaaaaatgtcattgcctgcattcagatggaactctacagaatctgtttttggatcacagtgtatttccagcagtcagaaccccagcgat
ctgtaagttatcccctgtcccaaaagtaagagctaagtgactatccttggaaatcccctttaagtactataccacagtgtgatcagtgggaatgaaaggatgtactgtaactcttctacaattattatttgtattatgtttattattaatactgttgccggtatccagtgcatat
gagcgttcccagaatcctcgcccctggctatgtccttctctgcataaataaatatccactgccatacagctatagtccaaggtgaacacagcatctactgcatcggtctattatattgtgttcattagagatattgtggtctaaggggtaacgtttctccttgtacctgacat
gccagtgtaaggaggcttataagccatgcaatgacattcttttgcgttgcctttttaacagttgatataatataatatattgatgccttcaatctttttagattctcctggaccccaccaaggtggaaggaaatccgtcatctgtatttgcaaccagaagaacggtcataatc
tttcaagtaaggacaaaactaacattttgggctcgattcattattgcatttgcacctttttatgtctagtcttaggtgtttttcatctgtttttcatttgctccttattcttctattaatttgcacctttttttaaatactattttagatgtggtcgcctggatttttttgttcaccatttggggg
cttccagatgtttccaattgatttttttaaaaagtttctaatttttcttgcaaatgaactccagtcctcctctttggtgtttttaatgcaaaacctgatttttttattttagcccaaattttgtgacattgtagtctaaggggtaatgttttttccttgtggagagtgacaagtcaagcctg
gcatgtcagagttaggagacttataagccatgcatgctcctctgggaaatgaaatatgcaaattgcctcttcagagaggaagaggactagaactctagtgccacctattggaagcagcgatcctaaaagtcaatatcgactttttaacgagccttgtcacatgactta
gataaaagccaaactagaatttcatgtaaaattttagtagaacttgtgactttttttgtgctaaaaacttgaagaaaaatataaagacaataataaagagcaatttttttgattttgcacaaaaatcataaacgaataccacaaaacaaaaagaaaaagtgactttgaaaaaa
aaaggtaaaagatgaatcgggcccttagaatattaaatgcatggccatctgacacataataatacatttccccctagtgtggctgctgaattgttggctgtagcctcaagagcttggcccttgataattagctcatgaccaagtggtcttaaaggataggtcatcaatatt
agataggtgggtgtgcaatatctgctacccccatcgatcagatgagaatattcaaacattcataggagctgttctgctgtacctggcagccgctgctacatattgaatggagctgcaccggcgctaataacagctgatcagtgggggtgctggtgtcagacctctac
agatttcatattgatgaccaattcttaggctccaataagagctaaccaattttggtgcaatttggcaaattttaagcctactctaagctacacctcttttttctaaccttttttaaactgtctccaaaggtataaaaagtgtctaaaaatataagaaatctggctcaagacattta
agaagagtttttttgtagttttgtaaatctatcttctttcttcattagatgtgtttttttttaaataatccgcaggaaaaaggcgccatatgagtgcacatcacaaaagtggtcctcatagccctgggtgtgccaggatctgcatgcggaaaagtggacgaggaggatccgta
cattccatacctttgtcatgaggaaacgttgaaaatttttctgtagaatccaggagtgtgaaaaatttacagtaactttttttttttcttaatgagccactccagtttacagcttacatacagatattatatatatataatcatataaagtcacattgtaggtagtttacatggcacatct
tagtctggattgtggcctatactatgctcagagctgcactcacaattctcctttgctttttttaagatagttttttgtcctgatgaagaaatgtataatgcttcgaaacgcgttgacaaaataaaaaagggtaaacccacttgtgtggacctggatttatcaattcgattagcag
cagcatggatcattcataattttcctttcccactttgtttgtatcactcagtggagcaattgaggcccccatacacatcaaacacatgttgttgaccaaacaatccgatttcggccaaccgcctctggtcttatgggttgtatggggtcctcctgactgcccccctgacaaat
gatgttgggtgacataatgatcggttggttccggttgaaattcaattattttgtacacatgacatggccatacagtgcctgttgtacaaatgaccctgcaatcactagaaaaacctctttttagagatagagccaaaaaagaatggtgagggattgctgctctgaagtttt
cagaactgttaatgcagctctggaatataacactgggatttataacattcaatttatactctcatttatatactacagagccatatataccacataaatgacaagggtgaaaatttcccagttaatataactccggaaaactaaaacttccaattttaataacaaaaagctgc
agtaaaaaaaaagattaaaggttcaaagagtaaagtgagctgttcagtaaaaccatttgtgttacatttttgtagtgcaggtttattgctctatctcacagtatttcttcatagtgttgtaatcaaccacggtcatattgttacttgcacatcttgttgtatttttacactattgcatg
agcacggcactaaactttatgagatattgtttgcattacaataaaaagtaaaaaattttgtacttgtttaattgtttttgtctctgattctatattttatgattatttatgatataatggtgcggcaacttgattctttggtttttggt

**C:** Sequence of cDNA (coding region in red and highlighted in yellow)

ttatttatctggagccaaggtgagagtacagcggctgtgagtggcagaaaacaaaagtatagagcttcccctgtgagaggagccaaagtgaagatgaaggcgaccacgatcttcatgttgtccgctgttctgatcctattctactgcttgaccactcaaggccgtc
acaattcaggaacatggcatgggaagtccggcgtagccagacggattgaacatggagattctcctggaccccaccaaggtggaaggaaatccgtcatctgtatttgcaaccagaagaacggtcataatctttcaagaaaaaggcgccatatgagtgcacatca
caaaagtggtcctcatagccctgggtgtgccaggatctgcatgcggaaaagtggacgaggaggatccgtacattccatacctttgtcatgaggaaacgttgaaaatttttctgtagaatccaggagtgtgaaaaatttacagtaactttttttttttcttaatgagccactc
cagtttacagcttacatacagatattatatatatataatcatataaagtcacattgtaggtagtttacatggcacatcttagtctggattgtggcctatactatgctcagagctgcactcacaattctcctttgctttttttaagatagttttttgtcctgatgaagaaatgtataatgct
tcgaaacgcgttgacaaaataaaaaagggtaaacccacttgtgtggacctggatttatcaattcgattagcagcagcatggatcattcataattttcctttcccactttgtttgtatcactcagtggagcaattgaggcccccatacacatcaaacacatgttgttgacca
aacaatccgatttcggccaaccgcctctggtcttatgggttgtatggggtcctcctgactgcccccctgacaaatgatgttgggtgacataatgatcggttggttccggttgaaattcaattattttgtacacatgacatggccatacagtgcctgttgtacaaatgaccct
gcaatcactagaaaaacctctttttagagatagagccaaaaaagaatggtgagggattgctgctctgaagttttcagaactgttaatgcagctctggaatataacactgggatttataacattcaatttatactctcatttatatactacagagccatatataccacataaat
gacaagggtgaaaatttcccagttaatataactccggaaaactaaaacttccaattttaataacaaaaagctgcagtaaaaaaaaagattaaaggttcaaagagtaaagtgagctgttcagtaaaaccatttgtgttacatttttgtagtgcaggtttattgctctatctcac
agtatttcttcatagtgttgtaatcaaccacggtcatattgttacttgcacatcttgttgtatttttacactattgcatgagcacggcactaaactttatgag

**D:** Amino acid sequence (signal peptide shaded)

MKATTIFMLSAVLILFYCLTTQGRHNSGTWHGKSGVARRIEHGDSPGPHQGGRKSVICICNQKNGHNLSRKRRHMSAHHKSGPHSPGCARICMRKSGRG
GSVHSIPLS

**Fig. S10.** Gene position and architecture (**A**), genomic DNA sequence (**B**), cDNA sequence (**C**), and encoded protein sequence (**D**) of the unannotated *cxcl17* gene in the NCBI reference genome of *Phyllobates terribilis*. The information was downloaded from the NCBI gene database (https://www.ncbi.nlm.nih.gov/gene).

**A:** Gene position and architecture

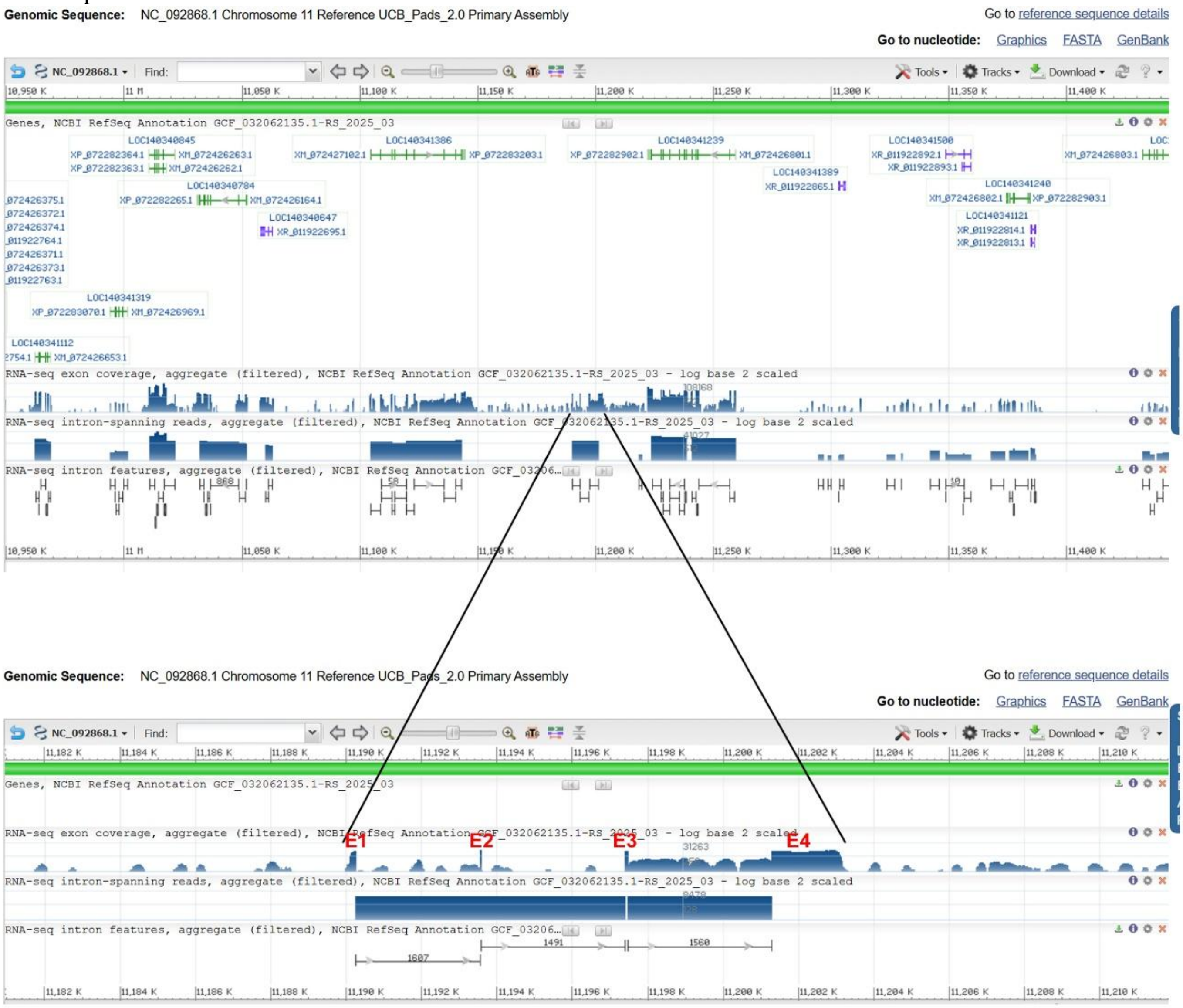


**B:** Sequence of genomic DNA (exons in red and highlighted in yellow, coding region underlined)

tggggggtaagtgatggacagatcatatcatctagggactttttgcctatggtgaaagagttaagatttggggggggggcacttgacattccatcagattttaaacctgttgtacggcccttccctgtgtgagtctgaagct
catgtaagagcaggacagagaaggaggcagcaaagagaaagtgctgctccccttacaagaatattgaaaccatgaagatagtcatgatttttctgtttgccgtggctctcaccctgttccactgctggaccgccgatg
ggcataaccacacaggtaattttaaatcaaaatcagaatgctgttaggagaagaaactgttgaccttagtaagtagaagaggctatctaaagtcctgaaagcccctctttgaccaaagacttttggtcatatttattgaagt
aaggaaacgggcaatgatacaatgtgacaacaaggaccggacacattcaaacaacactaggttcacgtcagtttgattaaaaaaactgtcagtttctgattggttgctatcagttgactaaaggcaatcaaacaacttct
ggtctactgcagttgggtgaaaaaatgctaaatgctgattgatttgagttgacgcctcagtgttatcttgactgagcatgactatttagataattgctaatacagaattgttaggaaaacagaaggactttgtaaaatggaat
ttatttaatcagattaatattagcaaataggcagtgatataaagtgaatatctgaggaacatacacgtaaggcaaatatttgtagacaattgtagcattcaatgatactttttttctctttttgtctgttggacatttttgagtgga
attttgtctaacataaaatgttataaaggtcaaggagagtcctaaaccttttttctcatacaacttttttgcctttaacatacacagtttcagctagggttctgcagccatggacatatactgtgtaactccaccatgatgctgcca
ttgctaatcatctcaaaatattagttttgttggacaaccaattattggcatttataacacatggatctaaaaattctctaccagaaaacgttgctaaaagtcagattcactgctttttattaggacttctcagaatgttgtaaactc
atttcatctttaaatattgaagcaactggattagcaggtcgagaagacaatgtactttcaaaaatatttgaatggataaagatttatcctatagcaggatttatttaaagtatatgtccaaccttatccttttaacttagaatgagg
aagaactataggccttatacgatttgaagcatgtaaataagagcagaaggaatattttgttattttatatttttattaggctatttatttatatcttgtattatatttccacataggatatttcccgcattgatggtaaaatatatataaa
tatatattaaaaatattatattatatatttaaatatattttagctccatgtacacaaaggaaaatcatagagctcaagtccaaattcataatagtaatgtcactgccttaaatgattcacaatggctcagaattgatatgattgaga
aaaagctgggaaaaattaaggttgacaaagcaccaggaaccgatggattacatccatgtgtcctcaaaaagctgagctcagttagtttaataattatttacaaattattacaaatacaagagactttttaatcactggcaag
gtaccgatggattggcgtaaggccaatgtggtttctatcttcaaaaaggtagcaaagtcattaccagttaactacagaaccattagtttaacatccatagttggaaaggtcctagagagtttgataaagaaccagataga
ggagtttcggctagaatataatattataagtgatagtcagcatggcttcaagaaagaccaaagttgtcaaacaaatttactctctttttatgaggaagtaagtaaacaggtagacagtggagtagcagttgatatagcgta
cttggactttgctaaagcatttgacactgtaccccacagacggtaatacgcacaatagggtcaattggttaataaaagtcaatctgtaaatggatagataactagcttaaagaccatatccggagagttgtaattattgatt
aatactctgaatggtctaaggttattagtggttcagtgttgggacctttattatttaacatcttcataaatgatatagagtttgtgactaaaagtaccatttctgtgtttgcagatgacaccaaactatgtaatagaattaagtcca
tacaggatgtctataatctacaagcagacctggatgtactgtttgattgggcagccaagtggcaaatgacattcaacatagataaatatacagtttttcatttgggggctaacaacatgcatgcttcatactgtctagtggg
aatacatttgtgggagtcagaaatggaaaaggatctgggggttctggtagatcatagacttaataacagcatgcaatgccaagctgcaatatctaaagctagtaaagtactttcttgtattaaaagtggaatagactgca
gactgcacactggagacataatcctagctgaactgaatctattctcccttgagaagagacgtttagggggatatgatcaccctgtataaatatataaatggtccgtatagagaactctcttccaaattttttacattgagatc
attacaaagaacaagagggcactctttgtgtctggaggaaaagaagtttaagctccggataaggaagggactcttcactgtaaggtctgtgaaaatgtgaagtttggctccctcaggaagtagtttcagcaaatactat
agattgctttaggaaaaaaatggatgcttttctggaaacacagactacaactggctattaggtctttaaagtaaaaatattagtgactttatgcagggaacacccgattgcctcatggaatcaggaaggattttgtttcccc
aatttgtaccagggtttttttgccttcttctggaacaactatgtcttatatggttttatatctgggatatgtttatttccctggtggttgaacttgatggattatagtcttttgttaacctaacctactatgtaactatgtaactatattata
tattatgatcatataaaatcttgttattgttgtttggagttctgctttaggactgggtttgcacactatgtagtgaaatgcagaaatgaatgaacaagtctgagctgattcatcatccagtggtgcatgactgacaaactcagtc
actgttgaggcaacactgtcattggaaaagaatttaggctttatttattcaagtgtgtagggagataaaatgtttttgtttcctaaacttccccatcaagttcagggcaatataattgtttgtgttatacactggacagtattatct

aaaagtaatgggattaaaacagagagttaaaagattatcaattaaggtccaatgcaggtgatattaccatatagagaaatgaacttgttctcttctgggagagcccaagagaagtactaaacactgtgttgtttttgtttgc
caggtaaaagaggtctacagaggaagacacaccacagaggtaagttaaaaatggaacactgcctaggattttaaggtataaagtatttgatgtaccagtatttattgtcataatgctgtcaggctttacatatttatataga
ttttttataataattcttagctgtagatctattaccagtgctgttgattgaagaaatgttcaaaggtggcggtatgtttgtgcatgtattgggggggggggagtaagtgggattgtgccaaacctcttgtgatccaggctgttgct
tgcttgtaagagtagcggtcagattctccaaggtgctgccaccatattttgtctttgcaacatttattaataattatattacgaaaaagaaaaaaacaacaaaaaaagcaaacctttttttcattttgtatgtgagccccttttgaa
ttacagtaaacataatgagggtccaatggggaacactggccctgtaataaatccataatatgcactttgttgacctttgtgatttttttttttactttaccacaatgcctgcccagacatgtctatgtaagcggggaggtagaca
gttaaataaagcaatatatatcttctcaacgacacttccatgacaatgcaatggtagataatgagagttgtgaataaagattagttgagataaaatatttagaaatagaaaataatgaagcatattgaatgaacatgaccat
cattgccttgttaatgatcaatgtagtccaataatgcacatatagatatagataatgcaaatcatgaaaatgtactctacaaaagataactttgttagacataatgaaactgtacatcctaacttgttgtgacaagaatgagat
gcaccatttagcataaggtatgtagccaaagaatacaaatagtttaaatgacctgcttggatatcttacagctaaacacagtgttgcagaacaataccattctcactgaaaactcaacaccataacctggcagcacatatt
ttgctgtggaatttatatgttttccacaattagttggcttacatgcaacagcagctccatctaatacactttagttaggatatgtgtagattaatattagaagttattattccaaatagtaccactgtagttcatgactttggaactt
gtactaaaaatcctaaagccctgtatataattgaaacaggtgtgagtagtccaaaactctttttgacatttttttgagaatcacttattgaaaataaagaaaccacaggtagcgcagtgttgttaagttagccagtacattttaat
aaaggaaaattaaacaaaatatactacataaaaaagagtgcacaaagtgtttttttactttagggattaagatactgcagatgacagcagtgtaaaattggccagcagcagaacaaatttttaatagcaaggctaggtcca
aattgacaggaaacaaaagctcatgctgaggatttactacaaaaataaagacactgttaagtttatagtatcacatttagggtgaggatgctacaagtacatataaaggatctaatctgtttttctatcttttatttattctaagg
gcattgggattacactctgtttttgcacatttcgcacaaacctgtctgcttgacctaattccctctgatctactctgtgcccattcctccaccctcgcccccgctctaaccaaactattcaacctctcgctttctactggtatcta
cccctcaactttcaaacatgccattattctccctatcctgaagaaaccctcgctagacctctccctaccttgtagctcctgtcctatctcccttctcccatatgttttcaaacttcctgagcaccttgtctacaaaagactcacc
aattacctgagcaccaactctctccttaaccctctccagtctggttttcgagctgcccattccctgaaacagcccttactaaagtggccagtgacctcatcagagctaagcctcaaggctgctttccccaccttctacttga
tctttcctctgcttttgacactgttgatcacccccttctaatacaaatcatgcgctccattggtatcagggaaactgatctctctggtttgctttctatctgtctgatcgctcctttcaagtttctttcaatggtaactcctccttgcc
ccctctcctccttgttggtgttccccaaaggtcagtccttggactggttccctttttctctgtacacctcctctctcggtagtctcatatcctcctttggcttacagtacgctgtatgctgatgacactcaaatttatctgtccactc
ctgacttttctccttcagccctggataaggtctcatgctgcctgttagccatatcatcatggatggctgacagatttctgaaactccacctggataaaacaaaactcatcatcttcccctctcaaattccaaattaccccctga
catattcctaactgttattcgtccctcacctctggcatgttgtcttggtgtcacctttaaatctgccctctcatttacccccctatattcaaaacatttccaggtcctgtcactttcatctacgcaacatctccaaaatctgcccctac
ctgttccctgagaccaccaaactccttgtaaatgctcttatcatttcttgcctggactactgtaacctcctcctttctggtattccactaactcaattctctcctctacaaactaatatgaatgctgcagccagactcatccatcc
ttcccaccgctcctcttctgctgcatctcttcatcgatgtccatttcacctttccccacatacctttctgacctgatagaaaaatactcccctagccgctctcttctctcttccattgacctactaatgacgttctcactcataacct
catcacacacaccggtccaagacttttctagagctggcccgactctctggagtggtcttcttcgtcctatttaccttgctcccactttctgctcatttaacagatcactcaaaacccatttttttcttacttgcctacccacctgtc
ttttgaaaccgtcactacttcccactattccatatctcccctcctattgtgtgttacttcccttacctcctagattgtaagctcttcagagcagggtcctctcctcctgtgtcactgtctgtatttttaacccctatttaatgtacagc
gtaatatgttggcactatataaatcctgtttttttatttttattattattattattattattaataataataattttagctagttacaacatgaaataaaaagataaaaagaattaaaatccccaaaaaacattgctaaaacaaaagtcta
tgcatgtaaagaaaaccttgcaaaagccgtcgcaaagcaaacttatggatagtccacaaaaaacaaaacaaaaaaataaagcaataggactgattgccaaaaagttttctaaatgtccctaatcagcattccttatccttc
ccttatgatatccttacctacaactacagtaagattcttttttctccccttcctataaagttaaaggactgatttatttaaaagcccccccttgttggcctttctaaggctgctgtaaatagttcaagggtcatacattgtatattgca
tacgaaaatgtaatgggcagttttgcacataaattcatcagggctgcaaattgtgactctccatgattttacttgaagccatggtagtggcaaacttcgcctttagtaatatgtgctacacaaagaaaatatattatgtgctttt
atcatgtttcaaattatttgattttcatattgtatatttttcatctctttgcagggcattatgtggtacctcattctaatgagaaaaggcttcttgaatgtacatgtgtgccaaggaaccctcatcatttacaaagtaagtaaaggttt
atagcacattctttctaactgccaatttaaaatatggacgaaaagctgttttattatatgtgtgtaggggcttgtgtcgatatttaattcacattttcatacaggtctgttagaattaaattagatcaaatgaagggtagaatgag
gtcaactacaggttaaatgcacctgtcaatgaattcggaagtatttcaaacaaatgtcaagcagatgaatactaatcatgggccccaacggtatttatttttagcaaatgatgctacattcctgacaaattgaaaaatattca
catatttatataaaaacacaggacatagcctcattcacaatatacataaaatgccagcaaactttattcaatcttatccattaaaagaagaactgataaaactaacatacaaaaggcaaaaatgaatctacagataagtctc
aaatcatcatgactcgtttcacagacatcttaagctgtgtacacacttccaatttttatcgttggaaatgaacgacgaacgaacgacgaacgatcgattgggcaaaaatcgttcgtaaaaaaagtaaccaacgacgccg
acgaacgaggatagtcgttggaaatgaacgaccggaccggcggatcggattggacgacgatcgttgaccatctatcgtgtgtatggtcgttcatttatcgtccatggtctgagcatgcgcggtgaacgaacgttcgct
cacttctgtggttcacgtcacttcctgtatcgttcaaacgatcgcatctatcatgtgtacaatatctgtgaacgatcgtgtcgttatctgtatgtacaggatcggtgctatacgattgttcacagatatcgtgcaggatcgttcg
tcgttcgtttaccaacgataataattggaagtgtgtacgtagctttagtttcatcaagagcattaagctctcctatttggcaagaagccatttcttcctcagagaatctcgggaaccaggctgtgatccatacttcaataaagt
aattgaataaaggaggtgactatctcccccacccacaataaagttttggcagtttgcctttcagtttgctttccagtttttacctttccctgtatggaacaatgaagccaggaactgaagaatgtccacaggatcagatggg
tactggttatttggtttttaaggtctcattcaatactcctgatgaagcggacagtgacgtctgtaaaacgcattgaggtatgaaaatttgaaacctttctgtcaatgaattcctgtatttttatgttaatttaaatgaattgagag
aaaagaaagaggtttgaggctcataggtttttcatgcaaaaagtgaattgtattaattgattattcagtacataagccatcaaatacacataaacatatattattttcatctctgtcctgttactatttcacagcacagagtagtt
atacatactgtatagaatggattttgtcattaagtattacatataaccataaaactgttcaaaaagccacgccgtggcacgcagtgatccgatgcgtttcgccacgaagggtttcctcaggtaaagatagtagtgcatgctt
atcggtccgaagttaggaacatgccccacgggggggggggggggggtctggtgtatatatatagttgctgctgatgataatgctaaagtttgaagtagtaatctatactcttgttagaggttttagcacaatttacaaatatt
tcatggatgcaatgaagttggtccctgcatccgtcgaccaatgtaataaataaatgaggagaagggtctccccttagaacttttggatttgcagaaataagcagtagggggatgaaaaagcagcagcctgctgttaaa
cccgaccttggttcggggtaaaaacacttgcaaaaagtgttaaacccgaacttttctcaggtgggtatacaaacattatattgcaatccgattatcatacattgtatgacaatcggattacaattgaaagaatatacttaccc
tttccccgcagctcctctggacacatctgtcctcttctgtcttctcccggcgagtgcagtgacgatctccggggtttcctggtgacgtcgctgcatgcgtcgctgcgggaggcggggcgggaaattcaaataactttgta
ttgagctcaatactaaaaagctgtattgagtccaaaacaaagaaatctttatataatatatatatataattgtattatatatatatattatataagctactgtacaattactgtacattatacactatttttttttttttttttaaagattttttttt
gttttataaaaaaaatttattattaaatttataaaattttggacatatttcggtgagttatgtggtaattagatgaattataagtaattattgagtaattctaggaattatagcctacaatctaaaatacatttccatgccaaaaacgta
acactttttgcatggaagtacagaagttcggaaagaattagaatacccggcattcgcgtacgcgtccctcggcaattcctgatgatgtccgtgcatgcgcccgtcgatgggggtcgtagcgggaaattcaaatattttgt
attggattgaatacaaagtcctgtatccagtccaatacaaaataatacaaaatatatttatgtggttttgtctataggtatgtgacgttggacggttggacactagggaggtgttttagaaaaatatattactatacagtatacc
gaattattgcattttcagtatttttgatttatttacacattcttgtttaagctgatttttgtgtttttttatttaattttattaaaagtaatttttttttttttttacatgattgtgtgtttttttaaacgtttttttatattcatgatatctactagaacccttg
tttatgtgtatttgatggcttatgtacttaataatcaatcaatacaattcactttttgcatgaaaaacctatgagcctcaaacctctttcttttctctctattcatttataccttaataccatatagaggtggctcaacacaaataacac
attgatatagcgaggaggagtaaaaccctttctatactaatttgtttatgttagttttatcagtttttcttttaattgaaaagattgaataaagtttccaggcagtttatataatttgtgtaagaatatatgtaaatttggtacctgaaa
agttcagttgtcttgtttcaaatttacacagaccctgaatagtgatggcaaaaattgtgtgccaaaaattgtaaaaaaggacaagttaaatgaagtctttttttggcttctcatttagcatcatgatagtagttggacagaaata
aaaaaaacaaaaaaaaaccaaagaaatataatataacaaaaacatttttaaatttctaatgaaagttttttttttaaatccccaaggagacactaacccgttgcttcttcaggccaatgtctgaagacacccctcccccccagg
aagctccatttatctattcacattcaaagcagcctctgaacccctccagagcagatttttgggtaggaggaaataccatgggtaggaggatgccctatataagtaccaacaccaaccacaccgtagcttagcttgacta
gcatggtgcagcaaggaggcctcaaggaagcttaacttgaatgtatacaggtttttgaaggggaatatacccatatatgacttatggacctaaattttatttagggaaaacaatgttcctgttctgccaggttattgccttatt
ttcctttgaaaccccaatatcgatgttttgtgaataatcttctatgtattgcaggaaaaagacgtcatatgagcggagctcagaagcgtaatcatcgcggacgcagatgtcaatgcaaacaatcaaagcgaaattctgag
ctctccataccattatcctgagtcttccttcctgcacattacagcagcagagaacatttttaattttcagtgtcatataaccatctttccaaagatttattttcatgctgcgcttgtctaagcctgtgtctgttagggtcgccctattt
acacagaacatttgcagtgggcctagacaatgatggatagataaaggcccttttgcacctgactcaaatcaagtaaaaaagactcctttaatccacatgtctgcacagcctttaaaaatggaattttaccctggttttgcct
ggtctagaatcttgtaaaaataataaacaaggctataggaaataagtttattattattaagtgcctcttgtaaatcagaccctgtttattctgtgtctttgattctgcaccatctggagatcttccccacagtacttctttttgccac
tagatgtcctcagcctaatagccaacaaaatttaacccacttcctccgagcctaggttgccctgaactcagccttgactgttgctaaacattaaaaagagaaacatcaatgtaatgagttcagctttcagtcaattgtatttc
tctccactattatgcttcttaacagcacatgaattgatttactttttttctctgcaggtaattcaaaaggcagatacagagttagatacttacactgcttgtgtttaaaaatgcatttaatcatctactaagttcagatttcaaagaaa
cttgggcattacattgagtttactttatctagttatacaaaaaaatgtaccctaaatgaaaatgttactgtataccctgatacttacccatatgtgaaatgaattgtgacaaaatacttattaatgaccatcatgagtaacactaatt
gatggaaaatcaaatcctcaacttaggatgtaactattgtaggtgaaacataaactaggtgtctcatttaaaaatatagcttgaaaagacaagaaattggcaaaaactgtcctgccattggatttgtatctctgtccattcag
tcctgagatttttttacaacactggaaggccgatgatcttgttctgacaaaatgacttaccttgttaagatgccctacctccaatactaaaatccccaccaccacctaaccctaactttttctgtctgatgcctctgcttaatctct
attctttgtgcctaatgttgacccatttcctgctgcctaacacccctaattccttatatatataccttactttatttgtgccccacctttacatattttggcagaagctaacctatccaaatcattgaaacaccaaccctacagaca
tctgtgtaacaggtaggttcaaagagtgggctattttggcagaacagaacctgagtttcctctacggtggttaaacaatgcatctagattatctcccaattccagcctgtgactggacagttagaatgtacaagagactga
gatcattattggcatatgtgaatactgaacacctagttggtccttgttcaataattatgttaattggtgttttctttttttatttttatcaaatagaacataactgagaaatattgtggtaacacatattagccacatgactgtcagcttt
tcaggatactccatcctaaatctcattggctgttaataaatgacctttgtatttttctgacaaccaaatgcttatgctttacattttccccctatttaactgtacctgtataccagatataagcttatttttatactattttataataaaaa
tcctgatcgtacaaatggtgctttgtgctgcaattcgtttgaaaaatatgtaccctctggctctgaacatgagcaagtcataccctattgcttggtttggcctaaattttttttagaaaca

**C:** Sequence of cDNA (coding region in red and highlighted in yellow)

agcaggacagagaaggaggcagcaaagagaaagtgctgctccccttacaagaatattgaaaccatgaagatagtcatgatttttctgtttgccgtggctctcaccctgttccactgctggaccgccgatgggcataaccacacaggtaaaagaggtctacagag
gaagacacaccacagagggcattatgtggtacctcattctaatgagaaaaggcttcttgaatgtacatgtgtgccaaggaaccctcatcatttacaaagaaaaagacgtcatatgagcggagctcagaagcgtaatcatcgcggacgcagatgtcaatgcaaaca
atcaaagcgaaattctgagctctccataccattatcctgagtcttccttcctgcacattacagcagcagagaacatttttaattttcagtgtcatataaccatctttccaaagatttattttcatgctgcgcttgtctaagcctgtgtctgttagggtcgccctatttacacagaa
catttgcagtgggcctagacaatgatggatagataaaggcccttttgcacctgactcaaatcaagtaaaaaagactcctttaatccacatgtctgcacagcctttaaaaatggaattttaccctggttttgcctggtctagaatcttgtaaaaataataaacaaggctata
ggaaataagtttattattattaagtgcctcttgtaaatcagaccctgtttattctgtgtctttgattctgcaccatctggagatcttccccacagtacttctttttgccactagatgtcctcagcctaatagccaacaaaatttaacccacttcctccgagcctaggttgccctga
actcagccttgactgttgctaaacattaaaaagagaaacatcaatgtaatgagttcagctttcagtcaattgtatttctctccactattatgcttcttaacagcacatgaattgatttactttttttctctgcaggtaattcaaaaggcagatacagagttagatacttacactgct
tgtgtttaaaaatgcatttaatcatctactaagttcagatttcaaagaaacttgggcattacattgagtttactttatctagttatacaaaaaatgtaccctaaatgaaaatgttactgtataccctgatacttacccatatgtgaaatgaattgtgacaaaatacttattaatgac
catcatgagtaacactaattgatggaaaatcaaatcctcaacttaggatgtaactattgtaggtgaaacataaactaggtgtctcatttaaaaatatagcttgaaaagacaagaaattggcaaaaactgtcctgccattggatttgtatctctgtccattcagtcctgagat
tttttacaacactggaaggccgatgatcttgttctgacaaaatgacttaccttgttaagatgccctacctccaatactaaaatccccaccaccacctaaccctaactttttctgtctgatgcctctgcttaatctctattctttgtgcctaatgttgacccatttcctgctgccta
acacccctaattccttatatatataccttactttatttgtgccccacctttacatattttggcagaagctaacctatccaaatcattgaaacaccaaccctacagacatctgtgtaacaggtaggttcaaagagtgggctattttggcagaacagaacctgagtttcctctac
ggtggttaaacaatgcatctagattatctcccaattccagcctgtgactggacagttagaatgtacaagagactgagatcattattggcatatgtgaatactgaacacctagttggtccttgttcaataattatgttaattggtgttttctttttattttttatcaaatagaacata
actgagaaatattgtggtaacacatattagccacatgactgtcagcttttcaggatactccatcctaaatctcattggctgttaataaatgacctttgtatttttctgacaaccaaatgcttatgctttacattttccccctatttaactgtacctgtataccagatataagcttatt
tttatactattttataataaaaatcctgatcgtacaaa

**D:** Amino acid sequence (signal peptide shaded)

MKIVMIFLFAVALTLFHCWTADGHNHTGKRGLQRKTHHRGHYVVPHSNEKRLLECTCVPRNPHHLQRKRRHMSGAQKRNHRGRRCQCKQSKRNSELSIP
LS

**Fig. S11.** Gene position and architecture (**A**), genomic DNA sequence (**B**), cDNA sequence (**C**), and encoded protein sequence (**D**) of the unannotated *cxcl17* gene in the NCBI reference genome of *Pyxicephalus adspersus*. The information was downloaded from the NCBI gene database (https://www.ncbi.nlm.nih.gov/gene).

**A:** Gene position and architecture

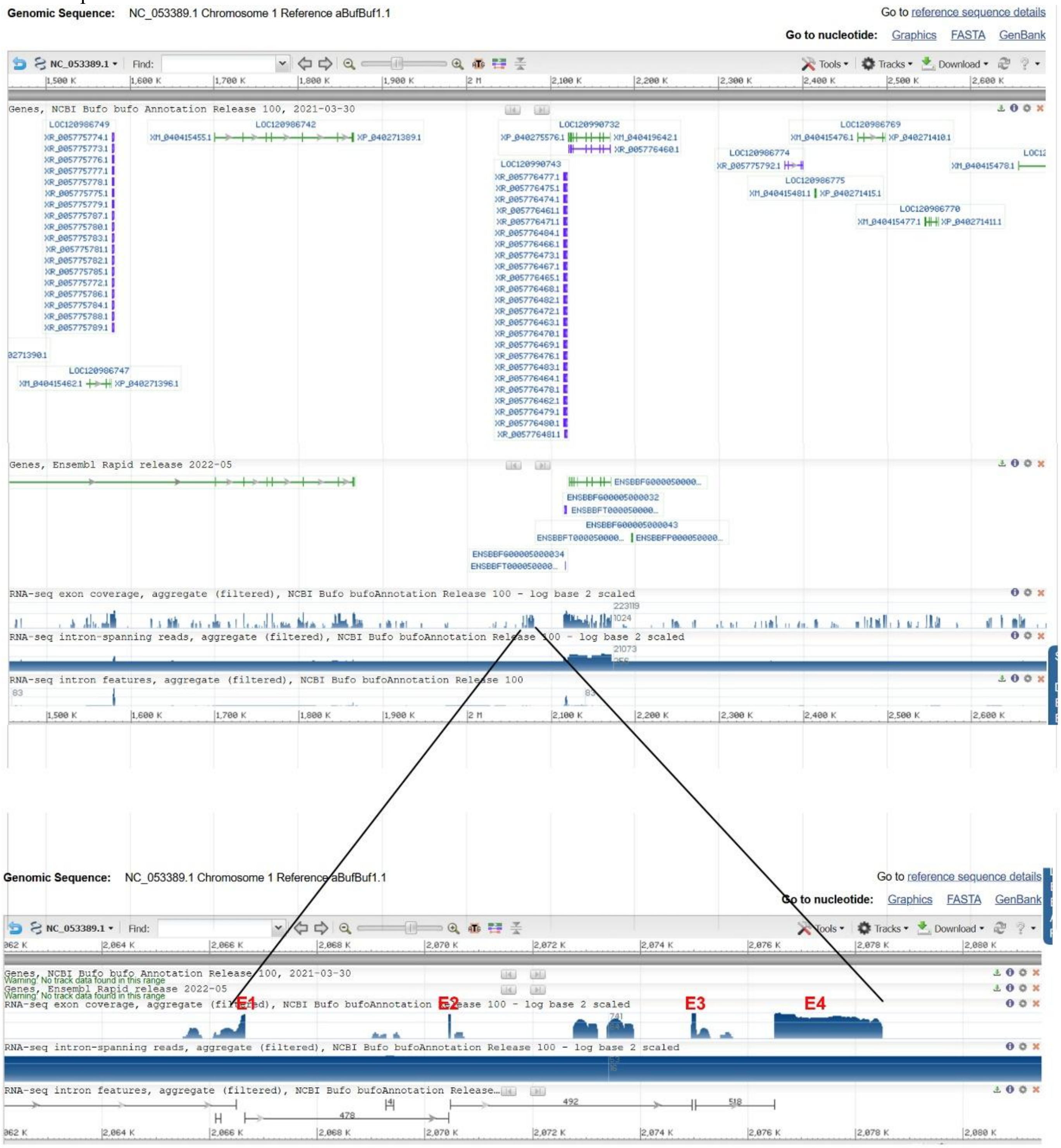


**B:** Sequence of genomic DNA (exons in red and highlighted in yellow, coding region underlined)

cccttatatatctgcagcctatgtgagtccacggaggttgtgaaaggcagacaacaaagaatgagagcgtctcttgtaagccaaggtgaagatgaaggcggccgcgatttttatgttgtccgcagctctgatccttgttttctgcttcaccactcaagggcgtaaaag
ttcaggtaacttgtaccaggagaatccaaacagcatgggctctaggagatacaaagtactgcatgggatcatggagcttacaaatattaatctatgggatctataggacactgtatgattatgggatctataggacactatgattatgggatctataggacactatatg
actccatgggatctataggaaactataggactctatgggatctgtaggacactataagactatgggatctataggacactatatgactctatggcatctatatgaaactatatgactccatgggatctataggagactatatgattatgggatctataggagactatatga
ctctatgggatctataggacactatattactccatgggatctagaggacactgtatgactctatgggatctataagacactatatgactatgggatctataggaaactatatgacttgatgggattcataggacattatatgtttctatgggatttataggacactatatgac
tctatgggatctataggacactatatggcttgatgggatccataggacactatatgactctatgggatttataggacactatatgacttgatgggatccataggacactatatgactctatgggatctataggacattgtggacattgcatgattctgtaggatttctaggg
tactatatgactctataggatctcttagacattatatgattatatgatatctataggacactatatgactctatggcatctataggacattatatgattccataggacttataggacattatatcactctatggcatctataggacattaaatgattctataggatttctaagacac
tattggggtcatttatcaatatggtgttgcccatagcaaccaatcagattctcactttcattttttcacaactttggggaatgatcccattcaatgggatccatatgacactatggtatctatatcacatatgattatatgtgatttataagacactatatgactctattgggattta
tagggcattatatgattcaataggacttataggacattctataactctatgaaatctataggacaatatatgatactataggatttctaagacattatatgactctatgggatgtaatggacattatattattctatgggatctataggacactatatgactctatgaaacctata
ggacattacatcatttcataggacataggaaattatatcactctatgtgatgtataggacactatatgactctatggcatctatagaacattatataattctataggagttataggacattatatgagtctatggaatctatagaacactgtatgatactataggatttctagga
cactatatgacactatgggatctatgggacattatatgattccataggacttatgcaacgttatatgactctatgggatatactgtataggacattatgtgactctacgggatatataggattctatataactcgatgggatctataggacaatatatgactctatatgatcta
tgggacattttatctccttccatcaattctctccagatttagttctaagagaaattccatcccatttccttccatcaaccatcactagcctcttattataagaccttccatcccatctccctccatcaaccctctccagcctcttattataagagaccttccatcccatctatccca
tctacctccaacaaccatctccagcctcttattataagagaccttccatcccatctgttacgaacgttggtagttgtagtaccttgttcacctattcctcacaagctgctgatcctcagggattatagttgcagttcgggtgtggataggaatggttctagacgaatgcagc
ttccaagtagttcctcccccagccagtagatagttacccaaataggagcccagacttcatgaagggtacaacaggaacctttaatggggctacatgtcagctttttatgcaggtcttccagcaagggacactcccatggggaccttgtggaagactgagacagtttt
ttacataaaccaattacatgcagttacaggcagaaaacacacacatgtgatacaattaactaacacaatgggataacgtaatcactcacaggcagaaactacacacttttccctttgtagaataatgagctgggggtggggtggtcgttggtcggtatattcctctac
cgaaaacataacatggcctcacaccatctccctccatcaaccgtctccatcctcttattataaggacatccatctcatttccctccatcaacccttttccagcatcttattataagatagaacttccattccatctccctccatcaaccatctccagcctctgattatatgacctt

ccgtcccatctccctccatcaaccctctcaagtttcttgttctaagagaccttccatcccatctccctccatcaaccctatccagcctcctattataagaccttccatcccatctctctccatcaatcctctccagactcttattataagaaagaccttccataccatctccttc
catcaaccatctccagcctctgattataagaccttccatcccatctccctccattaaccctttccagcttcttgttctaagagaccttcaatcccatgtccctctctcaaccttatccagcctctgattataagaccttccatctcatcaaccctctctagcctcttatgataaga
cctttcatcccatctccctccatcagcctcttataagactttccatatcatctccttccatcaaccctatccagcctcctattataagacctttcatcccatctctctccatcaatcctctccagcctcttcttctaagagaccttaaatccccccctccatcaaccctctccagac
tctgattataagaccttccatcccatctccctctatcaaccctctccagcctcttattataagaaagaccttccatgccatctccttccatcaaccctcttcagcctcttattataaagagaccttctatcccatcttcctccatcaaccctctccagctctttttcgagagatttt
ccatctctgaagtgtggagctcaaaggggtttgcccatcttggacatttggggcatatgtctaagatatgaccccaatatcagataggcgggggtgccagatctgggaaccacacatattttcagaatggagggcaaaattcaggagggcgcactgcacatgctc
cgccgtcctccattcattcctatgtgaagtcctattgaaatgaataggaagtgccaacgctcagctatttttggtgccccccttaggaataaatggtggcgggctgcgcatgcaacccctggcactttgcgggctccattctaagtataggtgcgagtcccagaggtg
gtacccatagctgtcagacattgagtgcatattctagtgatatgaccccaatgtcgaagatgggaacaacccatttaagccttaatcccgtattggagatatcctagttcaccagtgagaacatgtctgggcccatttccttaatctacagtccatggtgatctagtgga
tgtaaaccagaccctttttgtttaagacatcctggtctgcatcattagatgagaagaacaccatatgggatcaaccatctctctacagacccgttatagcagggcatgaatagagagctgtaccttttttttcccaattacaaactattctatcttttaggaagaacttccag
catagtcaagaggactgaccatggaggtaatacaatgagcattaaagactatagacacatttggaggcatttttttataattaattatttacatctattttgggctaaaatgtttttttgaatttttttaataaacaaatgttgctttggtttctgcagcacatttatttcataacac
ctagcagtccacagaaaaccactctgtgatgaatctgccagactggcactctgtcaggtccacagcccttcactcttttctactggctggcagttggatggctttgtttctagttcccctctcttctctactgctgacttgacaactcagtctcctgaaactctgtttgctat
ggggacctatttgacatattaggctccagcctttctctctcttctagctagctgtgtgatggcttggtctccagtgcctctctgttcttcactggtgggcttccagacagcttggtctccctcccaaccctttcttcactattggttgactgatgtcttggtatctagccctcctc
tttcctctgataactggctgtctgacagttcagtatccagccccctgtcttctctgttgagtagctgtttattggcttggtctccagcccctctctcttctctattggctgctatgtaatgccttggtatccagctctctgtcttctctgttaagtagttgtttattggcttggtctcc
agcccccagtctggctgtttattagttttgtctccagccatgtactctgctataccatgtgtctgatggtttggtttctagccattctcttctctactggcttactattgattagcttagtctttagcagctctctctagtatttgtctatctgacagcttgatctccccctcttacttttct
attcactgtcactgtattggctcagtctctatctcctctccctctctctcctttagtagctggctgtctgatggctcactttccagtccttctctattctttactggctggctgtctattagctcagtctccagtttctcagttattgtctaacattttactctcctgcccatatctcttct
atattagctggatatctgacagtttggtctccaggccctatcttctctctaatgactgtctggcagcacagtctccagttcctccctcttttctgctagcggtctaatgacttggtctccagcctctatctcctcaactagctgtctgcctgtctgctgatttagcctgtcgtcct
tctcacttctttactgattggttctatgatggcttggtctccagcatctttcttacctcaactgcctggatgtcttatagcttggtgtccagcctctctgtcttctctaatagtaggctgtctgacagcttggtctctatccttcttcaatgtctgtgtgatggcttggtgttcatttcc
tatcagttctatatttgctggatgtgtgatggctcagtctatagcccttctcttttttttactggctggctgcattttcgttcaggctccagtctatcccttctatagtagctggctgttgagtggctcactatcctgcacctctctctcatctactagatggaggtctacaacttg
gttgtcaaaccctcccttttatataatagccttctgtctgactgctcagtctccagtcccaatctcttctatacttgctgatggcttggtctccagctcctttactagtgtgctgtctaatggctcagtctacagcacctagctcttttacactgtctggatgtctattgactgagt
ctccagatcttctctctcctctagtagctggatgtctgaggaatcagtctccatcccctatatattctgtactagctggatgtctgagtgattggtctctagcccctatctcttctatactagctggatgtctgagggatcagtctccaaccccctatctttctatactagctgg
atgtctaaagagataagtctccagcccctatttctcctattgtagctggatgtctgagggatcagtctccaggacctatttatcctctagtagctggatgtctgagggatcagtctccagcccctatctcttctatactagctggatgtctgagggatcagtctccagccc
ctatttcttctatactagctggatgtctgagggatcagtctccagcctctatctcttctatactagctggatgtctgagggatcattctctagcccttatatctcctatactagctggatgtctgaaggatcagtctccagtcccaatttctcctctagtagctggatgtctgag
gaatcagtctccagcccctatctcttctatactagctggatgtctgagggatcagtctccagcccctatctcttctatactagctggatgtctgagggatcagtctccagcccctatctcttctatactagctggatgtctgagggatcagtctccagcccctatctcttcta
tactagctggatgtctgagggatcagtctccagcccctatctcttctatactagctggatgtctgagggatcagtctccagcccctatctcttctatactagctgtatgtctgagggatcagtctccagcccctatctcttctatactagctggatgtctgagggatcagtc
tccagcccctatctcttctatactagctggatgtctgagggatcagtctccagcccctatctcttctatactagctggatgtctgagggatcagtctccagcccctatctctcctctgatagctggatgtctgagggatcagtctccagcccctatctcttctatactagctg
gatgtctgagggatcagtccccaccctctatctcttctatacaagctggatgtctgaaggatcagtctccagtcccaatttctcctctagtagctggatgtctgaggaatcagtctccagcccctatctcttctatactagctggatgtctgagggatcagtctccagccc
ctatctcttctatactagctggatgtctgagggatcagtctccagcccctatctcttctatactagctggatgtctgagggatcagtctccagcccctatctcttctatactagctggatgtctgagggatcagtctccagcccctatctcttctatactagctggatgtctga
gggatcagtctccagcccctatctcttctatactagctggatgtctgagggatcagtctccagcccctatctcttctatactagctgtatgtctgagggatcagtctccagcccctatctcttctatactagctggatgtctgagggatcagtctccagcccctatctcttct
atactagctggatgtctgagggatcagtctccagcccctatctcttctatactagctggatgtctgagggatcagtctccagcccctatctctcctctgatagctggatgtctgagggatcagtctccagcccctatctcttctatactagctggatgtctgagggatgtc
tccagcccatatctcttctatactagctggatgtctgagggatcagtctccagcccctatttcttctatacaagctggatgtctgagggatcagtctccagcctctatctcttctatactagctggatgtctgagggatcagtctccagcccctatttcttctatacaagctg
gatgtctgagggatcagtccccagcccctatttcttctatactagctggatgtctgagggatcagtctccagcccctatctctcctctagtagctggatgtctgagggatcagtctccagcccctatctcttctatactagctggatgtctgagggatcagtctccagcc
cctatctcttctatactagctggatgtctgagggatcagtctccagcccctatctctcctctgatagctggatgtctgagggatcagtctccagcccctatctcttctatactagctggatgtctgagggatcagtctccagtcccaatttctcctctagtagctggatgtct
gagggatcagtctccagcccctatctcttctatactagctggatgtctgagggatcagtctccagcccctatctcttctatactagctggatgtctgagggatcagtctccagcccatatctcttctatactagctggatgtctgagggatcagtccccagcccctatttct
tctatacaagctggatgtctgagggatcagtctcagcccctggttcttctccactgtgtggctgtttaatggctcagtttctaacacttttctatcctttgcttaatggtcttctaagttgatttttaaagttgttagtttagtcatacactaaatcagtttggaaggttgttcagga
gatgagagagagggaatggagcgggaactgaaaatctatctatctatctatctatctatctatctatgtatcatccttctatccactaataagtgatgtaatatttttgggtggttccaatcctctagattcttctggatcgctgatgggtggaaggaagcaagttgtttgtat
ttgtgaccagaagaagcctcataatctaccaagtaaggataaagttaaccccagagcatacagatgcagtaaattttcacttgactttcagtagcttttgtcaggagcgatgtaaagtaatttgtttttgacttgtcaggtcgaggacactgaggccactctgctttctgc
ttgatgagaactatggagtccagctgcagtccacaggcagtaatcttgtggactgaagccgtaactaaagctgcatttgcacgagccaataatcgtccagattatcaggtcgttcctgacaatctgaccgtgtaaatgtcccgccggtcacccgataaacaatgctt
gttcctctggtgatcctgtcgttcatgcaggcaccaccggtcatactgtagtttctgggaagcagatcgtgtggtctaaacagtgatgttctgctacagcaatccctcatccctttactgtgaaggagatcgctgcatgtaaatgtgtctcctccactgaacgagcagc
ccactgccaacccgtgtaaattcacctcaagtagttaaaatctatccgcagcgcttcatgagtcaaagtgtagtcagggccggaagattgctgatctgtatataaagcacacacaaggtactgaaccgatactctgactccctgtaatccagaaactcttgctaataa
aacttatggcaccacactttgtacatggagtccatcctgccaatcagtgttttatatggaacttgcaaaaagaacggaatatggcggcactgctacaaatcaggtaaatgaaaccggcccggtctactggctcagcgccaacaatgcagaaatcagatcttcgtgtg
tggtgctcagcatagagaggggaagcaggatcgagtgcaacaatgtagaagaagtatcaaaagaacatgcggcactcaccgaagtagcaaaagtgtttctctttattctcccttcattcaggacagagtagatggggtgcgggggcggccaagctcttgcagg
tgatcagcagctacggccgtttcgcgcgttctatcgcgcttcttctggccgctctaatcgtcatcgcgcaatactgtgctttataaaggcgacgccccacagccggcatcatcagcaacagctgtgcgccgcaaataagagactttaaaaacatgcatgcgaagaa
gaggaactacatacaaatcggcaccacaggttggatcaatctcaatagtcatacgaacattttcatacaatgtatgtcactacatagtgacactactattttatcctaaccctaatgttcgtatgactattgagattgatccaacctgtggtgccgatttgtatgtagtgtttt
atatggtcctaagtgtggctgatatctgtggccagctgcagcctcaggttccaggcccaaggaggtcaccagactggcctcagatggtacgtggcagtagaggctcctttatcatgttgaccgcttcctccataatgtgacttttatgatccacaggaaaaagacgt
catatgaaggctcgtcgtagacacggtcatcataggcatgtgtgtggcaggatctgcttgacaaaaggtgcaccaggaggactggcggtgcccatacctttgacttgaaggaactctaaatgatgatgcttcacaaagaaaactcacaaaaccgtctcgttatcttt
tctgtagaatccgggaacatggacattacagaacatttttattaagggagcactccagtctacagtatatgcacatataggtataatcgatacatacaccttgttaatatatcacattaatcatcttgtaatgatcttaagttacagagtattattctccagagctgcattcac
aattctgctgctgattataagaaacttgtttttcagtcccctccccacaatatcttagctgagctcctataatatagcaggtcagtttgtctttggtgtagaaggctgtacagagcctggtggaaagacaacccttcccagaatgcaaatcaccagaacatattctatgca
aatgttgaaccagcacagaatggtgagggatttcagctctgaagcttgccgaactgttaattcagcccctagatataatattgggattcaaaagccatacatttaataacctactactgagccgtgaattccaaataatgaggctaaatccatgtttcagcctctactgtt
taacctgcaaaaatgattctaaagcataaaattagaaaaatccttcaaaattaaatttcacaacttaaataaaaaaggagctgaggaaagacaggagagagtttcccaaagtaaagtgaactcatctgtaagaaggcaacaagcttagatcgatactaacgtcaatc
tgttctcccttagtctctcggcagcacaataatgccttactcttactgggcagatggagttttgaataaacaaattagacattaaatagagccctttgaccctctaaaattgcatcccgcccccacctgctgtacagatgcagtcgagctaaacttgacggttaagtaaa
aaatgcctcaaaaaggttaactaatattttcaatgtatttcaatggcttttgtcacaagatggcgcgagataacactgtgacatgtgttatcagagttatcagagtcatttgtagctcggctagataagtatattataaacataataaagaacacaattattacagcagcatg
aatgtgctgcaggactaagggaaaacagacactctccacttaccaatgtcctaacagcggacattcttgcccctgtgtactagtaggacagatggaattaaacattaattagctagaggcccctgatccggaagaggacatgcagccccctccgtgtgttcattaa
aaagatgtaacttaccccggataggaaacgtgctgcagttgggactaaggaaaaacagattaaagtaaaaaaaatatatgcatcccctatgccctaccagcagcacaataatggagagttatcaagaagtatccccatggggaggaccatttcacctgcctgaac
tcaaaatcagaggataggttatcagtatctgattggttggggtccgatcagctgtttgagaaggcagaggcactcctgtggccttctctctgcttttcctaggccagcaacggtcatcttggctagtagcagctcggccccccttcactgatctgcgataccaagcaca
gccgctatacaatttgcggggccgggcttggtgagctgcaagaagcctgcagtgctcacattccactggttcttctaagtcatagagacccacattaatgtgcggctgataggcacagggaaaggccagatccttaggttctcgaatgatgcattctaccatgtata
ttatgtgtg

**C:** Sequence of cDNA (coding region in red and highlighted in yellow)

agagcgtctcttgtaagccaaggtgaagatgaaggcggccgcgatttttatgttgtccgcagctctgatccttgttttctgcttcaccactcaagggcgtaaaagttcaggaagaacttccagcatagtcaagaggactgaccatggagattcttctggatcgctgat
gggtggaaggaagcaagttgtttgtatttgtgaccagaagaagcctcataatctaccaagaaaaagacgtcatatgaaggctcgtcgtagacacggtcatcataggcatgtgtgtggcaggatctgcttgacaaaaggtgcaccaggaggactggcggtgccc
atacctttgacttgaaggaactctaaatgatgatgcttcacaaagaaaactcacaaaaccgtctcgttatcttttctgtagaatccgggaacatggacattacagaacatttttattaagggagcactccagtctacagtatatgcacatataggtataatcgatacatac
accttgttaatatatcacattaatcatcttgtaatgatcttaagttacagagtattattctccagagctgcattcacaattctgctgctgattataagaaacttgtttttcagtcccctccccacaatatcttagctgagctcctataatatagcaggtcagtttgtctttggtgtag
aaggctgtacagagcctggtggaaagacaacccttcccagaatgcaaatcaccagaacatattctatgcaaatgttgaaccagcacagaatggtgagggatttcagctctgaagcttgccgaactgttaattcagcccctagatataatattgggattcaaaagcc
atacatttaataacctactactgagccgtgaattccaaataatgaggctaaatccatgtttcagcctctactgtttaacctgcaaaaatgattctaaagcataaaattagaaaaatccttcaaaattaaatttcacaacttaaataaaaaaggagctgaggaaagacagga
gagagtttcccaaagtaaagtgaactcatctgtaagaaggcaacaagcttagatcgatactaacgtcaatctgttctcccttagtctctcggcagcacaataatgccttactcttactgggcagatggagttttgaataaacaaattagacattaaatagagccctttga
ccctctaaaattgcatcccgcccccacctgctgtacagatgcagtcgagctaaacttgacggttaagtaaaaaatgcctcaaaaaggttaactaatattttcaatgtatttcaatggcttttgtcacaagatggcgcgagataacactgtgacatgtgttatcagagttat
cagagtcatttgtagctcggctagataagtatattataaacataataaagaacacaattattacagcagcatgaatgtgctgcaggactaagggaaaacagacactctccacttaccaatgtcctaacagcggacattcttgcccctgtgtactagtaggacagatgg
aattaaacattaattagctagaggcccctgatccggaagaggacatgcagccccctccgtgtgttcattaaaaagatgtaacttaccccggataggaaacgtgctgcagttgggactaaggaaaaacagattaaagtaaaaaaaatatatgcatcccctatgccct
accagcagcacaataatggagagttatcaagaagtatccccatggggaggaccatttcacctgcctgaactcaaaatcagaggataggttatcagtatctgattggttggggtccgatcagctgtttgagaaggcagaggcactcctgtggccttctctctgctttc
ctaggccagcaacggtcatcttggctagtagcagctcggccccccttcactgatctgcgataccaagcacagccgctatacaatttgcggggccgggcttggtgagctgcaagaagcctgcagtgctcacattccac

**D:** Amino acid sequence (signal peptide shaded)

MKAAAIFMLSAALILVFCFTTQGRKSSGRTSSIVKRTDHGDSSGSLMGGRKQVVCICDQKKPHNLPRKRRHMKARRRHGHHRHVCGRICLTKGAPGGLA
VPIPLT

**Fig. S12.** Gene position and architecture (**A**), genomic DNA sequence (**B**), cDNA sequence (**C**), and encoded protein sequence (**D**) of the unannotated *cxcl17* gene in the NCBI reference genome of *Bufo bufo* (common toad). The information was downloaded from the NCBI gene database (https://www.ncbi.nlm.nih.gov/gene).

### A: Gene position and architecture

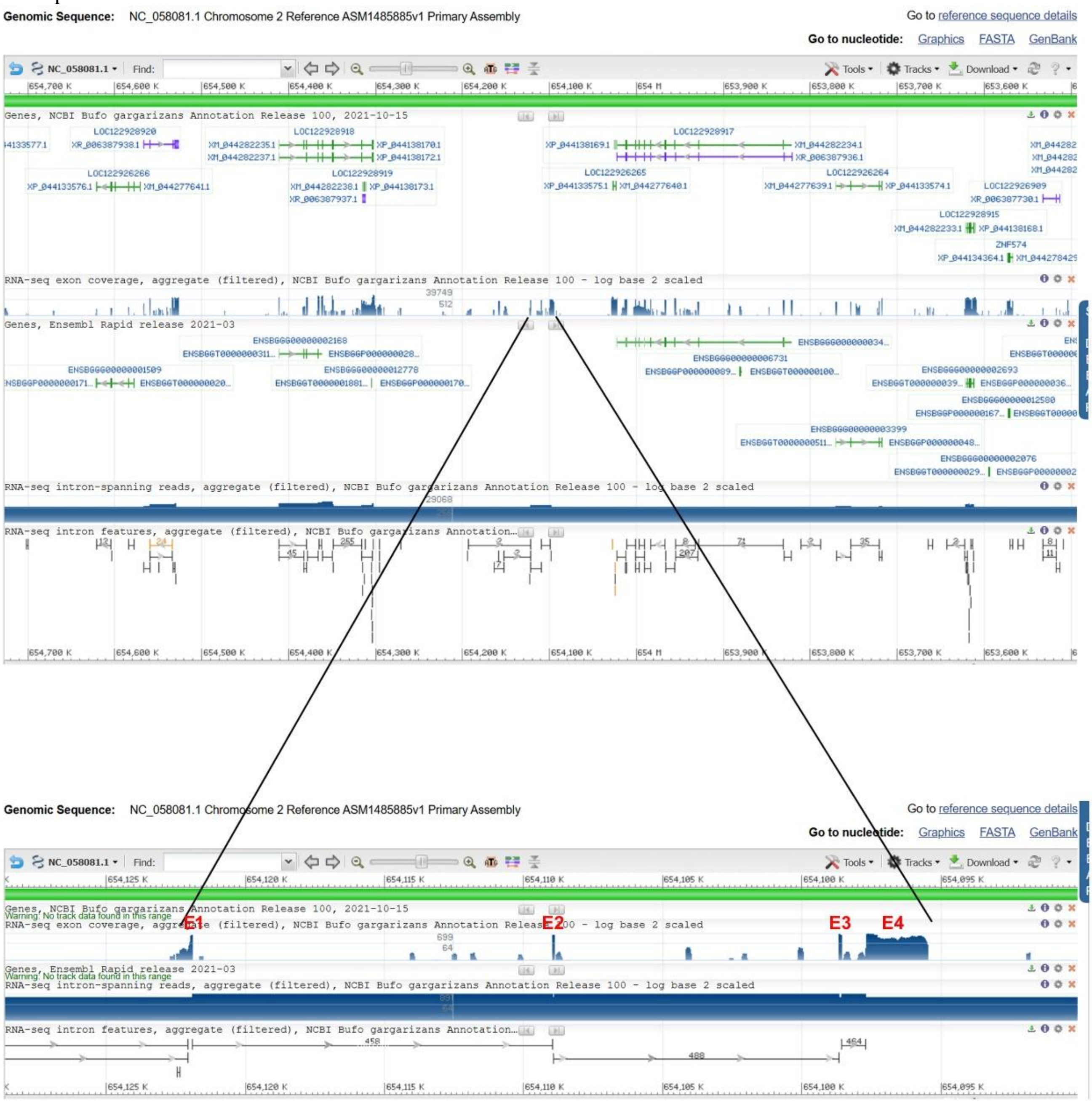


### B: Sequence of genomic DNA (exons in red and highlighted in yellow, coding region underlined)

cacttgatatttcattaatccttatagccctaccccttatatatctgcagcctatgtgagtccacggaggttgtgaaaggcagacaacaaagaatgagagcgtctcttgtaagccaaggtgaagatgaaggcggccgcgatttttatgttgtccgcagctctgatccttc
ttttctgcttcaccactcaagggcgtaaaagttcaggtaacttgtaccaggagaatccaaacagcatgggctctgggagatacaaagtactgcatgggatcacggagcttacaaatatgaatctatgggctctataggacactgcattattataggatctataggaca
ctatatgactccatgggatctataggatactatatgactctatgggatctttaggacactataaaacttgatgggatctataggacactatatgactctatagaatctataggacactatatgactttatgggatctatagggcactatatgactatgagatctataggacac
tatatgacttgatgggattcataggacattctattattctatgggatttataggacactttatgtctctatgggatctataggacactatatgactctatgggatctataggacattgtatgattctgtaggatttctaggatactatatgactctataggatctcttggacattata
tgattctatgatatctataggacattatatgactctatgtatggcatctataggacattatatgattccataggacttataggacattatatcactctatggcatctataggacattaaatgatactataggatttctaagacactattggggtcatttatcaaactggtgttgcc
catagcaaccaatcggattcaacctttcctttttcacaactcctttggggaatgatcccattcaatgggatccatatgacactatggtatctatatcacatatgattctatgtgatctataggacaatatatgactctatgggatttatagggcattatatgattcaataggact
tataggacattctataactctatgaaatctgtaggacaacatatgatactataggatttctaagacattatatgattctatgggatctatcggacactatatgactctatgaaacccataggacattatataatttcataggacataggaaatcatatccctctatgggatgta
taggacactatatgactctatgactagagttgagcggacacctggatgttcgggttcgagaagttcggccgaacttcccgaaaatgttcgggttcgggatccgaacccgatccgaacttcatcccgaactccattgaagtcaatgggacccgaacttttcggcact
aaaaaggctgtaaaacagcccaggaaagggctagagggctgcaaaaggcagcaacatgtaggtaaatcccctgcaaacaaatgtggatagggaaatgaataaaaaaaaaaattaaataaataaaaattaaccaaaatcaattggagagaggtcccatagca
gagaatcaggcttcacgtcacccaccacaggaacagttcattttcataaatttaggcccagcacacaggcagaggagagaggttctgtaacacagaatctggcttcatgtcaccagagaatcagtcttcatgtcatagcagagaatcaggcttcacgtcacccac
cactggaacagtccattgtcagatatttaggcccagcacccaggcagaggagggaggtcccgtaacagagaatctggcttcatgtcagcagagaatcattctgcatgtcatagcagagaatcaggcttcacgtcacccaacattggaacagtccattggcatata
tttaggccccggcacccagacagaggagaggttcattcaactttgggttgcctcgcaatataatggtaaaatgaaaataaaaataggattgaatgaggaagtgccctggagtccaataatatatggttaaggggaggtagttaatgtctaatctggacaagggacg
gacagatcctgtgggatccatgcctggttcatttttatgaacgtcagcttgtccacattggctgtagacaggcggctgcgtttgtctgtaatgacgccccctgccgtgctgaatacacgttcagacaaaacgctggctgccgggcaggccagcacctccaaggcat
aaaaggctagctctggccacgtggacaatttagagacccagaagttgaatggggccgaaccatcagtcggtacgtggaggggtgtgcacaagtactgttccaccatgttagtgaaatgttgcctcctgctaacacgttgcgtatcaggtggtaatgcagttagct
gtggcgtgttgacaaaacttttccacatctctgccatgctaaccctgccctcagaggagctggccgtgacacagctgccttggcgacctcttgctcctcctctgccttggccttgggcttccacttgttcccctgtcacatttgggaatgctctcagtagcgcgtatacc
aacgtgcgcttgtactcgcgcatcttcctatcacgctccagtgcaggaagtaaggtgggcacattgtctttgtaccgtggatccagcagggtggcaacccagtagtccgcacaggttaaaatgtgggcaactctgctgtcgttgcgcaggcactgcagcatgtagt
cgctcatgtgtgccaggctgcccaggggtaaggacaagctgtcctctgtgggaggcgtatcgtcatcgtcctgcctttcccccccagccacgcaccaatgatgggcccgagctgcgttgggtgccaccccgctgtgaccatgcttcatcctcatcctcctccacct
cctcctcatcctcgtcctccccgtcctccagtagtgggccgtggctggccacatttgtacctggcctctgctgttgaaataaacctccctctgagtcacttcgaagagactggcctgaaagtgctaaaaatgacccctcttcctcctcctcctcctcctgggccacctc
ctcttccatcatcgccctaagtgttttctcaaggagacatagaagtggtattgtaacgctgataacggcgtcatcgccactggccatgttggtggagtactcgaaacagcgcaacagggcacacaggtctcgcatggaggcccagtcattggtggtgaagtggtg
ctgttccgcagtgcgactgacccgtgcgtgctgcagctgaaactccactatggcctgctgctgctcgcacagtctgtccagcatgtgcaaggtggagttccacctggtgggcacgtcgcatatgaggcggtgagcgggaaggccgaagttacgctgtagcgca
gacaggcgagcagcggcaggatgtgagcgccggaagcgcgcacagacggcccgcactttatgcagcagctctgacatgtcggggtagttgtgaatgaacttctgcaccaccaaattcagcacatgcgccaagcaagggatgtgcgtcaaaccggctagtc
ccagagctgcaacgagatttcgcccattatcacacaccaccaggccgggcttgaggctcaccggcagcaaccactcgtcggtctgttgttctataccctgccactactcctgtgcggtgtggggcctgttccccaaacatatgagtttcagaatggcctgctgacg

tttaccccgggctgtgctgaagttggtggtgaaggtgtgtggctgactggatgagcaggtggaagaagaggaggaggaagccgagaaggaggaggtggcaacaggaggcaaagaatgttgccctgcgatccttggcggcggaaggacgtgcgctaaac
agctctccgcctggggcccagctgccactacatttacccagtgtgcagttagggagatatagcgtccctggccatgcttactggtccacgtatctgtggttaggtggaccttgccacagatggcgttgcgcagtgcacacttgattttatcggatacttggttgtgca
gggaaggcacagctctcttggagaagtagtggcggctgggaacaacatactgtgggacagcaagcgacatgagctggttgaagctgtctgtgtccaccagcctgaatgacagcatttcataggccagtagtttagaaatgctggcattcagggccagggatcg
agggtggctaggtgggaatttacgctttctctcaaatgtttgtgagatggagagctgaacgctgccgtgtgacatgcttaagacgcttggtgacggaggtggtggtgttggtggtatatcccctgtttgctgggcggcaggtgccaacgttcctccagaggcggag
gaagaggccgaggcggcagcagcagaagagaccgaggcggcggcagcagcagaagagaccgaggcggcagcagcagaagaggccgaggcggcagcagcagaagaggccgaggcggcagcagcagaagaggccgaggcggcagcagcag
aagaggtagcaggggggagcctgagtgacttccttggttttaaggtgcttactccactgcagttcatgctttgcatgcaggtgcctggtcatgcaggttgtgctcaggttccgaacgttaatgcctcgcttcaggctctgatggcacagcgtgcaaaccactcgggtct
tgtcgtcagcacattgtttgaagaagtgccatgccagggaactccttgaagctgcctttggggtgctcggtcccagatggcggcggtcagtagcaggcggagtctcttggcggcgggtgttctgcttttgcccactgctccctcttttgctacgctgttggctcggtc
tcaccactgcctcttcctccgaactgtgaaagtcagtggcacgaccttcattccatgtggggtctaggacctgatcgtcccctgcatcgtcttccacccagtcttgatccctgacctcctgttcagtctgcacactgcagaaagacgcagcagttggcacctgtgtttc
gtcatcatcagagacatgctgaggtggtattcccatgtcctcatcatcaggaaacataagtggttgtgcgtcagtgcattctatgtcttccaccgctggggaagggctaggtggatgcccttgggaaaccctgccagcggagtcttcaaacagctctgggcagcct
aaaaacagtgagcaattgagtatcagcagctcaatgacccacagctgcagatcgatcaatggatggagtcttttggaggagttaatcagcctaatctcgccctaacgtcgcagctgcaacctctccctacactgatcagagcagagtgacgggcggcgctacgt
gactccagcttaaatagaggctgggtcacatggtgctctggccaatcacagccatgccaatagtaggcatggctgtgagggcctcttggggcaagtagtatgacgcttgttgattggctgctttgcagcctttcaaaaagcaccaagaaagcgacgaacaccga
acccgaacttttacgaaaatgtccgggttcgggtccgtgtcacggacaccccaaaattcggtacgaacccgaactatacagtccgggttcgctcatccctatctatgacatatacagaacaatatatgattatataggagttataggacattatatgagtgtaacgag
cccctgctcgctctattcggcactccctgatgtgaccacaatatctgctggttccgacgttgatgatccggcctcaaacaaccgctcgtgtggttctaattctcccagaactaacacccgtcaggccgtatgtgagttcaaactgacaattctttattgaatgccccaca
tatatttatattgacagttggaagacctctttcaattggtaaattgtaaaaccccctctgattggctatgcaaatgaggtaagcagggattggttcaagagcttggctgggaggcagataggtgtggccattgtcacagagagattcaaaccatgacattacccccccc
tcccgggaaacagtccaacagccacagtgggaccaactcgaggatgttggaaaagcagtcttaagattccatgtctgtttgccgggataatccatccgcattccccttttgaggccctggccgatattgtatggtgaaattgtagggttgcaaagccaggctccac
cttagtaattggccattctctcccgccacccggtttagccatatcaagggattgtggtccgtcagccagcgggtgctccttgccgtcgtcgccaactggacttagtacggctcccagtccgaacaaggaggcgtctctatgtacgacaaagcgtttgttaggatcgg
gtgccgccagtatgggagcttgtgttagggtggttttcaactgttgaaatgctgatttgcactccggggaccacaggacctagggagattctttttggtcaggtccgtcagtggtttggctatgtcactatagtggggtacaaactttctgtagtacccggctgtcccta
aaaatgcaagtacatgggccttggttctgggaaccagtttaggatggcctcgatttttgcaggctccggcctctgcttcccacatcccactctatgtcccaaatattgcatctcagacatcccgatatcacacttgtctggcttcagggtcaggcctgcaactctgattc
tgtctagcactatgcccagctgtctgagatgctactcccatgtctggctatagattgcgatgtcctcaagatatgcacatgcaaaatcctgcagaccaccgaggagttaggaaagtagccggagcatttttcatcccgaatggcattacctggaactggtaaaggcc
aaacggggtgatgaaggccgactttggaatggcgtccgcctccaggggactctaccagtagcccttacataggtcaatagtcgtcagataatgtccctgagctattttgtccaacaattcgtccacttggggcatggggtaggcttctctcacagtacagtcatttag
tcacctatagtccacacaaaatcgggtcatgccatcccattttgggactaatactacaggggaagcccaggggcttgctgatggttcgatcaccccctaactcaagcatctccctaatctctgctctcatcccttcttgcacagctccgggaatcctataggctgattgt
ctcaggggcttctgcctgagggtctccatcttgtgcactgctaagtgagtgtatccagggagtgttgagaacatctctttctggtcctggagtaattccctgacctgttcttgttcctggggtcctaacccatctcctacctgaacatcttccacccctccagaacggtta
ctgattgcaggaaactccagcattggcaggtattcagagtccccggtggcaggagcgcatatggcggctactgcctcaggcctctcgtgataggccttcatcatattgatgtggaacgtcctcttaattcttgtgtttgcacagctggcgaccacgtaggtggtgtc
gcacagctgttttaccacctgataggggccctgccaggaagcctgtaattaattttgcttccagggttttagtactaggaccttctttccgacacaaaaagtgcaattccgagcagatcgatcataccaaactttctgctgactctgggctagctcggtaagttcccaca
gtcgatccctgagttccaagatgtaactaaggccttccggatcctctccccctttcccactgggtcctcaccaagtctaagggtcccctaacccttcgcccatacaacagctcaaatggtaaaaaccctgtggactcttgaggtacctctttgtaggcgaacagtaggt
ggggaagatatttctcccaatccttcctacttttaacgaaggtccgaagtagctgtttaagggtcccattaaatctctcgcacagcccgttcgtttgggggtggtagggtgcgctctggagggttttaattccacagaggcgccatagctgctgggtcagctcggcgg
agaactgatttccttgatcagacaatatctcctggggaaatcctaccctagtaaacacccacagtaaggcatcagctaccgtgtcagcctggatattggataggggaatagcttccgggtagcgggtggcgtaatccagcactgtaagaatatacttcttcccagtg
gctctggggcgggccagtgggccaataatgtctactgctatccggctaaagggttacctaattttcggcatggggtgcagagaggcttttggatgatcccccgcctttcctaccctctggcaagtctcacaggttcggcagtattctcatacctctgcatgtaaccta
ggccagaagaaagcacgagacaactgactccgggtcttagctatccctaaatgtctgatcaaggggatatcgtgagccagcctcaatagctcctgtctgtacttctggggtacgactagctgtttgtgggggccctcgtgcttcccgttgcccacccccttcagtgta
acgataaagaagccctccctcccactctattctgtcttcccctaaccccccctggatccttgcccaccctgtcccgatagcacgctaaagtagggtctctccgggtttcttgcctaacatcggaaggagaatcccaggacatgtggttgtcaagtcggggtaacgtgg
gaggattttgtcttacctgggtctccaaaggagtggagttgccttgtaggtggctttgttgtcgggtggtcacggacatggcagtctctggtgtaaatgaagaagtcaaatgtcccaagtcattccccaaaagtacttccgcgggtaactcacggagcatccctactc
gcacttgtcgcttccctgagccccaatgcaggtgaactctggcggtgtataaacgcagcaccttgccccttgctaccctcactgccacagtctggccagttcaggccgactgaggaaccatgcaaggctaaataagggcaagcgtagccccggaatcacgaag
tccctgggccggatgcccattaacccataccgcctgacgatggtgctgccgattatcttcagctgcgccctggacgggatcagccctcgaacagctcgaaaaactcttcctgaacatctagagggttggcctcattctggaggcaataagtcgccgacctggata
actgggcagtctcttggtaccttggggatcggttagggcaataacgctgcagatgacccgtctgattacatgtatagcacttaggtccgtagaaggttttgatcgggctaagggtctagaggcagcccactgagatactggagttgagctcaaggtcagtacactg
ggccatggtagtggatagcgctggcctccaatggatctcagagaatccagatattcatcggccaaagtcgctgcttgctgtaccgtctgtggccgcttatctcgcacccaatcccatactatcggaggggtgtggtcaaagaactgctctaagagaactagctgg
gtgatgtcctctatggtgtgagcatggcttccctggaaccagcccttgaggttccgctgcagatgatgtgcccactccacgtaggttagatagccttgtttccgggattctcaaaacttaatccagctggcttcaggggtaatggcaaatcgggctagcagggctgc
ctttacccggtcataatccatggcttcctcatcctccatggccagatacgcatccgcagctttgccagttaggttactcgccagtaaggcgacccgatcttgggacggcactttatgtatctgccacagtctctcaaagtcctggaggaactcttcaattttctcctcttct
tccttaaataatttgaatgcgcgaaagggcaacttacgcacatatgctatatctctaggtgctgtctctgatcgctgtatttccaccattctccactcatgttgtcgctctagttctctctcttgctgctcatcctgcagctggatgtctctgatgtcattctgtatggcggtctc
ggatgggttggcaccatataatgccaaccggtctctaactcgtcttcctcagaccgaggtccagcatcaccatccctctgatccatctcagctaactcactgatcaaggtggccttgttctttgtcccagcaatccctcctcggctctcaagtgaatctttcagcgtgtc
tctcctgccggctgcgtagctggtcattcctagttgtgagttggagtgtcccagtaactggagagccaatcctaccactgccaccaatgtaacgagcccctgctcgctctattctgctctcacggcactcccttatgtgaccacaatatctgctggttccgccgttgat
gatctgtatgtgagttcaaactgaccaaattctttattgaatgcgtcacatatatttgtattgacagttggaacacctctttcaattggcaaattgtaaaaccccctctaattggctatgcaaattagttaagcagggattagttcaagagcttggctgggaggcagatggg
tgtggccattgtcacagagattcaaaccatgacaatgagtctatggaatctatagaacactgtatgatactataggatttctaggacactatatgacactatgggatctatgggacattatatgattccgtaggacttatacaacgtgatatgactctatgggatatactgt
ataggacattatttgactctactggatatataggacactatatgactctatgggatctataggacaatatatgactctatatgatctatggcacattttatctccttccatcaattctcagcagatttagttctaagagaaattccatcgcatttccttccatcaaccatcactag
cctcttattataagaccttccatcccatctccctccatcaaccgtctccagcctcttattatcctacctattattactaccaatctgttacaaatgttggtagttgtagtaccttgttcgcctatttctcacaaactgctgatcctcagggattatagttgcagttcgggtgtggat
aggaatggttctagacgaatgcagcttccaagtagttcttcccccagccagtagatagttacccaaacaggagcccagacttcatgaagggtacaacaggaacctttaatggtgctacatgtcagctttttatgcaggtcttccagcatggggaccttgtggaagac
tgagacagtatcttacataaaccaattacatgcagttataggcagaaaacacacacatgtgatacaattaactaacataatggtataacgtaatcactcacaggcagaaaccacacacttttcccttgtagaataatgagctgggggtggggtggtcgttggtcggta
gattcctctaccgaaaacataacatggcctcacaccatctccctccatcaaccgtctccatcctcttattataagagcatccatctcatttccctccatcaacccttttccaggatcttattataagatagaacttccattccgtctccctccatcaaccatctccagcctctga
ttataagactttccatcccatctccctccatcaaccctctcaagcttcttgttctaagagactttgcatcccatctccctccatcaaccctatccagcctcttattataagagaccttccatcccatctacctccataaaccctatccagcctcttattataagagactttttatcc
cttctccctccatcagcctcttataaaactttccatatcatctccctccatcaactctatccagcttcctattataagaccttccatcccatctccctccatcaacccactccagcctcttattataaaaagaccttccataccatctccttccatcaaccctctccagcctctg
attataagacctttcatcccatctccctccatcagcccttttataagactttccatatcatctccctccatcaaccctatccagcctcctattataagaccttccatcccatctccctccatcaatcctcgccagcctcttcttctaagagaccttaaatcccccccccccctccat
caaccctctccagcctcctaatataagaactactatcccatctctctccattaaccctctccagtctcttattataaaagaccttccatcccttctccctccatcaacccccctccagcctctgattataagaccttccatcccatctccctccatcaaccctctccagcctctt
attattaagagaccttctatcccatctcgttccatcaaccttctctagcctcttattataaagaggccttctatcccatctacctccatcaaccctctccagctcttattggagagattttccatctctgaggtgactcagatctcctttatccttttttaacatgtggagcttaaa
gcggttttcccatcttggacatttggatcatatgtctaagatatgaccccaatatcagataggtgggggtgccacatctgggaaccacacatatacttagaatgaagcctgcaaagttcaggagggtgcactgcacacgccgtcctccattcattcctatggcagag
ctgacaatagctggctattttcggaagtcctattaaaatgaataggaagctcagccaccactccattcctttcaataaggtttactgaaatagccgagccaacgctcagccatttttggtgctcccttaagaataaatggtggcggcctgcgcatgcaccctctggcac
tttgcgggctccattctaagtatagatgcgagtcccagaagtgggacccatagctgtcagacattgagtgcatattctagtgatatgaccccaatgtccaagatgggaacaacccatttaagccttaattccgtattggagatatcctagttcaccagtgagaaaatgt
ctgggcccatttccctaatctacagtccatggtgatctagtggatgtaaaccagacccttttgtttaagacatcctggtctgcatcataagatgagaagaacaccatatgggatccaccatctctctacagacccgttatagcagggcatgaatagagagctgtacatt
tttttccaattaaaaactattctatcttttaggaagaacttccagcatagtcaagaggactgaccatggagggtaatacaatgagcattaaagactatagacacatttggaggcatttttttttataattaattatttacatctattttgggctaaaaactttttttgtatttcttgattaa
aaaaatgttgctttggtttctgcagcgcatttatttcataacacctagcagtccacagaaaaccactctgtgatgaatctgccagactggcactctgtcaggtccacagcccgtcactctatactactggctggcagttggatggctttgtctctagttcctctctcttctct
actgcagacttcttgacaactcagtctcctgaaactctgtttgctatggagtctcctgaaactctgtttgctatggagtctcctgaaactctgtttgctatggagtctcctgaaactctgtttgctatggggacctatttgacatattaggctccagcttttctctatcttctagct
ggctgtgtgatggcttggtctccagtgcctttcttttctccactgctgggcttcctgacagcttggtcccccaacccctttcttcactattggttgactgatgtcttggtatctaggcctcctctttcctctgctaactggctgtctgacagttcagtatctagccccctgtcttct
ctgttgagtagctgtttattggcttggtctccagcccctagtctggctgtttataagttttgtctccagccatgttctctgctataccatgtgtctgatggtttggtttctagccattctcttctctattggattactattgattggcttagtctttagcagctctctctagtatttgtcta
tctgacagcttgatctcccctcttacttttctatttgctgtccctctattggcacagtctctatctcctctccctctctcgcctttagtagctgactgtctgatggcttactttccagtccttctctcttctttactggctggctgtctattagctcagtctccagcttctcagttattgt
ctaacgtttgactctcctgcccctatgtcttctatattagctggatatctgacagtttggtctccaggccctatcttctctctaatgactgtctggcagcacagtctccagttcctccctcttctctgctagcggtctaatgacttggtctccagcctctatctcctcaactatct
gtctgcctgtctgttgatttagcctgtcgtccttctcacttctttactggttggttctatgatggcttggtcttcagcatatttcttacctcaactggctggatgtctcatagcttggtgtccagcctctctgtcttctctaatagtaggctagtaggctgtctgacagcttggtctc
tatccccttctcttcttcaatgtctgtgtgatggcttggtgttcatttcctatcagttctatatttgctggatgtgtgatggctcagtctatagcccttctctttttttttactggctggctttttgttcaggctccagtctatcccttctatagtagctggctgccgaatggctcacta
tcctgcacctctctctcatctactagatggatgtctataacttggtcttcaaacactccctcttatataatagcctgctgtctgactgctcagtctccagcccctatctcttgtattacttgctgattgcctgatggcttggtctccagctcttttactagtgtgctgtctaatggct
cggtctgcagcccctatgtcttttatactgtctggatgtctattgactgaataaccagctcttttctctccctctagtaggtgtatgtctgagggatcagtctccagcccgtatctcttctatactagctggatgtctgagggatcagtttccagcccctatctcttctatactag
ctggatgtctgaggaatcagtctccagcccctatctcttctatactagctggatgtctgaggaatcagcctccagcccctatctcttctatactagctgaatgtctgagggatcagtctccagcccctatctcttctatactagctggatgtctgagggatcagtctccag
cccctatctcttctatactagctggatgtctgagggatcagtctccagcccctatctcttctatactagctggatgtctgagggatcagtctccagcccctatctcttctatactatctggatgtctgagggatcagtctccagcccctatctcttctatactagctggatgtc
tgagggatcagtctccagcccctatctcttctatactagctggatgtctgagggatccgtctccagcccctatctcttctatactagctggatgtctgagggatcagtctccagcccctatctcttctatactagctggatgtctgagggatcagtctccagcccctatctc
ttctatactagctggatgtctgaggggtcagtctccagcccctatctcttctatactagctggatgtctgaggggtcagtctccagcccctatctcttctatactagctggatgtctgaggtatcagtctccagcccctatctcttctatactagctggatgtctgagggatc
agtctccagaccctatctcttctatactagctggatgtctgaggaattagtctccagcccctatctcttctatactagctggatgtctgagtgatcagtctccagcccctatctcttctatactagctggatgtctgagggatcagtctccagcccctatctcttctatactag
ctggatgtctgaggaatcagtctccagcccctatctcttctatactagctggatgtctaagggatcagtctccagcccctatctcttctatactagctggatgtctaagggatcagtctccagcccctatctcttctatactagctggatgtctgaggaatcagtctccagc
ccctatctcttctatactagctggatgtctgaggaatcagtctccagcccctatctcttctatactagctggatgtctgagggatcagtctccagctcctatctcttctatactagctggatgtctgagggatcagtctccagcccctatctcttctatactagctggatgtct
gagggatcagtctccagcccctatctcttctatactagctggatgtctgagggatcagtctccagcccctatctctgctatactagctggatgtctgagggctcagtctccagcccctatctcttctatactagctggatgtctgaggggtcagtctccagcccctatctc
ttctatactagctggatgtctgagggatcagtttccagcccctatctcttctatactagctggatgtctgaggtatcagtctccagcccctatctcttctatactagctggatgtctgaggaatcagcctccagcccctatctcttctatactagctgaatgtctgagggatc
agtctccagcccctatctcttctatactagctggatgtctgagggatcagtctccagcccctatctcttctatactagctggatgtctgagggatcagtctccagcccctatctcttctatactagctggatgtctgagggatcagtctccagcccctatctcttctatactat
ctggatgtctgagggatcagtctccagcccctatctcttctatactagctggatgtctgagggatcagtctccagcccctatctcttctatactagctggatgtctgagggatcagtctccagcccctatctcttctatactagctggatgtctgagggatcagtctccag
cccctatctcttctatactagctggatgtctgagggatcagtctccagcccctatctcttctatactagctggatgtctgaggggtcagtctccagcccctatctcttctatactagctggatgtctgaggggtcagtctccagcccctatctcttctatactagctggatgt
ctgaggaatcagtctccagcccctatctcttctatactagctggatgtctgagggatcagtctccagaccctatctcttctatactagctggatgtctgaggaattagtctccagcccctatctcttctatactagctggatgtctgagtgatcagtctccagcccctatctc
ttctatactagctggatgtctgagggatcagtctccagcccctatctcttctatactagctggatgtctgaggaatcagtctccagcccctatctcttctatactagctggatgtctaagggatcagtctccagcccctatctcttctatactagctggatgtctaagggatc
agtctccagcccctatctcttctatactagctggatgtctgaggaatcagtctccagcccctatctcttctatactagctggatgtctgaggaatcagtctccagcccctatctcttctatactagctggatgtctgagggatcagtctccagctcctatctcttctatactag
ctggatgtctgagggatcagtctccagcccctatctcttctatactagctggatgtctgagggatcagtctccagcccctatctcttctatactagctggatgtctgagggatcagtctccagcccctatctctgctatactagctggatgtctgagggctcagtctccag
cccctatctcttctatactagctggatgtctgaggggtcagtctccagcccctatctcttctatactagctggatatccgagggatcagtctccagcccctatctcttctatactagctggatgtctgagggatcagtctccagcccctatctcttctatactagctggatgt

ctgagggatcagtctccagcccctatctcttctatactagctggatgtctgagggatcagtctcaagcctctatctcttctatactagctggatgtctgagggatcagtctccagcccctatctcttctttactagctggatgtctgagggatcagtctccagcccctatct
cttctatactagctggatgtccgagggatcagtctccagcccctatctcttctttactagctggatgtccgagggatcagtctccagcccctatctcttctatactagctggatgtctgagggatcaatctccagcccctatctcttctatactagctggatgtctgaggga
tcagtctcaagcctctatctcttctatactagctggatgtctgaggaatcagtctccagcccctatttcttctttactagctggatgtctgaggaatcagtctccagcccctatctcttctatactagctggatgtctgagggatcagtctccagcccctatctcttctatacta
gctggatgtctgagggatcagtctccagcccctatctcttctatactagctggatgtctgagggatcagtctacagcccctatctcttctatactagctggatgtctgaggaatcagtctccagcccctatctcttctatactagctggatgtctgagggatcaatctcca
gcccctatctcttctatactatctggatgtctgagggatcagtctccagcccctatctcttctatactagctggatgtctgagggatcagtctccagcccctatctcttctatactagctggatgtctgagggatcagtctacagcccctatctcttctatactagctggatgt
ctgagggatcagtctccagcccctatctcttctttactagctggatgtccgagggatcagtctccagcccctatctcttctatactagctggatgtccgagggatcagtctccagcccctatctcttctttactagctggatgtccgagggatcagtctccagcccctatc
tcttctatactagctggatgactgaggaatcagtctccagcccctatctcttctatactagctggatgtttgagggatcaatctccagcccctatctcttctatactagctggatgtttgagggatcaatctccagcccctatctcttctatactagctggatgtctgagggat
cagtctccagcccctatctctgctatactagctggatgtctgagggatcagtctccagcccctatctcttctatactagctggatgtctgaggaatcagtctccagcccctatctcttctatactagctggatgactgaggaatcagtctccaacccctatctcttctatact
aactggatgtctgagggatcagtctccagcccctatctcttctatactagctggatgtctgaggggtcaatctccagcccctatctcttctatactagctggatgtctgagggatcagtctccagcccctatctcttctatactagctggatgtctgagggatcagtctcc
agcccctatctcttctatactagctggatgtctgaggggtcaatctccagcccctatctcttctatactagctggatgtctgagggatcagtctacagcccctatctcttctatactagctggatgtctgaggaatcagtctccagcccctatctcttctatactagctggat
gtctgagggatcagtctccagcccctatctcttctatactagctggatgtctgagggatcagtctccagcccctatctcttctatactagctggatgtctgaggggtcagtctccagccctatctcttctatactatctggatgtctgagggatcagtctccagcccctatc
tcttctatactatctggatgtctgagggatcagtctccagcccctatctcttctatactagctggatgtctaagggatcagtctccagcccctatctcttctatactagctggatgtctaagggatcagtctccagaccctatctcttctatactagctggatgtctgagggat
cagtctccagcccctatctcttctatactatctggatgtctgagggatcagtctccagcccctatttcttctatactagctggatgtctgagggatcagtctccagcccctatctcttctatactagctggatgtctgagggatcagtctccagcccctatctcttctatacta
gctggatgtctgagggatcagtctccagcccctatctcttctatactagctggatgtctgagggatcagtctccagcccctatctcttctatactagctgaatgtctgagggatcagtctccagcccctatctcttctatactagctggatgtctgagggatcagtctcca
gcccctatctcttctatactagctggatgtctgagggatcagtctccagcccctatctcttctatactagctggatgtctgaggaatcagtctccagcccctatctcttctatactagctggatgtctgagggatcagtctccagcccctatctcttctatactagctggatg
tctgaggaatcagtctccagcccctatctcttctatactagctggatgtctaagggatcagtctccagcccctatctcttctatactatctggatgtttgaggaatcagtctccagcccctatctcttctatactagctgaatgtctgagggatcagtctccagcccctatct
cttctatactagctggatgtctgagggatcagtctccagcccctatctcttctatactagctggatgtctgagggatcagtctccagcccctatctcttctatactagctggatgtctgagggatcagtctccagcccctatctcttctataccagctggatgtctgaggga
tcagtctccagcccctatctcttctatactagctggatgtctaagggatcagtctccagcccctatctcttctatactagctggatgtctgagggatcagtctccagcccctatctcttctatactagctggatgtctgagggatcagtctccagcccctatctcttctatact
agctggatgtctgagggatcagtctccagcccctatctcttctatactagctggatgtctgaggggtcagtctccagcccctatctcttctatactagctggatgtctgagggatcagtctccagcccctatctcttctatactagctggatgtctgagggatcagtctcc
agcccctatctcttctatactagctggatgtctgagggatcagcctccagcccctatctcttctatactagctggatgtctgaggaatcagtctccagcccctatctcttctatactagctggatgtctgagggatcagcctccagcccctatctcttctatactagctgga
tgtctgaggggtcagtctccagcccctatctcttctatactagctggatgtctgagggatcagtctccagcccctatttcttctatactagctggatgtctgagggatcagtctccagcccctatctcttctatactagctggatgtctggggaatcagtctccagcccta
tctcttctatactagctggatgtccgagggatcagtctccagcccctatctcttctatactagctggatgtctggggaatcagtctccagcccctatctcttctatactagctggatgtctgagggatcagtctccagcccctatctcttctatactagctggatgtctgagg
gatcagtctccagcccctatctcttatatactagctggatgtctgagggatcagtctccagcccctatctcttctatactagctggatgtctgagggatcagtctccagcccctatctcttctatactagctggatgtctgaggggtcaatctccagcccctatctcttctat
actagctggatgtctgagggatcagtctacagcccctatctcttctatactagctggatgtctgagggatcagtctccagcccctatctcttctatactagctggatgtctgagggatcagtctccagcccctatctattctatactagctggatgtctgagggatcagtct
caagcctctatctcttctatactagctggatgtctgaggaatcagtctccagcccctatctcttctatactagctggatgtctgaggaatcagtctccagcccctatctcttctatactagctggatgtctgagggatcagtctccagcccctatctcttctatactagctgg
atgtttgagggatcaatgtccagcccctatctcttctatactagctggatgtctgagggatcagtctccagcccctatctcttctatactagctggatgtctgagggatcagtctccagcccctatctcttctatactagctggatgtctgaggggtcagtctccagcccct
atctcttctatactagctggatctctgagggatcagtctccagcccctatctcttctatactagctggatgtctgagggatcagtctccagcccctatctcttctatactagctggatgtctgagggatcagtctccagcccctatctcttctatactagctggatgtctgag
ggatcagcctccagcccctatctcttctatactagctggatgtctgaggaatcagtctcctgcccctatctcttctatactagctggatgtctgagggatcagcctccagcccctatctcttctatactagctggatgtctgaggggtcagtctccagcccctatctcttct
atactagctggatgtctgagggatcagtctccagcccctatttcttctatactagctggatgtctgagggatcagtctccagcccctatctcttctatactagctggatgtctgaggggtcaatctccagcccctatctcttctatactagctggatgtctgagggatcagt
ctacagcccctatctcttctatactagctggatgtctgaggggtcagtctccagcccctatctcttctatactagctggatgtctgaggggtcagtctccagcccctatctcttctatactagctggatgacggaggaatcagtctccagcccctatctcttctatactagc
tggatgtctgaggggtcagtctccagcccctatctcttctatactagctggatgtctgagggatcagtctccagcccctgttcttctccactgggtggctgtttaacgttcagcctccaacacctttctatcctttgcttctattatctatctatctatctatctatctatctatcta
tctatctatctaccatccttctatccactaataattgatgtaatattttggggtggttccaatcctctagattcttcgggatcgctgccgggtggaaggaagcacgttgtttgtatttgtgaccagaagaagcctcataatctaccaagtaaggatgaagctaaccccaga
gcatacagatgcagtacattttctcttgactttcagtagcttttgttatgagcgatgtaaagtttgctgtaatttgtttttgacatgtcaggtcgaggccactcaggtgtcaggtcgaggccactgaggccactctgctttctgcttgatgagaactatggagtccagctgc
agtccacaggcagacagtgggttgtatgtaatcttgtggactgcagccgtaactaaagctgcatttgcacgagccaataatcgtccagattatcaggtcgttcccgacaatctgaccgggtaaatgtgctgccgattacccgatgaacaagtgaaatgcttgtccct
cgggtgatcctgtcattcatgcaggcaccaccggtcatactgtcgtttctgggaagcggatcgtgtggtccaaacagtgatctgctgctacagcaatccctcatccctttactgtgaaggagatcgctgcatgtaaatgtgtctcctccactgaacgagcagcccact
gccaacccgtggaaataagcagctaaaatctatcagcagcgctacatgagtcagagtgtagtcagggccgaagattgccgatctgtatctgaagcacacataaggtactgaaccgatactctgacatcctgtaatccagaaactcttgctaataaaacttatggca
ccacactttgtacatggggtccatgctgccaatcagtgttttatatggtcctaagggtggctgatatttgtggccagctgcagcctcaggttccaggcccaaggtggtcaccagactggcctcagatggtacgtggcagtagaggctggttcatcatgttgaccgctt
cctctataatgtgacttttatgatccacaggaaaaagacgtcatatgaaggctcgtcgtagacacggtcatcataggagtgtgtgtgacaggatctgcttgacaaaaggtgcaccaggaggactggcggtgcccataccattgacttaaaggaactctaaatgatg
acgtttcacaaagaaaactcacaaaaccgtctcgttatcttttctgtagaatccgggaacatggacattacagaacatttctattaagggagcactccagtctacagtctatgcacatataggtataatcaatagatttacaaggtgactcgactgtatatatcacattaat
cctcttgtaatgatcttaagttacagcatattattctccagagctgcattcacagttctgctgctgattataggaaacttgtttttcagtcccctccccacaatatcttagctgagctcctataatatagcaggtcagtttgtctttggtgtagaaggctgtacagagcctggtg
gaaagacaacccttcccagaatgcaaatcaccagaacatattctatgcaaatgttgaaccagcacagaatggtgagggatttcagctctgaagcttgccaaaactgttaattcagctcctggatataatattgggattcaaaagccatccatttaataacctactaccg
agccgtgaattccaaataatgaggttaaatctatatttcagcctctactgtttaacctgcaaaaatgattctaaagcatgaaattttaaaaatccttcaaaaattaaatttcataacttaaataaaaaaaggagctgaggaaagacaggagagagtttcccaaagtaaagc
gaactcatctgtaagaaggcaacaagcttagatcgatactaacgtcaatctgttctcccttagtctctcggcagcacaataatggggagtctccgccaccgcttactcttactgggcggatggagttttgaattaaactaattagacattaaatagagccctttcaccct
ctaaaactacatcccgcccccacctgctgtacagatgcagttgagctaaacttgacggttaagtaaaaaatgccacaaaaaggttaactaataatttcaatgtattttgtcacaagatggcgcgagataacactgtgacatgtgttatcagagtcaattgtagctcggc
tagatcagtatattataaacatcataaagaacacaattattacagcagcaagcatgtgctgcaggactaagggaaaacagacactctccacttaccaatgtcctaacagcggacattcttgcccctgtgtactagtaggacagatggaattaaacattacttagctag
aggcccctgatccggaagaggacatgcagccctctctgtgtgttcattaaaaagatgtaacttaccccggataggaaatgtgctgcagttgggactaaggaaaaacagattaaagtaaaaaaaaaaatgcttcccctatgccctaccagcagcacaataatggag
agttatcaagaagtatccctatggggaggaccatttcatctgcctgaactcaaaatcagaggataggttatcagtatctgattggttggggtccgatcagctgtttgagaaggcagaggcactcctgtggccttctctctgcttttactaggccagcaatggtcatgttg
gctaggagcagctcggccctcttcactgatctgcaataccaagcacagccgctatacaatttgcggggctgggcttggtgagctgcaagaagcctgcagtgctcacaggactcaagtggacttagctgaaagttggactttgatgaacgttggcctccttcatatg
ctcatgcagagagcgagggatgttaatcacagattagaacgttcttgagtagaacaccaagccaaacaaaccttctttaatagtccacactggacaggtggagcacaaggaggaggatcttgatgtcctttaagggcctaaaagtaattagtaattacttaattaaa
aattccatagagattccataagtcatagagacccacattaatgtgctgctgataggcacagggaaaggccagatccttaggttctcgaatgatgcattctaccatgtatattgtgtgtgggtttggggtctgtgtgagttcagtgtaatccttgtattgtcacatgcccgt
cttattgtgacacagaacatgtgattgtgtctccagttcaataaacatttttatattttggcactttctaattgtttttgtcaatgattttaaattacattcaatgatgaga

**C:** Sequence of cDNA (coding region in red and highlighted in yellow)

gagcgtctcttgtaagccaaggtgaagatgaaggcggccgcgatttttatgttgtccgcagctctgatccttcttttctgcttcaccactcaagggcgtaaaagttcaggaagaacttccagcatagtcaagaggactgaccatggagattcttcgggatcgctgcc
gggtggaaggaagcacgttgtttgtatttgtgaccagaagaagcctcataatctaccaagaaaaagacgtcatatgaaggctcgtcgtagacacggtcatcataggagtgtgtgtgacaggatctgcttgacaaaaggtgcaccaggaggactggcggtgccc
ataccattgacttaaaggaactctaaatgatgacgtttcacaaagaaaactcacaaaaccgtctcgttatcttttctgtagaatccgggaacatggacattacagaacatttctattaagggagcactccagtctacagtctatgcacatataggtataatcaatagattta
caaggtgactcgactgtatatatcacattaatcctcttgtaatgatcttaagttacagcatattattctccagagctgcattcacagttctgctgctgattataggaaacttgtttttcagtcccctccccacaatatcttagctgagctcctataatatagcaggtcagtttgtc
tttggtgtagaaggctgtacagagcctggtggaaagacaacccttcccagaatgcaaatcaccagaacatattctatgcaaatgttgaaccagcacagaatggtgagggatttcagctctgaagcttgccaaaactgttaattcagctcctggatataatattgggat
tcaaaagccatccatttaataacctactaccgagccgtgaattccaaataatgaggttaaatctatatttcagcctctactgtttaacctgcaaaaatgattctaaagcatgaaattttaaaaatccttcaaaaattaaatttcataacttaaataaaaaaaggagctgaggaa
agacaggagagagtttcccaaagtaaagcgaactcatctgtaagaaggcaacaagcttagatcgatactaacgtcaatctgttctcccttagtctctcggcagcacaataatggggagtctccgccaccgcttactcttactgggcggatggagttttgaattaaact
aattagacattaaatagagccctttcaccctctaaaactacatcccgcccccacctgctgtacagatgcagttgagctaaacttgacggttaagtaaaaaatgccacaaaaaggttaactaataatttcaatgtattttgtcacaagatggcgcgagataacactgtga
catgtgttatcagagtcaattgtagctcggctagatcagtatattataaacatcataaagaacacaattattacagcagcaagcatgtgctgcaggactaagggaaaacagacactctccacttaccaatgtcctaacagcggacattcttgcccctgtgtactagtag
gacagatggaattaaacattacttagctagaggcccctgatccggaagaggacatgcagccctctctgtgtgttcattaaaaagatgtaacttaccccggataggaaatgtgctgcagttgggactaaggaaaaacagattaaagtaaaaaaaaaaatgcttcccc
tatgccctaccagcagcacaataatggagagttatcaagaagtatccctatggggaggaccatttcatctgcctgaactcaaaatcagaggataggttatcagtatctgattggttggggtccgatcagctgtttgagaaggcagaggcactcctgtggccttctctc
tgcttttactaggccagcaatggtcatgttggctaggagcagctcggccctcttcactgatctgcaataccaagcacagccgctatacaatttgcggggctgggcttggtgagctgcaagaagcctgcagtgctcacaggactcaagtggacttagctgaaagttg
gactttgatgaacgttggcctccttcatatgctcatgcagagagcgagggatgttaatcacagattagaacgttcttgagtagaacaccaagccaaacaaaccttctttaatagtccacactggacaggtggagcacaaggaggaggatcttgatgtcctttaaggg
cctaaaagtaattagtaattacttaattaaaaattccatagagattccataagtcatagagacccacattaatgtgctgctgataggcacagggaaaggccagatccttaggttctcgaatgatgcattctaccatgtatattgtgtgtgggtttggggtctgtgtgagttc
agtgtaatccttgtattgtcacatgcccgtcttattgtgacacagaacatgtgattgtgtctccagttcaataaacatttttatattttggca

**D:** Amino acid sequence (signal peptide shaded)

MKAAAIFMLSAALILLFCFTTQGRKSSGRTSSIVKRTDHGDSSGSLPGGRKHVVCICDQKKPHNLPRKRRHMKARRRHGHHRSVCDRICLTKGAPGGLA
VPIPLT

**Fig. S13.** Gene position and architecture (**A**), genomic DNA sequence (**B**), cDNA sequence (**C**), and encoded protein sequence (**D**) of the unannotated *cxcl17* gene in the NCBI reference genome of *Bufo gargarizans* (Asiatic toad). The information was downloaded from the NCBI gene database (https://www.ncbi.nlm.nih.gov/gene).

### A: Gene position and architecture

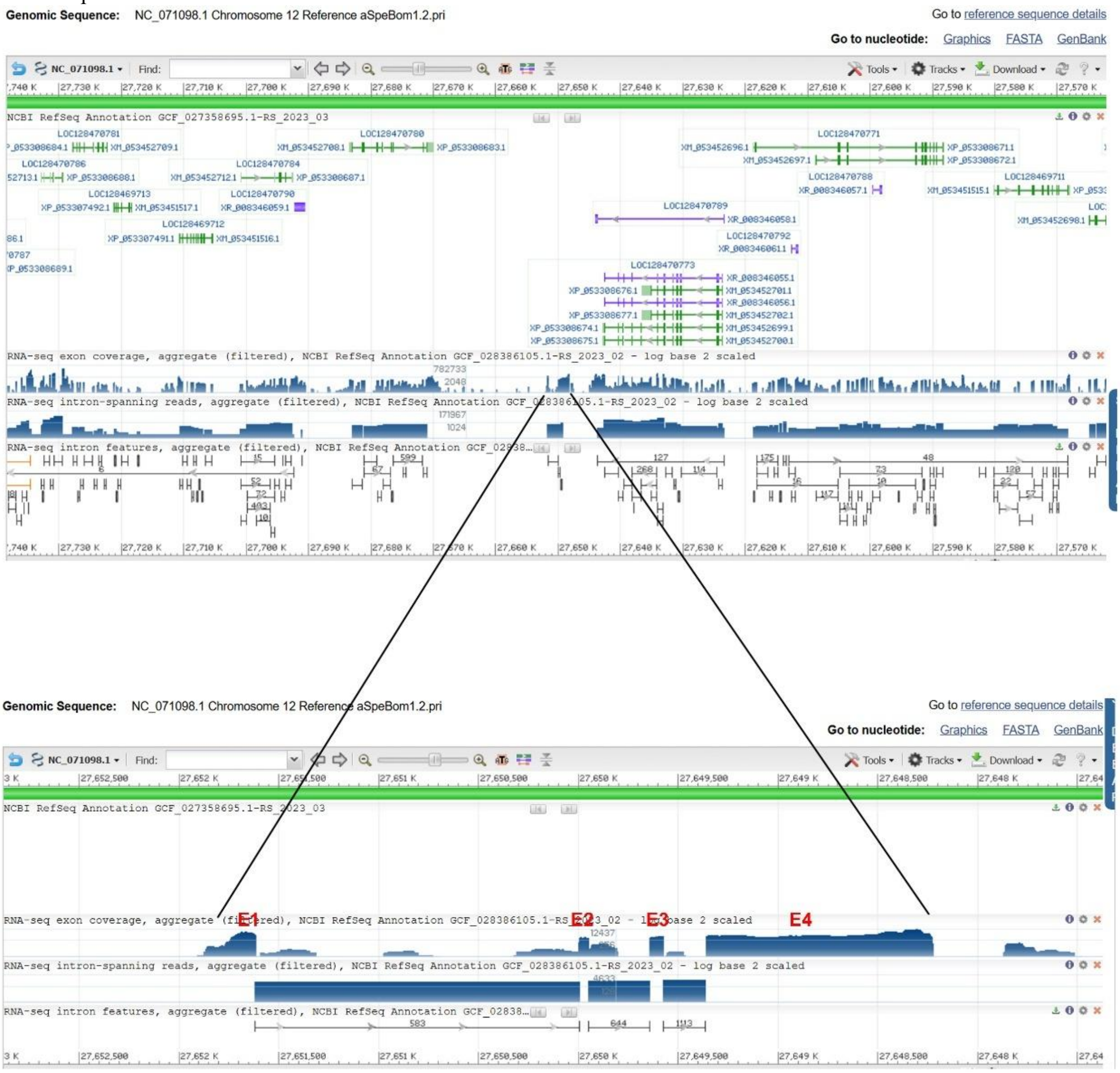


### B: Sequence of genomic DNA (exons in red and highlighted in yellow, coding region underlined)

tgtggcggagagggttaatgttggggatacttaacattccattgtgaaaacaaacctgctaatcgagacctgccccacaacatatgaagtgcaaggcggagagacttagagagtcaacgaggggtagctgtccttcacacctgaacttccgtgaggatgaagac
ggtcgctgtctttctgtttcttgcgtgcttcgccatctgcaattgctggacaccacatggacaggccaatgcaggtaaactgaaacctctaggactgaacaggaacgatgacatttgcaatctaggacccttctataaaaataacatggacgtctttcaaggggaatta
ggctactgacggataatatcatcattttgtaccggaggcaatggggtttacagtgattaaatggataaatgtggcattaacctaaccgctaaacatgtattaatgtgtaacgatccgcgatgatgtgccaaaatagcataaagcaatgtgagaagcttgcctgtgtgtt
atagggaaatgtataatttccgattatcaggattatcaggatccgcatgattctccccccttaattatttatctcaataagttttccctacctatataggactcttctataaaaataacatgtcaactgatggtggccatgataaatacggcgaattccgtctatgaccaactg
attctaaaaaattacatattctattgttcaacatttcagcttaaacaacagttctgaaaatcacttatactgtgcggtctataaatgtacatgtgcgtcgctaacaaaagagtgaatactttacaacataaatactggctttcaagctaagccagggatcaacgataggtatt
aggaagggtggtgtatccagcgactggagggtcagacgtgtgttttaatagtgtctctgaacaggcatcagatcatgtgcattatggtgatggtttgcaggccaaagctcactagagtggtgccaacagccatgtagtgcagactgaacacttacaggaaaccaa
cacgagcttctctggtggcagagctactagagaaccagaagagggagctttggttgctgattaaacactcaacattgcatatgtattttatttttaatttatgttgctaaccaggtagtgctggcatatcatgtgatttccatgcgtcttctgttgtttgtacctggactgga
agtttgtaactatgaagattgggatatctatcatgtctagttaaggtcccatctcatcaaaaaagtcttgatggtggaagagaaggagttattcattgggcaatagattgaattaagactattggacatggaaatatttccttactaatgtcccctcgcttgggtatattgta
gatcatacccaaggcttagctttatgtttaatgagtagttagttatacacatacctgggaactcactggttgaagcagcgcttcttcggttctctgggtcaacacccgaatcctccaggttgcggaagcagggtcttgacacaccctgcttcccgcccattttccctttaa
atgggggtgtttcccgctgacatcagcgggaccacccgctgatgtaagtgggcagtcctggccgcatcccacaagggaaggctccaggtagccggattgaagaagttcccaggtagggttctacgtatgaaagagcatttctattttagatttcaatattaaagtt
ctgtgcagcttaattaaaacattttgttgtggaatctaatgtatagcggatgtttaaaggttccagaatgcaatgaactgttcccaagatcctgcttgatagttttagacagagatacatatcagatggaatccataaatatattctcacattttcttcaacagggaagagcg
aggacagcagtctcgccataaaaaatacaaatataggtaatgtatgaaaaattgtggtcagattcttaattgcaaaactataaagagtcatcatagctaacccacacatgtacattgagaaatatggcattttagatcttctagatgtgacaatcaaaaagtaaaaacaa
caaaagcttggattaaaaaataaaataaaacataaaaataaatttaaaagtcttcaggtactcatttatgttaggggttcccgtaggtacatagttagctagaaccctttatactctaaagttgtgcaaataagataagaataagcacaagctatattattttctcattcagat
gcccgtggatatgaaagaaagatcataagatgcaactgcaagagtcacggaccccagaagctacatcgtaagacacaatttatcttggaatatgacaatttctttttattggtgaaggagtctgtcattattaatgtccatgtcagatggccacatcaatgagtttatga
gtttccatgtccaaaatatatatatatatatatatatatatatatatatatatatatatatataataccgcttgctactaacaaagcctttcattttcctgtccttcaaagagaaaagacgtcacaagaaagctgtctgctctaagccttcccatgatcatcgatgccgccatatctg
caatcggatccgtctgaagcatgggaacccctttgatgtctctatgcctctatcctgaagaacagtctccatgtgaagatgacggaaagccacctctctatctcatatactacttcacctcccattatcatcacagctttcacaaatcactgatatctatcgccgccacct
ggaacaccacatcacgcaaaaggatttctctttagaaacatattaggtaactgttacctactattatgatatatagtccattatacgctaagaatcacgtggcatataaactaatctgggagtttatatgtgaaaacctaaattaaataattccatttaggattatgatgggtc
aaccctcatggcctttacttgcaaacggttagtgaatacaaccctgcactataatgactagagtagaatatttacttggtctggccctggtagatataatggcaacaaaatataaatctagaaatatacttgctatacgtgaaacgctgcccccattggcactctccatc
gtgctccaaaattccatagaaatatacaataaggatggcgcaggatcattaaaaacaataaaacccataaaccatcatagtttatgtcaataaatctacaccaatacgtaatgtatcaatgatgcataaaggagctcatcaaaagagaaagcaatcagaaccaagat
attgcctattgtaaaacccaaatgcgtggtggttgagatcaaagatctcacttgtaactggcccattgtgcagcaagagaccaggggggcttgatggccaaagaaggtgactactgcctccccccaggtcctgccactctcgtatatgccgctgccgaccacctttta
catattgaatgcgtatatatctgaggtgaagatacagaagaagcttttgatgtagtagatatagcaaccaccggagacaatccaccctacgcatttcacttagtagacagacgtagtatttgctgtccatacatcccaagaacatctcatttctatattttacattcaattg
ctcccgtcattgtttcctattttctaataaaatgcttaagcacaaaaataattccagttctcctgttatttctgaacacacgtagtcagaattagtaaatgtgaagactttagtatatt

**C:** Sequence of cDNA (coding region in red and highlighted in yellow)

acttagagagtcaacgaggggtagctgtccttcacacctgaacttccgtgaggatgaagacggtcgctgtctttctgtttcttgcgtgcttcgccatctgcaattgctggacaccacatggacaggccaatgcagggaagagcgaggacagcagtctcgccataa
aaaatacaaatatagatgcccgtggatatgaaagaaagatcataagatgcaactgcaagagtcacggaccccagaagctacatcagaaaagacgtcacaagaaagctgtctgctctaagccttcccatgatcatcgatgccgccatatctgcaatcggatccgt
ctgaagcatgggaacccctttgatgtctctatgcctctatcctgaagaacagtctccatgtgaagatgacggaaagccacctctctatctcatatactacttcacctcccattatcatcacagctttcacaaatcactgatatctatcgccgccacctggaacaccacat
cacgcaaaaggatttctctttagaaacatattaggtaactgttacctactattatgatatatagtccattatacgctaagaatcacgtggcatataaactaatctgggagtttatatgtgaaaacctaaattaaataattccatttaggattatgatgggtcaaccctcatggc
ctttacttgcaaacggttagtgaatacaaccctgcactataatgactagagtagaatatttacttggtctggccctggtagatataatggcaacaaaatataaatctagaaatatacttgctatacgtgaaacgctgcccccattggcactctccatcgtgctccaaaatt
ccatagaaatatacaataaggatggcgcaggatcattaaaaacaataaaacccataaaccatcatagtttatgtcaataaatctacaccaatacgtaatgtatcaatgatgcataaaggagctcatcaaaagagaaagcaatcagaaccaagatattgcctattgtaa
aacccaaatgcgtggtggttgagatcaaagatctcacttgtaactggcccattgtgcagcaagagaccagggggcttgatggccaaagaaggtgactactgcctccccccaggtcctgccactctcgtatatgccgctgccgaccacctttacatattgaatgcgt
atatatctgaggtgaagatacagaagaagcttttgatgtagtagatatagcaaccaccggagacaatccaccctacgcatttcacttagtagacagacgtagtatttgctgtccatacatcccaagaacatctcatttctatattttacattcaattgctcccgtcattgttt
cctattttctaataaaatgcttaagcacaaaaata

**D:** Amino acid sequence (signal peptide shaded)

MKTVAVFLFLACFAICNCWTPHGQANAGKSEDSSLAIKNTNIDARGYERKIIRCNCKSHGPQKLHQKRRHKKAVCSKPSHDHRCRHICNRIRLKHGNPF
DVSMPLS

**Fig. S14.** Gene position and architecture (**A**), genomic DNA sequence (**B**), cDNA sequence (**C**), and encoded protein sequence (**D**) of the unannotated *cxcl17* gene in the NCBI reference genome of *Spea bombifrons* (plains spadefoot toad). The information was downloaded from the NCBI gene database (https://www.ncbi.nlm.nih.gov/gene).

### A: Gene position and architecture

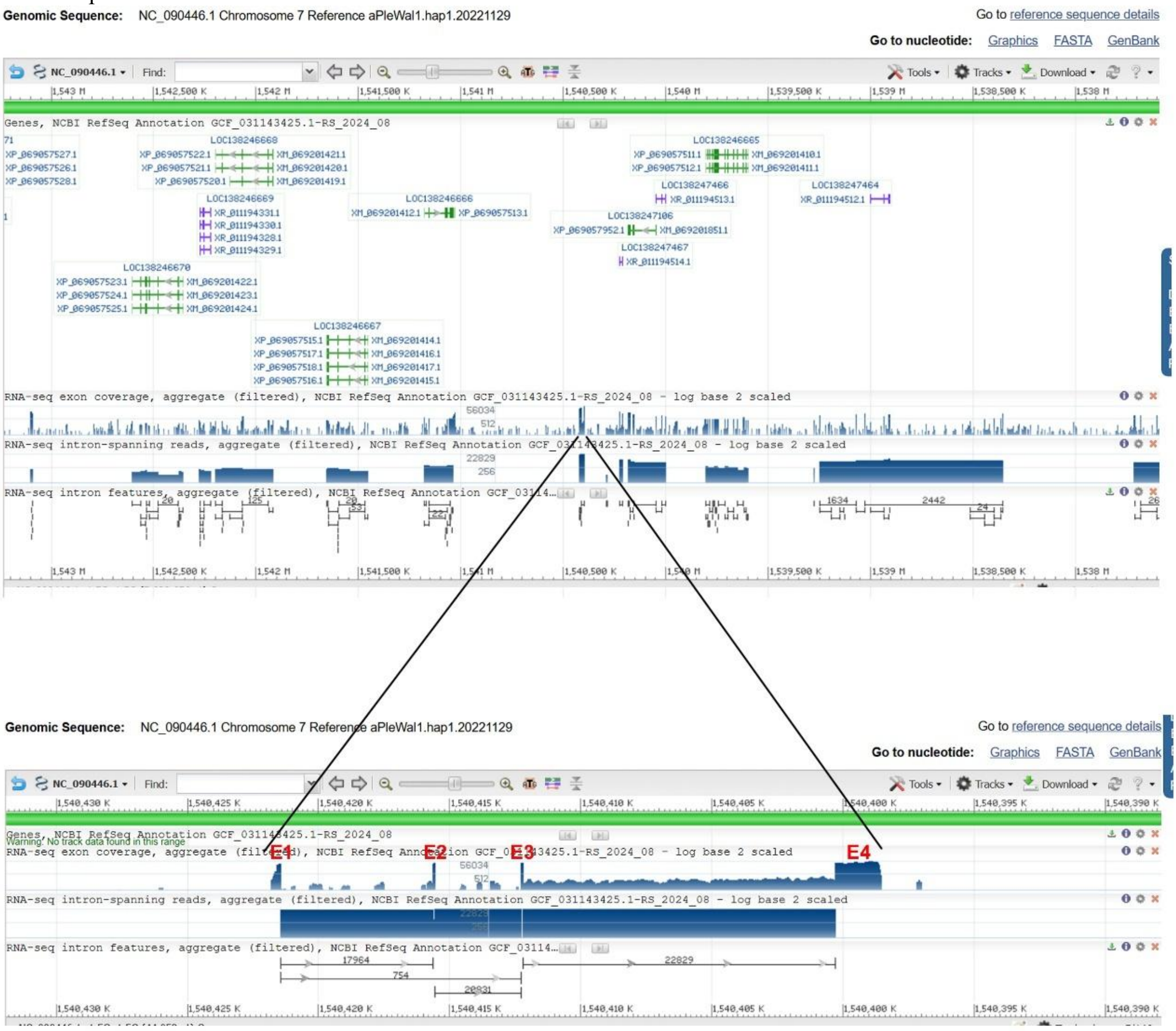


### B: Sequence of genomic DNA (exons in red and highlighted in yellow, coding region underlined)

ggcggggagagggggcggcctctgtgtccacaagccagtctgatcctgcctggtagatgtttgcttcgtggccagagatcttatcggtccgggctcctcgggggttgggccagtggccagcaggggtgcgagccgggctccttaaaagacaagccttcttgc
caagccaccttctttcggcgcagacgcccctgggtcggatagagcgctctaggggctcccattgtgtctgtggtcggtccctgcacaactatcgacctcactaggatgaggctctccctggcactggcggtgctgctagtggcttgtgtatccctctgctggggc
accgaggtaagacacccccgtcctccaaggtcgagtttgtgctattcccagctctggatgtaccgtgtgcagagtcccggcgctgtacattgatcgcatagaagccttgggatggagggaagcgctgtgcttgttttggggtgggtgaatgttgtgctttgcacgg
acgctttaggtgaggttgtgtcggggcctggctggcctagggttcccctctgaactcaatggatctgaggtcctatgaccccctaaatctcatatttgtcagttctatcccgagtgtaagtaacacaagaaactgagtgccccctgacgcacacacaccgggtcaccc
ccttgggctccctgcatgcccagggcagccacctgccccaggggtgggtgggattctgttcactagagggagctgtctggtccgatagggctcagtgccaccacacacacacgcgcgcacacacaccgggtcacccccctggggctccctgcatgcccagg
gcaaccatccgccccagggatgagtgagggtcgctgggaaggggatttctggtagggctgagtgccccctgatgcacacacaccgggtcacccccctggggctccctgcatgcccagggcgaccatccgccccagggatgagtgagggtagctgggaag
gggatttctggtagggctgagtgccccctgatgcacacacaccgggtcacccccctggggctccctgcatgcccagggcaaccatccgccccatgaatgagtgagggtagctgggaaggggatgtctggtagggctgactgccccctgatgcacacacaccg
ggtcacccccctggggctccctgcatgcccagggcaaccatccgccccagggatgagtgagggtcgctgggaaggggatgtctggtagggctgagtgccccctgatgcacacacaccgggtcacccccctggggctccctgcatgcccagggcgaccatcc
gccccagggatgagtgagggtagctgggaaggggatgtctggtagggctgagtgccccctgatgcacacacaccgggtcacccccctggggctccctgcatgcccagggcgaccatccgccccagggatgagtgagggtcgctgggaaggggatttctgg
tagggctgagtgccccctgatgcacacacaccgggtcacccccctggggctccctgcatgcccagggcgaccatccgccccagggatgagtgagggtagctgggaaggggatgtctggtagggctgagtgccccctgatgcacacacaccgggtcacccc
ctggggctccctgcatgcccagggcgaccatccgccccagggatgagtgagggtagctgggaaggggatgtctggtagggctgagtgccccctgatgcacacacaccgggtcacccccctggggctccctgcatgcccagggcgaccatccgccccagg
gatgagtgagggtagctgggaaggggatgtctggtagggctgagtgccccctgatgcacacacaccgggtcacccccctggggctccctgcatgcccagggcgaccatccgccccagggatgagtgagggtagctgggaaggggatgtctggtagggctg
agtgccccctgatgcacacacaccgggtcacccccctggggctccctgcatgcccagggcgaccatccgccccagggatgagtgagggtagctgggaaggggatgtctggtagggctgagtgccccctgatgcacacacaccgggtcacccccctggggct
ccctgcatgcccagggtgaccatccgccccagggatgagtgagggtcgctgggaaggggatgtctggtagggctgagtgccccctgatgcacacacaccgggtcacccccctggggctccctgcatgcccagggtgaccatccgccccagggatgagtga
gggtcgctgggaaggggatgtctggtagggctgagtgccccctgatgcacacacaccgggtcacccccctggggctccctgcatgcccagggcgaccatccgccccagggatgagtgagggtcgctgggaaggggatgtctggtagggctgagtgccccc
tgatgcacacacaccgggtcacccccctggggctccctgcatgcccagggcgaccatccgccccagggatgagtgagggtagctgggaaggggatgtttggtagggctgagtgccccctgatgcacacacaccgggtcacccccctggggctccctgcatgc
ccagggcgaccatccgccccagggatgagtgagggtcgctgggaaggggatgtctggtagggctgagtgccccctgatgcacacacaccgggtcacccccctggggctccctgcatgcccagggtgaccatccgccccagggatgagtgagggtcgctg
ggaaggggatgtctggtagggctgagtgccccctgatgcacacacaccgggtcacccccctggggctccctgcatgcccagggtgaccatccgccccagggatgagtgagggtcgctgggaaggggatgtctggtagggctgagtgccccctgatgcaca
cacaccgggtcacccccctggggctccctgcatgcccagggcaaccatccgccccagggatgagtgagggtcgctgggaaggggatgtctggtagggctgagtgccccctgatgcacacgcaccgggtcacccccctggggctccctgcatgcccagggc
aaccatccgccccagggatgagtgagggtcgctgggaaggggatgtctggtagggctgagtgccccctgatgcacacgcaccgggtcacccccctggggctccctgcatgcccagggcaaccatccgccccagggatgagtgagggtcgctgggaaggg
gatgtctggtagggctgagtgccccctgatgcacacgcaccgggtcacccccctggggctccctgcatgcccagggcaaccatccgccccagggatgagtgagggtcgctgggaaggggatgtctggtagggctgagtgccccctgatgcacacgcaccg
ggtcacccccctggggctccctgcatgcccagggcaaccatccgccccagggatgagtgagggtcgctgggaaggggatgtctggtagggctgagtgccccctgacgcacacgcaccgggtcacccccctggggatccctgcatgcccagggcagccatc
cgccccagggatgagtgagggtcgctgggaaggggatgtctggtagggctgagtgccccctgacgcacacgcaccgggtcacccccctggggctccctgcatgcccagggcagccatccgccccaaggatgagtgggattctgttccctggagggggggttt
gtggttcgataaggctgagtgcctgtcccaagcccccccccctcacacacacacaaaccgggtcacccccctggggctcccgcatgcccagggcagccacctgccccagggttgggtgggattctgtttccaggaggggggtgcctggtcctatagggctc
agtgttaccacacacacacaccgggtcacccctggggctcctcgcatgcccagggtgaccactcgccccaggaatggctgcggatcgatggagaggggtgtctggtccggtagggctcaggcccccacccccctcacacacacacaccgggtgggaca
tcagcaaagcttagttcactgtgcggtcttcccgagacctcgcgctgcggggttgatgtggttggttgcgcgttgtggggttgttttgtgatcccttccgtgttagtgagagcttgatgttgtgtggtggagttgtatggcggggtgtgtggttgggactcgtgatgccc
tgtgagctgtgaagatgtcccgggctctgtgtggttggcacacggccgggatttgttgtggtaacgtgtcaccttggatggtgagtggcgcagagaggtggccccggagcctaacacaagtgagggaagcctgcccgcataccccccccccataggttcac

agtaagacgctgatgttatgtgtgtgaagtgagcaccgctgccctggtgccactgcacctgcagcactgttggtagatgcaccctgggaactgtagagaagaggggtggggggtgaaggtggtgcgtggagggcaaatgggggctggtgctgtgtaccagg
cttgtgctgtgatgacttgtgcgttttgtgacagtgcctagcgggggctggtgctgtgtagtacgtgtatattgtggtggtttgtgggtgatgtagtgatgggtagtgtactcacaggtgtgtcttctgctcgaggagtaggttgtggtgagtgtgtactcacaggtgtgt
cttccacttggtgagtaggttgtggtgagtgtgtactcacaggtgtgtcttccgctcggtgagtaggttgtggtgtgagtgtactcacaggtttgtcttccgctcggtgagtaggttgtggtgagtgtgtactaacaagtgtgtcttccgctcgaggagtaggttgtggtg
agagcgtactcacaggtgtgtcttttgctcgaggagtaggttgtggtgagtgtgtattcacaggggtgtcttccgcttggtgagtaagttgtggtgagtgtgtcctcacaggtgtgtcttccgcttggtgagtaggttgtggtgagtgtgtactcacaggtgtgtcttccg
ctcaaggagtaggttgtggtgagtgtgtactcacaggtgtgtcttctgctcgaggagtaggttgtggtgagtgtgtactcacaggtgtgtcttccgcttggtgactaggttgtggtgagtgtgtactcacaggtgtgtcttctgcttgaggagtaggttgtggtgcgagt
gtactcacaggtgtgtcttcttctcgaggagtaggttgtgggtatactcacagatgtgtcctccgctcggtgagtagcttgcgatgagtgtgtactcacagttgtgtcttcctctcagtgagtaggttgtggtgagtgtgtactcacaggtgtcttccgctcggtgagtag
gttgtggtgagtgtgtactcacaggtgtgtcttccgctaggtgagtaggttgtggtgatactgtactcacaggtgtgtattcttctcggtgagtaggttgtggtaagactgcactcacaggtgtgtattctgctcagtgagtgagctgtggtgagtgtgtactcacaggtg
tgtcctctgctcagtgagtaggatgaggtgagtgtggcctcacaagtgtttcttccactcagtgagtaggttatggtgtgagtgtactcacaggtgtgtcttctctgtgagtagagtgtggtgagacggtactcacaggtgtgtattctgctcggtgagtaggttgtggt
gagtttgtactcacaggcgtgtcctctgctctatgagtaggttgtggtgagtgtgtactaacaggtgtgtcctttgctcggtgagtaagtcgtcgtgagtgtgtactaacaggtgtgtcctctgctcggtgagtaggttgtagtgagtgtgtactcacaggtgtgtcttcc
actcagggagtaggttgtgctgagtgtgtactcacacatgtgtcttccgctcgatgagtaggttgtggtgagtatgtactcacaagtgtgtcttctgcaaagtgagtaggttgtggtgagagtgtagtcacaggtgtgtcttctgctctgtgagtaggttgttggtgagtg
tgtactcacaagtgtgtcttctgcaaggtgagtaggctgtggtgagagtgaagtcataggtgtttcttctgctctgtcagtaggttatggtgactgtgtactcacatgtgcgtctctgttcctcaggcccccaaaaggacgagaaacgactagtcaagaagcacccct
ctcaatggaggcagcgccgggcacacagtgagtgctgggagggggctgagtaggtgtgtgtggggagggtagggaggtgtgtgtgtggggtgtttactagcttgtggaaagtggggaaatgtgtatgggagggggtgtatgtgaggatgaggaggtgttttg
ggggcaggggtgtgtgtgtttgggggcagggagggggtttgtgtgtgggggtgggaagggttgtgtatgtgtgggaggtggggagggctgtgtgtgggggtgggaaggggtgtgtgtggggggaggtgtgtgtgtgtttggggaggcgtgtatgtttggagg
ggagatgtgagtgtgaggggggaggggaggtgtctgttggggtgggaaggggtatgtgtgaaggcgggaaggtgtgtgttgggtggggaggtgtgtgtgggggtggcaaggtgtgtgtgtggggggggcagggaggtgtgtgtggggggggcagggagg
actgtgtgtgtgggagcagggaggtgtgtatggggggagtgggaaggggagtgtatgtgtgggagtggtgtgtgtttgaggggtgggaaggtgtgtgtgaggggtgatgtgtgtatgtgggggtaggtgtgtgtgtgtgtgaagaggcgtgtgtgtttggagg
gggagatgtgagtgggaaggtgtgtgtgaggaggggtgtgtgtgagggagggtaggagtgtttggttgtggggaggtgtgttttgcggctgcacggtgtctgtgtgggagcagggaggtgtgtgtgtggggatagagaggtgtgggtgggggcagggagg
agtgtgtgtggatgtgttgtgggggcagggaggtgtgtgtggggcagggagatgtgtgtaggtgtgggaaggagtgtgtgtggggggtggggaggtgtgtgtgcatggggtggcgatgagttgttatcagaaatgtttgtgtcattcccactctccgcagctctc
agtgttacacccacacacaccgcagcaggcacaacaaccccctcagctatctcagcttcaggtttcaccacatgcccgaggcttgtggcctcccagggagtgtgtggccccttgttgagaggagatgtcaccagggatttctggaaccaggggaaggttcacaa
ctcacaggaggtagagcccctgggggtggggggacgaggaggcgggacggccccgggtcgggaggcagaagtcacgctctattcacagaacaaggcggacaagcatcctccacagtcacagcatgtgctcaccgcagggcacactttcctctctgaga
agcggtgacgcacaggccgttggagcttcctgcgcatgcacaaacaaacactggtagtgcatagtggtgcaagcttcctgtgcacgcacaagcaaacacgggtagtgcatagtggtgcaagcttcctgtgcacgcacaagcaagccctggacgtacagtagt
ggtgcaagcttcctgtacacacacacacacaagccctgggcgggcattagcagggcaagcttcctgtacactcacaaggaagccctggacgtacagtagtggtgcaagcttcctgtacacacacacacaagccctggggggggcattagcagtgcaagcttcc
tgtacactcacaaggaagccctgcacgtacagtagtggtgcaatcttcctgtacacacacacacacacaagcccctgggcgggcattagcagtgcaagcttcctgtacactcacaagcaagccctgggagagcattagcagtgcaagcttcttgtacacacaca
cacacaagccctgggcggacattagcagtgcaagcttcctgtacactcacaagcaagccctggacgtacagtagtggtgcaagcttcctgtacactcacaagcaagccctgggagtgtgttagcagtgcaagcttcctgtacactcacaaggaagccctgcac
gtacagtagtggttcaagcttcctgtacacacacacacaagccctgggtgggcatcagcagtgcaagcttcctgtacactcacaaggaagccctgcacgtacaatagtggtgcaagcttcatttacacacacaagccctggacgtacagtagtggtgcaagcttc
ctgtacacacacacaagccctgggagggcattagcagtgcaagcttcctgtacacacacacaagccctgggagggcgttagctgtgcaagtatcctttacacacacacacaatccctgggagggcactccgttacacacactagcaagccctgtcctaagcac
aatcttgcaccgtatgagtagctgaatgcatgcttcctccagggctttaagtgggccgggtctgtcctggtctcagtcccggtagtgttttagaacggacctctctcttttttgcggctgattgtactaagtgcagagttggggtccagaggagattggttgtagggtgtg
tgggggtctcggccagcaccgaagctggagggggtgccctcgggttccgtactctgatgactttagaaagggtggtggggacatgtctggagctctcacacagggaatgttacattcagtttatgcagatgtcgccttgttcagtctttaggcgctagtcattccca
agccaacccatactgctctcagcatcacacgagtcatgctgacaatggcatgtagcgcatcattaacagtacagctggccttcaaggcgctgtgctgtcgcagtattgtcctagcagctcgacggatagcaataaatatagcgcttctgcggtgaagagatggtaa
taccctcagacctagaaggctcgcagccctcccactgaggcgcttcccatactcatgtggtaccaagccccgtcagtgcccggggctgccccatccgtgccaggggatgatctatcagtgccaggggctgccctatcagtgtaagggtcttccccatccgtgcc
aggggatgccttgtccgtgccaggggcttcctcatctgccctatctgtgccaggggctgccccatcactgccagggaatgatctatcagtgccaggggatgccctgtccatgccaggggctgccctatctgtgccaggggctgccccatcactgctaggggatg
atctatcagtgccaggggatgccctgcccgtgccaggggctgccttatcagtgccaggggctgccccatcactgccacggctgctccgtccgtaataggggtgccccatcagggggcaggctgtgtgcctcgtattgtactgtatcttttgacgttgttttccttca
cttcttctaactccaaggatgctttgtccttcagctgcccttgaagacaaaagacccaggaggatggaagaacccgtgagctcgaaaggggtgaaggactgtccgtgccacgtcctaaaacgtgagtaaaaccatcctctccatacgcacacacccaaaacagc
ccaacccccaatactgctgtaacaatggtccaaggtaccccttggaggaccagccggcttgtgatgctgccctgcaccccctgaagtgagtacggaagcgggggggagggtcacccctgtgattaactagtttaggggggtgagcattgcccgtggcccctcctct
gtcccagtccttatcccatgaggtgccccgcctcctggaagaggtttattttcacccaagcctctctgaagacacttccagccggtacttccggtttccactctactctccgcttcctgtatttatggcccctccccctctcccacctcctactgttccagtcccctggag
ggcttccgagtgagggaagcatttttaattaaatgctatggggcatagttacaagccccttacgtagctctagccgcatattgtcactctaaggcggtgctaaggtggcctttctcctgcaccagatttacagagtggcgcaatgcatgcattgcaccactttgtaacc
ccctgcgccatattatgtctccgtcaggcataatgtatgcaagggggcgttcctccattaggaggcccagaaaaacgacacactgaaatttacaagatttcactgcgtaatttttccgtcatttacattttcctgtcatttttaacgtctgctctgtactttgctgcactag
cgtcaaaaatgtttatgctattgtgctgatgactgtcgtggttcgtcttattataaatacgatgcaaccatggtgttgttagggagacgcaagaaaagtggtgcgttgggggcgatgcacaaatgtccccccatgggcctcatttacaagatgcatcggcccagatgt
gtcagatttcttgtgctgccctacaacccccctgatgaccccatggatacgctgtatttacaatatggagcttcatggcacacgtttgcacagatgcgtcagaaatcttgatgcagctgtggccctaaagtcccgcattgctagactggtggcacaaaaatgacgctac
ttcagcaatgagaggaaggccctcatgttaaaaagcccggcgtcaggttaactctgctctaaacaggcagtaaaaccctgacacaatgaatgcgcagaacaacgcagtgaaatgatctaaatttcactgcgccaggtctgtgtggctttttacgtgggaacgccc
accttgtaaatattatacctggtgcagggataatgtggcacaaggcttttgagaccattgcgccagaaagtaaatgtggcgcagtgtaggggactgcttagcaccacgtggcatccaaaaaaagatgcaatggtctcgctaggagcttgtaaataagccaatatatt
ttaatttgtggttgtattattgtgtatagcaagcggtggttcgagcgcttctgggcactttatcttctcccggtccacatgctgaggcgtgataggagtcttggagcaaaattactgctatgtaagtacctctctgcatagttaccaaagcaccctcgtagaagcgcagca
aactgttcccgctgtctcgaggtagcactctaggcactaggcgggctggggacaccttatagccatccaccccgtcccgtgacctgagcgacccttggctggccagccctctcataccccacacgttctacctgtgcaaagtttctgagccagcctgcacaatgg
tggtccacacatcccaaaccctgccctgagccctccttggctggccatcctcttccataccttgctcatcccacctgcacaaagtttcccagccatctcacactatggtggtccacacatccccatctcggatgtgtgccctcctcagctggccaccccatacctcttt
ctttttttgttatgttttatttgtattttatttcataaagcggctgaagcttgcgaagcttgcacacttttaaaaggaatcttgaagaatctgatgagggaaaacttctgtaactgtgcacaacactaacaacagacacggtgccctgcacggcgaggtaattagatagtaca
aaaatacacaaaacaaaaaagcaaatacaaatattaaaagctatgtactaaaaaaaaatctggatctaatttaatttaatgctaataagtgtgggaaggtaagattttaggaatcacttttgtggtagttgacacaaatgttgacagttacacaactcctattgtctacattta
ttaagaattaaatgtgccatccagtttcctttttttcatttagaaactagatcacaaacccacatctggcatattgcgggcataatcacaatgcataaaacgcaggagtcctgaccatgtgtctatcaatgcttgagaaggagggagcaggctagcgccatatgtaaca
tcgccagggtaggcagagccccaccttccaccctgattggcacaagcccttaggattaaagctcagtaccttttttgtgtctggctgtggtttgctttaccgctctgtagtttggtgcctgcccccgtctgtggtacgaacctatgtaagtggagatcttctggagtgag
ggtccttgtggagattggggtgcggtgaacgccgttatcctggctcctcaactgagagcggaaggggagaacaaacggggcgacttccatgttcctgtctcggcccagattcaccaaatagtcgcgcagcgcaatgcagcagtgagagacgccgcgctgctt
gaggaggcgggagcaggcgagcgccatatctacagaggtacgactctctcctcccctctccctagcgcccgcacagaaatccgctgccgtgcaccccgctcacacctttcaccgtactgcaagggtgtgtgcgtcagtctagcgtaggtttctatgggaaggg
aacccttccagtacaaaatactatgcgtagacacatgttacaactttgtctacattgtgcatgtgccgcacagtgcaacctgcatgcaaagtcagaaaacaaaggaaaaatgaaaacattagtccttttaaatgctgatttcacatggcgttaacctttgacacaaaac
caagtgcactattgtttgtaaatagggtcttgcatcaaaaggcatgggtgcttgcatgagaatgcccatataacacccccttggcactaagcaaagcagtgtttagcgctgccttgcgttacttttacgacactttccatcactttttatgatgcaaacccaccataaaatc
ttaaacacacccctagtatctccagagctccctcaggctttgtgggtcttcctgggcaggtcggggtcgtcagcggagcgtcactcctcgcggctcacaccaagggcgcaggaaccctggcccaccgtggaaggtggcctgtgtggcgaacgccatgatgac
cgtggctcccaaggacagcgtacctcgctgaagctctttcttccaggacagcgggtgctctcctgctgaagcacactttgctgtgtgtaactctccattgctaatttaacaaggggagatcacccatcctctggcctgggccacgatgtcaggcacagtattcagcc
tccagatgcatgtgaggtgccctggtggttccccggtgctttcattgtgtcatttggtgacaagtattacctgtggtacccctcacccttccctgatatatcacatctgtctcaattaaaggctgaagcaagacatacaccttgggctgtgctttatagtgggcaggggg
gtccatccgctgcttaggaccttctgaagggctcccagtgggggatgcaacctatcggagggtacagtcctgggggatcctcttccctctgagggattaccagtggggaatgtgtggccagacaggtgtttcaggtgatgctttgtggttttacagctgttcaagtc
ctgagtgactgagaagcccaacctgagctttgcagggaagaaaaacattctgttagtggttcaaagtttcactgattgtgctgccaatgagtgatttcagtgagggaactacagcctgtcagctgtctaggcccttacaaactgagtaccctgtctggcattcaagtc
aaggaatgcatcctgccagctgctggtgctcagacaaagtgtgtacccacctggcacagaaggtgatgggatcatacatctcctgtgtattggttacacaataacacactgtgctacttgtctcattgaaagtgatgggagcacacacgtcctgggagttgttcaca
tcttaacacactgtgctaactgcctggcactgaaagtgatgggagcacacacatcctgtgtgctgtggacacacatcttgtgtgttgttcacatcttaacacactatgttgcctgtctggcattgaacgtgacgggatcacacacgtcctgtgtgttgtgcacatcttaa
cacactgtgttagctgtctggcattgaaagtgaccggagcacaccagtgcttaatttcttaataaaaacgtgcctgtgcctaaagccctcctctgaaacacgcagctgctgcaaagaaatgtgcgaacacggaatactgcggcggcgtaatcctgaagccatctca
ggcctcttcaatgcatttacagccactccctgccccttcagctcactcttgcagctttcttctttttctcattgtgacgctttttcgtttttctcttcctccgtctttcccaaacgtgtcttttgctcgcagcaaatgcttgaggcagaagaataagcgcaggcccttaaaaataa
gtgtcggtgctcagcgccggaaacaacaagcacaaatcaagcactggcgcacacacgtgctgtgagttgtgcacaccttaacacactgttacctgtatgtgttgcctgtctagcattgaaagtgacgggagcacacacgtcctgtgtgttgttcacaccttaacac
actgtgttgcctgtctggcattgaaagtgaagggggcacccacgttctgtgagttgtcacatcttaacaacactgtgttgcctgtctcattgaaagtgatgggagcacacacctcctgtgagttctcacgcgttaacaggctgatctacgttcctggcattccagagga
agaagcctcaatcttcgccccctcacacagacatgacatcctttctagagtttcagacaccgaggtgcggcttgtcagctcttctcaccctgacaatcttcattaaccaaccagcggtcagtaaggaggctcgtgcctgctgctcaaactctgagacttggaggtag
aggtgaaccaagcatctctgcccaggcaggactcagcgctggcggcgccctgtgcgacaatcccctcccgccctcccctaccacccaggacctcctcctcggactccctcactaccacctgccccccacccatccagagcccctctctcacacacacgtcatt
tgttttaaagcttcggtaaaggctgactttactgatccactcagctaacacataaaatacagatctgttctgtgcaacaggcacattaaccctctgtgctactttatggctaaaaagtaccactggagaaaactccatctctctctagatctgggacattattgacgaaagt
tatctcgacatttttattgctgcctgaaagctggacacaaaagaactctgcagcaggtgctttaaaatcgaaagttattgctcaagacagcgcccccccatgaggtcagtgccccccctccaggtcagcccctagtgcgactgcaccggtcgcaccctctaaagcctg
ccctgtatccgccgtttgctctcagtgagctttaactgccaggttatccttgccagaggcggtgatctgggacacttccttccacgaaaagcaatttactccagccattgacaggagtaaaccttcgagccttctcagcttggacatgggcagaaaagaatcagtgg
gcgctcctggacagtaatgtccgagctgtcaccaaacttccatgccagccctccacgccttcacccccccaatactcacctcctctgctgattcacagcctgtcataggagtgactgcggggtgatattacagcctgcgggtgtcctgcagcccacagaggcagg
agtttacgattctagaagtggtttcacgagtaaacagtgtattcataaatccaaccccttcccatggatgtgtatatggtttgtgtacattgcacacgtaatcttgctattttgatatttcacgttgtcacgcctgctgttttcggtcagacattacgcatttttagtacattttctgt
attattgtttttggctatcataatgaagtaatgaaaatattattgtgtaatcattcgtagtgtaaataaaataaaaccgccataatgacgaattaggttgtgaaaaccggcagaacgttgttcttagtacagagaacgaaaaaggcatcaaaacatttcattagaaacatag
aacaagaagaaaccaaccgcggctattcagaaaggtaactcgcactgcaaagttactccacagtgtagacctgatacaaggcgcacagccaaggttacagcgcgagttaccaatgccacaaatgaaagcgcgcagcgccacacacaggccaccactggtc
ctgcaacagccccatacagaagaagggactgaaactaaaacaccgtaggaatttacctacaaatatttaacaccctttgaaataccagtaaatatacttgaatgtcaaacaatttgaaaataaaacatatactatttatttataacgtatttaactaccttttttactttaaaa
gttcaggtaacacatttcataaaaatattcaagctcattacaaggggcccgtaaacagggtgatgtttattgcgcttgataaaatctgaccccatagagtaagagtttatcaagtgcgataaatatcacctgattacacatttttgaggaataatgtggattttggtgcttat
cacatcgaatctgcatgtgctggtcattaccgaacagtcctctctcccaataaatgttgtcgggtttcacctgtcggagtttttcagggtaaaactctgtggctgtgatagttatcagacgactcataattcggctcagcacggggatcaggatttaaccccgctgtccg
cgtgtgagggggactccatagaatagatgagaggtgatggtggggatttacattcgtaggtttgtgaatagcattttggaatactttagcagtagtactgtagtaatgataagtacacacctaaatgtcatttgtggatagggcccaatgtgttcaccatcataagtact
gcacaaacaaatcaagaaacattatcaaagcattatttacatttttgtatatgaataaagttaaaaagtaactataacatgtaatttgaaacggaatgaaaaaaacagtagcagaagaatgtgattagcactccatgcttgagagaaagaaaacatctaaaatagagcc
tcacgtgcagagcacatcaaagcccagggaacccacgtccgagaagcgctgcctgtcctctcactctgagaattaggtcctgatgttggcatgaacgctcccattgattcctcgcgcatgctcagtcgagaaggacgcactgtctaccggcagcagagctgcat
caaatgtctgcactttaggtgcctctcagagacaccccacgtcactggggattgtctcatgtttacagtaagaacttgtgggtgacacccaagcccacacgtcgcgccttcctctgggggtgtgatacactcctgtccgagtctcagtgggctccgacagtccaggt
gtgagggggcacagaccaggaacaccagctgaggaagagtgggcggggaatcgttgggtgctgggcaccccagtaaagccaaactagacctcatggtgtaggcccttcaaagagatgttctaaacattcaggtacatggacgtcatgttacgtgtgactaca
cattgtggtactagacacttacatcagacgtgaccagatagtggggcactggcctcttgagtcaggtgtgaacagatagttgggtgctgggcccttcagtcaggtgcgaccatattttggggtgctggcctcttcagtcagctgtgacagatagtggggtgctggg
cccttcagtcaggtgcgaccgatagtggggtgctgggcacttcggtgaggtgtgaacagatagtggggtgctgggcactcagtcgggtgagaacagatagtggggtgctgggcagtcagtctggtgtgaacagacagtgggatgctaggctattcagtcagg
tgtgacagatagtggggtactgggcacttcggtgaggcgtgaacagatagtggggtgctgggcactcagtcgggtgtgaacagatagcggggtgccgggcactgagtcaggtgtgaacagatattggggtgctgggcacttcggtgaagtgtgagcagata
gcggggtgccgggcactttggtgaggcgtgaacagatagtggggtgctgggcactcagtcgggtgtgtatagatagcggggtgccgggcactgagtcaggtgtgacagttagtagggtgctggccccttcagtcaggtgtgagcagatagtggggtcctgg
catcttcagtcagctgtgacagatagtggggtactgggcccttcagtcaagtgcgacagatagtggggtgctgggcccttcagtcaggttccaccatatattggggtcctggcatcttcagtcagctgtgaaagatagtggggtgctgggcccttcagtcaagtgc

gacagatagtggggtgctgggcacccagtcgggcgtgaacagatagtggggtgcagggcactcagtccggtgtgaacagatagcggggtgctgggcactgagtcaggtgtgaacagatagtggggtgctgggcacccagtcgagtgtgaacagatagtg
gggtgctgggcactcagtcaggtgcgacagatagtggggtgctgggcacttcggtgaggtgtgaacagatagtggcatgctgggcactcagtcgggtgtgaacagatagtggggtgctgggcacttccgtgaggtgtgaacagatagtggggtgcttggcac
tcagtcgggtgagaacagatagtggggtgctgggcagtcagtcgggtgtgaacagatagtggggtgctgggcccttcagtcaggtgcaacagatagtggggttctgggcacttcggtgaggcgtgaacagatagtggggtgcagggcactcagtccggtgt
gaacagattgcggggtgccggccactgagacaggtgtgaacagttagtggggtgctgggcacttcggtgaggtgtgagcggatagcggggtgccaggcactttggtgaggcgagaacagatagtggggtaccgggcactttggtgaggagtgaacagata
gtggggtgctgggcacccagtcgggtgtgaacagatagcggggtgctgggcactcagtcgggtgtgactagatagcggggtgctgggcacccagtcgggtgagaacagatagtggggtgccgggcactgagtcaggtgtgaacagatagtggggtgctg
ggcacttcggtgaggtttgtgaagatagcggggtgccgggcactgagccaggtgtgaacagatagtggggtgctgggcacttcggtgaggtgtgagcagatagcggggtgccggacacttcggtgagacgtgaacagatagtagggtgctgggcactcag
tcaggtgcgacagatagtggggtgcagggcacttcgtgaggtgtgaacagatagtggggtgctgggcactcagtcgggtgagaacagatagtggggtgctgggcagtcagtcgggtgtgaatagatagcggggtgccgggcactgagtcaggtgtgagca
gatagtggggtgctggccccttcagtcaggtgtgagcagatagtggggtgctggccccttcagtcaggtgtagacagatagtggggtgctgggcccttcagtcaggttccaccatatattggggtcctggcatcttcagtcagctgtgaaagatagtggggtgct
gggcccttcagtcaagtgcgacagatagtggggtgcatggcacccagtcgggtgtgaacagatagcggggtgctgggcactgagtcaggtgtgaacagatagtgcggtgctgggcacccagtcgggtgtgaacagatagtggggtgctgggcactcagtc
aggtgcgacagatagtggggtgctgggcactttggtgaggtgtgaacagatagtggcgtgctgggcactcagtcgggtgtgaacagatagtggggtgctgggcacttcggtgaggtgtgaacagataatggggtgcttggcactcagtcgggtgagaacaga
tagtggggtgctgggcagtcagtcgggtgtgaacagatagtggggtgctgggcccttcagtcgggtgcaacagatagtggggtactgggcacttcggtgatgcgtgaacagttagtggggtgctgggcactcagtcgggtgtgaacagatagtggagtgctg
ggcacttcggtgaggtgtgagcggatagcggggtgctgggcactcagtcgggtgagaacagatagtggggtgctgggcacttcggtgaggtgtgagcgcatagcggggtgccgggcactcagtcgggtgagaacagatagtggggtgctgggcacttcg
gtgaggcgtgaacagatagcggggtgccgggcactcagtcgggtgtgaacagatagtggggtgctgggcactcagtcgggtgtgaacagatagtggggtgctgggcactcagtagggtgtgaacagatagtggggtgctgggcactcagtagggtgtgaa
cagatagtggggtgctgggcacttcggtgaggcgtgagcagatagcggggtgccgggcactgagtcagttgtgaacagatagtggggtgctgggcacttcggtgaggtgtgagtggatagcggggtgccggcacttcggtgaggcgagaacagatagtgg
ggtgctgcgcacccagtcgggtgtgaacagatagcggggtgctgggcactcagtcgggtgtgaacatatagtggggtgctaggcactcagtcgggtgtgactggatagtggggtgctgggcacccagtcgggtgagaacagatagtggggtgccgggcac
tgagccaggtgtgaacagatagtgaggtgctgggcacttcggtgaggtgtgagcagatagcggggtgccggacacttcggtgaggcgtgaacagatagggggggtgctgggcactcagttgggtgtgaacagatagcggggtgccgggcactcagtcggg
tgtgaacagatagttgggtgctgagcccttcagttaggagcgaccatattttggggtgctggcctcttcagtcagctgtgacagatagtggggtgctgggcccctttagtcaggtgcgacagatagtggggtgctgggcacttcggtgaggtgtgtgtggatagcg
gggtgccgggcacttcggcaaggcgtgaagagatagtggggtgctgggcactgtgaacagatagtggggtgctgggcactcagtcgggtgtgaacagatagtggggtgctgggcacccagtcgggtgtgaacagatagcggggtgctaggcactcagtc
gggtgtgaacagatagtggggtgctggacactcagtcgggtgtgactagatagtggggtgctgggcacccagtcgggtgagaacagatagtggggtgctgggcactcagtcgggtgtgaacagatagcggggtgccgggcacgcccctcgggcacgtttc
tcacctctggacaaggtgggctcctgcaataaatgcatacaaccccccagagagcagatgttgaacttgccgtgtcccccacactcactgaccttcatctgctccacaggaagacgccggcacaacagaccttctcggaataaacacatctcgcaggagtgcc
ggagacagtgccgggggcagtggaagagccgcggctgccagcatgaaccctgcaggatggaggtggcgatgcccttgtcataacacctgagagaatgtgggaccacgacaggtgcccctctggcacccccatgatacttaggccctggtttgtcaaactcc
aaacctgaccccgagtctctttttcatgttcagctctcacctacagacatggtcagcattcgtgaaactccaggggtaacttccatcaactgacacaaaatatgtgcagtccaggggtggactgagtgacacagtggtgagctgggcagtccttgtgctgcagacg
cagggtttgtgctgtgtggttatctgcaccgggccacccctgagtacttatgtaacgcattagtctctgacagtgcttaatttgtaaataaaaacgtgccggtgcccaaagccctccccttaaataggtggctgctgaatttaaatgtgggaacacagaatactgagcc
atctcgggcctctttaatccatttagagccactccctgccccttcggctcactcttgcagctttctgcttactatctttgtgacggtttttcatttttctcttcctccgtctttaccatgtgtcttctttctgctcgcagcaaatacttgaggcagaagaataagcgcaggccctta
aaaataagtgccagtgctcagcaccgccaacaacaaacacaaattaagcagtggtctctgatcttgccaagttcagtggtgcgagagatgtttagacagaggctgcgaatccatacgaccattccttgtttatacttaaagccagtccaataaatggacacttcatat
ctccaaacttctgcccatctgcatgcgtaaccatgtcgatgtcccttaggtctctcctaagacagcacatgctctgcagcttgtaacattctgtttactagaaggggctttggagcccacctcttgcttaccattttctgtattcactgtcattcttatgaaccatttcctcatt
ggccagagcttgccagtttcatgtcctcttctttcctgtggagcatggaccagctactgattgcttcagtttgattagtgtcctcctgttctgaattaggtacttttttttgttcttcttcctgtcgtgttgcttctcaactccgacctgcccattatgtgtgttgtttctctgtctgtgtt
tctccttcgttcccacactttcgttgtatttgttcttctcgcctacctcatacctcccttttttcatgttacttgtaagcctcttgttcttgttgcctgtgtcgcttgtaagcctcccgcctcctgttggtcttgttccttgaaagttgtccgtgttgtttcttcaccccccccacccctgct
gcttgtattgtggttccacctcaacttgagaaaacaaaattgctttgatgctggccagcacttttgtcaaacttttagtcatgctgtacagccatgaacgattggctttaccaatgcttgttacatttgttctcaaaattgatttccctcattacagattgttaaactggcataca
caggttattatgcaagtaattacattggtgtattacgcgtgccagctttttgggccttctgttattgaaaatgttgtatcttctgtctttgaaaataaagacatttttaccaaaaaactataaatgtgttcttctactttgtattctgttttacaggctgttacccaggtgtggtatta
gttgattttcatgtaaaacttgagagcggtggcactggctacgctagttagacctacttgtgcgacttttgaattagaccaatttttccgtttttatatt

**C:** Sequence of cDNA (coding region in red and highlighted in yellow)

ggcgcagacgcccctgggtcggatagagcgctctaggggctcccattgtgtctgtggtcggtccctgcacaactatcgacctcactaggatgaggctctccctggcactggcggtgctgctagtggcttgtgtatccctctgctgggggcaccgaggcccccca
aaaggacgagaaacgactagtcaagaagcacccctctcaatggaggcagcgccgggcacacactgcccttgaagacaaaagacccaggaggatggaagaacccgtgagctcgaaaggggtgaaggactgtccgtgccacgtcctaaaacgaagacgc
cggcacaacagaccttctcggaataaacacatctcgcaggagtgccggagacagtgccgggggcagtggaagagccgcggctgccagcatgaaccctgcaggatggaggtggcgatgcccttgtcataacacctgagagaatgtgggaccacgacaggt
gcccctctggcacccccatgatacttaggccctggtttgtcaaactccaaacctgaccccgagtctctttttcatgttcagctctcacctacagacatggtcagcattcgtgaaactccaggggtaacttccatcaactgacacaaaatatgtgcagtccaggggtgg
actgagtgacacagtggtgagctgggcagtccttgtgctgcagacgcagggtttgtgctgtgtggttatctgcaccgggccacccctgagtacttatgtaacgcattagtctctgacagtgcttaatttgtaaataaaaacgtgccggtgcccaaagccctcccctta
aataggtggctgctgaatttaaatgtgggaacacagaatactgagccatctcgggcctctttaatccatttagagccactccctgccccttcggctcactcttgcagctttctgcttactatctttgtgacggtttttcatttttctcttcctccgtctttaccatgtgtcttctttc
tgctcgcagcaaatacttgaggcagaagaataagcgcaggcccttaaaaataagtgccagtgctcagcaccgccaacaacaaacacaaattaagcagtggtctctgatcttgccaagttcagtggtgcgagagatgtttagacagaggctgcgaatccatacga
ccattccttgtttatacttaaagccagtccaataaatggacacttcatatctccaaacttctgcccatctgcatgcgtaaccatgtcgatgtcccttaggtctctcctaagacagcacatgctctgcagcttgtaacattctgtttactagaaggggctttggagcccacct
cttgcttaccattttctgtattcactgtcattcttatgaaccatttcctcattggccagagcttgccagtttcatgtcctcttctttcctgtggagcatggaccagctactgattgcttcagtttgattagtgtcctcctgttctgaattaggtacttttttttgttcttcttcctgtcgtgtt
gcttctcaactccgacctgcccattatgtgtgttgtttctctgtctgtgtttctccttcgttcccacactttcgttgtatttgttcttctcgcctacctcatacctcccttttttcatgttacttgtaagcctcttgttcttgttgcctgtgtcgcttgtaagcctcccgcctcctgttggt
cttgttccttgaaagttgtccgtgttgtttcttcaccccccccacccctgctgcttgtattgtggttccacctcaacttgagaaaacaaaattgctttgatgctggccagcacttttgtcaaacttttagtcatgctgtacagccatgaacgattggctttaccaatgcttgttac
atttgttctcaaaattgatttccctcattacagattgttaaactggcatacacaggttattatgcaagtaattacattggtgtattacgcgtgccagctttttgggccttctgttattgaaaatgttgtatcttctgtctttgaaaataaagacatttttaccaaaaaactataaatg

**D:** Amino acid sequence (signal peptide shaded)

MRLSLALAVLLVACVSLCWGHRGPQKDEKRLVKKHPSQWRQRRAHTALEDKRPRRMEEPVSSKGVKDCPCHVLKRRRRHNRPSRNKHISQECRRQCRGQ
WKSRGCQHEPCRMEVAMPLS

**Fig. S15.** Gene position and architecture (**A**), genomic DNA sequence (**B**), cDNA sequence (**C**), and encoded protein sequence (**D**) of the unannotated *cxcl17* gene in the NCBI reference genome of *Pleurodeles waltl* (Iberian ribbed newt). The information was downloaded from the NCBI gene database (https://www.ncbi.nlm.nih.gov/gene).

### A: Gene position and architecture

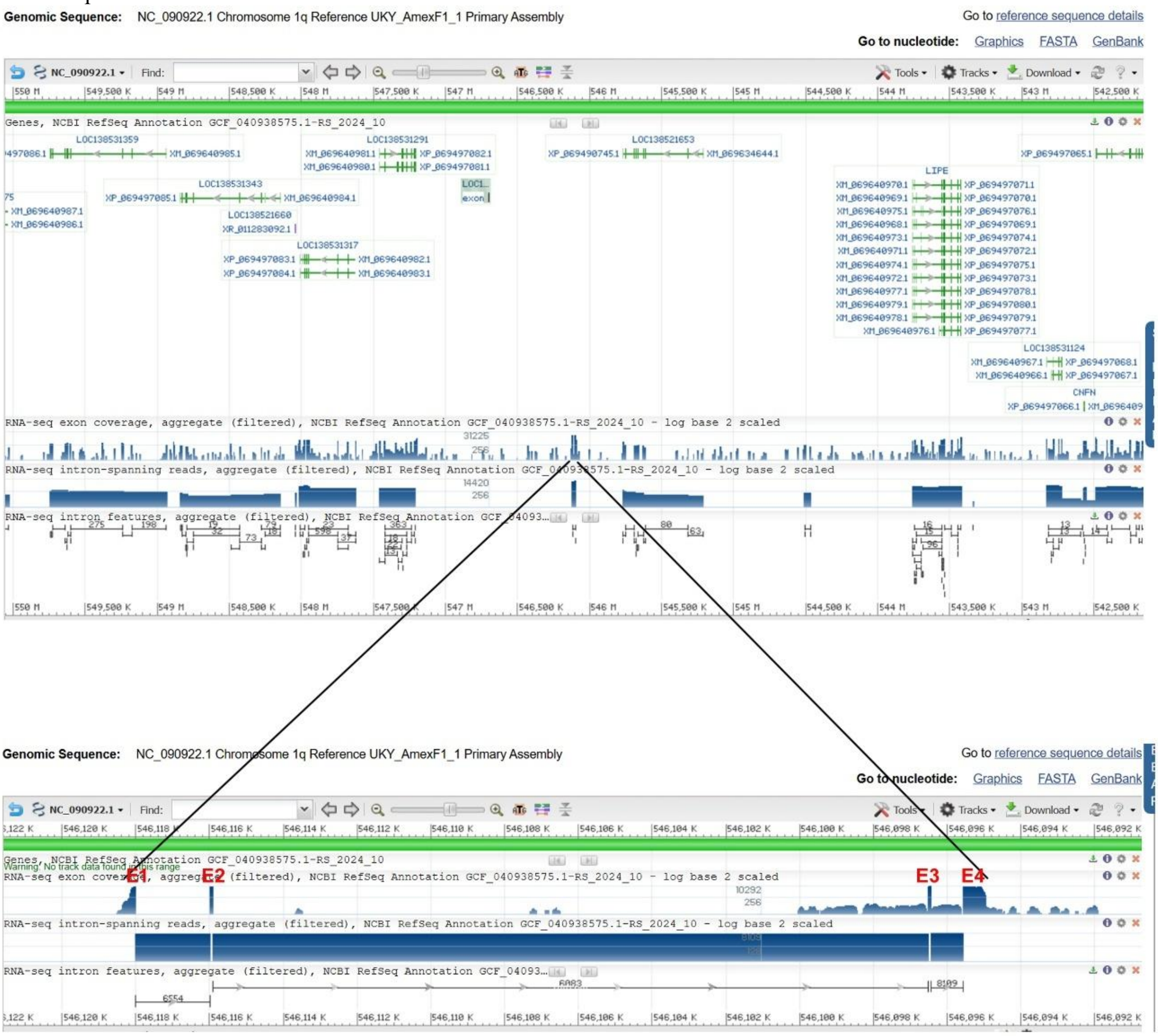


### B: Sequence of genomic DNA (exons in red and highlighted in yellow, coding region underlined)

cttctgataggatcttcgccccataagggagggcctctagtgctcggggctgtcccagtagccagcaggggggtgttgggcggggtcttgttttctgggtttttcgcagggctttttgtttcgtttacttcaggtttcaaggtgatcggctgagggcttggatcagtagtg
ctgtgccttttcacttggccctacaccttctcagcgagcggctgcaatgaggaccaccctggcgatggcactggtgcttctgctctgcctctctcactgcacgggccatagaggtaagaactcgtgcatgctttacttcatacgtagagcgttgtacgtcgtgcgtgc
attcagtttcgccgcgctgtttctgactaaaaggccctaatttaggcggcgtcgctcccttgggaatcctgatctataacagcctttcaataagtgcaaagcctcccagaggctttcatggcaccacccgacaaaaaatgcccttctgcaaaaacgtcacactggca
gtgcactccccacccccctaaagactaaacacgctgagaggacgtacttcttaatgtataacaaggcgtgcatcttacgttctgcctcatatggtgtcagaacgtacggcgcttgactgggcacactgacgtacaagcacttctaggcacatatatcagatcgttaatt
ttaaactgttctaaaggttcctgcgttctgatcaaggaagatacgccgtatcctctcagtatttcccgctggagggccgggaaaagtaaatgtacagtatgtttctcttcaaacacgtgcaatgagcacaggggggcttgccagtacgtgtaaatcaccgagatatta
ggattgaaatagccgggctttcaggagaagctttggtaaaccattgaaaatatgctattacgattcactggacccaaatgattaagagtatgaggcctcctcctggttgttgtcatgtagccttcaacagacgagacgcagtccatccagtgactgcatctcgtcttttc
aaggctgtgattgcattaccaggaaacgtaaatatgcaattaaaccaacagatgtggagcagttgggaccttagagcctcatttgaaaaatgttaaaagcccccgacgccgtccgtcagcgcacccctacaaccaatgcagggagctcaagttgtctcattgcaa
ggtagattaccacagtgcgtttgcatttccaccaaacaaaaaaaaaacgtatgtgcttactgcgaccatttgcaatttctcatgtggcaagctttacacattaccccgtggctcagtggtggggaaatgggggacagtgccagcctcgggtccgccagcgccctgt
gtgaattatccgccgaacaacaaaaccgagccccctgctgttcctgtgaagtgtttttaagggcctcgcccagcgcagcgtgattctttaacctttgataggcgctgggccagaccttgcctccagctcagtgtcatgtacacttaattacatgacacgcagatgga
ggcaaggcgccctttatttgccgcccgtgccatattatgcctgtgtgcctagagatgagctttcacttcagagataatcccgcgtatgcaaattcgcaccacttgactccagaaaaaagcaccttgttccccgaaaaggtggtttaaaggtgtgtccagcaggtgttt
cttgtggctcggtgtggtgaagataggatggctgcattttcccgaatcaaaatcccgggacaattttacaaagttaatcatagagaaaggactacaggtgcacattggccccgtctattcttccctctctcagatcccagctgccattgcgtgaactcctgctccttcct
gtccctctcataaaatgccgtgggctttattttcccccatgacttgtccacgcggtgtttcatatctcagttgctgtaatcacttcttggtgcgttcaaagcttttgtctttccagcaccaataagagcggcttatattgattgccccctctccctccctccccccacagaagg
ctttccctttgctggcagctggcccacattcaccttttttttttaaatcagagaagtgctctaataaggatctcgttttctaaaactggggtctttgactgtttagcataagcttgaaaaactggtactacacccatatcccgccaaaaaaaattaacctatgtgaagagg
agcagactttccaacgttatatgcgtggatgctaagagaatgagtttcaggatgtactccgatgtggggctttttgcaggttattttttacactaacaaaaccccttttctcattcccaggttccctcaggcatgagaagggcctgaccgagcaccagtccttacccggg
cgacatcaccatggacacggtaagcagcatgtcccctggcctgcagagcacccaaagcaaaatactatttactggctatctgcggggcagcacatgctggtctagcttgattgcgggctctgctccagctgtgcggtgctctgccctccactgctgtccgcactg
ggtcctgtgtgcctcgtgatttgtgctgtgcgccctatgacccgcatgatagaggagtagggtttccgctgtacctgctggagacatatgtataggcggtcagggctttaagtatggcagggcttcccacagcgcagccctggcagtctcttttttttttaattcagtta
gttaaatataatcaaccttacaataacaaaatatctacatcataaataatttcataataaaatcttaattgcataaaaatcaatcaacattaggtacgtgttaggaaagatttcgtaattgtaggtaattttccttccatgtggcaaccccctaaactttttgctgttgttttgcacct
tgttcttaactttgccagagaagtgcagccttctgctgcactcctctgggctttccctccttcttacggggaaactgacatgcccttaagacagtcagtaccggggaacgctgtttcccgtagtttcttttatcgtgttgcggctttcgtgcaacatgacacaagagggca
ctgtaagctaagttacagctctcttgggtctcttgccttctacaggccctagatactcttttaggatggtatttgatcggtgcaagtcattcagggactcgaatggtggcagcgatcgttttgtttattttgagcttcttttggcgttctgagcacgtttttcgcactgaaccca
aaatggtggcctggctcccaaaatggccagaccacgaggctcctttgtcttgtccactgaccggcttgctcccttcaccttccccaggtcgagcgacccggggccttccttatttgggcatgtactttcccgcgaaacctggttgctttcgcgggcttctacatgtcgt
ttgtgtgcctccagggtgtgggctgggatgtctggtcccaatacacatcctgggtgccagagagtctagggtagtgtgaatgcctgggaccaccgtggtccgtgattggtccacgtatggtgtgtgtgttgatgagcggatctcccattggctcccctccgttttgg
cgcgaagggaaccgtggataagaggggggctggaaacaccactaccatgtcgatctcctgagttctcacgaacaaaggaatacaggagccaagtatcctgcaaccacgaggctggaccgaggaatcgctggtgacccgctgaaggaatcctcctttgcactg
ctgtcgtcctgtgaaaccctttcctcatttgatcgaaccagaaggcctgtactggtgcttcgacctttggaatagcaggaacagctattcccggcgacttcatcatctcctgccaagtaagccttcggaagtgtctctgggccatccagttgactgaccagcttacaag
ggagtcctgctctgaaaccgccgccaggttgtgttgtatttcgagtcctggtcactctcccacgcttcgtacacttggcgttcccaagtctgaagggaggagtgtacatttggctcggatacacacggccgctccgttgtcctgctgcggcaacaaactgtgctggtt
tcgtcgattgggacgtctttttgctctacacgactacgaggcgttccctgctcttctgaacctgccagctgctgaaactaacgtggagaagcctgggtctgcttgcttcggggaccggtgagtagtttagcaaggatctggtcgccctcctgcagtagcccattgcttt
tccccgctgtgtaaaaacagcacccttttgtccgtactcgcgctctgcttctgctcgctgtggaggctgcgcttgagttgcggaacctgttctccttctgctgggcactaacccgaactgcctgacttcctccgaagagcttcccctttgccagttggcactggcgtg

gcaggtactttgagcacaaggctccactcttcgcagggctcggacttccatctagttaggtgcctctgaaggcgaactgagtctaaagcgttatcgcactgcctgttttccttcccttgcaggtggactacacatttcgtgcttataacgctagcgactgttgtatttgta
tatatattgtgttgatatggtatgtgttggtttaatattgtgatattgtgattttctcgctacggtcattgttttaaaggccttaaaataaatgtgtatctatttttgatataacaactgtttggagtagtttgtagcgctcttgctttgtgtgctcattgcgggctttcagtcaccctca
cccctcttgagacctagtcttgaccgccgcgatcgctaccttgattggtctggctcgtccagaggttaccctgttccaagtaattaggttggccttctgaacttctgcgccaccaggacagagttcagaaatccgtcttaacaaattggtgtacagcggtgggatacg
actgaaggtatccagtgttgcgctcgaactaaggcatttcaggttattccaaccagtacacgtagtgaattgttgtagtataactttaaaaaggccatagtcctatctaacagtatggaaacaaataccttgttaactttaaatgcttccaaatcagaaacattgaagaag
atgtgtgcggagagaggtgtttcctctcaaggcacgaaattggaaatgatccaaagaatcgtggcgtggcaagaagtgcagcttgacgctataacttctgaggacagtgcagaaggtggtggggctgctctcactggtagaaccgacggttctgccgctgcac
cgatccatcgcgaccacgaggacagggaggatgtgatgtctaatgtttcctctgccagtgctgatggggaggctatgcttgaattcaagaagcaagagctgcggtttaggcagcaggagcttgaattcagcatagccaaactaaaaatcgaacaggcagaaaa
acagaaagacagggatcaccagttggaactggccaagcttcaggttgcaaatgggtccaagggtcctggacatgggtctggggcctcctctcatccacggtttctgaaagatctgctcagtgtgtatgaagaaaggaatgacattatggactggctgaatgcgtt
tgagtatgcatgcaaggttcaagaagttgacaatgctgataagattgcgtggttgctcagtagactgccccattctggctgtagtgccattatggaaatttcaaaggctgagcgagctgatttcgcaatgttaaagcaaactctcatttttaagtttgggaagactccgg
cccaatatttaaagcgttttcgagagtataagaaaaaggaaggtggtacttgggtatcttatgtcgctgaatctttgaaaaacttaactggctgggtcgaaggttataaggtttctaccttcgaaggcatgataaatcttgtcatgctggaacatatttttgcttcttcttccc
cggagcttaagcagcatttggccgattcgaaggagatagatcctaagaagctggccattgcagcagacttatgggtgtcacatcgagtgtcccataaggacaaaactcatcctggtaagcaggaggaagggaaaaagtccaaagatatggagggtggagaat
ctgctaaatcttcttcccataagggccacaaaaagaaaaacaagggtaagacacaagagaattcacagggtagtggaggttcgcctccgaaacagaactctggttacaataagcctccctatgtaaacagacccactggggcttgtttttcttgtggagcaactga
ccatgtgaagggggatccccggtgtaaggttagacctttgggggtccactccgctaatctggccttctcctcctacgaggaagaagagattacaggagcatccttctcctcagactcatttcccagcgggatagaggctgaggtagttttagtgtccctgggttcct
gggagcctaaagtcgcagaggcttatgctaggattccggataataccaagcactactttaaagacagtgtgctgatcaatggagtggatactccagctcttaggaatagtgggtgcagtcgcactgtagtggattctaccttggtagactcagaggaagttgccag
agctactctgatgcccactaagttggctgacggccccctccgtatgtttcccgtaatacctgtaaatcttactttaaatggtacagcagtcaggattccagtgagcattcagagaaacgtcccgggaaaggtcttgttaggtaatgatctcagagctttagggctcgta
ggagacactgccgaaagaccctacacacggtctcaagcacgggcagcattggccaagtccaatacgccgccccagccaaaggggaaacccaaggcgaaaggagtctacaagtcaaaacgagggaaagagaatattcccccgagctcgcctaccactgc
ggtggaggaagttggtcaatcgcccgggcctgacgaggaggtggcgatgcctgaagtattcgaccccaacgatcatcctgaactaaaagacttgttagttgaaggaggtcctgatagggacaaatttagtaaaggccaacgtgagtgcccatctctggaaggc
ctaaggagacaggcctgctcccagcttgagggtgaacctcctgggatgcataggatttattgggaaggtggactcctgtatagtgagcccacagagtctacagaaggagaccataggaggttggtggttccccttgctgttaggcccttcctcctgcagttggcg
catgaagttccccttgctggtcaccttggtatgaggaggaccaagcaaaggttctccatgcacttctactggcccaagatgggggagcaagtagaaagtcactgcaggagttgtgccgcctgccagttgtctggtaaggttggcgcgacaatccaagctccgttg
gtaccgctcccagttgtgggggtaccatttgaaagggtatggatcgacatggttggtccccttgacccgcaaacatcctcgggcaataggtttatcctggtgctggtcgaccatgcaaccaggtatcctgaggccatcccatggaaaactgtaactgccaagtcag
ttgcagaagctttactcggcatatttactagggttggcttccctaaagaagttgtgtcggatcatggcagcaactttatgtcgaagtataggcaggctatgtggaaagaatgtggggtcacctacgggttctcttctgcataccacccccagaccaatggtctggggg
ggcgtttcaacaggacgttaaagagcgtgattggggctctggaggagcctcagagaagaaaatgggatcttcttgtttcatgcttactgtttgcctaccgagaggtaccacaggagggtgttggcttcagccccttcgaacttctgttcggtcatcccgtgcgggga
cccttagccatcgtcaaagagggatgggagatgcccactccccctgttactaccaatgtagttgattatgtgctatggctcagaaaacggatggtacttctgatgggcttggctagtgataaattgaaggcagggcaagcgctggtgaaacattggtacgaccaga
aagcttccctcatcaactttacacttggtcagcaagtcctgatcatgaatcccataaggcctcgagctttacaggacaaatggtcaggcccttacccagtggtggagcccctgggtgaggttagctattcagtggaccttggaaggcctggacaggctctaaaagt
gtttcacgtgaataggctgaaggctttcggcccaagcgtcgaaatcgcagcactgctactggcttcagacgttgcgggagaggagtctgaacctctccccgacctgttcccaagcagtccttcagatagctcttttgagggtgcccgagtggcccctcaccaaaa
tggacaaggctcagtggtttaccttggacatgaagtaggtggagggaggagacagcctctactcagtcaggtacaagcagtacaagactggcctacccccactacgcaaacccaagtgagagcctttttaggactcacggggtattaccgcactttgtggcag
actatggcaccatagctgctcctctgactgccctcacaactccgaagaaacccaagaaggtaacttggaccgacgagtgtgagaaggccttcaatggcttaaaagactccttgtgccaggcccctgtcctcagggtccctgattatcaaagaccttttatcgtgca
gacggatgcctctgacactgggataggagctgtactatcacaagtagatgagaagggtaaggaacaccctatcggttttattagtcgccagcttaaggacagagagctcagatggtccacggtagagaaagaatgtttctctgttgtgtgggccctaaggaagct
gaggcccttcctctatgggacccacttcactgtgcagactgaccataagccgttaaagtggctaatgaacatgaaggggggagaatccaaagttgcttaggtggtccataagtctacaaggcttcgacttcactattgagcacaggccaggtgtccaacatagcaat
gctgatgggttgtccagaatgtttaaccgacctgtcaagagactcatccaggggtggattagttcccccctgtccatagtctgggagggggcgagtgttaggaaagatttcgtaattgtaggtaattttccttccatgtggcaacccctaaactttttgctgttgttttgcacc
ttgttcttgactttgccagagaagtgcagccttctgctgcactcctctgggctttcgctccttcttacggggaaactgacatgcccttaagacagtcagtaccggggaacgctgtttcccgtagtttcttttatcgtgttgcggctttcgtgcaacatgacacaagagggc
actgtaagctaagttacagctctcttgggtctctagccttctacaggccctagatactcttttaggatggtatttgatcggtgcaagtcattcagggactcgaatggtggcagcgatcgttttgtttattttgagctccttttggcgttctgagcacgtttttcgcactgaacc
caaaatggtggcctggctcccaaaatggccagaccacgaggctcctttgtcttgtccactgaccggcttgctcccttcaccttccccaggtcgagcgacccggggccttccttatttgggcatgtactttcccgcgaaacctggttgctttcgcgggcttctacatgt
cgtttgtgtgcctccagggtgtgggctgggatgtctggtcccaatacacatcctgggtgccagagagtctagggtagtgtgaatgcctgggaccaccgtggtccgtgattggtccacgtatggtgtgtgtgttgatgagcggatctcccattggctcccctccgtttt
ggcgcgaagggaaccgtggataagagggggctggaaacaccactaccatgtcgatctcctgagttctcacgaacgaaggaatacaggagccaagtatcctgcaaccacgaggctggaccgaggaatcgctggtgacccgctgaaggaatcctcctttgca
ctgctgtcgtcctgtgaaacccttttcctcctttgatcgaaccagaaggcctgtactggtgcttagacctttggaatagcaggaacagctattcccggcgacttcatcatctcctgccaagtaagccttcggaagtgtctctgggccatccagttgactgaccagcttac
aagggagtcctgctctgtaaccgccgccaggttgtgttgtatttcgagtcctggtcactctcccacgcttcgtacacttggcgttcccagtctgaagggaggagtgtacatttggctcggatacacacggccgctccgttgtcctgctgcggcaacaaactgtgctg
gtttcgtcgattgggacgtctttttgctctacacgactacgaggcgttccctgctcttctgaacctgccagctgctgaaactaacgtggagaagcctgggtctgcttgcttcggggaccggtgagtagtttagctaggatctggtcgccctcctgcagtagcccattgc
ttttccccgctgtgtaaaaacagcacccttttgtccgtactcgcgctctgcttctgctcgctgtggaggctgcgcttgagttgcggaacctgttctccttctgctgggcactaacccgaactgcctgacttcctccgaagagcttcccctttgccagttggcaccggcg
tggcaggtactttgagcacaaggctccactcttcgcagggctcggacttccatctagttaggtgcctctgaaggcaaactgagtctaaagcgttatcgcactgcctgttttccttcccttgcaggtggactacacatttcgtgcttataacgctagcgactgttgtatttg
tatatatattgtgttgatatggtatgtgttggtttaatattgtgatattgtgattttctcgctacggtcattgttttaaaggccttaaaataaatgtgtatctatttttgatataacaactgtttggagtagtttgtagcgctcttgctttgtgtgctcattgcgggctttcagtcaccctc
acccctcttgagacctagtcttgaccgccgcgatcgctaccttgattggtctggctcgtccagaggttaccctgttccaagtaattaggttggccttctgaacttctgcgccaccaggacagagttcagaaatccgtcttaacaaattggtgtacagcggtgggatac
gactgaaggtatccgtcttaacagtacgtgtgatataagttagatgtccctttcttaggaatggaatcacttaaggaagagaagaaggagaagaataaaggagcaaacataacaaataactaatatcaacatatttgtaactcatgatctttacatatatactcatgattc
atagttcattcctggtacattcatagttcacgatttgttaataattgtttcaacacatagtttacagtcagtaatgatgaacttatttacaatcaatcgatcaggacatgatgatgaaacgttttgagaaatgcactatacagtctccaaatatttccctccagcccgggctctg
tcagggctctccgaactcacttccctcagcacatttcttgaaaaatctgaatggcaatactgctatttgccactagatggcccttccgtaatgtttagaatcacacgatttaaagatttgtgaaatcaggtacttttcaaataagaaaggggaattataagagaatgcatg
gtttgatttttttttctccacacatcgtagaacagtattatcattaaagtgggttaacaaaggtagctggcgcccctctctcgacagaacaaggcctgataacctgttcagtggttattaaccacttgagagttcgcagccactgcttttactcactgtgcctttcacagatg
ggaaatatccatttgcatatcagtctttgtcctgcccaggggcagtccagcaccgaaatacaatggcaattccaatctgaacgagagtaccaccagcaaccccatacacgtgtttcacccttgttggggatcatcagtgaggtgaagctgatgcctctctaagcaca
gtgagagggagaccacgtctggttgcccccaggagacctagggttaccaatcccaggttccttgaaatttagtttacaaaggtagcgggcgcccctctctcgacagaacaaggcctgataacctgttcagtggttattaaccacttgagagttcgcagccactgct
tttactcactgtgcctttcacagatgggaaatatccatttgcatatcagtctttgtcctgcccaggggcagtccagcaccgaaatacaatggcaattccaatctgaacgagagtaccaccagcaaccccatacacgtgtttcacccttgttggggatcatcagtgaggt
gaagctgatgcctctctaagcacagtgagagggagaccacgtctggttgcccccaggagacctagggttaccaatcccaggttccttgaaatttagtttacaaaggtagcgggcgcccctctctcgacagaacaaggcctgataacctgttcagtggttattaacc
acttgagagttcgcagccactgcttttactcactgtgcctttcacagatgggaaatatccatttgcatatcagtctttgtcctgcccaggggcagtccagcaccgaaatacaatggcaattccaatctgaacgagagtaccaccagcaaccccatacacgtgtttcac
ccttgttggggatcatcagtgaggtgaagctgatgcctctctaagcacagtgagagggagaccacgtctggttgcccccaggagacctagggttaccaatcccaggttccttgaaatttagtttacaaaggtagcgggcgcccctctctcgacagaacaaggcct
gataacctgttcagtggttattaaccacttgagagttcgcagccactgcttttactcactgtggctttcacagatgggaaatatccatttgcatatcagtctttgtcctgcccaggggcagtccagcaccgaaatacaatggcaattccaatctgaacgagagtaccac
cagcaaccccatacacgtgtttcacccttgttggggatcatcagtgaggtgaagctgatgcctctctaagcacagtgagagggagaccacgtctggttgcccccaggagacctagggttaccaatcccaggttccttgaaatttagtttacaaaggtagcgggcg
cccctctctcgacagaacaaggcctgataacctgttcagtggttattaaccacttgagagttcgcagccactgcttttactcactgtgcctttcacagatgggaaatatccatttgcatatcagtctttgtcctgcccaggggcagtccagcaccgaaatacaatggca
attccaatctgaacgagagtaccaccagcaaccccatacacgtgtttcacccttgttggggatcatcagtgaggtgaagctgatgcctctctaagcacagtgagagggagaccacgtctggttgcccccaggagacctagggttaccaatcccaggttccttgaa
atttagtttacaaaggtagcgggcgcccctctctcgacagaacaaggcctgataacctgttcagtggttattaaccacttgagagttcgcagccactgcttttactcactgtgcctttcacagatgggaaatatccatttgcatatcagtctttgtcctgcccaggggca
gtccagcaccgaaatacaatggcaattccaatctgaacgagagtaccaccagcaaccccatacacgtgtttcacccttgttggggatcatcagtgaggtgaagctgatgcctctctaagcacagtgagagggagaccacgtctggttgcccccaggagaccta
gggttaccaatcccaggttccttgaaatttagtttacaaaggtagcgggcgcccctctctcgacagaacaaggcctgataacctgttcagtggttattaaccacttgagagttcgcagccactgcttttactcactgtgcctttcacagatgggaaatatccatttgcat
atcagtctttgtcctgcccaggggcagtccagcaccgaaatacaatggcaattccaatctgaacgagagtaccaccagcaaccccatacacgtgtttcacccttgttggggatcatcagtgaggtgaagctgatgcctctctaagcacagtgagagggagacca
cgtctggttgcccccaggagacctagggttaccaatcccaggttccttgaaatttagtttacaaaggtagcgggcgcccctctctcgacagaacaaggcctgataacctgttcagtggttattaaccacttgagagttcgcagccactgcttttactcactgtgccttt
cacagatgggaaatatccatttgcatatcagtctttgtcctgcccaggggcagtccagcaccgaaatacaatggcaattccaatctgaacgagagtaccaccagcaaccccatacacgtgtttcacccttgttggggatcatcagtgaggtgaagctgatgcctctc
taagcacagtgagagggagaccacgtctgtttgcccccaggagacctagggttaccaatcccaggttccttgaaatttagtttacaaaggtagcgggcgcccctctctcgacagaacaaggcctgataacctgttcagtgattattaaccacttgagagttcgcag
ccactgcttttactcactgtgcctttcacagatgggaaatatccatttgcatatcagtctttgtcctgcccaggggcagtccagcaccgaaatacaatggcaattccaatctgaacgagagtaccaccagcaaccccatacacgtgtttcacccttgttggggatcatc
agtgaggtgaagctgatgcctctctaagcacagtgagagggagaccacgtctggttgcccccaggagacctagggttaccaatcccaggttccttgaaatttagtttacaaaggtagcgggcgcccctctctcgacagaacaaggcctgataacctgttcagtg
gttattaaccacttgagagttcgcagccactgcttttactcactgtgcctttcacagatgggaaatatccatttgcatatcagtctttgtcctgcccaggggcagtccagcaccgaaatacaatggcaattccaatctgaacgagagtaccaccagcaaccccataca
cgtgtttcacccttgttggggatcatcagtgaggtgaagctgatgcctctctaagcacagtgagagggagaccacgtctggttgcccccaggagacctagggttaccaatcccaggttccttgaaatttagtttacaaaggtagcgggcgcccctctctcgacaga
acaaggcctgataacctgttcagtggttattaaccacttgagagttcgcagccactgcttttactcactgtgcctttcacagatgggaaatatccatttgcatatcagtctttgtcctgcccaggggcagtccagcaccgaaatacaatggcaattccaatctgaacga
gagtaccaccagcaaccccatacacgtgtttcacccttgttggggatcatcagtgaggtgaagctgatgcctctctaagcacagtgagagggagaccacgtctggttgcccccaggagacctagggttaccaatcccaggttccttgaaatttagtttacaaaggt
agcgggcgcccctctctcgacagaacaaggcctgataacctgttcagtggttattaaccacttgagagttcgcagccactgcttttactcactgtgcctttcacagatgggaaatatccatttgcatatcagtctttgtcctgcccaggggcagtccagcaccgaaat
acaatggcaattccaatctgaacgagagtaccaccagcaaccccatacacgtgtttcacccttgttggggatcatcagtgaggtgaagctgatgcctctctaagcacagtgagagggagaccacgtctggttgcccccaggagacctagggttaccaatcccag
gttccttgaaatttagtttacaaaggtagcgggcgcccctctctcgacagaacaaggcctgataacctgttcagtggttattaaccacttgagagttcgcagccactgcttttactcactgtgcctttcacagatgggaaatatccatttgcatatcagtctttgtcctgcc
caggggcagtccagcaccgaaatacaatggcaattccaatctgaacgagagtaccaccagcaaccccatacacgtgtttcacccttgttggggatcatcagtgaggtgaagctgatgcctctcaactctcaagtggttaataaccactgaacaggttatcatgtct
gttcttgcgagagaggggcgcccgctacctttgtaaaacaatttttaaggaacctgggattggtaaccctaggtctcctgggggcaaccagacctggtctccctctcactgtgcttagagaggcatcagcttcacctcactgatgatccccaacaagggtgaaac
atgtgtatggggttgctggtggtactctcgttcagattggaattgccattgtatttcggtgctggactgcccctgggcaggacaaagactgatatgcaaatggatatttcccatctgttaaaggcacagtgagtaaaagcagtggctgcgaactctcaagtggttaata
accactgaacaggttatcatgtctgttcttgcgagagaggggcgcccgctacctttgtaaaactatttttaaggaacctgggattggtaaccctaggtctcctgggggcaaccagacgtggtctccctctcactgtgcttagagacgcatcagcttcacctcactgat
gatccccaacaagggtgaaacatgtgtatggggttgctggtggtactctcgttcagattggaattgccattgtatttcggtgctggactgcccctgggcaggacaaagactgatatgcaaatggatatttcccatctgttaaaggcacagtgagtaaaagcagtggc
tgcgaacgctcaagtggttaataaccactgaacaggttatcatgtctgttcttgcgagagaggggcgcccgctacctttgtaaaactatttttaaggaacctgggattggtaaccctaggtctcctgggggcaaccagacgtggtctccctctcactgtgcttagag
aggcatcagcttcacctcactgatgatccccaacaagggtgaaacatgtgtatggggttgctggtggtactctcgttcagattggaattgccattgtatttcggtgctggactgcccctgggcaggacaaagactgatatgcaaatggatatttcccatctgttaaagg
cacagtgagtaaaagcagtggctgcgaactctcaagtggttaataaccactgaacaggttatcatgtctgttcttgcgagagaggggcgcccgctacctttgtaaaactatttttaaggaacctgggattggtaaccctaggtctcctgggggcaaccagacgtgg
tctccctctcactgtgcttagagaggcatcagcttcacctcactgatgatccccaacaagggtgaaacatgtgtatggggttgctggtggtactctcgttcagattggaattgccattgtatttcggtgctggactgcccctgggcaggacaaagactgatatgcaaat
ggatatttcccatctgttaaaggcacagtgagtaaaagcagtggctgcgaactctcaagtggttaataaccactgaacaggttatcatgtctgttcttgcgagagaggggcgcccgctacctttgtaaaattatcattaaagtgcctttaatatgtgtgactaatggattt
cttgcagaagaactttgtgttggcatggtgtaattattttggtcaacatttttacaaacattaaccatgtttaaatacatcttttaaaccagaatgtttgccctttcattgcacacatgtaaactcgattggctctgttgatcactcctttaattacacaggccacgcctcttttaaa
gcttcattaactacttgggattttaggttacctgagaccttttatagggccctctgacaaatctcgggaggatagctcagcggcaacgtgcttgccttggacgcaaaacgacatggagcaacacaggttcgatccacaggtccgtgcttagaaacctcctgttattaa
gcacattataaataacctaccatcatcaaatattgcatcacgacacattagagctttcgtgcacttcgaccactgcttctttccaagctcgacatgggtcggaagaatgaaaggctgagcgcctcatttctgggagtaaacctgttatttcaggggacacatgtttctct

gcagccccttctcatggcaatgatctgtgctatcctgtgcatggctctgctgtggtttaatgctggaatcaaggcaaattgcgacttttgagaatggcggcaaatgagagatgttattatcgacacacagatttacaattggggggacatattcagattttaaaaatgagt
cgatgggtaacggtagctctgcacccgcattcccccccttttttttagaacagtgcaagacgggcagtgcctgcgtcttgcgcaaaacatgcatattatgggctaaacctacacaccccaaatgtatcaaggacagtgcaagctatcccggtatagaaaccaagt
gcaatacagcacgggcatggcaacctgtgcttgcactgcacaaaacagctctgtactgatcagtgcctggttctatcacacagaactgttactatatcgtccgtgcaagaaccgcttactttctgcaagtaagaatgcagactgcaagctttcctggcgacagcatg
gactgcgcaaggccccaccaagcaccacaagcactgcctggacagtggcacctcatctcacacagaacatggagactgcaagcctccctgacacaaagtgcgcgatgacgacatcagcccttgccacggtcacttgtgagggacaccagaaaggcacga
agtgagacagggcgcagtgccagggtccagaagaactgggctcctaatcctgggggtcctttgccattcccgggcatttacgccagagtgacatgtaattccgggtatttaaatgtagatgacaagcgtcacttccggtcttaatgaattcccagtgatgctcaac
atcacgtctaggtagattcctttcgagttaacgtccacctctggtagttgttactgccaggaaccctatttttgctcacctgctctgcggggagcgtaagccaagctgctcctttcattcctccagcgtcatgcgttttccccgtcatctatcctcatagtggtcccttcgta
aatcttccaacgctcccatggattcattaatccccacattcatattcatttacccacgccacctgcatccactgttcactccctctgagttccccatgacatttccggagtctgcctacatgtgtctttgatcatgacgttttcatcacacagcagtcaaaactgttgtttttgt
caatgtttgtgtgagttttggaaagtaggcgtctcacaatagtcattcggatcaacaggaactcagcaaaacaggtattttaaaaaaacaacctttctgccattttgaatggttgtctgggccaattagtttcttaaaaggccttcctggtcccgtaagatttactgtttcttc
taaatccaacaagacatagaatgtcctttaatatatacctaaatccctagtggccgaaggatcgggggggagtgagggagagaagggggggcggtgggggtggaggtattgattgccctcccggaatgggagatacgctcccaggggcctgtaggcagcgg
acatttttatcataatttcagaaaacctttcgatcccattggacatcatattttgtctccgtgtaacaggacccagtggtcttttacacaatgacgggcatccttgtatgtgatggattattacttgaagtccctgccccactccctgttggctactttagtataattgcactctt
gccatcagacagccattggaaagtggcactcacagatcctgtaaagaagcactatgctactatatcatgctatgggatgtggagagtgaggagatagtgttgacccgtcaggattccggccctcgaggcccggagttggacatccctgtgttagtgcattaatcc
cattgtgcattcatctgaaattcctttatctcaacctccccataaagtacaggattaagttctggcttccagatgcaccatctatgagcctaagccgaacaagcagtggccaagccaacagttttagcttgggatttcccattcaacgccatgtactactatgccataaaa
tgacagcagctggaaatggtgtaagtacctgggcatatgatgtcagtgcctggcaaatgtctatctaatacatatacataagccaaaatcgttagctctgccaatgtttttcaagtgttcattgaaaagaattatgcaaggatacacatatttctaatacatcaaggggc
tgaatgcactctcgtttaagaccttttatttcatgaacaactcatttactgctgtaattaaaccccttaaccgcaggaagagtgcttcaggagaacagtggcaggcggtatcaggtaatattatccccaccaaaaatatccccgtatatatggtcgtctggaacgccatag
gtgacattttttgggggataatttgtgggagcgggggttgcgggggtgtcccccacacatttaaaaataatgggggtttgcgggggtgtcccccgcaggttgcatgcctcaatatcgtcgcaaaaatatggcgatatttttgtatcgcaatatttttgcggggatatta
acacatggaacctggcaggctccatacaacacacagtttgatgtgtctgtgcgaaagatctagctctaggagaaccctggactgtataatatgggtctggaaggcaggtaggtgggaatggtcatagggaatagcatcgctatggatgctcaattgtatttattattt
atttatttattctgatttgtaaagcgcgcaacagctcggatgcgttgtggcgctggttacataaggttacataaggggggtgacaacttagttacataggtagcagtaaaaagaggtcacaacttagttacatcgacagcagaagcaaaacttagttacagtggtcgca
ggcaaagtagaatactcattgaggaacagacagtttgtagcaaagctagtggagaggtttaattaaagagagagtaaataaccaagcactggccataatctgtgtatccctccacacatccgaccttcctcatatctgtactccaatgcaatagtgttggcttacagg
tggtgctaaaagaccacctattaaccaaaactaaaaagggtttgctatgtgtatccctcacaaccatgtaagctttacttatataggcaggagaatcaaatcaccaacatttcagttctgagggtgctgggataatagatgcatcgccgtgctgatccccctgaactagc
tgtagcaaactagcctgtgctactgtttcttattttctaatacttctgaaactatgtatcctctaggtgccattgaacaggcgagtcctggattagaggaaagcgttgtgcgccgaaaagaggcaaaggattgcaactgttccctcttccgcgtgcgtaagtaatgttag
ggcggatacaactttctatcgaaatgacttcaagatgggatatagagagctgccagggccccagcacccctttccactctcctgcacctggtagacgtcagcacgcatcagtcagaagtgacggggggggtgtggcactatctaaattcaggattctctctgcattta
atgcgttcatgaagatggcctttactttttctgattgaatggaggttatgctcaacaaacctctagcgttcagcgtttagtgccccgccctattgaggtaaattgatgtaagcttcctaatccattaacaagttatatgcaatcaaacgtttttttccaatgtgtaaaatgaaca
tttctggtagtacatgattttctgttgaatttagataacacaagatttcactacatcaatgctagtcagtcaatattaaagctttggtaaaagcccaagttctgaattttgatataagattatagaggagcacctcagcgaagcaaaataccagtccaccccaaactcccag
atagacaatacaccattactgttgtataagaggttatctgcattgattgttctaccacaatgtacagttctttgccctgattacccggcagctgtggccactcatcccttgtggatacatagaggagaaacattggttgcatgtgggagggcaatagtcgttgtggtgca
agtggctctgcgcaggcatgtcttgcaattcttgaggttgcatatgggcttgtgaatcatttgtattatatctgcacatacacagtccgaaagagtgaatcatattgacataaaagccatatcactcaaacttacaaatgtctcctctgcctcatagggagacggcgaca
cggcaagatgtcccgcaacaagaacaagacccggaagtgcatgcagcattgcaaaggaatactaaagagaagtggctgcttccgagaaccatgcaacatggacttagcaatgcccttgtcataaactcctgagggatgtggcattacattgtcaggataacaa
cttgagggatactgttctttccgtttttagatgatctattccagtagagcaaagccaagttgggccacgttccccatggcctgcccagtggtcaggagtgaccatattgtagcgataacaatttcagtattgatgttgtttggaagacttacatagctcggctcacatatg
ggaggctaactctgcagtactgagtagatgtagctatgggaggcaaggctccaatctaggcattgttagaatgaatctaggtattgtatgctagaagaattcaaataattatgtaagtcagtgcgatcagaagggaaccagcagcattattgatctctttgacgctact
gctgggggtttggcggatatggcagtgtgtggttgccacatggtttctaaagataacagatagttgaagtctgtcgttataataaagctgtctgttgcctaacaaatgcgtgtttttgggagttttggtacatatgttggaggttgtgccacctgtggccaaataattgcca
atgaaaagtgaacaaaagaaacttatgaatgaagaaggatctttggttggcatggcaattgcaatacttgttgcatgtattgactgaagta

**C:** Sequence of cDNA (coding region in red and highlighted in yellow)

Aaggtgatcggctgagggcttggatcagtagtgctgtgccttttcacttggccctacaccttctcagcgagcggctgcaatgaggaccaccctggcgatggcactggtgcttctgctctgcctctctcactgcacgggccatagaggttccctcaggcatgagaa
gggcctgaccgagcaccagtccttacccgggcgacatcaccatggacacggtgccattgaacaggcgagtcctggattagaggaaagcgttgtgcgccgaaaagaggcaaaggattgcaactgttccctcttccgcgtgcggagacggcgacacggcaag
atgtcccgcaacaagaacaagacccggaagtgcatgcagcattgcaaaggaatactaaagagaagtggctgcttccgagaaccatgcaacatggacttagcaatgcccttgtcataaactcctgagggatgtggcattacattgtcaggataacaacttgaggg
atactgttctttccgtttttagatgatctattccagtagagcaaagccaagttgggccacgttccccatggcctgcccagtggtcaggagtgaccatattgtagcgataacaatttcagtattgatgttgtttggaagacttacatagctcggctcacatatgggaggcta
actctgcagtactgagtagatgtagctatgggaggcaaggctccaatctaggcattgttagaatgaatctaggtattgtatgctagaagaattcaaataattatgtaagtcagtgcgatcagaagggaaccagcagcattattgatctctttgacgctactgctggggg
tttggcggatatggcagtgtgtggttgccacatggtttctaaagataacagatagttgaagtctgtcgttataataaagctgtctgttgcctaacaaa

**D:** Amino acid sequence (signal peptide shaded)

MRTTLAMALVLLLCLSHCTGHRGSLRHEKGLTEHQSLPGRHHHGHGAIEQASPGLEESVVRRKEAKDCNCSLFRVRRRRHGKMSRNKNKTRKCMQHCKG
ILKRSGCFREPCNMDLAMPLS

**Fig. S16.** Gene position and architecture (**A**), genomic DNA sequence (**B**), cDNA sequence (**C**), and encoded protein sequence (**D**) of the unannotated *cxcl17* gene in the NCBI reference genome of *Ambystoma mexicanum* (axolotl). The information was downloaded from the NCBI gene database (https://www.ncbi.nlm.nih.gov/gene).

### A: Gene position and architecture

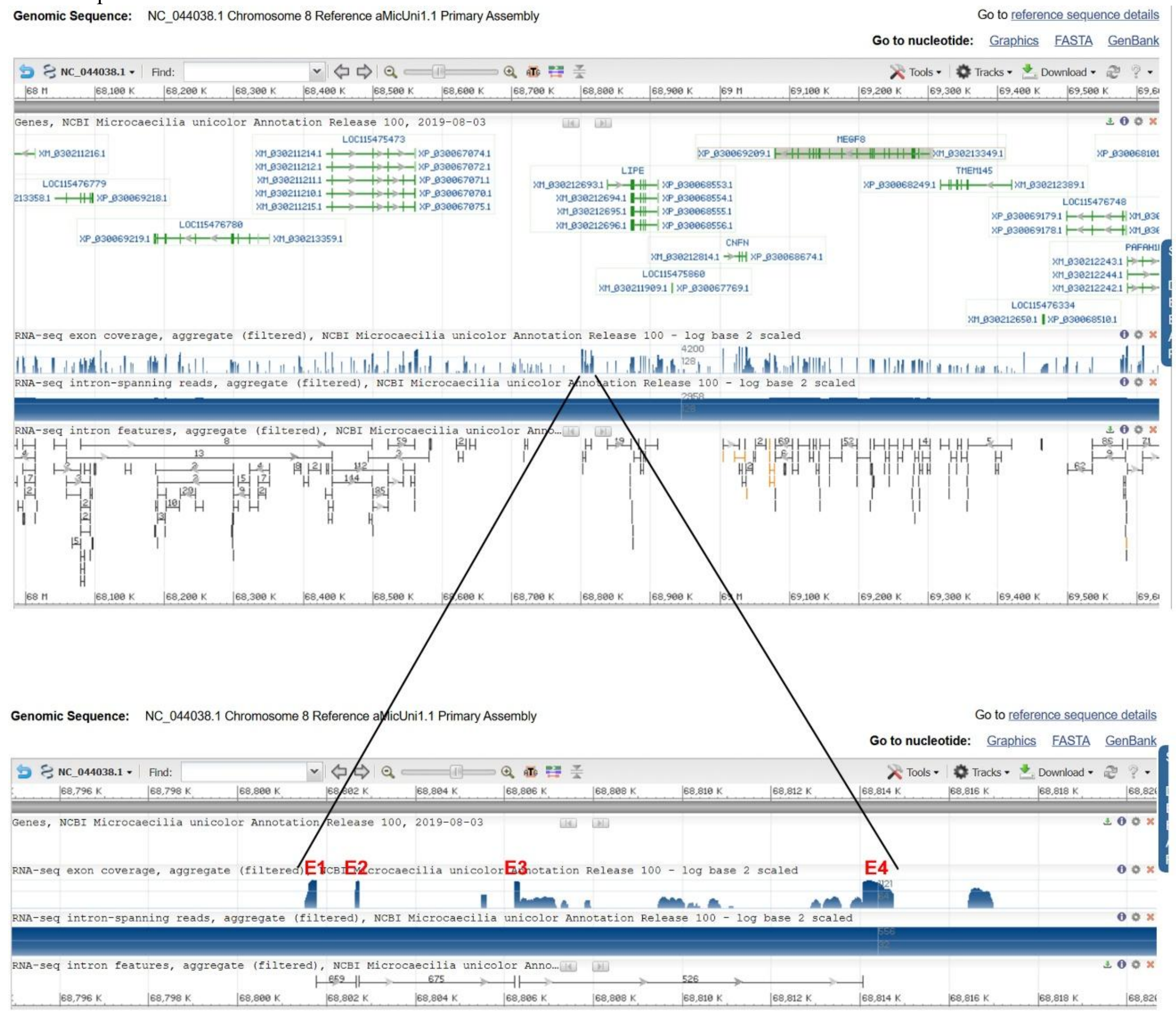


### B: Sequence of genomic DNA (exons in red and highlighted in yellow, coding region underlined)

gagggccaggcaggcgggggagggttcacttaatttctgaacaagttagttctgtttcaaaaggctttgagaattattcacagatgattaccatccctttcaatgactttgcgtcttgtatttatataagacctgcctcttcttagttatgggtcagtctgaacttagggtgg
cggagacggactgtgggactgtaactgccgtatcaaggaaacaagttttgtcctgtgggttttgattagacaccttttgtgatacaggttttgtcctagctcagcatgagggccctagctgtccttgtcctggtgatttcagccatcactgtctctctctgtcatgaggag
aaaggtaagcaacattattcatcaatattaaaatatgaaccgtgagagtcccttttaccgaaatgagtgctgagaatcccccattcctgaactgtgttatataaaggcctctaaacttttgagttgcccaaacttgagatctgtgttttctcttatatgcagtatgagttctcc
acaatacctaaacagagaactatctatatctcaattgtgaaacttccactcacagctgaactgggagatctttttacattatatcatgtgttgtattgggttattcagaattatattaagagctgggacagatagaagtattaaaagagcagaggtatagcgattatcccatt
ggagatattgaaatgttgaggtataaagatgaaggaaaaaattccatagataaaatccaggagacaaaaagcagaactagacagaaagaggtgaaggaaccacaacagtttattcatggggcaacccgacacagttgtgttttggcatcttcatcaggagttcag
tatccgccttgagtgaactccttcaaaaaggtggtaattaaatgctaataaataaaaagtatctattaagagtttttaaatataattttctactcccttatgtgacagcggattggattccaaattgactaatagtgctgtcagccagcagcatctcctgacacaggccattta
tgccgaaatatggctgtgtcaggtcacctcatgaatgaactgttgtggttacctcacctttctggctagttctgctttttgtttccttgatgacgaaatgttgacgatggttcttcctgaggacacaactccaagctctttaatggggaattctgtgctgtcatctactaacag
gaaaatatttcttctcacaacagattcccagaggcagcagcatgggctggcagggaggccctcagaggtctcaagcacacatggagaacaaaagattggtgagttcaagcatgaatagttattccacttcttcctcttctggtgtccttgtcacactggatgaccttg
gccattgcttaagaacataagaatagccatattgggtcagactaatggtccatctagcctagtatcctgcttccagcagtggccaatccaggtcacaagtagctggcagaaacccaattagtaacaccattccatgctactaactccaggccaagcagtagcttcct
ccatgtccatctcaataatagcacttatgccagccacagagccgatttaaatgctcatgcctaaattgttacattatgcttataaattatagtatttcagaattacatgtgttactatgtgccccacccattccctgccagactctgcccgtatgtacatccaccttaaaacta
cataccatcgcatttaggggcccttttactaagacatgtaggcaacctatgtgtgtcaaattagaactgctgcccagtaattctaattttgacacacatccaaaacgcgcgacagaaaataatttctattttttaccacatggggctaacctggtggtaattggcaagatc
ataagaagcaccacagactacctccagttcagaaaaaaactaaaaacctatcttttcaagaaattctatgattgatctaagcgcactaataatattacattatcactctgaaataacaatgtaccaaacaaacatgtaattatagtttatctatgtaatctcaaataactgtta
aatgtaagccacattgaaccgaaacttttggataattgtgggatacaagcatgaataaataaaataaataaaaataaaaagtaaagaaacactcagatatactgtatgtgcaatgcataattcatagagtagagaaaagaacaaaagaaagagataaacctctcgtc
cagctgctctcagtatcaaatttcaggaggtacatcgactgttattgaggtggacttggataaagccaccggtttgacacggttccccacaggaggctcttaaataaactcgatgggctgaagataggtcccgaagtggtgaactggattaggaactggttgacgg
acagacgacagagggtggtggtaaatggagttcgctcggaagagggaaaggtgagtagtggagtgcctcagggatcggtgctggggccgattctgttcaatatatttatgagtgacattgccgtagggttagaaggtaaagtttgcctatttgcggatgatacta
agatttgtaacagagtggacacccgggagggagtggaaagcatgaaaaaggatctgaggaagctagaagaatggtctaaggtttggcaattaaaattcaatgcgaagaaatgcaaagtgatgcacttagggagtagaaacccaagagagacttatgtgttagg
cggtaagagtctgataggtactgagggggagagggatcttggggtgatagtatccgaggatctgaaggcaacgaaacagtgtgacaaggcggtggctgtagcgagaaggttgctaggctgtatagagagaggtgtgatcagcagaagaaagaaagtgttga
tgctcctgtacaagtcgttggtcaggccccacctggagtattgtgttcagttttggaggccgtaccttgcgaaggatgttaaaaaaatggaagcggtgcaaagaaaagctacgagaatgatattggatttgcgttccaagacgtatgaggagagacttgctgacctg
aacatgtataccctggaggaaaggaggaacaggggtgatatgatacagacgttcaaatatttgaaaggtattaatccgcaaacaaatcttttccggagatgggaaggcggtagaacgaaaggacatgaaatgaggttgaagggggggcagcctcaggaaagat
gtcaggaagtattttttcacggagagggtggtggatgcttggaataccctcccgcgggaggtggtggagatgaaaacggtaacggaattcaaacatgcgtgggatatgcataaaggaatcctgtgcagaaggaatggatcctcagaagcctagccgaaattgg
gtggcggagcaggtgggggggaagaggggttggtggttgggaggcgaggatagtggagagcagacttatacggtctgtgccaaagccggtgatgggaggtgggactggtggttgagaggcgggaagcactgctgcgcagacttgtacggtctgtgccctg
aataaggcaggtacaaatcagggtaaggtttacacatatgttgtcttgttgggcagactggatggaccgtgcaggtctttttctgccgtcatctactatgttacccattagaggcagtagcatggaaccttggcactttttgagattctgctagatactcgcgatctagat
tggtcactgttggaagcaggatactggtgggctagatcagtacttctcaactctttcctggagacacatctagctaatcaagttttcaggattgtcacaatgaatatgcatgagacaaatttgcatattgtagagacccattatatgcaaatctatttcatgcatattcattgt
gatgatcctgaaaacctgactgacaagatgttcctccaagagagagctgaaaagcactaggctacatggacctttggtcatacccaatatggcagttttatgtctcatcccagtgattcccaaacctggtcctggaggcaccccagccagtcaggttttcaagttatc
tactgccaccactgcatgcaaatctctctcatgaatattcattgtggatatcctgaaaacctgactggctggggtgcctccagaaccaggttcgggaatcactgtcttatgatgtgagaatgaaggagcgatagagaaactgtggtgctgcctcagtcccagaggtg
ccagagatacatcgtgctgctgatggtacttggtagttctctctcttatcttactttttttctttttttctttttttacatccataggggttgaattagcccaagtatgaaactcatgagcaacagcactaatcatgacaatttcaaccctaggcactttccattcttgcattcataatt
cagttgcctttaaatatgcaaagtaatattcacttacttcttggattgataagaagagctgggaatggtagctcagagctctgttaggctcccatttggaaagttaaatatcattgtttatgtattgagcagggctccctcaacagaaatagaaaatagagagtgtgggtg
ctgttgtcattcctaagatggcctgttagcaatcaggtctttttgctcatataattatttaatctagggtttgaggagagcctgttgggtgtagtagagggggaggcttgcccctttggggagctgaagagtctctatgcctggttaactttttaagttcctgtctctgccattt

ctccctctttacagccttgcttcctcttctgtaacagcttgagtggtttgtgtcttttcagattcactgggagctgggaaccaggtgacaaatgaattgagccaccgaaggagagagagagagacatcagatgctactgtcagaagacggcgaaacttactagaagtaa
gaagccagcctgagggagcagcccaagggtgatcctaccacagcataacagaggaggctctaatccaaggggagaaagcccgtgtcaactcaaagggaaaccctagccagtggtgtaatgacagggaggttgggagtgggccaggacccccaggtgc
cagcttgaaaggggtgctgaaggaggagtgatgcgagcccctaggcatctgcctatactgtgaattctgtgcacaactaattcacagtctccatggcctgccagatagacagcctctactacaagctctcagctacagcctatcctgcctgaccaatccgggtcgt
gaggcagcccacttgcagtctatactgcccctccaaggctattacagttctttaggtttcctagagacatgtgcagataacactcagacagttaatataggtaataaaacaacttttatttacttactgaagaattggtgaatctggcatacaaggtgcggtgtaagctca
cctgccaggaatcaaggattataggagcatataagaacagcaaatccaacaatccttcttgtctgtcttcggaattttgcctagctacagagtttcttttgcataagaaaaaggtcagtagtcccaagaaaagatgttattgcaataaatgagaaggcagagagtcag
ggaaaaaggggtgccaaaacaaataaaaggcacttaaggatatacctcagctctgaccatcctaggctggctatcagccaatagggaaatccctggattggacaggaaggataggctgtagctgagagcttgtagtagaggctgtctagaccatgaattagatc
tgcacagaattcacagtataggagtttacttcaaggaactggaaggtcaaaaaaaaaaaagcacgggggggggggggggggggggagacccatggtaatgccactcaccgtcactcaaggagaatcatctcccagcttaaggatgatggtcacagccaaa
gaggcagaatcaagcaattagagaaacccaatcccagcaagaatagattatcaaaagaacttttattcacaaacacataaaacacaatataaaaacaattcatatgcagcccaaataattcaaaagtcaatatgatagtcacttagtccattgttctacatctttcagta
agaaaataagaacataatagctatactgggtcagaccaatggtccaactagcccagtgtccttttttccaacagtggtcaatccaggtcacaagtacctggtagaaacccagatagtagcaacattccatgctaccaatcccagagcaagcagtggcctccccatgt
ctatcttcatagcatactatggacttttcctctaggaacttgtccaaacctcagataagctaaccactgttaccacctcttccggcaacaagttccagagtttaatctttgagtgaaaatatatttcctcctatttgttttaaaagtatttcaatgtaacttcattgagtgtcccct
ggtctttgtactttttgaaagaggaaaaaaatcaattcacacttacccgttctacactactaaggattctgtagatctcaatcatatctcccctctaccgtctcttttccaagctgaagagccctaacctctttagccttttcatatgaaaggagttccatcccctttaatatttt
ggtcgctcttctttgaaacttttctaatttcttttctgagaaatggcaaccagaattgaatgcagtactcacagtggggtcgtaccatggagcgatacagaggcattatagtattcttgttcatatttaccatccctttcctaataattcctagtatcctgtttgctttgtttggtca
ctgaagcacaatgggtagacgatttcatctagatctttttttttgggtgctgactcctaaggtggaccatagcatcaggtaactgtgattcagattattctttccaatgtgcttcacttagcatttgtccacattaaatttcatctgccatttggatgcccagtcttccaatttcct
aaggtctccctgcaatttttcacagtccgcatgtgttttaacaactttgaacttgtcgttctgctttccagatcatttataaatatgtaaaatagcatcggtcccagtacagatccctgtggccctccactattcgccctcctccattgagaaaaatagccatttaacactacc
ctgttttctgtccaataaccaattcccaatccacaacagagcattgcctcctatcccatgacactttaattttctcaggagtctgtcatgaggaactttgtcaaaagctttctacaaatctagatacagtacatcaacaggctcacctttatccacatgtttattcacgccttca
aggaaatgaagcaaattggcgaggcaagagttcccttggctgaacccatgctgactctgtcccattaaaaccatgtttatctatatgttccataattttattctttataatattttccactattttgcccggcactgaagtgaggcttactggtctgtaatttccaggatcacc
cctggaacccttttaaaaaatcagcgttacattggccaccctccaatctccaggtactacagacaattttaatgacaggttacagatcattaactgcagatcagcaatttcatatttaagttctggatgcatgccatccggtccaggtgatttactactattctttaatttgtc
aatttggctcaatacatcttccagcttcaacaagatttctttcagttcctctgtatcatcacccttgaaaagcatttctggtacaggtagatttcttacatctttgtccataaagaataaagcaaagaattcattaagtctttccactatggccttgtcctcgctgagtgcccttttt
gctccttcatgatctaatggtcccatggtttccctcacaggttttctgcttctgatgtacctgaaaaagttgttactatcagtttttgcctctgcagcaagcttctcttcatattctcttttagccttctttatcagtgctttgcatctaacttgccagtgcttatgttgcttcttattttct
tttgggtcctttttccattctttgaaggatgttcgtatggctctaacagcctctttcacttcaacttttatccacgctggctgtcatttgctattctttccagctttgtaaagaagcccaatttgcttactgagatttattccaatttatcagcctgatctaatttccttctttgtatacgg
ttgtagctacaagatatattcaaataaatgaccattgttgttctaaattctatgagtttattttattgggcagactgggtggaccgtgcagggctttatctgctgtcatctactatgttactatgatactagcccctgaagcaggcaatgtcactgaaacatgccgggcttcc
attgaaagatgtagaacaatagtctaagtggttatctttgtggtttttgtattatttggactgcatatgaattgtttttgtattgtattttatgtgtttgcgaataaaaattctgtatttttcagttaggattgtgttcctctgcttgcttcattgtgcttctgatatttgtgcggttggaattt
gtttttcttctgtgcacagccagctaaagggagaactctggtgaccttaccgtggtctagaggttcaaaggaattctttaattattgtatggggttaattagttaaaggcctacctgatagtataaactgaaatctcccctgtacttaagcacagagagcctccgtcagag
ctaaacagtaacacagagtttttgaaaggtgaactaccaggtgaattaggctataggaggacgaaggaggctctaggccagtgtttcaccttttttcagctcacaacacatgcacataaataaaaaaatattgtggtacaccatcccatctctacccccagtgcatgata
gagacctcactgatcccagcacaatattatattgtatagttgactaatttattttctgtgtgagatcagagcttgctttgatgtacccagtctctttatcctggcaggaattggtgacagtcaaacttaggggtgcttttactaacctgccagcacaggttttactacacacta
gccattagcacaggcctgttctaacagctaccacgtgctaaaaccagccagtgtggtgaaaatcccacactaactgcttagggatagggttaccatatgtctgggtttccctggacatgtcttcctttctggtgactatcagggcatccgggcggtttttgctcatgac
agtttttgtctgggttttaattccgggagcctgtgcctgtaccgtgtacgttttcctttcatttcccgtgtgctccagaagtggaagcagagaggcaaggctgacatcaactcgctcccctctgtgctgactcagagccaatgccaggactcagttgctgaaccccaca
accggtggtggtaagtccccagagtgagagtctgactccacgcagacacacacaagctgtgctgttcaccctctcccccactgaaatcgcactgcaagaatggcttcatatatcgggctgtttcttcctcctccttttgtctgcctagaggcaattgcttatgaaccgc
aggggtctctgtacagcctcccgcggttcacatgagcagtcagttctctaggcaggcacaggggggagaaaacagtccagtatgtagccgttcaggctctcctcctcttgcaatatgccttcctgtggctgtgtatagcaggcacatgtggaggggcataatcgaa
cgcctatctccatgggcgtctatttccgaaaacgggtacgtgaagaggcgggacagactgtattatcgaaaaaatggacgtttttcagctgggcgtttgtttttttttagcgataatggaaactaaaaacgcccagctcaaaaacgtcctaatccgagccatttggtcat
gggaggggctaggattcgtagtacactggcccccctgatatgccaggacaccaactgggcaccctagaggtcagtgcggtggacttcagaaaaagctcccacatgcatagctcccttaccacgggtgctgagcacccaacccccctcccccaaaacccact
acccacaaatgtacaacactaccatagctcttaggggtgaagggggcacctacatgtgggtacagtgtgttttggagacctcccatttaccagcacaagtgttataggtaggggggggatgggcctgggtccacctggctgaagtgcactgcggtacccactaa
aagtgctccagggacctgcatacacgcaggtctctaggacttgttgctgctatataacattggcacaccagttgacacctgaagactaatctctccgaaaatgtcctttattggaataagcacgcttactcacagttaactgcagatcagaggttgtgccccactggc
aacgagtctccctggtactgagatgagcagtaggtcagagctggcagaatggtgtacaatgccctctttcagccacattcaaggtaagaactaagttctgtaacgtggctaacacatgaaagggatctaaaactggcttacaaaaatggccactacctcatggact
accggaaacaaaacagggcacactctgacccagtaagcagggggaaaagcaccatgggagtagagcctaccaactaccaacatcgtgagcatttgccacaagctagtggaatcacggagcccaataccctacacccacataacagaaaaggtgtcacact
cacccgagagccacatcaaaaccagggaacggctgtcagaggatagaacacattctgctgtcatggaggtgggtacggcatttgaggctggcataaaggctggaaaaaacgtttttaaagtgggttttttttaatgggaggggggttagtgaccactgggggagt
ccagggaggtcatccccgattccctccagtggtcatctgggcagttggagtactgttttgggacttgttcgtgaaaaaaaaagggtccaaaaaaagtgacccaaaattgcggtaaaaatgcctttttttttgattatcagctaaagacgcccatctctccttggccgata
accacgccccacttccgccttcaccacgcctccgacacgccccgtcaactttacccgtttccacaacggattgcagttggaaacacccaaaatcggctttcgattataccgatttgggcgcccacgggagaaagatgcccatctcccgatttgggtcgcaatatag
gcgtttttctatttcgattataagcaggatagtcacctgggggggggggggggcaaagcacggtgactgagcaccatatgggggggtgccaaccatttttatattttgtctccacaaatatggtaactctaggtaggggcaggttatcagcagaaaaagggcagag
cggaggagtgcccaccagagttaactagcacataggtcggtgctgcatgctgccaaaactgccagctgtgtagctgtacataatactacatcgggggggggggggcattaaatgcagtcccactaccatattctggaagtgcagacacggccctatagggggag
gaccatcaaccacactctccaagaacagaaacccccaagagctgtaccccacctgatccaaaaccctcttaattcaacacccccccttgctctaatcctccttctaacaggctcatacatttgtgccctattttcagccaaattttaggcacataaaatttcagttgaaaat
tcaattaggagtttaaaagctaggttcataaatgtaggcctgccatttatttgcctagacttaagagcctaatttatgaatctagtgctgaaaatcagtgctaagctcctaactgcttttccccactcaaatccacccctgttatcatccactttttatactccaaaatttaaga
gtttagtgaaattttcagtaaaacatttatgcactcagtcctagtaaatattcatatgtggagttcttttcattatccacttctgtattgattgtatactgctttgattttttattgtgaatgacagtacatcaagagcctaaaataaatacatacatacattttcagagctgccacttt
ttgagcttaactctcaaggttaggagcataaatcctatcaggcccatggtatttactgcattggctgttaacagtagaggagtggcctagtggttagagcactggtcttgcaatccagaggtggccgccggttcaaatcccactgctgctccttgtgatcttgggcaa
gtcacttaaccctccattgcctcaggtacaaacttagattgtgatccctcctgggacagagaaatatccagtgtacctgaatgtaactcaccttgactgctgaaaaaggtgtgagcaaaatcaaaataaataaatataaacacagctgtgcactaatggttaacctgca
ctaaaatctactctaattgcttatcacacttaagtgatcctaatgaaatctaaaccccaaatccaagccaaataagaccccaaacccacaccagcctgtccataaacctctaaatccactcccaggtctgtttgggtttatccataaagcaaaacctgcaaatgagcct
cctctgttgaagtctggaaagatgagctactgcctaggaagaagaatctaacccttttctgttctctgtgtctcaggaatgaagagaaatgggaaagcaccaagcaagtataaccaaatgaagtgcatcaaaaaatgccacagagtgaagtggaaacagcctatg
atccccctctaaaaggttgatttccctgtactaagggggttctcttctgactctaggatccccttctgactgatgaatttcactgatcctgctgtctgttatgaatctgtctatggtgcattttcaacctacagtaatgtctgctgacttaatatctgtggcttttgtaataaactc
ctagaatttatcactgtgtaatgagcagaaaaataatctttctcaagaatctgtgactgagtttggctgcggagaaccaaagcatgcatgagctatgagtaagagatatgtaataagaatgtgaatgtataatccacaggaggagtgtcataacacatatgtaaagtat
ttgtatatcaaaggcactagtaatacatggacagcaaatatttgatgtataaataaacattaacattgtaataagtgcatttatcttactatcagagaaaaccctagaattaggtttcatatccatgtgccagttccctgagctatccagtcctgtttccagattcaccctata
gggtagagaggtgtggtagccgtgttagtccacttttaaaggtaatcaatagaaataaaacaaaataaaattatacctttttttattgggcataacttaatacagttcttgattagctttcgaaggtttcccttcttcgtcagatcggaaataa

**C:** Sequence of cDNA (coding region in red and highlighted in yellow)

Tgaacttagggtggcggagacggactgtgggactgtaactgccgtatcaaggaaacaagttttgtcctgtgggttttgattagacaccttttgtgatacaggttttgtcctagctcagcatgagggccctagctgtccttgtcctggtgatttcagccatcactgtctct
ctctgtcatgaggagaaagattcccagaggcagcagcatgggctggcagggaggccctcagaggtctcaagcacacatggagaacaaaagattgattcactgggagctgggaaccaggtgacaaatgaattgagccaccgaaggagagagagagagacatc
agatgctactgtcagaagacggcgaaacttactagaagaatgaagagaaatgggaaagcaccaagcaagtataaccaaatgaagtgcatcaaaaaatgccacagagtgaagtggaaacagcctatgatccccctctaaaaggttgatttccctgtactaaggg
ggttctcttctgactctaggatccccttctgactgatgaatttcactgatcctgctgtctgttatgaatctgtctatggtgcattttcaacctacagtaatgtctgctgacttaatatctgtggcttttgtaataaactcctagaatttatcactgtgtaatgagcagaaaaataat
ctttctcaagaatctgtgactgagtttggctgcggagaaccaaagcatgcatgagctatgagtaagagatatgtaataagaatgtgaatgtataatccacaggaggagtgtcataacacatatgtaaagtatttgtatatcaaaggcactagtaatacatggacagca
aatatttgatgtataaataaacattaacattgtaataagtgcatttatcttactatcagagaaaaccctagaattaggtttcatatccatgtgccagttccctgagctatccagtcctgtttccagattcaccctatagggtagagaggtgtggtagccgtgttagtccactttt
aaaggtaatcaatagaaataaaacaaaat

**D:** Amino acid sequence (signal peptide shaded)

MRALAVLVLVISAITVSLCHEEKDSQRQQHGLAGRPSEVSSTHGEQKIDSLGAGNQVTNELSHRRRERDIRCYCQKTAKLTRRMKRNGKAPSKYNQMKC
IKKCHRVKWKQPMIPL

**Fig. S17.** Gene position and architecture (**A**), genomic DNA sequence (**B**), cDNA sequence (**C**), and encoded protein sequence (**D**) of the unannotated *cxcl17* gene in the NCBI reference genome of *Microcaecilia unicolor*. The information was downloaded from the NCBI gene database (https://www.ncbi.nlm.nih.gov/gene).

**A:** Gene position and architecture

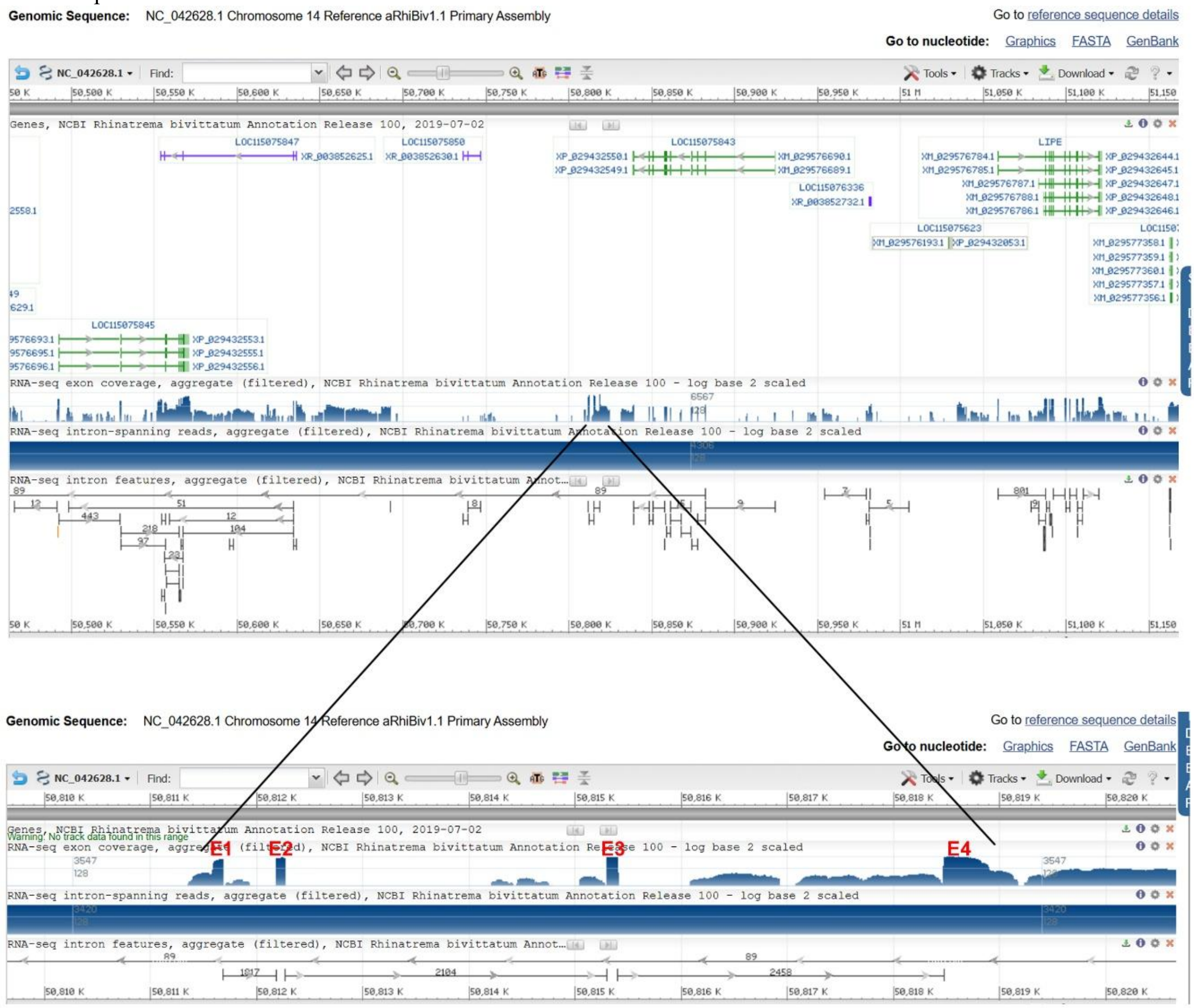


**B:** Sequence of genomic DNA (exons in red and highlighted in yellow, coding region underlined)

atggctttgcctcttgcaacactaaaagacccgcctcttcgtagacataggccagtgtggaggagactgacaggttgagggattatattataaccatcctcccacggaggtgattcgtccggtgcattttgtccttgctcaggatgagagcagtggccgttctcttcct
ggtgagcgctgtcattgctctctctctgtcttgaagagaaaggtaagagaccctccacgtcacctgcactaactgaagtataaaaacctccaccccctatcagagctgtgagagtctccttttacctgaatggtgaaagctcccccaggtacctgagtgttgagaac
ccccatacccgagttgtgtgagcctcctgtacctgagatatgaaagcctccatatacttcacctttgagagttgaagtgtatgaattttcccatacttgagttatacgttttctctaatactggagcagcgagctctctcatagctgggctcacaaactcctgctaatataat
atgtgttatattgcatcctagattcaattctgtctgtgctggaatagctatgaggcaagtgttataatagcagaggttactgggaggaagctggcagtgggatagtgaaatgttgacaatgtttctacctgagaccacagttcaaaggtctctaatgggagatgtttttgtt
ctcctcaactgacagaaatatgtgtcttcccacaatagattctcacaggcaggcacaggagttggcagggaagcactctgagttcttgagaagacctggggagcaagagattggtgagttaatgtgtgataagtcattctgtttcttcttcctcgtctcgtgtccttgca
ttaggctggatcactgtgcacaagtgatggttcttgtagttgctgtagtatggtccttcttcatacaggtttccctaaagcatcaataaacaagcaggtgcttcgtgatctgcacgtttatcctatatactctttgttattgatttcatcaactaggactaatctaatctttgattt
gccagtttgcatgcattggaaattcaggcattgccatgttgaccattctgcaagaaagtataagtgatagaaaagagcttaccggtagtaagagactggtgaatgtatccatgggctattagtaaaaatttgtaacttatcattaaaatcatccattgtacctgaagactg
gaggatagcaaatgtaaccccaatatttaaaaagggctccaggggtgatccgggaaactacagaccggttagcctgacttcagtgccaggaaaaatagtggaaagtgttctaaacatcaaaatcacagaacatatagaaagacatggtttaatggaacaaagtc
agcatggctttacccagggcaagtcttgcctcacaaatctgcttcactttttgaaggagttaataaacatgtggataaaggtgaaccggtagatgtagtatacttggattttcagaaggcgtttgacaaagttcctcatgagaggcttctaggaaaagtaaaaagtcat
gggataggtggcgatgtcctttcgtggattgcaaactggctaaaagacaggaaacagagagtaggattaaatgggcaattttctcagtggaagggagtggacagtggagtgcctcagggatctgtattgggacccttacttttcaatatatttataaatgatctggaa
agaaatacgacgagtgagataatcaaatttgcagatgacacaaaattgttcagagtagttaaatcacaagcagattgtgataaattgcaggaagaccttgtgagactggaaaattgggcatccaaatggcagatgaaatttaatgtggataagtgcaaggtgatgc
atatagggaaaaataacccatgctataattacacaatgttggattccatattaggtgctacaacccaagaaagagatctaggtgtcatagtggataacacattgaaatcgtcggtacagtgtgctgcggcagtcaaaaaagcaaacagaatgttgggaattattaga
aaaggaatggtgaataaaacggaaaatgtcataatgcctctgtatcgctccatggtgagaccgcaccttgaatattgtgtacaattctggtcgccacatctcaaaaaagatataattgtgatggagaaggtacagagaagggttgccaaaatgataaggggaatgg
aacaactcccctatgaggaaagactaaagaggttaggacttttcagcttggagaagagacgactgaggggggatatgatagaggtgtttaaaaacatgagaggtctagaacgggtagatgggaattggtttatttactcttttggatagtagaaagactaggggggc
acttcatgaagttagcatggggcacatttaaaactaattggagaaagttcttttttactcaacgcacaattaaactctggaatttgttgccagaggatgtggttagtgcagttagtatagctgtgtttaaaaaaggattggataggttcttggaggagaagtccattacctg
ctattaagttcacttaaagaatagccactgccattagcaatggttacatgcaatagacttaggtttttgggtacttgccaggttcttatggcctggattggccactgttggaaacaggatgctgggcttgatggacccttggtctgacccagtatggcattttcttatgttct
tatgttcttaaaatgtagagatgatggctggagagtacatgggaaatgagagactgctgaattcaggcatgggttggtcacattgcaggaatgcattagtgattgtgacaaaatgaacatggagtaagaaaagaaatactgatgaattcatatgtgggtcagagaca
cataggagtgcaggcacgatgggggagatgggggaagagagcatgtagggaatatgatattgctgctgaattcatgtgtggacctgtcacactgagatagtgtgcagaagtgaagagagcacagggtagggagaaattgaagaattcgtacatgggtatttca
cactgtggaagagcagctatgatgggaagagactggaaatttcacgcttggttagattatgctgtggggggtacaggaatgatgggagagagctcctagggagtcagagttgtgcattcatgcatggattcatcatactgcgcgaaagcagggatgatgatgactg
tatgctacctcggccccacctgtccctgtgtttctgttcaaggggctcaggatttggctattttaccttatatttataaacagagaggggttgaattagcataaatattagcagcaccaatcgtgaaaacttcaacctgggcagctttccattcatgcatctttcataagcat
tcattactttaaaaaatacaaaacaaatctatcaaaagtaaaagcaaatcttccattcacttggatggataagaagagcttggaattctcccttggagctcctacagcgttcttgttaggatatttaaatatcaatataaaagtttccatcggtgcccttattcccaaatcatt
gacaaatagagagaatatgggggctgatctcgtatctggaatggtctgttagcttagcaatgggctcttgtgactcagagaattatttattctacagcatgaggtaagtctgctggataggtggagggaggaggctgccaactgtagggaattgaagaatctctgtg
ctcagttaactctgtaactaatctcaggtcctaccatttctccctcttcacaaacttgtctcctctgcactgcctgtaacaactggaggtatttgtgtctttgcagatacattgggagctgggaagtgggaggtgagaaatgaattgaatcaccagaggagagaacgag
ccatcagatgctactgtcagaagacaccaaaacatcccagacgtaagaaagcagcctgacgggccgatacagtaaaatcgcgggagagcaggcgagcacccgctctcccggcgtgcgcataggccactctcctgtgcgacaatacagtattttaatttattaa
aattagggccagtagtaaaaagaggcgctagggacactagtgcgtccctagagcctctttttggctgtcaaacccagctgtcagtgggtttgacagccgacgctcaattttgccggcgtcggttcttgagcctgctgacagccacgggttcggaaaccagacgc
caacaaaattgagcgtccggttttcaacccgcgagccacgggcccatttaaaaattttttttacatttttgactttttttaacttttggggcctccgacttaatatcgccatgatattaagtcggagggtgcacagaaaagcagtttttactgcttttctgtgcactttcccgg
tgccggcataaattaacacctacctttgggtaggcactaatttcttaaagtaaaatgtgcggcttggctgcacattttgcttactgtattgcgcgggaatacctaataaggccatcaacatacatttgcatgttgcgggcgctattaggttcgggggggttggacgcgc
gttttggatgtgttattacccccttactgaataaggggtaaagctagcgcatccaaaacgcgcgtccaatcgtgggctaacagtgcgctccgccggagcgcactgtactgtatcggcccatgagtggggcaaatccagctcagggaggatgatcctacccaagca
taaaggggggagaagacaaccccaacccaagctggggtgattggctgccaggtcatgccaggttaccaactcggcacgtttcttaacttacttccatttccccattgtgaaaaagtggaaagtggaggctgtcgctggatccaacctggtaccttttttatttagaaaat

tactaccaagcacaaggggtgggggagtggctccagcccaagaggtgatcttacactagcccattgcaaatccctgatattacatattaaatatgccaataatcagagggcatcacaacccaaacagaaagagtcaaatgtaactccacacatctaaggcaagc
aaaaataaaaggggaagacaatactaagaagtcttaaatcaaagctagcattcaccaaaatttgtcacaatatacacaaggtgtcagcaagccggacggtgtgtctgataacacaaaccaccccgatcaatcattcaccttgtttgaaaaaatattttttagaaacat
ttttttaataaaaattatttgaaataaacaagtcattatatctttttgaaagccctttgaaagtattttatccaggtaagtagcagcaaaccaagccagatagcaatataagtactgacttatgcagcgatctggcagctgcttcccactgcccattagatggataaaccaat
atttatccaggtaagtcagttattattttggtagtggctacagctgccgggacttacccagagagtttttgactatttttgaaacttttaaaaaactttttactcgtttttcataacagctggaaacttatttatttaaaagtatttatataccatttttcaaaacagggtttttgtca
aaacggtttacagagtaaaaataaaataaaattaaagtaaaataataattaaaataatgttaaaatacaaaaactggtatatacctaaaaaggtactgatattaacaaaagtctaactaataatggagccggctctaaccaggggttaactttccagcactataaaagg
caagctggccagatgcatcctctctgcatcaggaggtaatgcttaatggcctcacctgcatgggatttccatgccagggcactaagcagtactcccgtttggaaatgcacattatctgtgcgcaaaactcctttctgcataatgtgctttgctaggccaagtccccac
ctataccctcaaattcctctatcctgcttaaaaagtatgatggttggcatcagcctccacctgcctgcaacatctgaaatcgatgaacccccccccatacttcagaaaccccttctgcgctgggattgtggtaaagtgcgatcctggcagtttccaaggttaattaaa
ctgcagcactagaagtgtaattactagtgttgcattgatttcatttatttatattctgtttcagcagttcaaagcagattacattcagttactgtaggtattcccctacttcctgaggcaaaggagggtgaagtgacttacccaaggtcacagggagcagcagggggattt
gaaccctggcttccctggtttgcagtcccctgctctaaccacgagggcttctcctaaccagggaagggagtggggaggatcagtgctgatcatcctacttttctttgaaggggtaggaggcttggcctggcaggacactttttatatattgagtggggggggctggg
aattggaccaatagttgccacttaattggctacattcttttgaatattgctggttaagccttaggactgctctgagaaaggcctaacgttatcttgctaacttagctgattttcagctgtatctggctaagttagtctgatatattagctccaccccagaacacctctattttatc
cagctaaaatttagctggatgagttccctagatattcaaaagtaggcattcagccaaataactcacaattatcccactaaatgcctttgaatgtggaccacttggtgtacaatccataaacccagatccagatctagccatacattctaatccaagccaaggtttgtctat
agacccctaaacccccactcaggtctatccatagacccccaaacctatgtccaggagtgtccatggaccttcaaacccaagcttgggtctggccatagttcccccaaaccacaatcaggtctgtccatgatattcacatctgtccataaattgaaatctacacctgg
gcctggagagaggagatgcagcatgggaagaagaacctaactctccccctttctctgtgtcccaggaagaagacggcatgtgaaatcactgatcaagaaaaagcagattaagtggctgtgtaccaagaaatgccaaaagtataatatatgggagcctcctgtc
atccccctctgataggtggatcgtcttgtgacccccctcaatgatcccccatttagccactcctaaccttactaaaatgatcctcctttgatctatgtgtcccaggctccctccccttccatccggttccctggatgcctgagaagtggagaggagggggctggatcagg
aagctttttaatcaggcatacaatactaacccaaactaagagactaaccagagtatcgttatgcagatgctatgcacagaagatctatttttggtttaaatgtttttattgttaatgtcataataattttataatgttagataaactactgcttatttgattgttatgtgtatttattga
ggtatatgttctaaatgccctcatgagttttaatggtatgtgtgtaaaccgatgtgatggttgtttaatctaaacaccggtatataaaaaaaaccgtaaataaataaacaaagagctattccctgctgagtaagatgcagcacagctgga

**C:** Sequence of cDNA (coding region in red and highlighted in yellow)

Tccggtgcattttgtccttgctcaggatgagagcagtggccgttctcttcctggtgagcgctgtcattgctctctctctctgtcttgaagagaaagattctcacaggcaggcacaggagttggcagggaagcactctgagttcttgagaagacctggggagcaaga
gattgatacattgggagctgggaagtgggaggtgagaaatgaattgaatcaccagaggagagaacgagccatcagatgctactgtcagaagacaccaaaacatcccagacgaagaagacggcatgtgaaatcactgatcaagaaaaagcagattaagtggc
tgtgtaccaagaaatgccaaaagtataatatatgggagcctcctgtcatccccctctgataggtggatcgtcttgtgacccccctcaatgatcccccatttagccactcctaaccttactaaaatgatcctcctttgatctatgtgtcccaggctccctccccttccatccg
gttccctggatgcctgagaagtggagaggagggggctggatcaggaagctttttaatcaggcatacaatactaacccaaactaagagactaaccagagtatcgttatgcagatgctatgcacagaagatctatttttggtttaaatgtttttattgttaatgtcataataa
ttttataatgttagataaactactgcttatttga

**D:** Amino acid sequence (signal peptide shaded)

MRAVAVLFLVSAVIALSLCLEEKDSHRQAQELAGKHSEFLRRPGEQEIDTLGAGKWEVRNELNHQRRERAIRCYCQKTPKHPRRRRRHVKSLIKKKQIK
WLCTKKCQKYNIWEPPVIPL

**Fig. S18.** Gene position and architecture (**A**), genomic DNA sequence (**B**), cDNA sequence (**C**), and encoded protein sequence (**D**) of the unannotated *cxcl17* gene in the NCBI reference genome of *Rhinatrema bivittatum* (two-lined caecilian). The information was downloaded from the NCBI gene database (https://www.ncbi.nlm.nih.gov/gene).

**A:** Gene position and architecture

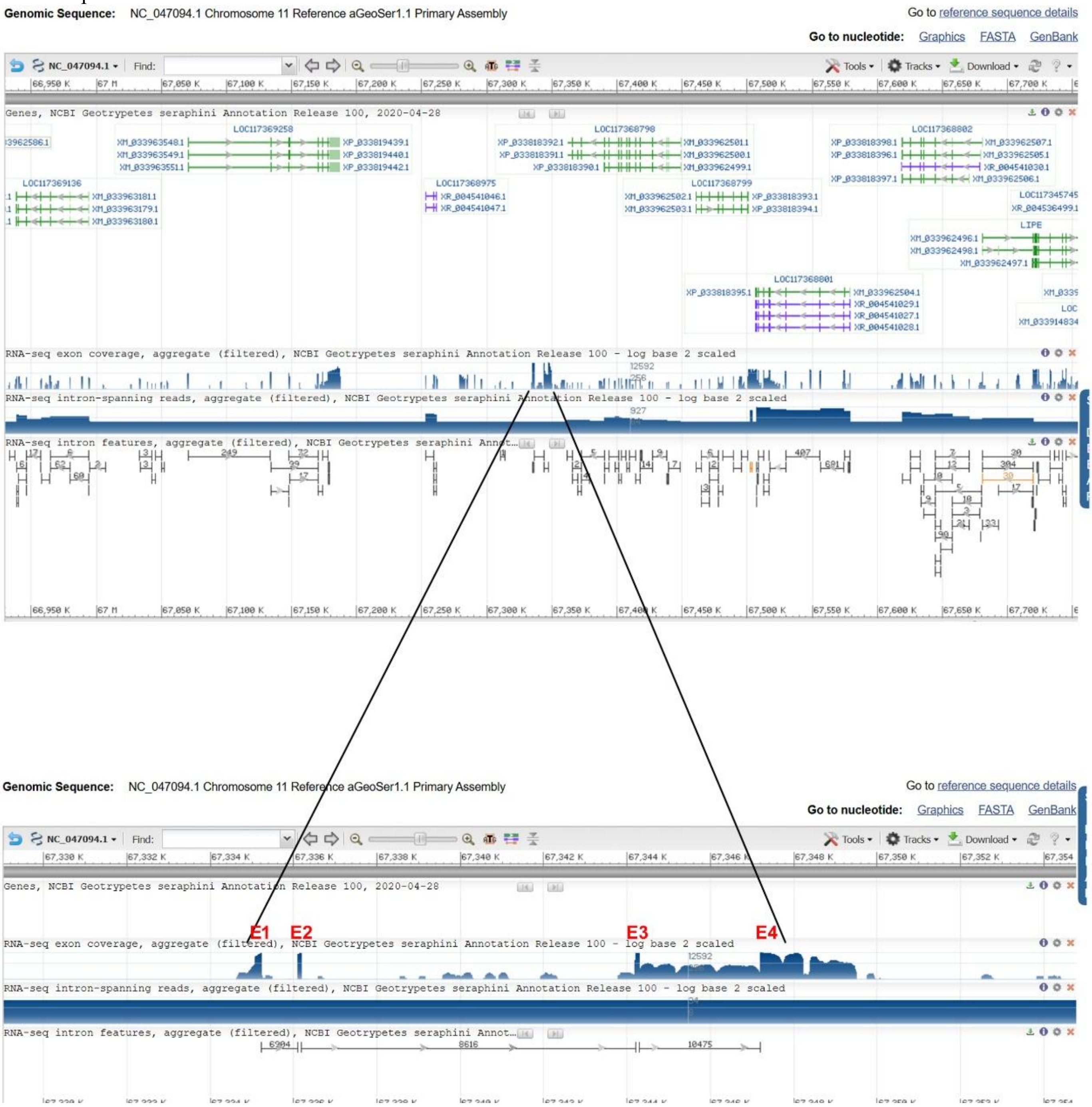


**B:** Sequence of genomic DNA (exons in red and highlighted in yellow, coding region underlined)

tcatgccattcaatgactttgcgtcttgtgtttatataagacctgcctcttcatagtcaggggccagtctgaagagacagacggacagactgcgggattgttactgctgtatttttgtcttgtagattttgattagtcacctttgcaattcaggttttgttctggttcagtatgag
ggttatccttgtcttagtgatttcggtcatcactgtctctctgtcatgaggagaaaggtaagagacactattcattggtactaactaaattatgagatcattcaacccttcttgtgctgtgagagtcccctttacctgaatggtacaagccacctctatacctgactgttg
agaattccccattcttgcactgtgtgagacttatgtatgtgagatataaaggcctacaaaatttgagttgcctaaccttgagctctgtgttttctcttatatgaggtatgagttctcctcgatactaagcaggaagtctctacatttgaattgtaaaaattccagtcataactga
attgagagatctttttatgataaatcatgtgttgtgttgtattattctaaattatattaggaaccatgacagatagaagtattaaatgtataatgattaatccattgggcatattgaagtgtcaagataaaaagatgaaagaaaaaattccacagataaaattcaggcaataag
aagcataactaaccagaaagaggtgagggaaacacaacagtttatttatggggcgacccaacacagttgtgtgttcacatggtctacatcagagatccagtatctattaagagtttttaaatgtaattttctactccctaatgtgacaaggtcagctagatttcaattgac
taatagtgccagcagcatatcctacaggccatttatgccataatgtggctgtgttgggtcaccccatgaatatactgctatggttctctcacctttctggttagttatgctttttgttttctagatgttgaaatattgccaatgattcttcttgaatacacattttcaagttctccaa
gggggggattctttgctgtcattcattaacagaaaaaatgttgcttctcacaacagattcccaaaggcagaagcatcggctagcaagaaggcctttaaaggtctcaagcacacatgggcaacttgaaattggtgagttcatgaatgaatagtcattctacttcttcttcc
tcttctggtgtccttgtcacacttgatggccttggtgactgcttcattaatgtcctcttttatataggcctccctatacttttctgcacttgctgtgctctggctaagagagtggtcccatgactatgtgaatctttctcattgtgaaagtgagagagccaatggcgtacctagc
atatttgatacttggggtgattattttttaacattcccctataaaaaaatattatttatagtaataattgacaagtcatacaacaaggatgcacctaggaaaaggcagcattttaaacgctgcagtgagcactagaacatcaatatatccatacccattgtaaaattaaata
agctagattagtacagatcaatcccgcataatcaatgctaacagaaaaccatgttttttcatacatatagaacacagatataccctcacccagtatggaataactaatcacaaactaaaattaagaaatatgtagacaaaagttaaattgaaccaccaagaagtcaga
ctctgcatacagggcaacaccacagaaacagaaatagtgatgtatgtcccctaatactatgcagaataataatgtggaaacgaagttcttggtgtaaaaattttatacaattcttcacaggtgaaaacaaggtaatgcaataaaaacagtacatagtacacacaacaa
agattgtggtttggacccaacatggtccgtgtttcacttttggtcaccttcttctagggtccaaatatcagatgccacaagattaagttatgcgaaaagcatccagtaatgtgtatatgtgaatgttgggtatcattaagaagggtatcacgaccaggacaaaggaagtc
atcctgccactgtatcgtgctatggtgcgcccgcacctggaatactgtgttcagttctggtcgccgtacctcaagaaggacatggaggtacttgagagggtccagagaagagcaactaagctaataaagggcatggaggacctctcatatactgacagactgaa
aaagctagggcttttctccctggaaaagcggagacttagaggagacatgatagaaaccttcaagatcatgaagggcatagaaaaaatagacagggacagatttttcaaattaaggggatcaataagtacaagggggcactcagagaaattgaaaggggagag
gtttagaacaaacgccaggaagttctttttcacacagagagtggtggatacatggaacgcgctaccagaggatgtgataaacaggagcacgctacaggggttcaaagaagctttggataggtacttggaagacaaagggattgaggggtacagataagagtag
agggatgggattagaggtaagttacaaaattaatcagggaccactgttcaggcactaggcctgatgggccgccgtgagagcggaccgctgagcaagatggacctctggtctgactcagcgggggcaacttcttatgttcttatgttcttataaaaatgccagaaa
ctgacttgtgctcactgcacataaatcctcttctggtattcgatatttggaccccagaagaaggtggccaaaaccgaaacatggaccgtgttgggtccaaaccacaatctttgttgtgtgtattatgtactgtttttattgcattgccttgttttcacatgtgaagaattgttgt
tattacatttttacatcaagaaatttgtttccacaatactgttctgttttgttgttggttttcccttgttgttctgtggaatttgttggagttcttgttctggtggaccttaatactatgcaaaatataaatataaagatagcagacacaaatttgaaaaaaaaatgacaaataatcac
tactttacaaattaacaaatagaaatacaataaaagattgaaaataagaaaataccattttattggactaatacatttttcagttagctatcagagaccaaaacctccttccttaggttagtacagtgtacagcttttatagtatcctgtcctgacacgaggaaggggatttg
ggtctccaaatgttagcaagaaaatgtattaaaattaatccaataaaaagattacctttttttcaattttctatttataaacatttaacaatacagctataatactactttattctaatgaaacaaaacattttttgaccttttgtcctctctgatttctgctttcctcatcttctactca

ctctcttctttccatccagtgtctgccttctctctcctttcatccagtatctaccctctttctctgctccttccatccacagtccaccctctttttctgcccttccatacattgtctaccctttttctgctccttacatcgacgtctgccttctttttatacccctttccataaactgtcta
ctctctccccctttccatccaacctgtgccacttctctccattatatatgatttatttcagcttcaccctctctccattttctctctcctctttgtctccacccttcccccatgctctggcatctctctcttctcctttccttccttctaccccacctatcccaccccatggtctagcatc
tctgtctccttctttttccccatgccctggcatctctccctctccctccatgatctggtatctctgtctccttcccttccctccccccatgccctggcatatctccttcttccttcatagtctggcatctactttattttctttcctttcatccctctctccattccttccttctgccctcc
tttctgcatggtctggtatctctctctcctctccctttcctggtcttccttttctctctccctctctcctccctcccacacccaaaatcaggtgcatcatttctcttctcccttcctccctccatcctcccaatgggtatctacttccctctctctctgttacccccctcttgaaaacttg
ccttttattattggcagcgcagccttcacaactcactactctggctgacttcaggccatcatctaccgcctatttcttcttctgtgaaggcaggacctagcagagaggaaggcccagcgtcggccagagctgcgagttgtgaaggctgccgctgctgctgcaagaa
ggggaaaaagttcccagacaatgggggccccctggagctgtagagaaccccccttatgatctgcatccgggagtggattgccctcccgcccccctcaccaccttggtatgccagtgaagggatcaatgactggagagcttgtgggcaatgaaacactattgaac
tcactaatgggtttatcacacagcaagaatacagaagtgatattgataaaatgcacagggaattagaggttggtggattcatgtgtgtgtatcagtcttcagagtataggaagaaaacacacaggcaacctgactgcagaatgtctgtacaggcttgtcgaactgtg
agagtgtaggagtgatagggaaagaatttttttaaaaaaaatctttattgattttccaaacttcccaatgtgcaaacaaacaaattgaagatacataacaatacaagaaaagcacattaaacttatagacaataatgaacaaatatttccccccctccccccaccccccac
caaataatcagaaaaaaggatgcctacaagaaactacccatatgttccctgaattaaacgccagtgcaaaccccctcccccctcccacccacccaccctggatgtgtatgctgaaaggttaacaaaataaaataaattatcattccttacaatacttagccaatggttcc
caaacataacagggaaagaattagaatgaagaatatgtatgtggcttggacatatggaataaaaacaaagaaatactgtggaaggagatgttggtgattcaggtatttatttggtcaacaaatgatgcaggcaacagttccaagcaaattacctttttttaaacccaga
tacaataaccactgttaccacatcctctggcaatgagttccagagtttaacttcgttttgtgaaaaaaagctttcctcctatttgttttagaagtagttccatataacttcattaagtatcccctggccattgtggtttttgaaagagtaaaaaaaatcaattcctggaagtacag
gaatgatgagaacagcatgaactgaaaataaaagaatgaggaagttatgcttggtttggtcatattataagcgtacaagagggataggagtgagttccccaaaagtaacctatagtataggtgaattcatgcatgggtgtaatatgaaattgctggagataaaatac
actagacttggggctcctttttacgaaggtacactagtggaaaatctatcacctgctcaaaagggagtggtagcggctagtgtgtgcggcaattcagcacgcgctattctgcgtgttaaggccctagcacgcctttgtaaaaggagcccttggagaaaatcactggtt
attcatggaagtcagtagcatggaatcttggtacttttggggattctgctagatatgacctggtttggtcactgttggaagcaagataccagtgggctatatcagtacttctcagccttctcctggaggcacatctagctgtcaagttgtcaggatggccacaatgaatat
gtatgagggttcttcaaaacatttctgcactttaatattttgacgggaaatgggcggggtgggagaagtggtaagaggacatgttaggacgctggaaaatgtatcgcaaaacagggtgattatgtgaagaagtgatgtaatttgcctttcagaaatttaatagatagc
gtttaaaaaaaattaaagtgtggaaactttttgaagatccctcgtatgagtcaaatttgcatattgtagagacctattacagtggtaccttggtttaagagcataattcgttccagaagcatgctcttaaaccaaaacactcatatatcaaagcaactttccccatagaagata
atggaaacttgaacaattagttccaccacccatctcccccccccgaggctaccggcgctgctccatcacccccctccccccgctctaaccggcatcgcacccccctgaaccggcatcgcaaacacccccccccccagcgaatgccccccgcgtgacccgcaa
agcaccccatgctgaaagtggcatccgcccgcccttcctcccaagcacggtccttacccccttctgctcgttgtaagggtgaatgtgagctcccgcctcttgcctgggctgggccttgagcatctgcgcatgctcaaggccttctggctctcgttctctcggagatct
cgttctctccgctagaacgagagccaaaaggccttgagcatgcgcagatgctcaaggcccagcccaagcaagaggcgggagctcacattcacccttacaatgagtagaaggggtaaggaccgtgcttgggaggaagggcgggcggatgccgctttcagc
acggggatgctttgcggttcttgcaggggggcattcgcgcgggggggggttgcgatgccggttcagggggggtgcgatgccggttagagcggagggggaggtgctcgtacaccagaacatgctcagtttgcgaggcaaaagtttgctgagtgttttgctcttctt
gcaaaacactcgcaaaccgggttactcgcaaaccgaggttccactgtatatgcaaatctatctcatgcacattcattgaggcgatcctgaaaatctgactggcaaggtgcctctaggagagggtagaaaatcactagactagatgaacctttggtctgatccattat
agtagttttatgtcttatcccaggggtaggcaattccggtcctcgagagccggacacaaatttagaccaggtcgacgctaagcatgattctatacagtgcgtctaacatttatagaatcgcgtttggccctgtactagtcggcgctgattttctaggcaccatgcatag
aatcagggcctaacttcttacctggattgataagcagaactgggaatagcagctcagagctctgttgggctcccatttggaaagataacataacataagaacataagaattgccgctgctgggtcaggccagtagtccatcgtgcccagcagtccgctcatgtggc
agcctacaggtcaaagaccagtgctctaaatgagtccagtctcacctgtgtacgttccagtttagcaggaactcgtccaactttgtcttgaatccctggaggttgtttcccctatgacagactccgtaagagcgttccagtttctaccactctctgggtgaagaagaact
tccttatgtttgtatggaatatgtcccctttcaactttagataatgctctctcgttctccctaccttggagagggtgaacaacctgtctttatctactaagtctattcccttcagtatttttaatgtttcgatcatgtcccctctcagtctcctcttttcaagggagaagaggcccag
tttctccaatctctcactgtatggcaactcctccagccccttaaccattttagttgctcttctctggaccctttcgagaagtaccatgtccttcttcatgtatggcgaccagtgctccaggtgagggcgcaccatggcccggtacagcggcatagtaaccttctccgatct
gttcgtgatccccttctttatcattcatagcattttgctcgctctttttgccaccgccgcacattgcgcggatggcttcatcgactagatcataactctcaagtctctttcctgggaggtctctccaagtaccacttctcccgccccgactatttattggtaagatcttcctcaa
cagaaaagagagagagtgtgggcagtggcgtagtgaagggtgagaggcacccctcccccaccctcatctctgccacccccatgctccttccccattccccccgccacacgcacccccttccattcccccgtacctctagttgaagttgttgctcacagcggtcaac
cacattctcctcgcgagcccgtcaactctccctctgatgtcactttcgatgcgcggcacccagaagtggcgtcagcgggacagctgatggggacgcgaggagcacaatgttgaacactgcaagcaacaacttcaactagaggtacagggaagggggcatac
gtgtggcgaggggaggagtggaaagaggcaggggtggagagaaggaggggtgctggtcagtggcgtatcaagggcggggtggggcggggtggggagggcagatcgccctggttgcatgccctaggggggtgcacagccggcccggtccggaacc
tcatgggacctgcacttcagcatggctgccggtccaccccgcggcaaagaaaaaggctaagaagctggcgggagcagcgaaagcgtcctttaaggaacatcctcccgatctggctctatcccgcctccttcgagtcgcttctggagaatgttctcacgaagg
gggcggcatagagccagatcaggagggtattccttaatggacgcttttgctgctcccgctggcttcttaggctttttttgaaccatagcagtggcataccaagggtgagagtgtgtgtcaaaaaaaatgatgggccaggggggtgtgtgtgtaaaaaatgatgggtccc
gggtatcacatacccctaggtacatcactggtgctggtgcacccattagcatggtgcccggggcagaccacccccttcagtgtcgtagtaagggggggggggcccttactacgccactgagtatgggagctgttgtaactcctgagatgatctgtcagcaattaggt
ctttttgctcttagaattatttattctagggtttgaggagaatctgttgggtgcagtggagggggggaagcgtgcccctttgaggagctgaagagtcgctgtgcccaatgaactttttaaattcctagctctgccatttctccctctttacagctctgcatcttctgtaacaac
cagtggcatgtgtcttttcagattcactgggagctgggaaccagatgagaaatgaattgagtcaccaaaagagacagagaggcatccactgcaactgtcagaagatgacaaaacagtctagaagtaagaagccatcctgatggagcaacccaaggggtgatc
ctaccactgtatagcaggggaaactctagtactaggggagagagcccctagccagtggtgtaatgacagagagggagggagggaaccatgaccccaggtcaccctatgcagaatctgaccttaagtccactgtttacacttttcaatggaagcttgtcatttttta
gtgccatcacctcttcaggagccaatagcatagcgaaggtgagtggcgcccaagatggggaccccccctcccctgccctattccctgattcccccccgccgcacacgcccctgtccaatacctttttaacttcagagcgagcagctaccaacttactgcctgcgtca
gcttgggcactctctctgacgtcacgtcctagttgcgggtcctgaaaatgacatcagagagagtgccgaagctggcgtgggcagcaagttggtggctgctcgccctgaagttaaaaatgtactggggaagggacgtgcgtacacacacacacacacacact
cttccggcagtgggagagatgtccaatttctcctgctgctaacagacacacctcaccagcaggaggggccttaaaaatctgggccaatcagcctcagacccctctccagatgcatctggggaggggcctaaggctctgattggcccagatgcataaggcccca
cccataagaggtcagtcagagcctcaggcatctccccggtgcatcccaagatgaactggggaggggaaggcctgccattttgaagaggtggatttgctggtgggagagaataggcatccttctggccagccatctgaaaataaggtaaaggggagagggtg
gggtcatcgtaggcatgggggaagggtgttgcatggttgtgggagagcatgggcaactctcatgctgccaagggggggttagaaaagagtgttggttggcggcaggagacatcgggcatctctcctgctgctggggagggtgttggttggcaggaggaaag
actgggcatctctcccgttgccaaaggtggctataatacacgtgctcaagaaacatgcttttactattctcaccaataataaaatagatgacaatttatgattcttcatgtaatttattttttttttcataagccacagtcaaaacacacaccacaagatgcacttttagcagtg
ctgacactataccagcatccccggatcagtcactgatttttgttgccagaagctgatgccactcttggagaatggctcggcaatttttaagaatccaccggagtgctcatttcaatactgaagagcccagttgcatggcagtgtcggagactgctagaaagctcatta
aagaccgagatacgccattttgataatccgccactaaaatgcatgcaggctaaactgtctggaaacagtttagtgactgcattaaagttttgagaatctgggtcaataggcgcatctagccgaaatagcttaatttttttaattggttcaatagaccatgttaattgaacag
cgctattaagaaccaatttaaaaaaaataataattttgggctaggcacctaacatggtaggtgccatgtagatgtcctatggtgtagctagggtaggaggctcccagggtggtggtgcccctgagccctcctctctgccccctgctccttccgtgtctccaccatcac
actcacaccctccattcccaccccctaaagttcttcaaatcttcatcagcacaagcaacttctctggcctgcttcttgtgctggctttccctctggtgtcacttcctggccccacgatccggaagtaatttcagagggagccaaattggcatgagctacaagctagagta
gttgctcatgctagtgaagattttaaagaggtaccagggaggggcagggaagggagggtgcaagcgtggcctgggggacggagaggtgctgacatccccaccaagactgtgcccagggtggtccacccccgccctcttaatatgccactgtaggtatct
acaccaaggcgcctacaagcatggattgtgttaggtactggtattttaggtcatgaaagcgtctgtaccgtctctgattcctattggcacctaatctgagttaggcaccgttaggcgcaattctataaatagcggctaattttgaatgacttgcagtaggtgccatttcctt
aggtgtctattgatttaggtgctgtttgtagaaacagggccttagctcttaacttctcttccccaccctacatccattcctggttactacctactttttatactcttaaatttagttgaattttcagtaaaaatgaatgcgctcattcctagcaaatattcatatttagagttcttttcc
ttactgacttatgtattgattgtaaaacgctttgatttttttcttgtgagcctaaagtaagcatatgcataaatttgttgagtccaatttttaagcataactctcagacttagcataaatcttatcagacccatggtatttactgcactgttttttttagcatacagtagctgttagcac
agctgtgctataatggttaacactcactaaaatctatgctatttagcacactttagtaaaaaggttccaaaggttcagctctatccataaatctaaacccagatctacacctaattatatacaccaatccaagccaaggtctgtccataagccccatattccaggcctaga
tctgtctatagatccctaaacctcactcccaggtctatctagattgtccaggtctatccttaaggtgaaatctaaaaatgaggctcctctgttagggtctggagagatgaagtactgccagggaagaagaatctaacccttttttttctgtgtgtctcaggatcaaggagg
catggaagagtagcgccgaagtacaggcatatgatgtgcatcaagagctgccaggcacatttgcccaacaatttgaactggaatggtcctataatccccctctgaaaggtcattctcatgattcccctctgaaaggtcgatctccctgtactaagaggattttcttga
aactctaggatccccttctgcttggtgaatttcccactgatcctgccagtggcgtagtgaagttgagaggcgcccaggatggtggcactcctcccccaccctcttctctgcccccacctccacacgctccttccctgcccctcctgccgcatgtgccctgcttccctt
cccccgtacctctgtaatgttcctggcatgagcagcaactcccaacctactgtcacgccaacaataactcttcctctgacatcacatcctaggtgagggtccaggaagatgaagaaccaacactggcgtgacaacagttttgggttgctgcttgtgccagtaatgtt
acaaaggtatgggggaagggaagtgttgcctgcgtggcagtgggggaggagggaggaaggagtagggggggggcagggaggtggatgggtgcctgcgccctcaccaagtcagcgcacggggaggaccgccctccctttactacgctactggatcctgc
tgtctactatgtcagttatgaatctatgtgcagtaca

### C: Sequence of cDNA (coding region in red and highlighted in yellow)

Cagtctgaagagacagacggacagactgcgggattgttactgctgtatttttgtcttgtagattttgattagtcacctttgcaattcaggttttgttctggttcagtatgagggttatccttgtcttagtgatttcggtcatcactgtctctctctgtcatgaggagaaagattc
ccaaaggcagaagcatcggctagcaagaaggcctttaaaggtctcaagcacacatgggcaacttgaaattgattcactgggagctgggaaccagatgagaaatgaattgagtcaccaaaagagacagagaggcatccactgcaactgtcagaagatgacaa
aacagtctagaagatcaaggaggcatggaagagtagcgccgaagtacaggcatatgatgtgcatcaagagctgccaggcacatttgcccaacaatttgaactggaatggtcctataatccccctctgaaaggtcattctcatgattcccctctgaaaggtcgatct
ccctgtactaagaggattttcttgaaactctaggatccccttctgcttggtgaatttcccactgatcctgccagtggcgtagtgaagttgagaggcgcccaggatggtggcactcctcccccaccctcttctctgcccccacctccacacgctccttccctgcccctcc
tgccgcatgtgccctgcttcccttcccccgtacctctgtaatgttcctggcatgagcagcaactcccaacctactgtcacgccaacaataactcttcctctgacatcacatcctaggtgagggtccaggaagatgaagaaccaacactggcgtgacaacagttttgg
gttgctgcttgtgccagtaatgttacaa

### D: Amino acid sequence (signal peptide shaded)

MRVILVLVISVITVSLCHEEKDSQRQKHRLARRPLKVSSTHGQLEIDSLGAGNQMRNELSHQKRQRGIHCNCQKMTKQSRRSRRHGRVAPKYRHMMCIK
SCQAHLPNNLNWNGPIIPL

**Fig. S19.** Gene position and architecture (**A**), genomic DNA sequence (**B**), cDNA sequence (**C**), and encoded protein sequence (**D**) of the unannotated *cxcl17* gene in the NCBI reference genome of *Geotrypetes seraphini* (Gabon caecilian). The information was downloaded from the NCBI gene database (https://www.ncbi.nlm.nih.gov/gene).

A

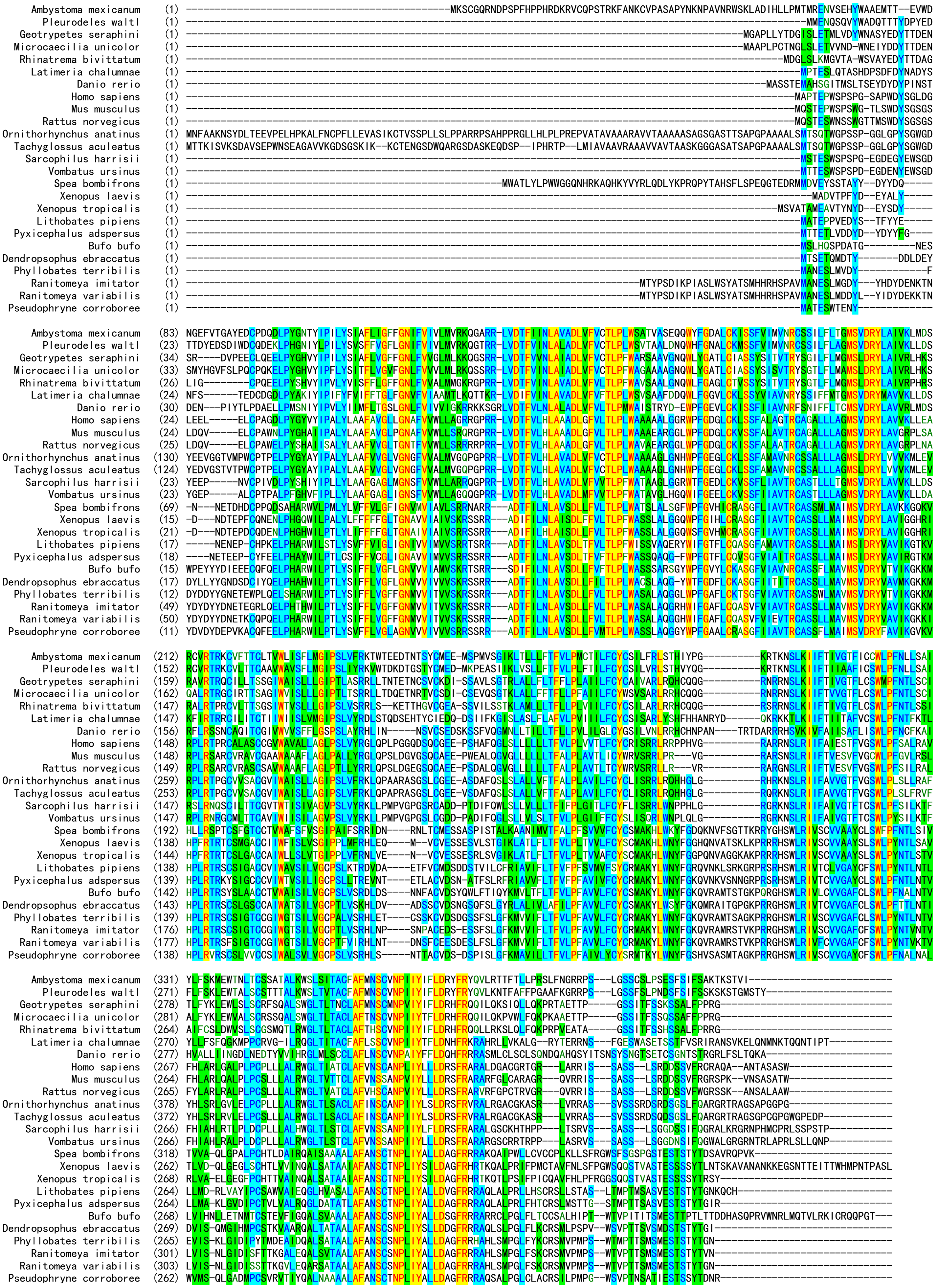
Ambystoma mexicanum
Pleurodeles waltl
Geotrypetes seraphini
Microcaecilia unicolor
Rhinatrema bivittatum
Latimeria chalumnae
Danio rerio
Homo sapiens
Mus musculus
Rattus norvegicus
Ornithorhynchus anatinus
Tachyglossus aculeatus
Sarcophilus harrisii
Vombatus ursinus
Spea bombifrons
Xenopus laevis
Xenopus tropicalis
Lithobates pipiens
Pyxicephalus adspersus
Bufo bufo
Dendropsophus ebraccatus
Phyllobates terribilis
Ranitomeya imitator
Ranitomeya variabilis
Pseudophryne corroboree

**B**

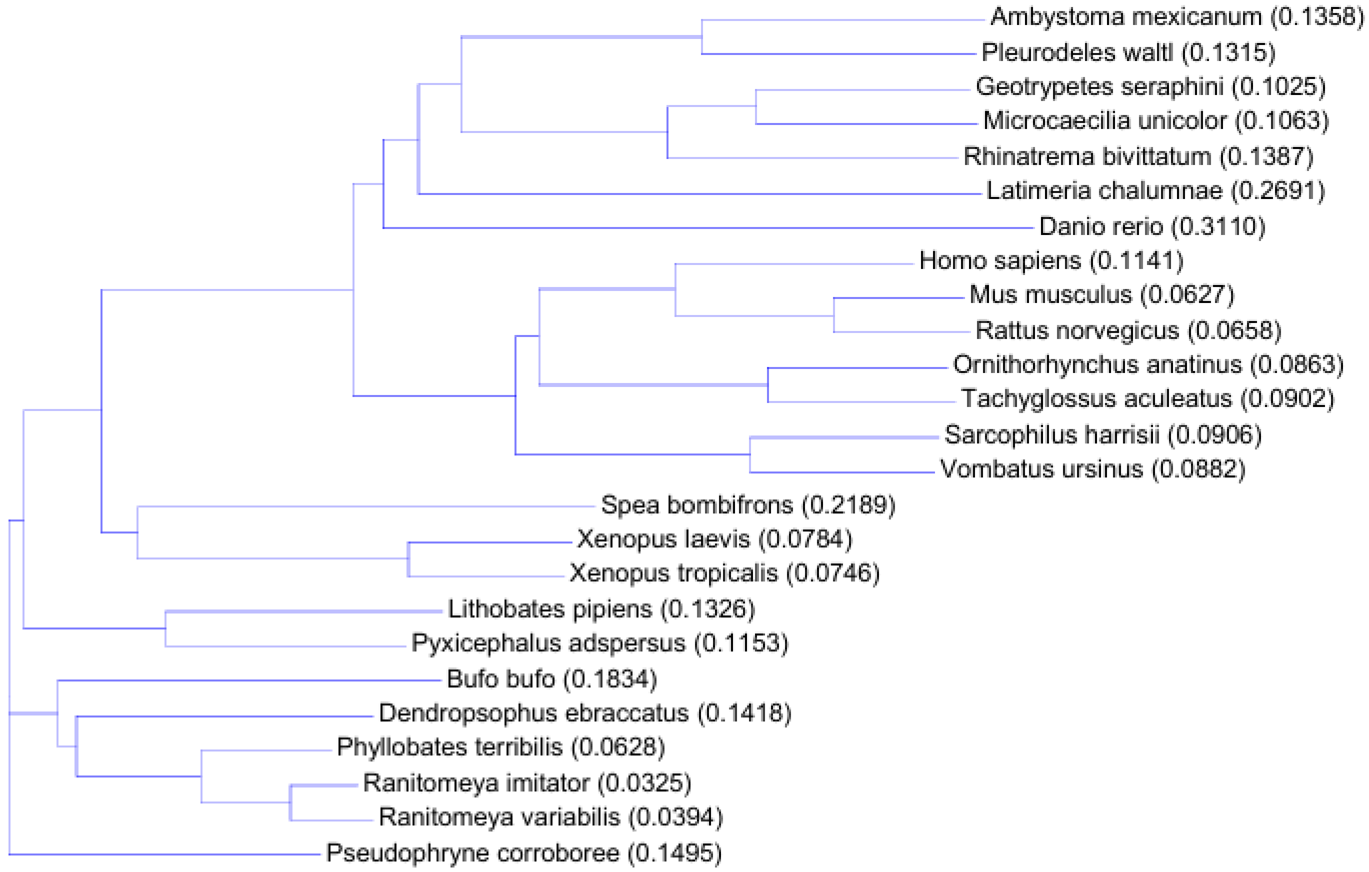


**Fig. S20.** GPR25 orthologs from mammals, fishes, and amphibians. **(A)** Amino acid sequence alignment. (**B**) Phylogenetic tree. Information about these GPR25 orthologs is listed in supplementary Table S2. These sequences were aligned via AlignX algorithm using the Vector NTI 11.5.1 software.
